# Polymer physics of nuclear organization and function

**DOI:** 10.1101/076661

**Authors:** A. Amitai, D. Holcman

## Abstract

We review here recent progress to link the nuclear organization to its function, based on elementary physical processes such as diffusion, polymer dynamics of DNA, chromatin and the search mechanism for a small target by double-stranded DNA (dsDNA) break. These physical models and their analysis make it possible to compute critical rates involved in cell reorganization timing, which depends on many parameters. In the framework of polymer models, various empirical observations are interpreted as anomalous diffusion of chromatin at various time scales. The reviewed theoretical approaches offer a framework for extracting features, biophysical parameters, predictions, and so on, based on a large variety of experimental data, such as chromosomal capture data, single particle trajectories, and more. Combining theoretical approaches with live cell microscopy data should unveil some of the still unexplained behavior of the nucleus in carrying out some of its key function involved in survival, DNA repair or gene activation.

## 1 Introduction

The eukaryotic cell nucleus separates the genetic material carried by the DNA molecules from the rest of the organelles as opposed to prokaryotes where there is no membrane separation [5]. The cell nucleus is certainly the ultimate key subcellular compartment to understand, because its organization is essential for a myriad of DNA based transactions such as transcription or DNA repair. Understanding its complexity is a joint effort of both theoretical and experimental approaches. We review here recent polymer models and their analysis, computational methods and numerical simulations, statistical and data analysis approaches for extracting information from large amounts of experimental data and predicting the main driving forces responsible for nuclear organization and function.

Physical models driven by diffusion at different length/time scales [119] and polymer models can be used to describe DNA and the chromatin molecules [226, 153], aggregation-dissociation with a finite number of particles [123] or the time for Brownian particle to find small targets [55]. The mathematical analysis of these models is used to estimate short- and long-time behavior of rare events, such as molecular binding, DNA looping, DNA repositing to the small location or DNA search for homology. The asymptotic analysis of the model equation reveals how key physical parameters control the spatial scales of the nucleus starting at a molecular level.

We recall that the nucleus contains the genetic material folded in multiscale but yet unknown organization (Fig. 1). Genetics assays that introduce mutations to the DNA can perturb its function and change the nucleus organization. While these local changes are often difficult to resolve spatially, they can however be studied using polymer models and analyzed by stochastic processes [89].

**Fig. 1:**
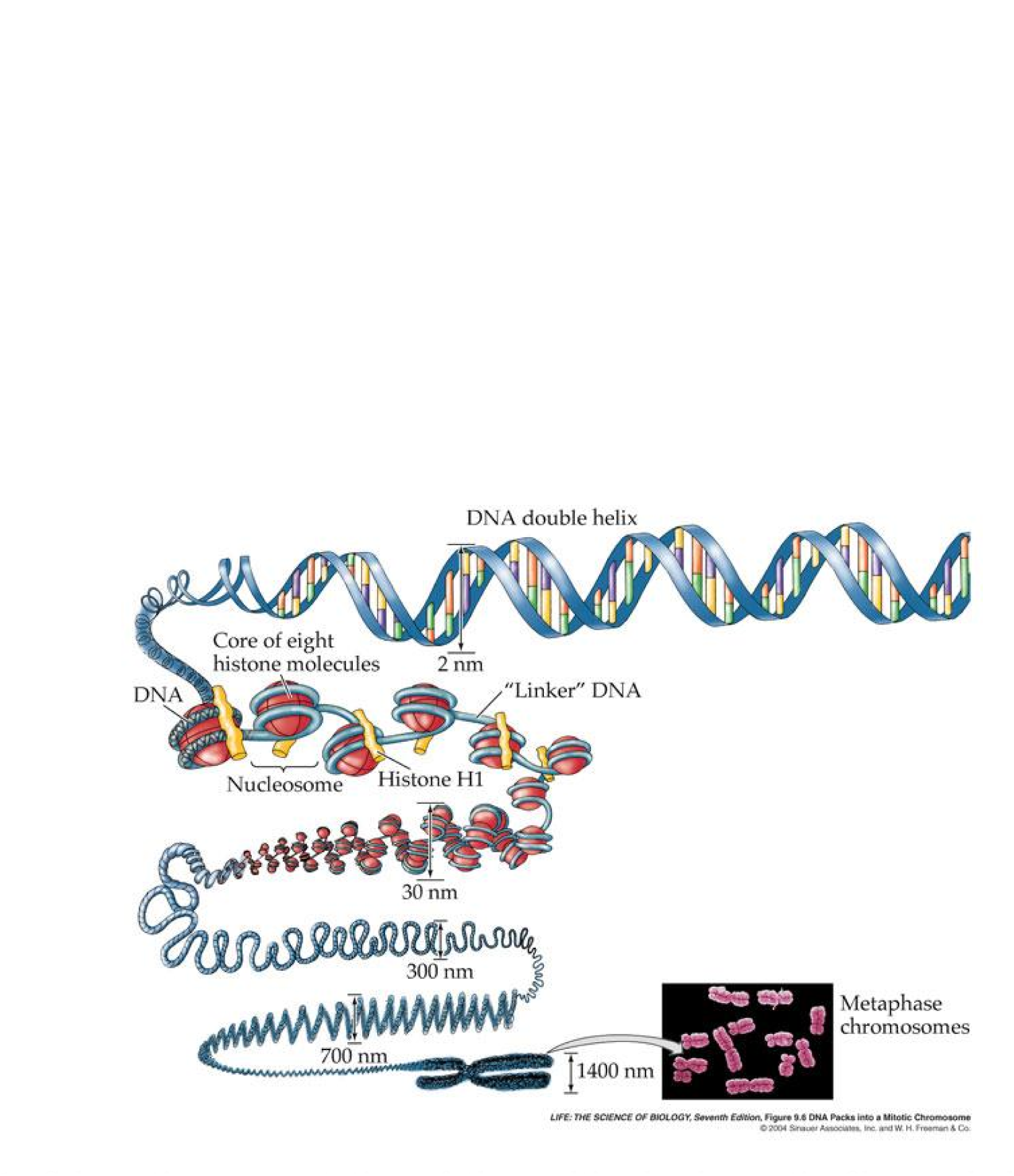
**Schematic representation of the multiscale chromatin fiber and chromosome structure (adopted from [210]).**

The multiscale organization starts at the molecular level (the four bases A,C,T,G). Eukaryotic DNA is wrapped around histone proteins, mediating mechanical and electrostatic interactions, to form a unit called a nucleosome (Fig. 1) [165]. The spatial scale above the nucleosome diameter is often referred to as *beads-on-a- string*, where nucleosomes form beads connected by linkers. Nucleosomes are further compacted by different proteins such as linker histones to form another structure called the *30 nanometer ber* [77]. The fiber is organized in a hierarchical structure whose nature is now being revealed, until the very coarse-grained level of a whole chromosome [80] (Fig. 1). Integrating all these interactions from molecular to the chromatin level is certainly a challenge for modeling.

While the 10 nm level scale is supported by the nucleosome on which the DNA is wrapped, the organization and folding at higher scales is still under debate. This multiscale representation has also been recently challenged because the nucleus is highly dynamic and chromosomes are constantly moving, opening or reorganizing [59, 199], an activity that can be directly monitored using live cell microscopy at an unprecedented spatio-temporal resolution of tens of nanometers and milliseconds [243, 155].

Live-cell imaging of the chromatin and polymer models are now converging. If chromatin locus can now be routinely tracked at a spatial scale of tens of nanometer resolution and at a time resolution of hundreds of milliseconds or less [40, 172, 25], in parallel, polymer models and their numerical simulations are constantly increasing their accuracy by incorporating physical and mechanical properties of the DNA or the chromatin structure to reproduce their local organization [173, 222, 187, 223, 272, 291, 274, 10].

Statistical quantities derived from Brownian motion theory [39, 69, 181], such as the Mean-Square-Displacement (MSD), computed along single particle trajectories are used to analyzed chromatin properties. The MSD is used as an indicator of spatial confinement [170, 111, 68], but it remains difficult to give an final interpretation, because the motion of a single DNA locus cannot be reduced to Brownian motion. The motion is indeed much more complex, because a locus is a bead moving with the entire polymer. Thus polymer models are needed to interpret the statistical analysis and bridge the gap between the output of statistical analysis obtained for a single locus and the reconstruction of the underlying chromatin organization. Additional polymer properties are clearly necessary to interpret the motion of a single DNA locus, especially when the entire DNA molecule is moving in a confined environment, or when a small promoter or DNA site is searching for a specific partner that can be relocated on the nuclear surface (Fig. 3). In most cases, statistical analysis has so far revealed deviation from diffusion called anomalous diffusion, for which the interpretation still remains unclear.

The effort of revealing the nucleus organization does not simply prevail for mammalian cells, the chromosomal architecture in bacteria is quite complicated and appear also to be modulated both spatially and temporally. High-resolution single particle trajectories have revealed rare but rapid ballistic trajectories. While the origin of this motion is still unclear, polymer models are used to examine whether subdiffusive chromosomal dynamics exhibits these rapid movements or they are due to external forces (see [129, 128] and [175] for a general review on physical interpretation of bacterial DNA based on sub-diffusion theory.)

Another difficulty inherent to the cell nucleus is the absence of clear main driving forces. In physiological conditions, ionic concentrations vary from tens to hundreds millimolar, and thus an average electrostatic force cannot act at a distance larger than few nanometers, as measured by the Debye length [58]. This electrostatic force has a fast decay due to a screening potential induced by the statistical properties of ions in solution. Consequently, two charged molecules cannot directly interact unless they are moved in close proximity, a process mostly driven by diffusion. Thus, electrostatic force should not have any role in long-range forces in the nucleus. Another source of chromatin reorganization and motion is due to actin fibers in the cytoplasm outside the nucleus, that is physically connected to the nucleus membrane and can affect directly the nucleus motion.

The genetic material organization seems to be dominated by random forces at various scales, but it is unclear how fast and how far these local forces can affect the long-range chromatin organization. In addition, it is also unclear how the local chromatin loci are affected by diffusion. Local diffusion is generated by random collisions, but the overall random motion also account for transient interactions with surrounding molecules and thus does not simply refer to classical Brownian motion of a particle. Random motion of a locus probably includes random forces that originate from the thermal noise in a crowded and organized structures such as the chromatin fiber. This subject is not new in stochastic processes, where models are used to study correlated noise and how random collisions in uence large molecular structures [236, 238, 237]. Complex noise can also be characterized statistically using the power density spectrum [237] for single particle trajectories, but here again the interpretation of the data should be based on the statistical properties of polymer models. In summary, chromatin is constantly moving driven by various cellular sources leading to random fluctuations inside confined nuclear sub-domains and restricted by the nuclear envelope in eukaryotes. Accounting for these forces is a clear challenge for polymer physics and for the interpretation of data.

In this review, we present several polymer models to account for the physical forces underlying nuclear organization. We shall first summarize the classical polymer models (section 3) followed by an analysis of polymer encounter events that are responsible for generating the large spectrum of time scales involved in nucleus organization (section 4). Indeed, the intrinsic polymer structure is certainly involved in the genesis of long-time correlation between the different spatial scales. Thus, the description of a monomer or a polymer dynamics does not fall into the classical random walk or the idealized Brownian mathematical framework. In addition are events arising from stochastic polymer processes such as random looping, search for a small target or long-range reorganization is probably the appropriate framework to analyze chromatin organization. A classical example is polymer looping [202, 209] (Fig. 14), where two sites are brought together for possible interaction. Analysis and simulations permit computing the rate of looping, which is the reciprocal of the mean first encounter time (MFET) between two monomers. Such rate provides great constraints on polymer dynamics and predicts the frequencies of encounter between two given loci on the chromatin. The analytical expression of these rates constitutes the emerging polymer physical laws for DNA and chromatin, that clarifies the dependency of many small and large parameters, especially when it becomes necessary to interpret the motion of a single tagged locus located on a (non labeled or hidden) polymer and to extract biophysical parameters (section 6). Polymer models can be used to analyze anomalous diffusion of a single locus, characterized by a parameter *α*, which is the exponent of the time-correlated function for small time, approximated by ≈ *t*^*α*^ [104, 34, 275, 284]. As we shall see, this anomalous behavior is generated by the collective motion of the DNA, again captured by polymer models and reviewed in details in section 3.

The chromatin is constantly remodeled, during cell cycle or under different stress conditions [255], preventing the nucleus from reaching a steady-state equilibrium. Telomeres which constitute the chromosome ends (fig. 5) can associate and dissociate in small cluster dynamically. Chromosome sites do encounter with different frequency depending on their locations, while gene regulation and DNA repair are possible based on constant re-organization of the chromatin [67, 265, 159]. These encounter events can be characterized by their frequency distribution that we review in section 5. Interacting sites coming into proximity is a random and rare event, with a long time scale compared to single particle diffusion [119]. The exact implications of these encounter events are still unknown, but over the past decades, many experimental methods have been developed. One critical and very prolific was the chromosome capture method [61], from which the encounter probability of DNA sites is extracted and the contact frequency can be estimated [83], as discussed in sections 5 and 6.

A down point of the theoretical approach is the actual performance of numerical algorithms to simulate chromatin dynamics at an atomic level. These performances are still insufficient to reconcile them with the need of following the nucleus reorganization over minutes to hours, as it would involve simulating too many components. Simulating the encounter between DNA-sites located far away is still hard because they are rare events, requiring long simulations and asymptotic formula (see section 4). But analytical and coarse-graining modeling methods are valuable tools especially to estimate fundamental time scales involved in encounter processes of macromolecules or during DNA repair (section 9). They are also used to extract biophysical properties of single particle trajectories of chromatin locus using live cell microscopy approaches [265, 155] (section 8). Interpreting data produced by these dynamical methods continue to benefit from polymer models, but reconstructing the chromosomal organization from these data remains a challenging hurdle.

In summary, this review aims to describe polymer physical models, numerical simulations and data analysis in order to understand chromatin organization in the nucleus of mammalian, yeast cells or even inside bacteria. It is organized as followed. Part 1 (section 2) is a very short description of the cell nucleus organization, summarizing basic properties of physical modeling. Current methods employed to explore the nucleus include live cell imaging, single particle trajectories approaches and more recently chromosome conformation capture methods, producing million by million matrices of encounters between DNA pairs. The second part (section 3 and 4) describes polymer models, starting with Rouse and other associated polymer models used to described chromatin. Asymptotic methods are used to derive formulas for the encounter time of two monomers during DNA-looping in a free and a confinement domain. These formulas reveal the structure of the parameter space and the role of singular parameters, which are usually the bottle necks of long numerical simulations and probably the underlying biophysical processes they represent. These analytical methods are based on eigenvalues expansion of the Fokker-Planck equation in a high dimensional space (the dimension of which is the number of beads of the polymer). The associated numerical simulations are based on stochastic dynamics, also used to reconstruct the chromosomal organization in the nucleus and to interpret chromosome capture data (section 5). In section 6, we discuss some modeling approach of the chromatin as a heterogeneous strand, aimed at interpreting the formation of topological domains in chromosomes and the relation between the epigenetic state of the chromatin and its structure. We describe in chapter 7 the encounter dynamics and organization of yeast telomere. In section 8, we summarize various approaches where polymer models are combined with live cell imaging data to unveil physical properties, such as normal and anomalous diffusion of a single chromatin locus trajectories, depending on the time scale of experiments. We mention several criteria to identify the appropriate time scale and nature of the underlying motion. Section 9 discusses applications of polymer models to various scenarios underlying major events in the nucleus, such as double-stranded DNA break repair, which involves a still unclear search process for a homologous sequence. In the final part 10, we summarize the role of polymer physics in interpreting the chromatin organization and discuss open questions.

## 2 A brief description of the nuclear organization

The nucleus of eukaryotic cells contains chromosomes and many other components such as nuclear pores and the nucleolus, a subcompartment composed of proteins and Ribosomal RNAs (rRNAs) [52] (Fig. 2, references are www.lifesci.sussex.ac.uk/home/Julian_Thorpe/tem29.jpg see also vle.du.ac.in=mod=book=print.php?id=9345#ch13971 and www.pinterest.com=pin=83316661827063673/) and Fig. 4). These components are implicated in proteins production, mRNA regulation and cell homeostasis. Yet, how these elements are organized in the nucleus remains poorly understood. The nucleus is not segregated in expressed and non-expressed genes, but the organization is certainly involved in the regulation of gene expression and repair mechanisms.

**Fig. 2:**
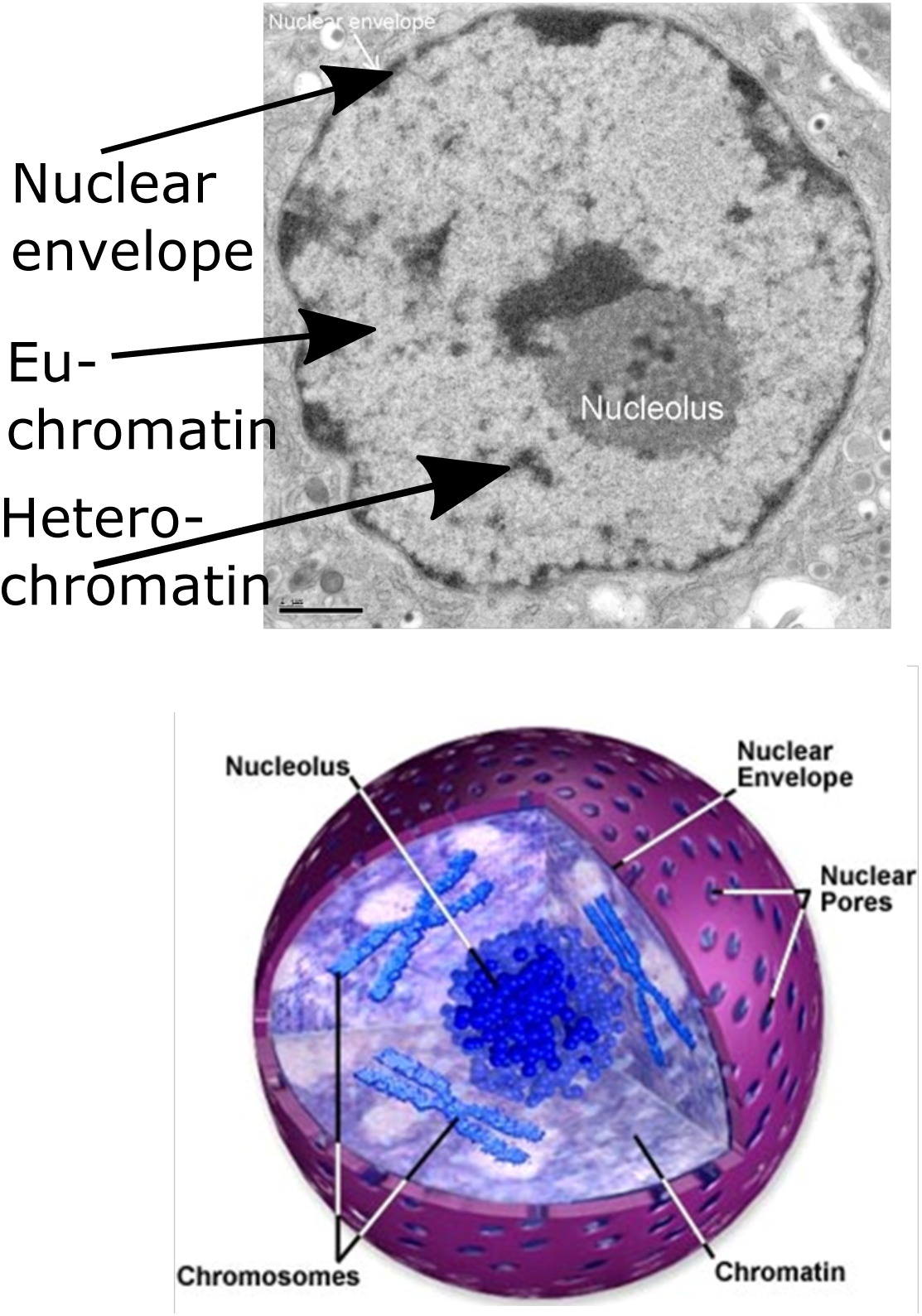
Organization in the cell nucleus. Upper: Electron-Microscopy image of the nucleus. We have indicated the different region: heterochromatin(dense black regions) and euchromatin (white region) and the nuclear boundary **Lower:** artistic representation of decondensed chromosomes (blue) floating in the nucleus. Nuclear pores are located on the surface.

**Fig. 3:**
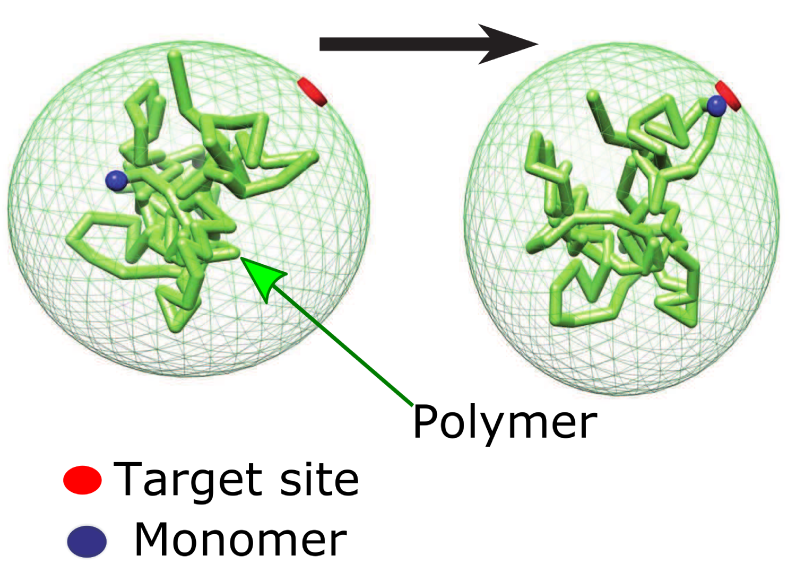
Schematic representation of a polymer. A monomer (blue) encounters after some time a small target (red) located on the nuclear boundary (figure from the group).

**Fig. 4:**
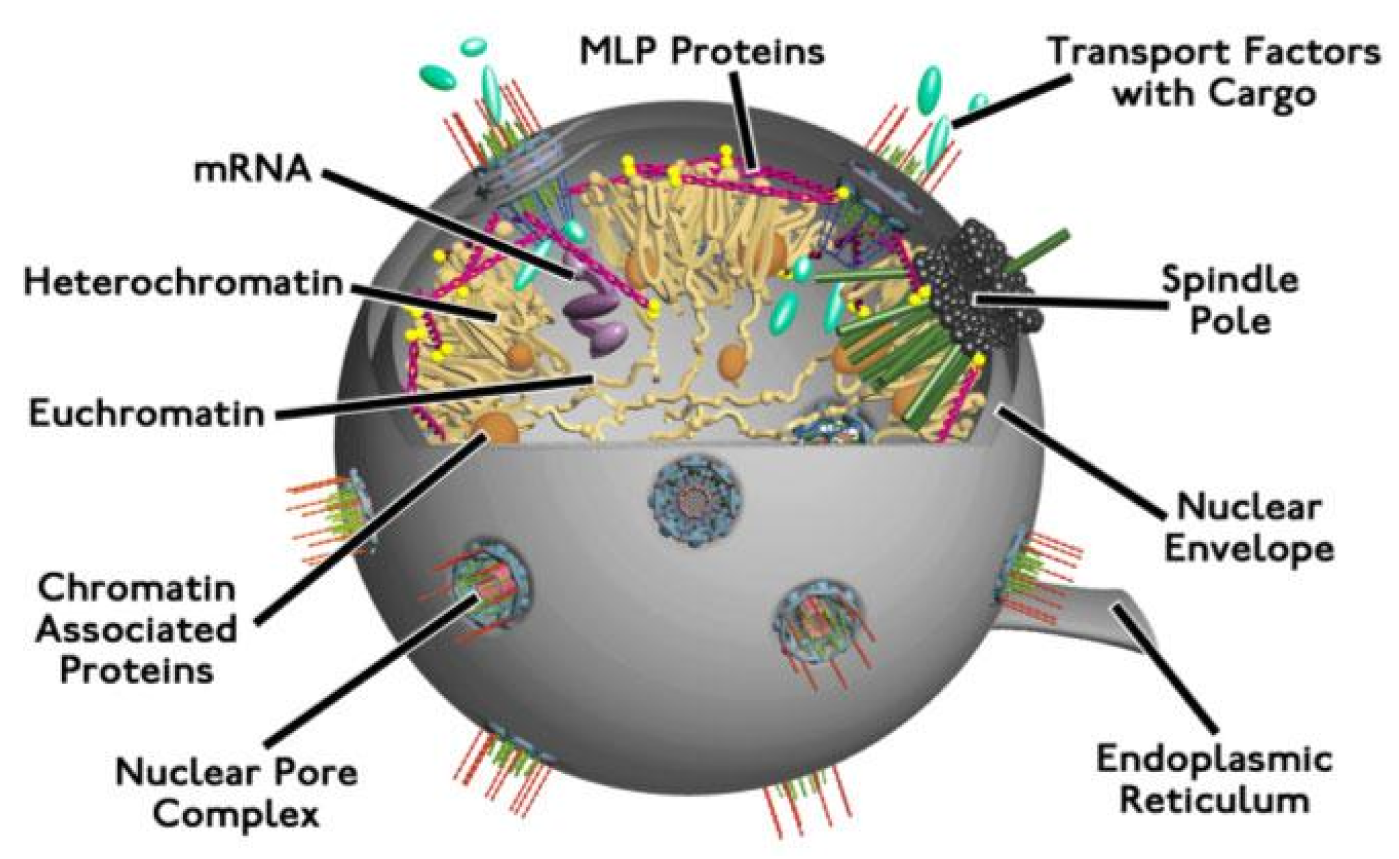
**Schematic representation of the yeast nucleus with various components (Rocke-feller University (http://lab.rockefeller.edu/rout/resproj3).**

### 2.1 Eukaryotic chromatin

The DNA is usually covered with an ensemble of molecules, forming the chromatin, which has an intrinsic structure, imposed by cylindrical protein complexes called nucleosomes [209, 165, 77] (see also [53] for a recent review on the physics of epigenetics). Beyond a 30nm scale, the chromosome structure is thought to be non-uniformly distributed in the nucleus. In yeast, the chromosomes fold back, called the Rabl configuration [131], resulting from the attachment of microtubules to a structure (the kinetochores) assembled at centromeric chromatin [213]. Microtubules are connected to a structure called the spindle pole body [265] (Fig. 4). In Metazoan, the nucleus is more dense (Fig. 6) and chromosomes adopt a globule shape. Interestingly, during cell division chromosomes are pulled by microtubules through the kinetochores into daughter cells.

### 2.2 Organization of telomeres

The ends of the chromosomes are made of repetitive sequences, the size of which changes after each cell division [27, 87]. In yeast, telomeres are located on the nuclear periphery and form stable but transient clusters through the interactions of proteins (Sir4 and yKu70/yKu80 proteins) [264, 265, 225, 124] with other proteins of the nuclear envelope such as Mps3 (Fig. 5). Telomeres are not uniformly distributed at the nuclear periphery, but form small dynamical foci [94] and their number and composition evolve over time [225, 124]. These clusters can promote gene silencing [225, 124], depending on (the Sir) proteins and on the spatial position of telomeres. Telomeres tend to cluster according to their chromosomal length, such that telomeres belonging to longer (or shorter) chromosomes cluster together [269]. Telomeres of mammalian cells are distributed more or less evenly over the nucleus. They can form clusters at the periphery [186], but also internally on nuclear bodies.

**Fig. 5:**
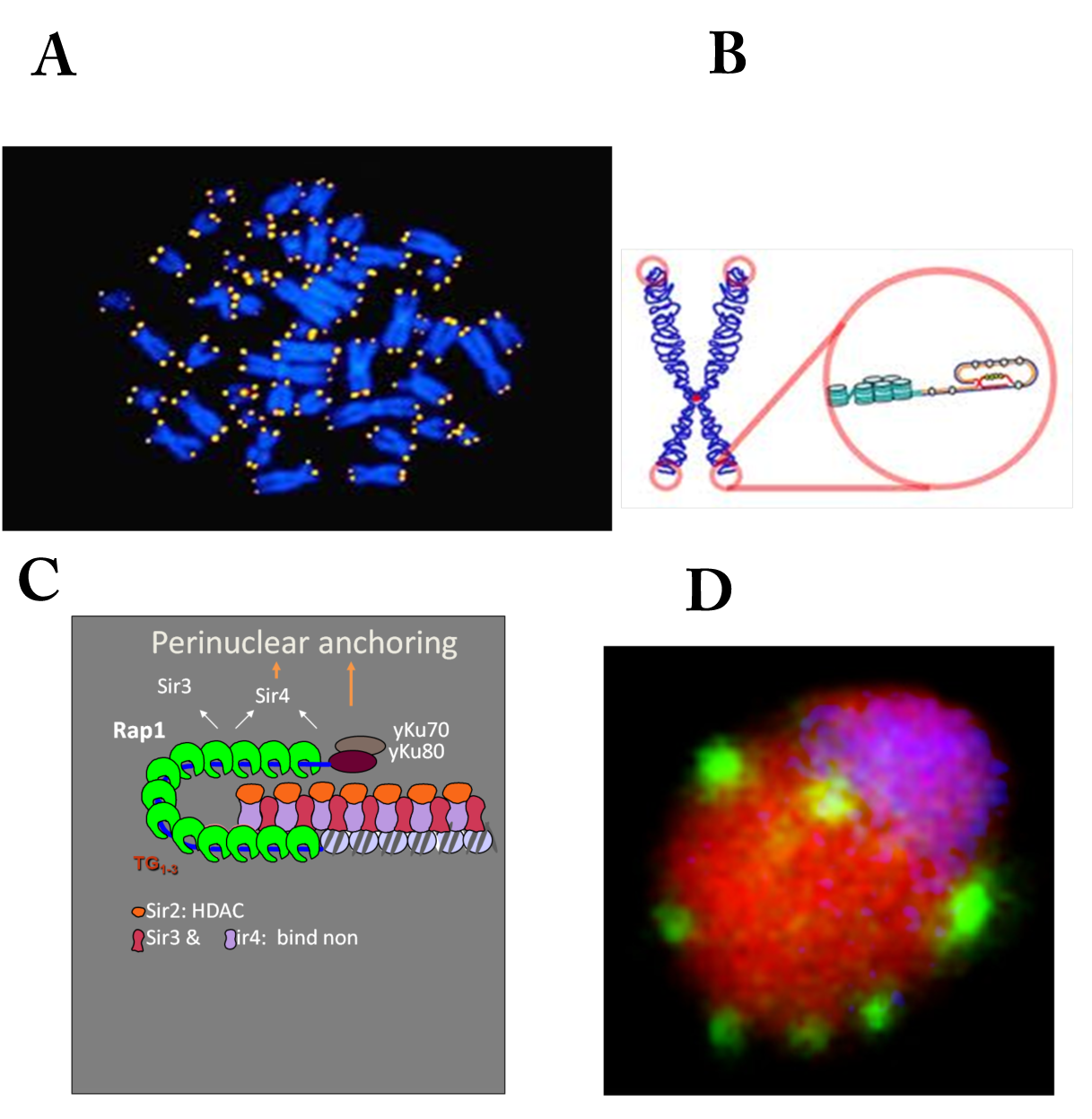
Telomeres (located at the end of the chromosomes) in the cell nucleus. **(A)** Telomere are marks in yellow. **(B)** Magnification of the telomere geometry.**(C)** local molecular organization of telomeres **(D)** Telomeres organized in clusters at the surface of the yeast nucleus [265].

### 2.3 DNA organization in bacteria

In bacteria, DNA, RNA and proteins are present in the same single cell compartment and the chromosome is a single DNA circular molecule further organized with complement proteins (the nucleoids). Despite the absence of intracellular membranes, bacteria have a high degree of intracellular spatial organization (see [175] for a recent physical and biological review). In the two bacteria E. coli and Caulobacter, fluorescent tagging of chromosomal loci reveals linear spatial arrangement of loci according to their chromosomal coordinate, the presence of chromosomal fiber and a highly non-uniform chromosomal structure, suggesting a compartmentalized structure with boundaries. Four macrodomains of a few hundred kbs in size have been identified, corresponding to regions surrounding the replication origin. 200-400 chromosomal domains have been identified in E. coli, characterized by macromolecular crowding, local electrostatic forces, supercoiling and nucleoid proteins are contributing to DNA compaction and organization [175].

### 2.4 Chromosomal territories

Live cell microscopy data suggest that chromosomes are organized in distinct geographical territories with heterogeneous shapes and sizes [54] (Fig. 6). This organization prevents chromosomes to mix. This organization is not prominent in yeast, probably because the nucleus is not dense enough with chromatin, however it becomes prevalent in higher eukaryotes, where the nucleus is larger and contains a higher density of DNA, generating a crowded environment.

**Fig. 6:**
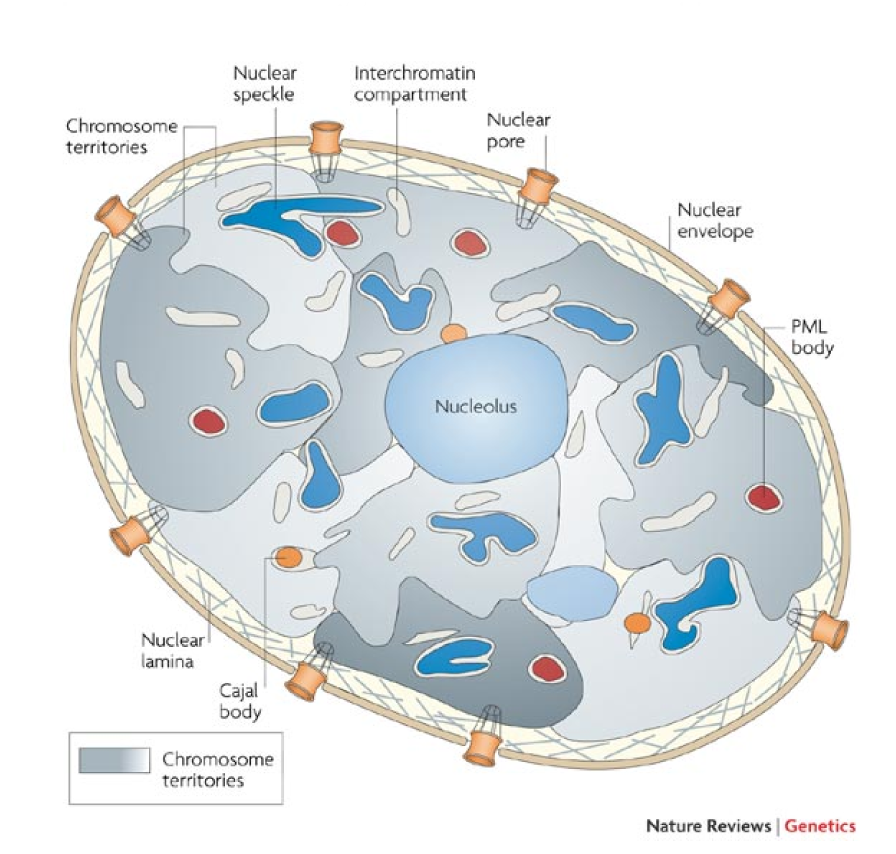
Representation of the cell mammalian nucleus organization. The nuclear envelope contains small pores and intermediate filaments. Nuclear sub-organized regions cluster to form nucleoli, near chromosome territory (gray) and interchromatin compartments. The chromatin is organized in distinct territories (reproduced from [150]).

Interestingly, changing chromosomes organization by clustering them together or part of them or several parts of different chromosomes can activate or repress genes. This reorganization could allow the simultaneous regulation of multiple loci in specific substructures, called 'transcription factories', although the existence of these factories is still debated [259, 51]. Gene expression can be modulated by regulatory elements, such as enhancers [209], which are often located far away from their target genes along the DNA strand. Many enhancer proteins are thought to associate with the promoter sites on the DNA molecule that they regulate, but a chromatin loop can also bring a molecule close to its promoter site, as observed using fluorescent markers [166].

Gene position affects transcriptional activity, but there is no general rule across eukaryotes: in yeast, genes relocate to the nuclear periphery after activation [266]. In mammalian cells, they relocate from the periphery towards the interior of the nucleus once they have been switched on [176]. Similarly, a number of highly transcribed gene-dense regions for example the MHC class II locus are expelled from chromosome territories, once they are activated after the formation of a large chromatin loop [281].

In summary, the organization of the genome is characterized by interactions of chromatin with nuclear sites, influencing gene regulation depending on the location. However, many genes do not change their expression level based on their position and active genes can be found both at the periphery or deep inside a chromosome territory.

### 2.5 Spontaneous DNA loops

When a DNA molecule bends enough to bring into close proximity two sites located faraway, a loop is formed [209]. Surprisingly, genes can be regulated by random looping events, when a promoter site interacts with an operator sequence. The classical example is the Lac operon system [148], where the lac repressor functions through clamping of two out of three target DNA sites (operators). Between the two sites, the formed loop can interfere with RNA polymerase while reading a gene sequence [174]. This process has been investigated experimentally [66, 76], numerically [47] and analytically [288, 261, 202, 13, 9] revealing the rate at which looping occurs. This rate is the limiting step for many other biological activities. A large part of this review will be dedicated to these looping events.

### 2.6 Motion of a single DNA locus

The motion of a single DNA locus is monitored using live cell imaging microscopy in bacteria or in eukaryotes and is described as random [265] or directed [111]. However, a unified physical description is still lacking. Interestingly, glucose depletion or the addition of protonophore to deplete the membrane potential, abolishes large range movements [170]. These effects suggest that interphase chromatin movement is sensitive to various ATP energy levels. Chromatin remodeler molecules rather than RNA or DNA polymerases [265] underlie the ATP-driven motions. Moreover, chromatin movement can be modulated depending on the cell phase. For example, it decreases as cells enter in the S-phase, which may be related to the binding of molecules such as cohesin to the chromatin [111, 70].

### 2.7 Chromosome conformation capture data

Chromatin organization in the nucleus was analyzed by estimating the frequency by which two pairs (small DNA fragments) come into close proximity (Fig. 7) [61]. In these experiments, the spatial chromosome position and interactions are detected by literally freezing the nucleus by chemical ligation of near-by sites [61, 60]. By averaging over a large cell population, a map of encounter frequencies summarizes the detected interactions in a large two-dimensional matrix (Fig. 8). There are several refinements, depending on the number of site and resolution. The interaction probability for the interactions between two sites is called 3C, between one site and multiple others (4C) or many sites interact together (5C) and Hi-C, when a low resolution map of all sites is obtained [73, 157, 75]. It is a challenge to reconstruct local physical interactions and the underlying chromatin organization from these averaged maps. Analysis of chromosome capture (CC) data revealed that chromosome distribution in yeast is not uniform. Some chromosomes almost never interact, whereas others show favorable interactions [221]. Hi-C data rely on ligation assays that capture only an instant of the chromatin conformation, which can be a rare event in the large ensemble of chromosomal configuration space. The probabilistic nature of the Hi-C maps reflects both inherent chromatin fluctuation and technical aspects of the methods (cross-linking, digestion and ligation reactions). For that reason, chromatin reconstruction relies on population averaging, which certainly destroys some local organization [200]. However, new techniques of performing single cell CC [189], allow studying the cell-to-cell variability. Features within the chromosome conformation ensemble have recently been found, characterized by increased interactions. These regions are called topologically associating domains (TAD) [197, 91]. As we shall see in this review, modeling ensembles of chromosome conformations has relied on numerical simulations of polymer models [212, 246], with constraints imposed by proximity ligation data [157, 22]. Polymer models are also used to explore the configuration space and DNA unpacking inside the nucleus [200, 89, 272].

**Fig. 7:**
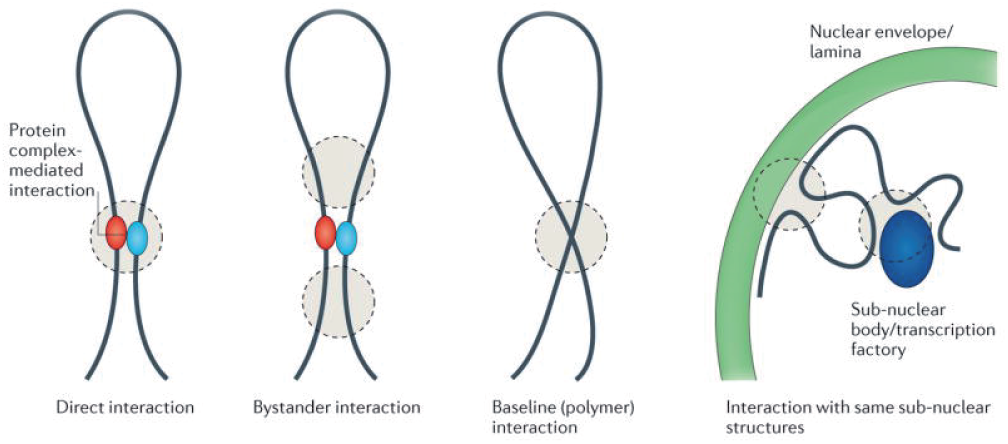
Schematic representation of the 3C-based technologies capture loci, located in close proximity. There are several possibilities for loci to be in close spatial proximity (proteins mediating interaction, Bystander interaction, polymer interaction). All these scenarios are accounted for in the 3C data (reproduced from [60]).

**Fig. 8:**
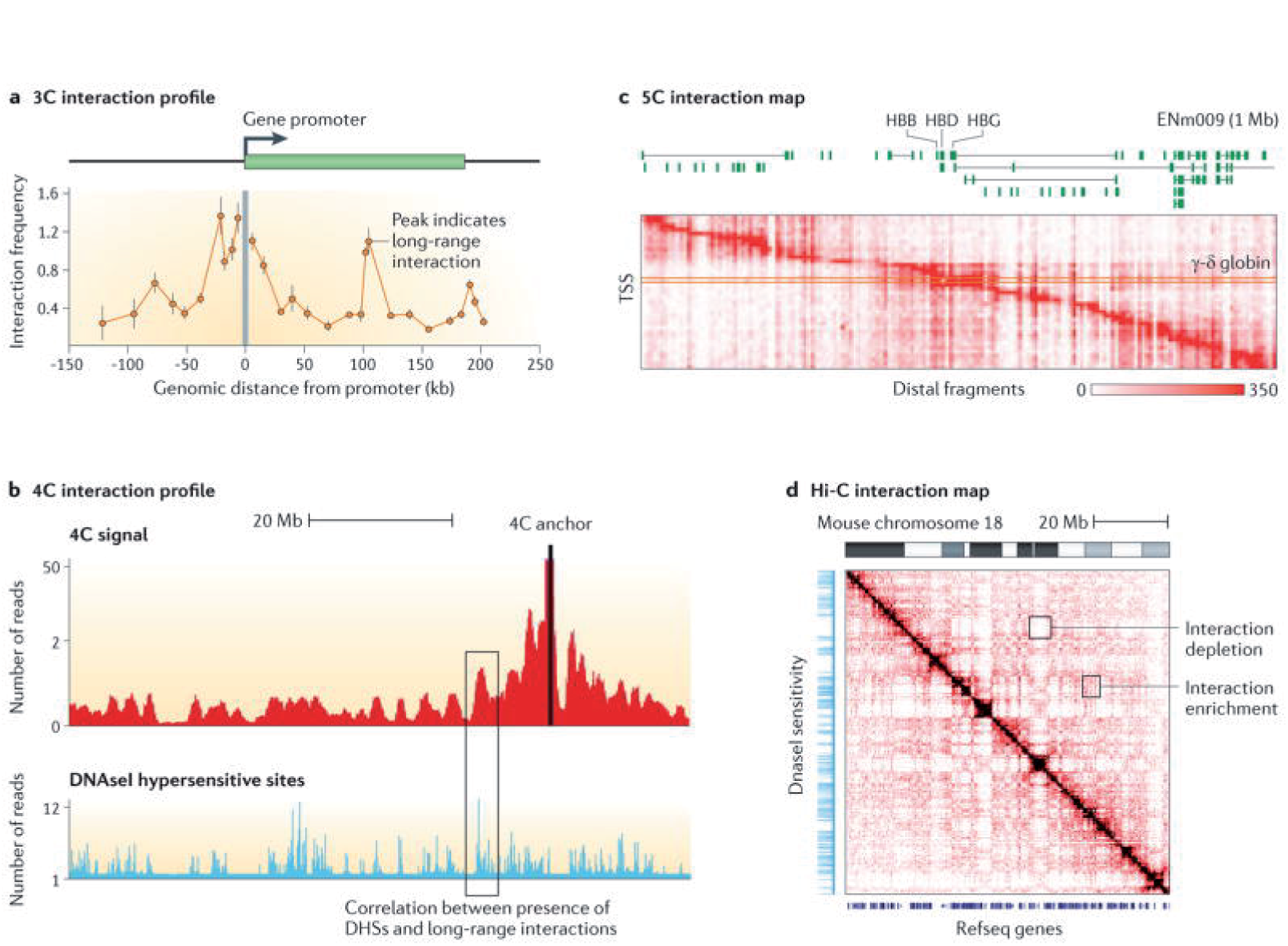
3C, 4C, 5C and Hi-C datasets. A. 3C data B. 4C data. C. An example of a 5C interaction map for the ENm009 region in K562 cells 46. Each row represents an interaction profile of a transcription start site (TSS) across the 1 Mb region on human chromosome 11 that contains the beta-globin locus. D. Hi-C. 3C and 4C data are linear profiles along chromosomes and can be directly compared to other genomic tracks such as DNAseI sensitivity. 5C and Hi-C data are often represented as two-dimensional heat-maps Other genomic features, such as positions of genes or the location of DNAseI hypersensitive sites, can be displayed along the axes for visual analysis of chromosome structural features (reproduced from [60]).

**Fig. 9:**
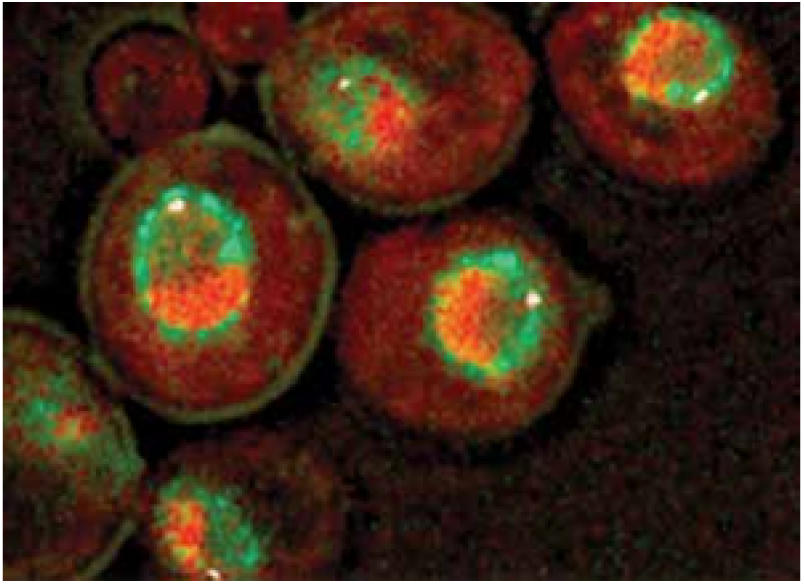
Nuclear organization of the yeast. Visualization of different elements of the yeast nucleus: the nucleolus (red), the yeast spindle pole body (white) and the nuclear envelope (green ring) and a GFP-tagged telomere (brighter green spot). The cells shown here are at different stages of the cell cycle and illustrate the organization within the nucleus (reproduced from [94]).

### 2.8 Spatial nuclear organization for gene regulation, DNA maintenance and repair

Spatial nuclear organization is certainly crucial to maintain genome integrity and it plays a key role in double-stranded DNA breaks (DSBs) repair. There are two modes of DSB repair: one is called Non-Homologous-End-Joining, where a break is repaired by a direct relegation [93], the second one is homologous recombination (HR) [160], whereby an unbroken template is used to correctly repair the break (further described in section 9). Interestingly, regions of repair are called foci, that can be visualized using a fluorescent marker - tagging a protein Rad52 involved in DNA repair by facilitating the DNA-strand invasion (see [188, 41] for the peripheral repositioning of the break).

When an artificial break, induced genetically cannot be repaired, the break persists and relocates to the nuclear periphery [188, 201] where it can be anchored. Irreparable breaks that have been processed by the HR pathway associate stably with nuclear pores. At a nuclear pore, a break may be repaired. Possibly the three dimensional configurations of the chromosomes near the nuclear surface play a role in such a process. In summary, DSBs repair involves large-scale motion of the chromosome allowing the break to search for the homologous sequence and for copying the base pairs that have been deleted. During repair, a broken DNA locus scans a large fraction of the nucleus [69, 181]. Interestingly, generating additional damages can affect the dynamics of the homologous-strand, although it is not directly affected. The different steps involved in this repair mechanism remains unexplained. Finally, after a DSB is initiated in the heterochromatin, it moves outside this region to particiapte into the recombination process [49].

There other classical examples where the geometrical organization of the nucleus is involved in regulating its function, including long-range motions in the mammalian nucleus during cell differentiation. In that case, the two X chromosomes needs to meet and to interact. This process can result in X-chromosome inactivation [172]. Large-scale motions are also observed in telomere reorganization and in cancerous cells. Another feature of the geometrical organization is the position of genes: gene regulation depends on their distance to the nucleus surface [266] and in general active genes tend to be found inside the nucleus, as opposed to inactive ones that are located at the periphery.

## 3 Polymer models for modeling DNA dynamics

The renewed interest of classical polymer models [57, 95, 72] answers the need for interpreting, simulating and ultimately understanding the dynamics and structure of the DNA molecule and chromatin. New experiments have called for novel and more accurate polymer models [91] [157]. Basic properties of polymer models have been summarized in [95] and will not be reviewed here and we refer to the classical literature [57, 95] for static and elementary polymer properties, including translocation and reptation [286] through a cylindrical tube.

We recall that DNA is composed of repetitive units of the four fundamental bases guanine (G), adenine (A), thymine (T), or cytosine (C), joint by covalent bonds [282] and can be modeled to the first order as a linear chain composed of connected beads or monomers. A polymer model for a macromolecule is described as a sequence of repetitive structural units. A monomer is a unit representing a sequence of arbitrary length. Once the length is fixed, it defines the scale of the modeling. Thus the same polymer can be used to model a chain with a length ranging from one to hundreds of nanometers. Polymer models have also been used to describe circular DNA (found in prokaryotic cells), where elementary and tractable computations can be performed [134].

How chromatin shall be modeled? are hydrodynamic interactions relevant? For single and double stranded DNA, the local interactions of the chromatin are inferred experimentally from the dynamics of a florescent locus. At a time scale where the chromatin spatial fluctuations is larger than the persistence length *L*_*p*_, single-stranded DNA (*L*_*p*_ ~ 2 *nm*) was suggested to behave as a flexible chain, although the monomer motions are coupled via hydrodynamic interactions [247]. This behavior is well accounted for by the Zimm model [72]. Due to the larger persistence length of double-stranded DNA (*L*_*p*_ = 50 *nm*), the inherent molecular rigidity (rod-like behavior) influences the short-time dynamics that are modulated by hydrodynamic effects [113, 205]. At intermediate times, DNA dynamics is quantitatively described by Zimm theory for a semi-fliexible chain [113, 114]. More elaborated models take into account bending elasticity to force the alinement of three consecutive monomers, as well as rotational energy of the strand [143].

The contribution of hydrodynamic interactions for describing accurately chromatin dynamics is still under discussion [295], but long-range coherent motion of the chromatin over large distances of the order of several *µm* is probably synchronized by ATP-dependent motors [35]. At 100ms and more, where independent forces are applied to chromatin, Rouse and generalized type model are used to describe chromatin motion.

In this section, DNA, chromatin and chromosomes are modeled as polymers, discretized into monomers connected by springs. Analysis and stochastic simulations of the models are used to explore the ensemble of conformations. Polymer configurations obtained in a confined domain depend on the confinement itself [72, 8]: the average end-to-end distance of a polymer is smaller in a spherical domain [103] than in a free space [9]. We first review properties of polymer models starting with Rouse (Fig. 10), which is a quite accurate description of chromatin, as confirmed by microscopy experiments [105, 4]. Interestingly, recent CC experiments suggest that the Kuhn length is less than 5kb [231]. This length scale confirms that the bending elasticity may be neglected for modeling chromatin above this small scale. We further discuss more elaborated polymer models, that include interaction energy such as Lennard-Jones or bending elasticity, phenomenological models, based on self-similar properties and characterized by a scaling exponent [183], loop models [29] and the new class of *β*-polymer model [10] are further discussed for modeling refined chromatin properties. The second part of this section is dedicated to anomalous behavior properties of a monomer.

**Fig. 10.**
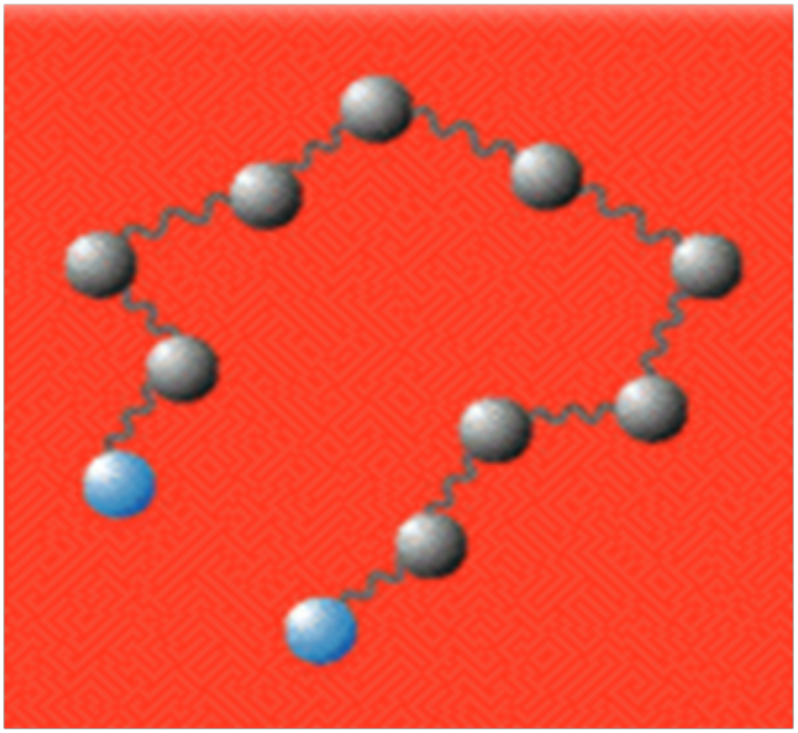
**Schematic representation of a classical Rouse polymer model, where monomers are connected by identical springs.**

### 3.1 Elementary polymer chains

Since the mid-20th century, polymer models were built based on a linear chain, where each monomer is connected to only two adjacent monomers. A *N*-step random walk can be seen as a realization of a polymer of length *N*, where a monomer is centered at each step point. The result of this random walk is a snapshot of a polymer conformation. When the random particle is not allowed to return to the same point, the realization is interpreted as a self-avoiding polymer (in a good solvent). For such realization, the end-to-end vector (*R*_*ee*_) for example, which measures the size and compactness of the polymer is defined by

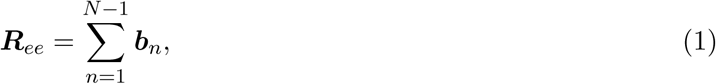

where *b*_*n*_ is a step vector of length *b* (Kuhn length). When the steps are independent, the mean square end-to-end distance is given by [57]

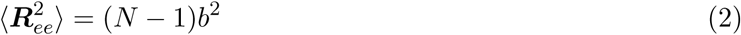

and for long polymers (*N* ≫ 1), the probability distribution function (pdf) of the end-to-end vector *R*_*ee*_ was thought to be a Gaussian variable with zero mean and variance *Nb*^2^. A measure of the polymer size is the *radius of gyration R*_*g*_, which accounts for the position of all monomers: it is defined by

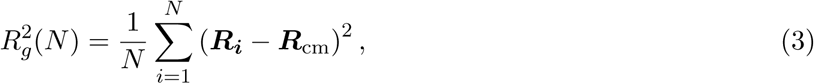

where *R*_*i*_ is the position of particle *i* and the center of mass is 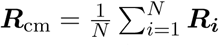 As the number of monomers *N* increases (in a free space), the sequences 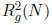 converges to the mean value [57]

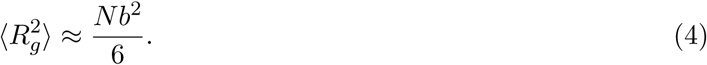

A *Gaussian chain* model is a continuous polymer model where the distance between two neighboring monomers is a Gaussian variable of zero mean and standard deviation *b*. The energy of a polymer configuration is given by the sum over the spring energies

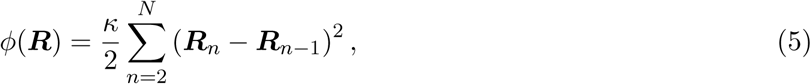

where ***R*** = (***R***_1_***, ***R******_2_ *, …, **R***_*N*_) is the ensemble of monomer positions, connected by a springs of constant 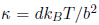 with standard-deviation *b* of the distance between adjacent monomers [72], *k*_*B*_ is the Boltzmann coefficient, *T* the temperature and *d* the dimensionality (dim 2 or 3). The equilibrium pdf of the polymer is given by the Boltzmann distribution

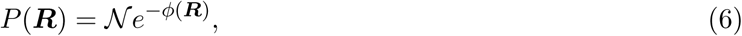

where the normalization coefficient is

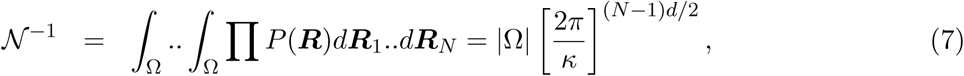

where |Ω| is the volume of the domain, when it is finite, in which the polymer evolves. The normalization constant is computed over the polymer configuration space, in which case the integral is performed over n-1 variables only. The normalization formula 7 is valid for domains large enough compared to the apparent volume 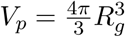 where *R*_*g*_ is the radius of gyration [8].

Another statistical quantity of interest is the distribution of distances between two monomers *m* and *n*, which is the sum of Gaussian variables. The mean square separation distance depends on the distances along the chain and using relation 6, it is given by

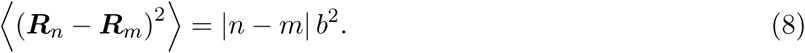

### 3.2 Polymer with self-avoiding interactions in confined domains

In a general polymer model, monomers can interact not only with their nearest neighbors along the chain, but with every one [149, 81]. It is also possible to account for an impenetrable or exclusion volume around monomers, represented by a repulsion force. The associated excluded volume interaction energy *ϕ*_*EVI*_ is described by a short-range potential

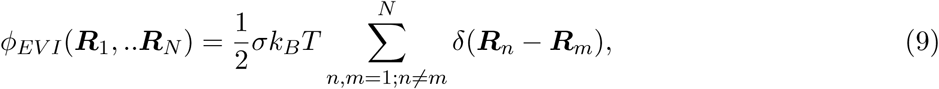

where σ is the excluded volume, which is computed from the radius of the monomer and δ is the classical Dirac-function. In one-dimension, excluded volume interactions lead to a stretch polymer, because it cannot overlap anymore. In higher dimensions, the interactions lead to a significant swelling of the polymer size. For ideal polymers, the end-to-end distance follows the scaling 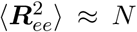 (formula 8), but with additional exclusion interactions, it can happen that:

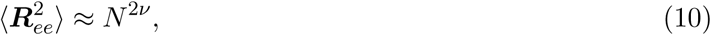

where the exponent ν (Flory exponent) depends only on the space dimension: ν = 3 */*4 for *d* = 2 and ν ≈ 0.588 for *d* = 3 [57].

Elastic and dynamic behaviors of a self-avoiding chain confined inside a cylindrical pore of diameter *D*_*cyl*_ have also been studied using a phenomenological Flory approach and molecular-dynamics simulations. This approach is based on minimizing the free energy for the longitudinal end-to-end distance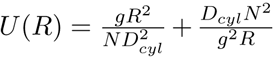where *g* is the number of monomers inside a blob 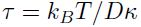 and the excluded volume around each monomer scales like *a*^3^. The chain size was predicted to verify 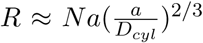 and the slowest relaxation time of the confined chain in the absence of hydrody-namic effects is 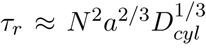 [133], that should be compared with the classical Rouse slowest relaxation mode 22 also proportional to *N*^2^.

The worm-like chain (WLC) model describes a semi-flexible polymers such that the orientation correlation function defined by the tangent vector 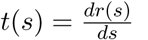 decays exponentially along the chain

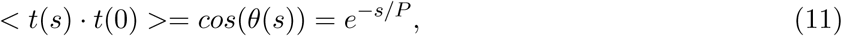

where *P* is the persistence length. We refer to section 3.7 for an elementary construction of the polymer model using the Langevin description. A mean-field approximation model was used in [85] to replace constraints of confinement with a harmonic well approximation (see section 4.6). The simulations of polymers in confined geometries (surface and the interior of a ball cell) allows computing the pressure of DNA packaged in viral capsids. Steady state properties for strongly confined semiflexible polymers, trapped in a closed space or compressed by external forces are further analyzed [229]. Using simulations and scaling argument [42], the escape time of a polymer from a spherical confinement (the polymer has already entered the small hole when it escapes) is predicted to be 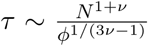 where *v* = 1/2 and the monomer concentration is 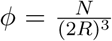 (*R* is the radius of the cavity). This estimate does not account for the mean time that a polymer end finds the small hole on a cavity, where the polymer will eventually exit.

### 3.3 The Smoluchowski limit of the Langevin equation for polymer dynamics

We review now polymer models in the context of stochastic processes that involve a dynamics description, which is the basis of analysis and stochastic simulations. We recall that the motion of a particle in a bath of colliding random particles at equilibrium is described by the classical Langevin's equation [237, 219, 238]: for a particle of mass *m*, the damping force is due friction and is proportional to the particle velocity *v* only through the friction constant. γ It is assumed to be linear and given by the relation *F* = −γ *v*. The friction constant γ depends on the particle geometry and the viscosity of the fluid. The random and friction forces have a similar origin: in a solvent, the random particles collide with each other and the imbedded particle. When the particle of interest moves in a given direction, more solvent particles are colliding with it from that direction, than from the opposite one, generating the effective friction force.

When the dissipation-fluctuation theorem applies [237], which described the particle moving in a bath of particles at equilibrium, the Newtonian equation of motion [45, 237] becomes

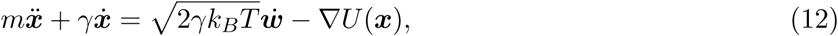

where *U* is a potential energy. In the over-damped limit γ → ∞ of the Langevin equation, the damping term is much larger than the inertia term and equation 12 becomes [237]

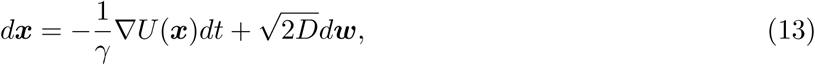

where the diffusion coefficient is *D* = *k*_*B*_
 *T*/γ in dimension 3 [237]. The expression for the diffusion coefficient in dimension two for a cylindrical protein embedded in a membrane depends on the viscosity of that membrane [228]. In this form *w* is a Gaussian noise with mean zero and variance 1.

### 3.4 The Rouse model

The Rouse model for a polymer is a collection of beads connected by springs. The monomers are positioned at *R*_*n*_ (*n* = 1, 2, … *N*), following Brownian motion and the spring forces are due to the coupling of the two nearest neighboring monomers. The spring force originates from the potential energy defined in eq.5. In the Rouse model, only neighboring monomers interact [224]. In the Smoluchowski's limit of the Langevin equation 12, the dynamics of monomer *R*_*n*_ driven by the potential *ϕ*(*R*_1_, ‥, *R_N_*) described by eq.5 and generating the force −∇_*Rn*_ *ϕ*(*R*_1_, ‥, *R_N_*) is

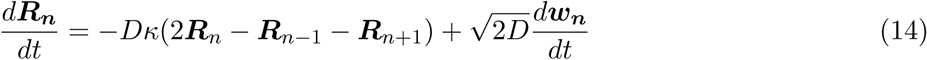

for *n* = 2, ‥ *N* − 1 (the reader should adjust the expression for the first and last monomer). At equilibrium, all beads are centered at zero, but the variance of the distances in a polymer realization is given by

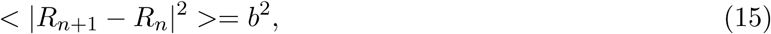

where *b* is the standard deviation of the bond length, *κ*= *dk*_*B*_ *T/b*^2^ is the spring constant with *d* the spatial dimension, *k*_*B*_ is the Boltzmann coefficient and *T* the temperature. A freely-join-chain polymer is a generalized Rouse, where the energy between monomer is given by

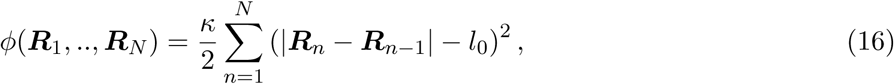

leading to a steady state configuration, where the mean distance between neighboring monomers is < | *R*_*n*+1_ − *R*_*n*_| > = *l*_0_. Taking *l*_0_ = 0 is the classical Rouse model. Starting with a given configuration, the relaxation of Rouse polymer to steady state configuration in a free space can be analyzed using the Fourier space (normal or Rouse modes) [72]

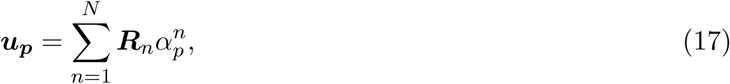

where

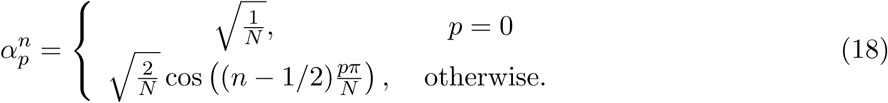

*u*_0_ represents the motion of the center of mass and the potential *ϕ* defined in equation 5 is written with the new coordinate as

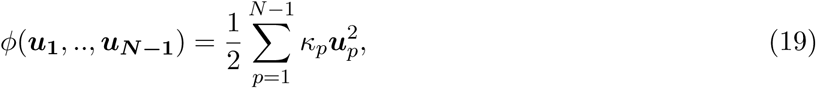

where

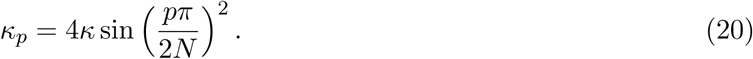

Equations 14 are now decoupled and we obtain a (*N* − 1) *d*-independent Ornstein-Uhlenbeck (OU) processes [277]

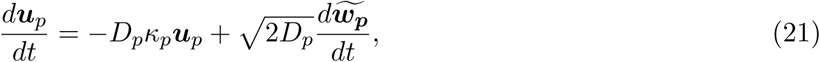

where each 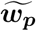 is an independent *d*-dimensional Brownian motion, with mean zero and variance 1 and *D*_*p*_ = *D* for *p* = 1‥ *N* - 1, while *D*_0_ = *D/N* and the relaxation times are τ_*p*_ = 1/ *D*κ_*p*_. The center of mass behaves as a freely diffusing particle. Starting from a straight line, the characteristic time for a Rouse polymer to relax to steady state is constraint by the slowest time constant given by

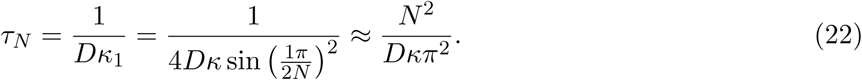

### 3.5 *β*-polymer as a generalized Rouse model

What should be modify from the Rouse polymer model so that the the mean-square-displacement (MSD) for a monomer *R*_*c*_ behaves for small time as

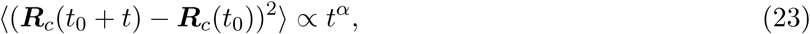

where 〈·〉means averaging over configurations and α *>* 0? is it possible to built a polymer model with a prescribed exponent α? Such construction is possible and the result is a *β*-polymer [10]. Prescribing the anomalous α exponent imposes intrinsic long-range interactions between monomers, beyond the closest neighbors of the Rouse model.

Starting with the Rouse equations in Fourier's space (eq.21), the coefficients κ^~^_*p*_ are modified to

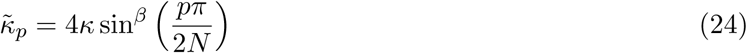

with a free parameter *β >* 1, while *D*_0_ = *D*_cm_ and *D*_*p*_ = *D* for *p >* 0 (see definition of the previous paragraph 3.4). When *β* = 2, we recover the Rouse model. The polymer is reconstructed from the inverse matrix transformation eq.18 between the original and the Fourier space. In the procedure, the coefficients α_*p*_^*n*^ are not changed, only the exponent in the eigenvalue of equation 24.

This procedure defines a unique ensemble of long-range interactions: modifying the eigenvalues results in long-range monomer-monomer interaction as revealed by computing the potential energy which differs from eq.5. Indeed, using the inverse Fourier transform, we get

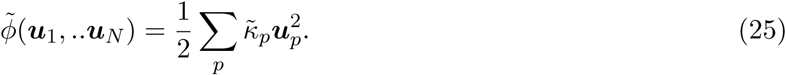

The Rouse transformation in eq. 18 leads to an explicit expression for the interaction between each monomer:

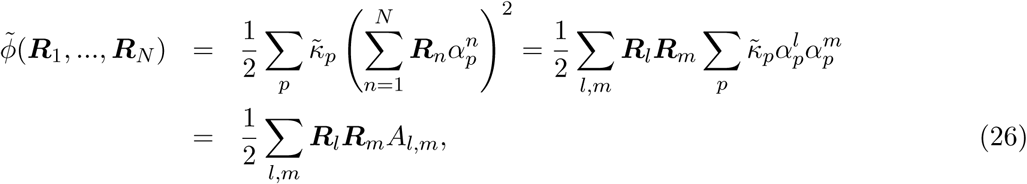

The coefficients are

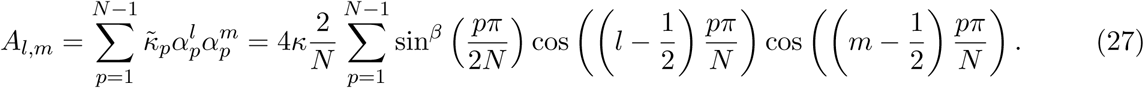

For *β* ≠ 2, all monomers are now coupled and the strength of the interaction decays with the distance along the chain, as shown in Fig. 11a-b. For example, the coefficient *A*_50*;m*_ between monomer 50 and *m* depends on the position *m* (fig. 11b for a polymer of length *N* = 100 and *β* = 1.5). The coefficients *A*_*n,m*_ obtained for various *β* are summarized in table 1.

**Fig. 11.**
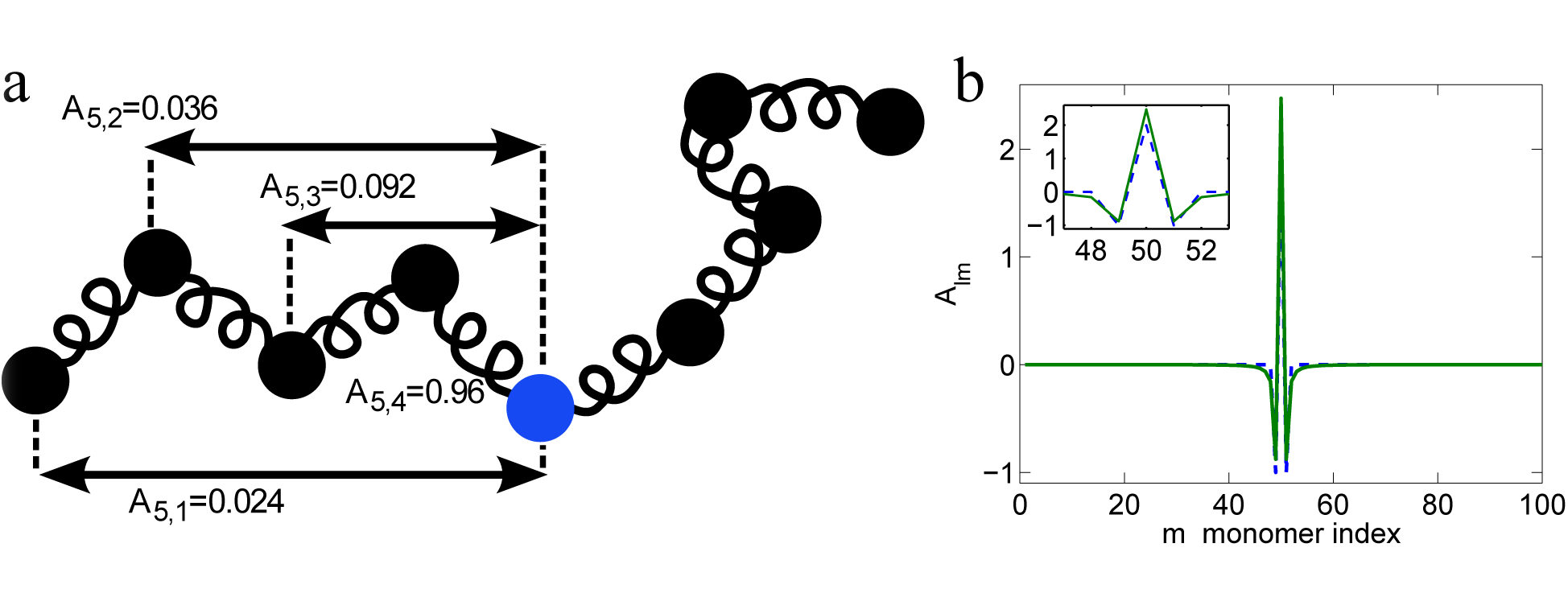
The *β*-polymer. (a) Representation of a *β*-polymer, where all monomers are connected together with a strength that decays with the distance along the chain. The central monomer (blue) interacts with all other monomers in a chain of length *N* = 9 for *β* = 1.5 (interaction unit κ = 3/ *b*^2^).(b) Monomer-monomer interactions in the modified Rouse polymer model (*β*-model). The coefficient *A*_*lm*_ (in units of κ) measures the strength of the interaction between two monomers. Shown are the coefficients *A*_*lm*_ for the polymer with *β* = 1.1, where *l* = 50 and *N* = 100. All monomers interact with each other and the strength of the interaction decays with the distance along the chain (reproduced from [10]).

**Table 1:** Values of the coefficients *A*_*l,m*_ in units of for the middle monomer in a polymer chain of length *N* = 101, for different values of *β*

| $\beta$ | $A_{51,51}$ | $A_{51,50}$ | $A_{51,49}$ | $A_{51,48}$ | $A_{51,40}$ | $A_{51,30}$ |
| --- | --- | --- | --- | --- | --- | --- |
| 2 | 2 | -1 | 0 | 0 | 0 | 0 |
| 1.5 | 2.22 | -0.95 | -0.087 | -0.029 | $-1.07 \times 10^{-3}$ | $-2.2 \times 10^{-3}$ |
| 1.2 | 2.40 | -0.90 | -0.14 | -0.054 | $-3.05 \times 10^{-3}$ | $-7.9 \times 10^{-3}$ |
| 1 | 2.55 | -0.85 | -0.17 | -0.073 | $-5.48 \times 10^{-3}$ | $-16.68 \times 10^{-3}$ |

The long-range interactions between the monomers in the quadratic potential is linked to pair monomers interacting by

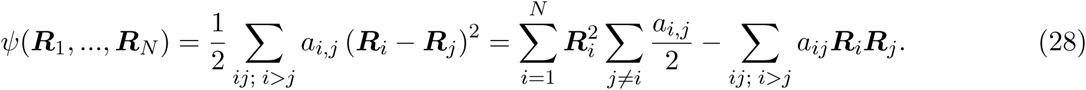

The coefficients *a*_*i,j*_ are related to the *β*-polymer *A*_*l,m*_, (see eq. 26) by the following formula

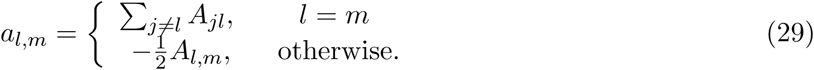

Expression 28 for the potential can be used to reconstruct the amplitude of the interactions from the coefficients of the *β*-model. In section 3.10.2, we show that the anomalous exponent α (eq. 23) is directly associated with the exponent *β* of the polymer, by relation ±= 1-1/β In summary, within a *β*-polymer model, all monomers are coupled and the strength of the interaction decays with the distance along the chain.

### 3.6 Numerical implementation of polymer simulation using Euler's scheme

The elementary method to simulate stochastic polymer models is based on Euler's scheme [238], where the simulation time step ∆ *t* should be chosen such that each monomer moves on average less than the smallest length scale ∆ *x*^⋆^ of the system. This length ∆ *x*^⋆^ can be for example the diameter of a small target. In practice, the small spatial scale imposes a constraint on ∆ *t*. If we used ∆ *x*^⋆^ = *f* × ε where *f* is an extra con dent parameter, fixed at 0.2 and ε is a small parameter, then the time step ∆ *t* should verify the condition √2 *Dδt* ≤ ∆ *x*^⋆^. This is usually a serious limitation of the stimulation, that can be relaxed by using an adaptation time step, that can be big far away from the target and refined close enough. In practice, the numerical scheme is

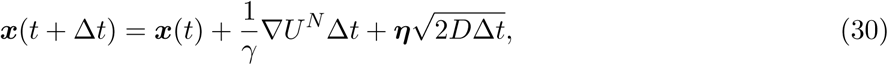

where *U*^*N*^ is the potential 16. In that case (except the end monomers),

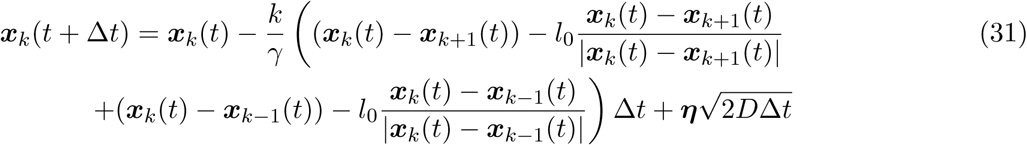

where η is a Gaussian variable with zero mean and unit variance. At a reflecting wall, all monomers are reflected according to the classical reflection for Brownian motion, which follows the Snell-Descartes re ection [238] for an isotropic diffusion coefficient, otherwise the co-normal reflection needs to be used [249].

### 3.7 Implementing excluded volume interactions

Additional interactions such as bending elasticity, which accounts for the persistence length of the polymer and Lennard-Jones forces (LJ) [156], describing self-avoidance of each monomer pairs allow representing more realistic polymers. To account for the LJ-forces, the energy potential is modified to

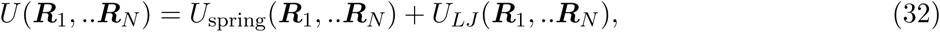

where the spring potential is defined by relation 16 and *U*_spring_ = *ϕ*. The notations are *r*_*i,j*_ = *R*_*i*_ - *R*_*j*_, 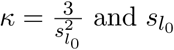 is the standard deviation of the bond length. The Lennard-Jones potential is defined by

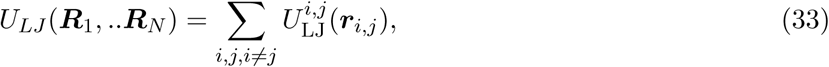

and 

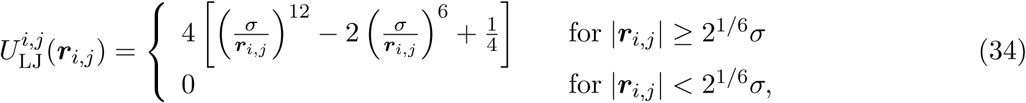

where σ is a cut off, for example the size of a monomer. For the chose *l*_0_ = σ, the springs which materialize bonds cannot cross each other in stochastic simulations of eqs 13 with potential 32.

### 3.8 Modeling stiffness

To account for some stiffness in polymer model, we can add a bending energy to the total energy of a polymer defined as

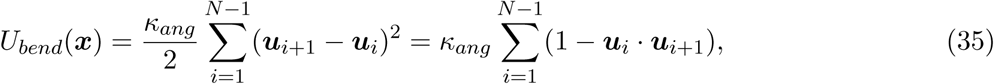

where 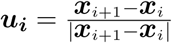 is the unit vector connecting two consecutive monomers and *x_i_* is the position of the *i*^*th*^ monomer. This potential depends on the angle θ_*i*_ between two successive monomers with the relation *u*_*i*_ · *u*_*i*+1_ = cos θ_*i*_. The bending rigidity κ_*ang*_ is related to the persistence length *L*_*p*_ of the polymer by [31]

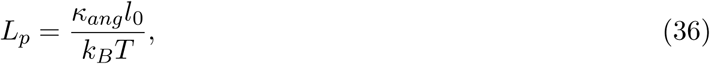

where *k*_*B*_ is the Boltzmann constant, *T* is the temperature, *l*_0_ has been defined above. The persistence length quantifies the stiffness of a polymer and can be characterized using the unit tangent vectors *t*(*s*) and *t*(0) at positions *s* and 0 along the polymer. Averaging over all starting positions, the expectation of the angle *ϕ*_*a*_, which is the angle between *t*(*s*) and *t*(0), decays exponentially with the distance *s* along the polymer

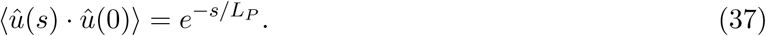

Using the stochastic equation

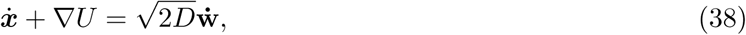

for the total potential

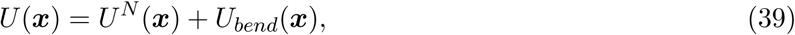

*U*^*N*^(*x*) is the elastic potential defined in Eq.16, the polymer dynamics can be simulated, as shown in figure 12. The parameters are κ_*ang*_ = 5 [257] and *L*_*p*_ = 250 *nm*. In the coarse-grained simulations of chromatin [8], the two energies *U*^*N*^ and *U*_*bend*_ (relation 39) have similar order of magnitude: indeed for the maximum extensibility of DNA about 10% of its total length [37], the length | *r - l*_0_| ∼ 5 *nm*, and *U*_*el*_ = 1/2 *k*| *r - l*_0_|^2^ ∼ 2 × 10^-18^ *Nm*, while the maximum energy between three consecutive monomers due to bending is 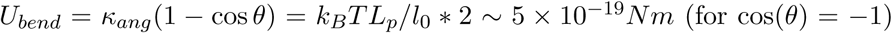 Here *k*_*B*_ is the Boltzmann constant and T is the temperature. A snapshot of a stiff polymer is shown in Fig. 12 (see also [8]). In comparison, a Rouse polymer would not show bending near the sphere, as *p* the polymer configuration is more compact with a gyration radius proportional to √ *N* (parameters are presented in table 2).

**Table 2:** Parameters of the simulations

| Parameters | Description | Value |
| --- | --- | --- |
| $R$ | Radius of the circular/spherical domain | 250 nm |
| $a$ | radius of the absorbing window | 50 nm |
| $l_0$ | Polymer persistence length | 50 nm |
| $D$ | Diffusion constant | $4 \times 10^{-2} \mu\text{m}^2/\text{s}$ |
| $\gamma$ | Friction coefficient | $3.1 \times 10^{-5} \text{Ns/m}$ |
| $k$ | Spring constant | $1.75 \times 10^{-2} \text{Nm}^{-1}$ |

**Fig. 12.**
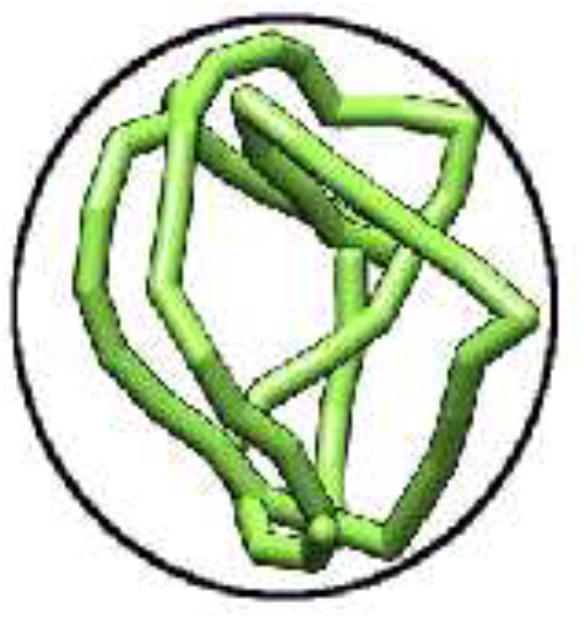
Stiff polymer in a ball. Snapshot of a semi-flexible polymer, locate near the spherical boundary.

### 3.9 Rod-like model

A crude model of polymers is composed of *N* rigid segments where the angle between two consecutive segments is a random variable θ, uniformly distributed in the interval [0 - 2π]. The polymer model consists of a freely jointed chain (see [72] p.8). Each monomer is connected to two neighbors by a rigid bond of length *b*. This dynamics results in long-range correlations between monomers motion, which is different from Rouse. With this model, it is possible to derive close analytical formula. Indeed this model is tractable and the derived asymptotic formula results can be used to guide intuition for other models [12].

In two dimensions, the rod-polymer model is a collection of *N* +1 monomers (points) at positions (*R*_0_, *R*_1_, …, *R*_*N*_) in the complex plane, such that | *R*_*i*_ - *R*_*i*-1_| = *b* (*i* = 1, … *N*) while *R*_0_ = 0. The position at time *t* of the *k*-th monomer is

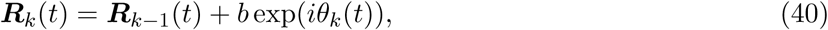

where θ_*k*_(*t*) ∈ [0, 2π] is the angle between the *x*-axis and the vector *R*_*k*_(*t*) - *R*_*k*-1_(*t*). The position of the *k*-monomer at time *t* is the sum of exponential of Brownian motions

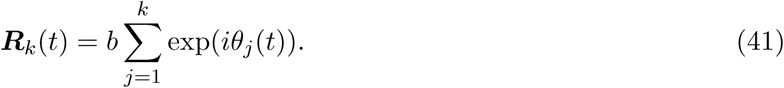

The angles θ_1_. θ_2_, … θ_*N*_ are independent Brownian variables on the circle, satisfying the stochastic equation

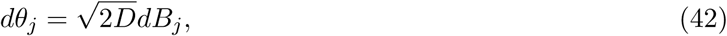

where *D* is the rotational diffusion coefficient (in *s*^−1^), *B*_*i*_ are Brownian variables on a circle of radius 1 with variance 1. The dynamics of the rod-polymer end is represented by the mapping 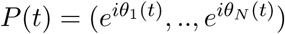 on the *N*-dimensional torus, *R*_*N*_(*t*) = *b*Ψ(*P*(*t*)) where

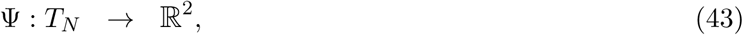

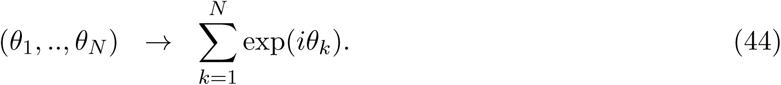

This approach can be generalized to any dimensions (see [294]).

### 3.10 Anomalous diffusion in polymer models

Before describing the property of a single monomer, we recall now the general description of anomalous diffusion. The motion of a small molecular probe located at a position *R*(*t*) at time *t* is characterized by the statistics of its second moment time series given for small time *t* ≪ 1 by 

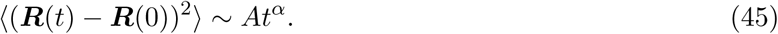

where 〈·〉 means averaging over realization and *A* is a constant. The exponent characterizes the deviation from normal diffusion: when α *<* 1 (resp. α *>* 1) it is the subdiffusion (resp. superdiffusion) regime. Anomalous diffusion was reported in Richardson study (1926) [216] for turbulence, where the exponent in equation 45 is α = 3. Motions for which α > 2 are described as ballistic. The dispersive transport of amorphous semiconductors [233] showed that charge carrier exhibits also subdiusion.

Particles in percolative systems [142] and porous media [140], collective diffusion on solid surfaces [163] and in bulk-surface exchange controlled dynamics in porous glasses [38] also exhibit superdiffusion. Micelles traveling by motor proteins are characterized by an exponent α = 3/2, while DNA-binding proteins diffusing along a double stranded DNA can perform subdiffusion [250].

A protein that detaches from a DNA chain and connects to another segment nearby in three dimensions but far along the chain is modeled as a Lévy flights [33, 161]. When a chromatin locus is modeled as a Rouse polymer, the anomalous exponent is α = 1/2, but other polymer models lead to other exponents. For the Zimm model, accounting for hydrodynamic interactions, the exponent is α = 2/3 [296] while for a reptation-polymer, α = 1/4 [57]. Analysis of single particle trajectories of chromatin locus reveals that the anomalous exponent is α = 0.33 [104]. Determining the origin of anomalous diffusion in a crowded environment, the cytoplasm or the nucleus remains a challenging problem, because different physical properties (polymer intrinsic mechanical properties, external forces applied to the polymer, etc.) are reflected in that anomalous exponent.

Theoretical models to describe anomalous diffusion are continuous-time random walk (CTRW), obstructed diffusion (OD), fractional Fokker-Planck [138, 139], fractional Brownian motion (FBM), fractional Levy stable motion (FLSM) or fractional Langevin equation (FLE). Phenomenological models [284] based on fractional Langevin equation leads to a MSD that exhibits a power law. The construction of the associated polymer model relies on the Langevin equation with a memory kernel that decays algebraically (see relation 58). This kernel accounts for the properties of the viscoelastic fluid, which slows down any loci dynamics.

We start now by the generalized Brownian motion called *continuous time random walk* (CTRW), a classical model of subdiffusion. In such model, the waiting time between steps and the length of the step are taken from probability distributions φ (*t*) and λ (*x*) respectively. Their corresponding mean time and variance are

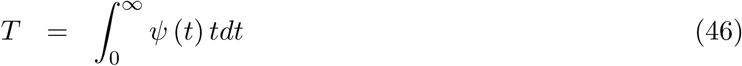

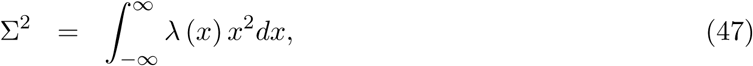

which can be either finite or not. When the distribution of waiting times is Poissonian and the steps length are Gaussian distributed, it is possible to reproduce in the long-time limit the regular Brownian dynamics. When the waiting time distribution has a long-tailed asymptotic behavior 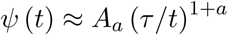 with 0 *< a <* 1, the mean time is in finite. These conditions are used to model a particle trapped for long time (amorphous semiconductors [233]).

For a Gaussian distribution of λ (*x*), the probability distribution *P* (*x, t*) of a particle *X*(*t*) to be located at position *x* at time *t* satis es the *fractional Diffusion (or Fokker-Planck) equation* (FDE)[141]

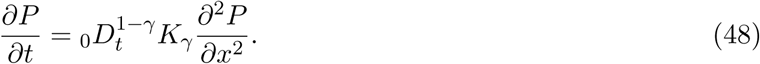

which reduces for γ = 1 to the classical diffusion equation. _0_ *D*_*t*_^1-γ^ is the Riemann-Liouville operator defined for 0 *<* γ *<* 1 by

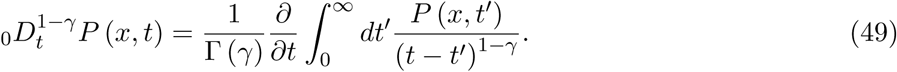

The fractional differentiation of a power *q* ∈ R is

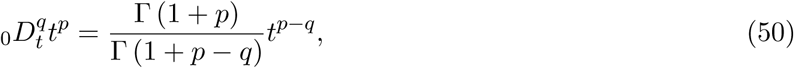

Where Γ is the Euler's Γ-function. Computing the mean square displacement from equation 48 with initial condition *P* (*x*, 0) = δ(*x*) [141] gives (with *x*(0) = 0)

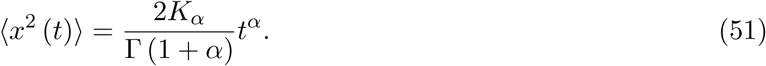

The Lévy ights (LF) model, which is a Markov process with a large jump length distribution is characterized by an asymptotic power law

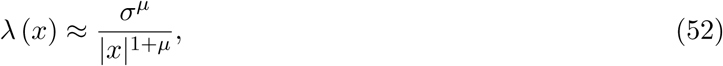

where the variance is in finite. Trajectories are characterized by an exponent μ [179]. Unlike Gaussian random walk that ‘fill’ the area (μ = 2), LF-trajectories consist of self-similar clusters, separated by long jumps. The Fourier transform of the function λ is proportional to | *k*|^μ^ (equal to | *k*|^2^ for a Gaussian process). The fractional derivative is defined using the Fourier transform [179] as

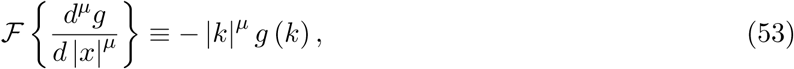

where 1 ≤ μ < 2. Lévy process can be defined though a fractional diffusion equation 

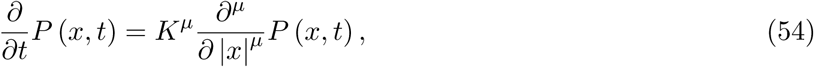

with a generalized diffusion coefficient 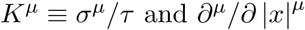is the Riesz-Feller derivative.

The probability density of a LF decays as a power-law 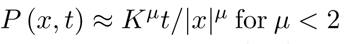 Consequently the mean-square-displacement is in finite. A Subdiffusive process *B*_*H*_ (*t*) can also be generated by the FBM [230, 130]. It has the following properties

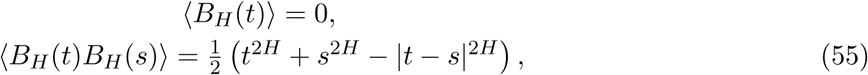

where the parameter *H* is called the Hurst exponent (0 *< H <* 1). This process is constructed using a generalized Langevin's equation [144] defined as

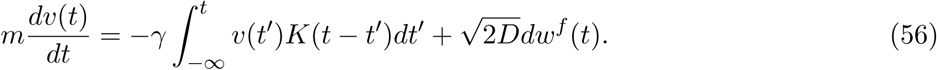

This equation is not written in the strong damping limit, for which the inertia term remains (*m* is the mass). The memory kernel *K*(*t-t*^*′*^) relates to the fractional noise *dw*^*f*^(*t*), following to the autocorrelation function by the relation

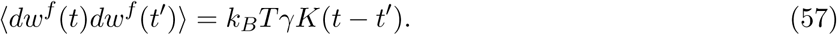

When the kernel is a δ-function, the classical Langevin's equation is recovered. However, for a kernel with a long-time decay (such as a power law), a sub-diffusion behavior is found in the FBM. With the kernel

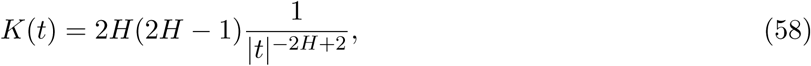

the anomalous diffusion exponent is α = 2 *H* and the fractional noise is defined by [145, 63]

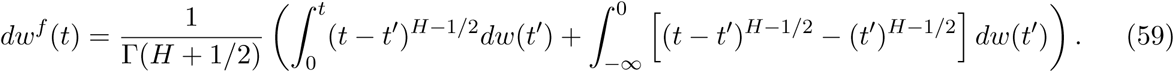

Finally, the variance of the position of a particle governed by eq.56 is

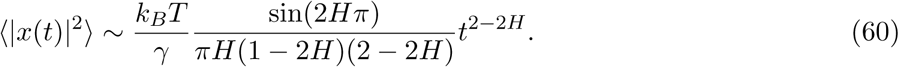

To conclude a particle with the characteristic described by eq.56 performs anomalous diffusion with exponent α= 2 - 2 *H*. The FBM is used as a phenomenological model [284] to describe chromatin locus dynamics. The power-law decaying kernel accounts for the motion of the loci in a viscoelastic fluid. In that medium, the motion of a locus as well as the chromosome dynamics are slowed done, resulting in a sub-diffusion regime. Relating the exponent *H* to the local chromatin properties or the nuclear environment remains a challenge.

#### 3.10.1 Normal and anomalous diffusion regimes for a single monomer

The statistical properties of the k^*th*^-monomer motion of a polymer containing *N* monomers can be computed empirically from its time position *R*_*k*_(*t*). Its motion is however non-Markovian, but the motion of the ensemble (*R*_1_(*t*), ‥, *R*_*k*_(*t*), ‥ *R*_*N*_(*t*)) is a Markovian process, satisfying the stochastic equations 14. A fundamental characteristic is the MSD (relation 23) for small and intermediate time *t*. We present here analytical MSD computations for a Rouse and β-polymer. Similar expressions are not known for general polymer model with additional physical forces such as bending elasticity or LJ forces.

#### 3.10.2 Anomalous motion of a Rouse polymer

The position of monomer *R*_*c*_ of a Rouse polymer is given by

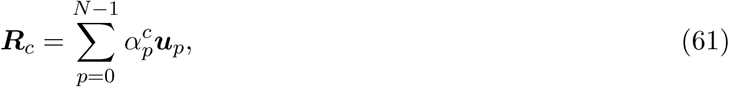

where α^*c*^_*p*_ are described by relation 18 and the vectors *u*_*p*_ satisfy eqs.21, which form an ensemble of *p*

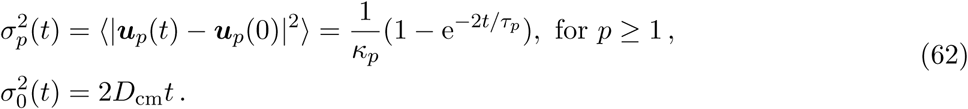

The relaxation times are defined by

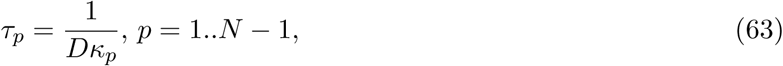

while the diffusion constant is *D*_cm_ = *D/N*. The shortest time is τ_*N*-1_ ≈ 1/(4 *D*κ) which is half of τ_s_ = 1/(2 *D*κ) during which a free monomer diffuses a mean square distance between adjacent monomers (*b*^2^ = 1/κ). The center of mass is characterized by the time scale τ_0_ ≡ *b*^2^ *N/D*_cm_ = *N*^2^/(*D*κ), that a particle diffuses across the polymer size. For long polymers τ_0_/τ_1_ ≈ π^2^. Using relation 62, the MSD of monomer *R*_*c*_ is a sum of independent Ornstein-Uhlenbeck (OU)-variables:

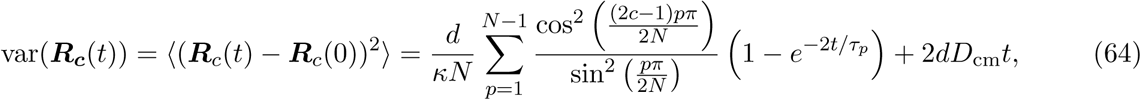

where *d* is the spatial dimension. Formula 64 shows the deviation of the MSD compared to Brownian motion. There are three distinct regimes:

1. For short time 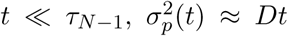, which is independent of *p*, (as shown by a Taylor expansion of the exponentials in eq. 62). The sum in eq. 64 gives

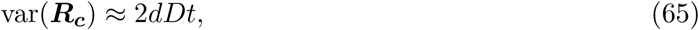

which characterizes diffusion.
2. For large time *t* ≫ τ_1_, after relaxation, the exponential terms in relation 64 becomes independent of *t*. Only the first term in Eq. 62 corresponding to the diffusion of the center of mass gives the time-dependent behavior. This regime is dominated by normal diffusion, with diffusion coefficient *D/N*.
3. For intermediate times τ_*N*−1_ ≪ *t* ≪ τ_1_, such that 2 *t/*τ_*p*_ > 1, the sum of exponentials contributes to eq.64. The variance 64 can be approximated by

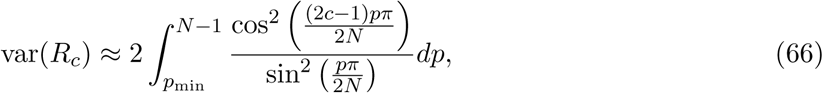

where *p*_min_ is such that τ_*p*__*min*_ = 2 *t*. We shall see below that var(*R*_*c*_) ∼ *t*^1/2^. Due to the behavior of a Rouse polymer, all internal relaxation times τ_*k*_ contribute to the intermediate time, leading to anomalous diffusion for a Rouse monomer. The time interval can be arbitrarily long with the size *N* of the polymer.

We present below the analytical derivation of the power law behavior.

#### 3.10.3 Monomer motion characterized by a power-law decay time

To describe the large range of anomalous exponent of chromatin locus diffusion, the Rouse potential *ϕ*_Rouse_ (relation 5) is introduced in eq. 56 [284], so that every monomer is characterized by a subdiffusion regime. The motion of a monomer is described by

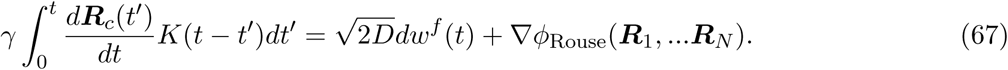

The power-law kernel *K*, described in 58, accounts for the motion of the loci in a viscoelastic fluid, which slows done the chromosome dynamics, leading to a sub-diffusion regime. For the middle monomer, the long-time asymptotic of the MSD is given by [284]

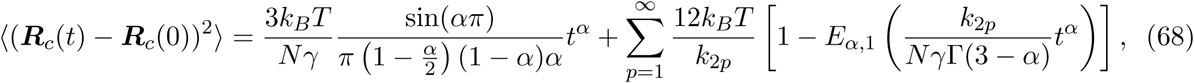

where *E*_α,1_is the generalized Mittag-Leffler function 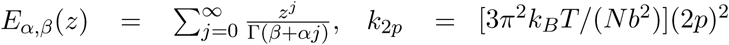 Here, for short times, the MSD of the middle monomer is

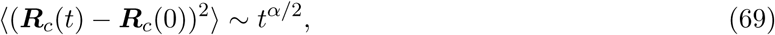

where 0 < *α* < 1. At long times, the motion is governed by that the center of mass leading to

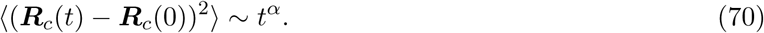

#### 3.10.4 Correlation function for a Rouse and *β*-polymer

We have already introduced the polymer *β* model in section 3.5. This class of polymer resolves the following question: construct a generic polymer model for which the anomalous exponent of a locus is prescribed. A large variability of the anomalous exponent for the dynamics of a chromatin locus cannot be accounted for by the viscoelasticity alone, subdiffusion with an exponent α > 0.5 can also appear in some polymer model [11], a case that involves a deterministic force or directed motion or other monomer interactions (self-avoiding and bending interactions).

We present here the computation of the cross-correlation function for monomer *R*_*c*_ using eq. 61. For *N* ≫ 1 and intermediate times τ_*N*−1_ ≤ *t* ≤ τ_1_, we show that

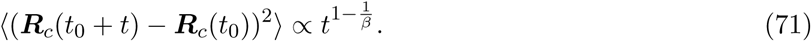

Using 25, for any time *t*_0_ and *t*, a direct computation from eq. 61, with 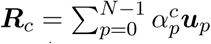 and given the correlation function of *u*_*p*_ which is 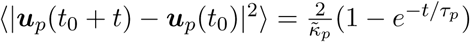 we get 

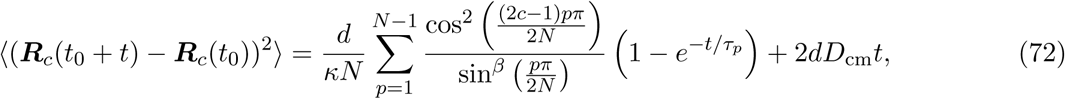

where 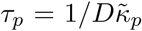. We plot in fig.13b the cross-correlation function of a monomer computed from Brownian simulations of a polymer with length *N* = 128 and compare the cross-correlation function for *β* = 3/2 with the regular Rouse model (*β* = 2). In the intermediate time regime, where 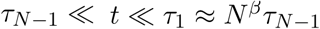 (anomalous regime), eq.72 can be estimated. For large *N*, using Euler-Maclaurin formula, the cross-correlation function is approximated for 1 ≤ β ≤2 by [10]

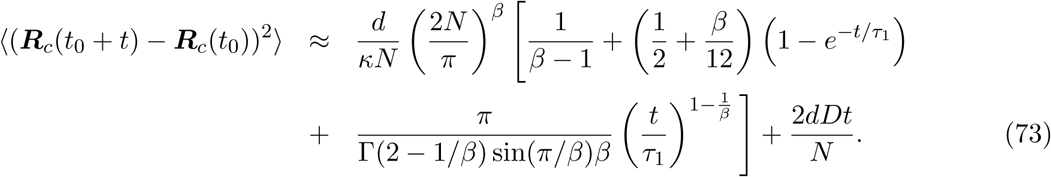

Note that this formula uses that 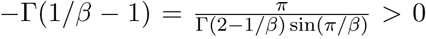 and Γ (2−1/ β) *>* 0, because β ≥ 1 *>* 1/2. We conclude that for intermediate times (τ_*N* − 1_ ≤ *t* ≤ *τ*_1_), the center of mass diffusion does not contribute to the process and thus the cross-correlation function scales with time as shown in eq.71: while the exponent α is an indicator of anomalous dynamics, it is a challenging task to relate it to the hidden underlying driving forces, local geometrical organization and crowding organization.

**Fig. 13:**
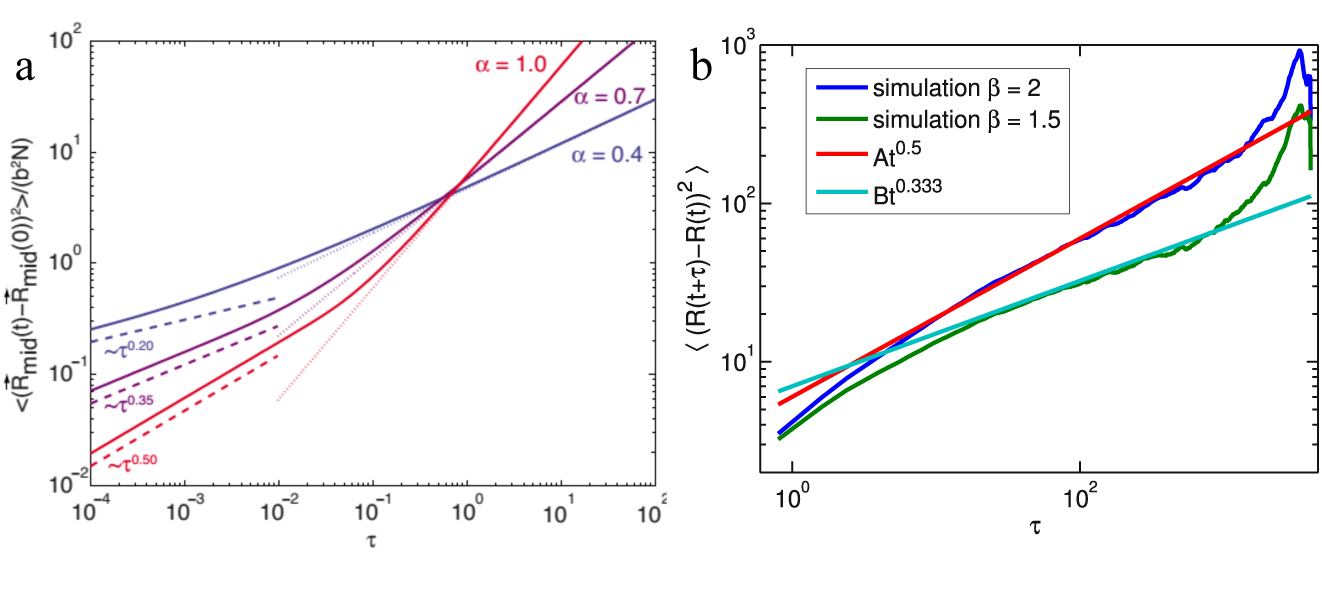
Anomalous diffusion properties of a single monomer. (a) MSD computed for the midpoint monomer ∹(R_mid_(*t*) − R_mid_(0)^2^)〉/ *b*^2^ *N* of a fractional Langevin motion for a polymer versus the dimensionless time τ = *t*/ [ *N*^2^ *b*^2^γ/ *k_B_T*]^1/α^ for *α* = 1.0 (red), α = 0.7 (purple), and *α* = 0.4 (blue) see [284]. Analytical solution of (eq.68) (solid lines) are represented for the three values with the long-time asymptotic behavior (dotted) and short-time scaling of *τ*^α/2^ (dashed). (b) Brownian simulations of a polymer (length *N* = 128) for different values of *β*. The correlation function is computed from simulated trajectories. For *β* = 3/2 (green line), the anomalous exponent is *α* = 0.33 at intermediate times (eq.71). The trend line (cyan) is also plotted. For the Rouse model *β* = 2 (blue line), the anomalous exponent is *α* = 0.5. Also plotted is the trend line (red) according to eq.71.

## 4 Polymer looping and the search for a small target: a mean passage time study

This section is dedicated to polymer looping and the estimation for the rate of this process. We start with early models [202], followed by non-Markovian dynamics [287, 288, 261, 64, 254, 99, 100] and then asymptotic approaches [13, 9, 14]. We also present simulation results for the search for a small target and the associated Mean First Encounter Time (MFET), as asymptotic computations are still lacking. For a recent review of the search process for a small target by Brownian particles, we refer to the Narrow Escape Theory [119, 118, 55]. We recall now the context of chromatin looping.

DNA short and long-range loops are observed in chromatin at various times and spatial scales (as we will discussed in the next section 5). A loop is formed when one piece of the chromatin is brought into close proximity of another one, although the two parts could be located far away in terms of the genomic distance. The continuous change of chromatin configuration that lead to looping can result in gene direct interaction [209] and expression [148], modulation, activation or repression. DNA and polymer looping have been investigated experimentally [66, 76], numerically [47] and analytically [288, 261, 202, 13, 9] over the past 40 years in the context of polymer physics with recent renewed interest in the recent years. Some early computations for the time a polymer knot become untied [286] are not discussed much here, but these computations are quite relevant to interpret data about two-tagged locus located on the same DNA.

The questions to consider are: estimating the mean looping time in free and confinement domain as well as the search time for a polymer locus to find a small target, which tends to infinity as the size of the target tends to zero. Deriving analytical formula allows exploring continuously the parameter space (where variables are the target size, the polymer length, the bond strength, etc…) at low cost compared to numerical simulations. In the context of the nucleus, such formulas provide estimates for the rate that two loci come into close proximity or the time a DNA fragment meets a small target driven by random collision. They are used to interpret data (see section 5).

The mean time for two polymer ends to meet (Fig. 14), starting from any given configuration has several implications for DNA looping because a gene can be activated when a transcription factor bound far away from the promoter site is brought near the active site [66, 209, 76]. The mean first encounter time (MFET) is de ned as the rst arrival time for the end monomer into a ball of radius *ε*, centered at the other polymer end (Fig. 14c). The MFET depends on the radius *ε* and the polymer length *N* (measured in the number of monomers). For a Rouse polymer, characterized by the relaxation time *τ_p_* [72] where the slowest is proportional to *N*^2^ ([254] and section 3.4), the MFET depends both on the initial end-to-end distribution [261, 287, 82] and the radius *ε* of a ball located around one end (Fig. 14c). These two time scales are reflected in the asymptotic regimes, identified *p* numerically [47, 78] and characterized by the ratio 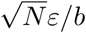 (*b* is the standard deviation of the bond length). When this ratio is of order 1, the MFET depends on *ε* and scales as *N*^3/2^, when it is ≫ 1, it is dominated by *N*^2^ and is independent of *ε*. In general, the MFET shows both scales [273] in *N* as we shall see here.

**Fig. 14:**
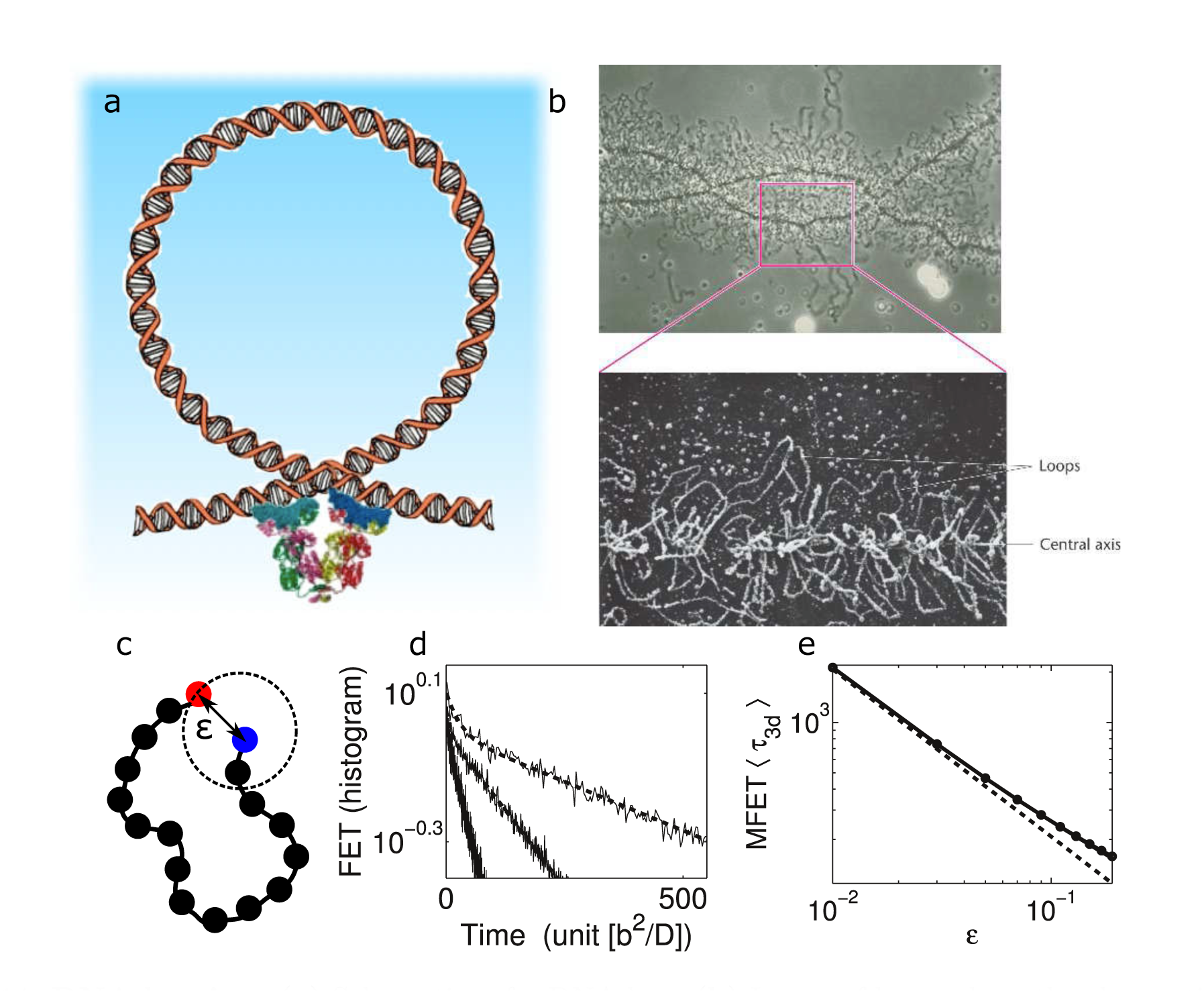
DNA looping: (a) Schematics of a DNA loop (b) Image of loops along the chromatin (c) Scheme of a polymer loop, occurring when the two ends are located at a distance *ε* from one another. (d)) Histogram of the rst encounter times (FET) obtained from Brownian simulations in three dimensions (full line) and tting with two exponentials (dashed line) for different polymer length (*N* = 16, 32, 64 (left to right) and *ε* = 0.1 *b*. The distribution of looping time is well approximated by a single exponential for small *N*, but two are necessary for larger *N*. (e) MFET as a function of the radius *ε* in three dimensions. Comparison of the Brownian simulations (full line) with the reciprocal of the rst term in the expansion of the rst eigenvalue (eq.114)(dashed line) and eq.175) (circles).

For a worm like-chain model [125] with hydrodynamic forces, self-avoidance and coulomb forces, the MFET depends on several parameters such as the polymer length and the bending elasticity, as shown numerically [206, 256]. The MFPT has been estimated using analytical methods [288, 261, 202, 273, 82], numerically [47] and experimentally [218, 7] and also when one end was tethered to a surface [271].

To derive the dependency of the MFET on *N*, we shall recall its formulation as a mean first passage time to a small boundary, [237, 202] which is a boundary value problem for the second order Laplace equation in a high-dimensional space, defined by the polymer configuration space. The MFET is computed by expanding the first eigenvalue for the Fokker-Planck operator associated to the stochastic Rouse dynamics. Although the Markovian aspect of looping and search process for a small target by a polymer locus was established in early models [202], the non-Markovian aspect was recently considered [254, 99, 100], but the search time is in fact well-approximated as a Poissonian process [8, 9, 14].

### 4.1 Brief analytical formulation

The first looping time between two monomers is the First Encounter Time *τ_e_* for two monomers *n*_*a*_, *n_b_* to come into a distance *ε < b*. It is de ned by

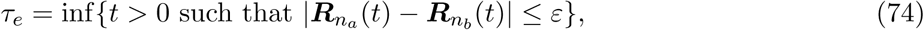

where *R*_*n*__*a*_ and *R*_*n*__*b*_ follows for example the Rouse equation 14. The goal of the section is to present several analytical studies to compute the Mean First Encounter Time (MFET) 〈 τ_*e*_ 〉.

### 4.2 Szabo, Schulten& Schulten approach and later analysis

Applying the first passage times theory [236, 237, 238] to diffusion controlled reactions, in a pioneering article Szabo, Schulten& Schulten [261] studied intramolecular reactions of polymer end groups by using differential equations. Depending on the boundary conditions, chosen to be absorbing, or radiative boundary condition (partial reflection), they derived various asymptotic expressions for the chemical reaction time. For a diffusion process *X*(*t*) satisfying the stochastic equation

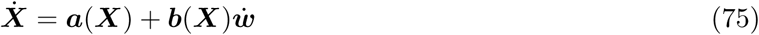

where *a* is a deterministic drift, *b* the diffusion tensor and *w* the classical Wiener δ-correlated noise, the MFPT for the arrival time 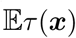 starting at position *x* = *X*_0_ is solution of

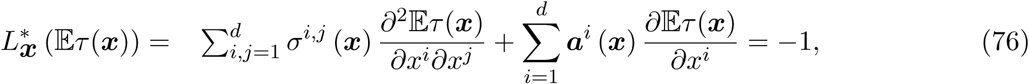

in dimension *d*, where σ(*x*) = *b*(*x*) *b*^*T*^ (*x*). An absorbing boundary condition is imposed on the target and reflecting boundary condition are given on other part of the boundary. In particular, in spherical coordinate in *d*-dimensions, for a gradient field *a* = − *Dβ∇U*, equation 76 becomes

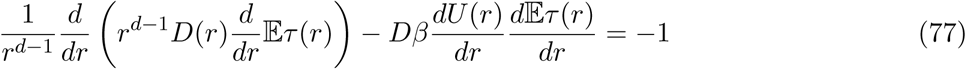

with reflecting boundary condition at *r* = *R* and absorbing at *r* = *a*. After a double integration, the mean time is

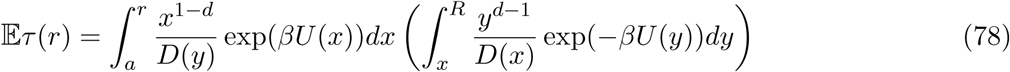

and the mean 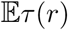 averaged over the steady state distribution 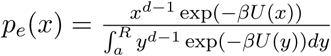

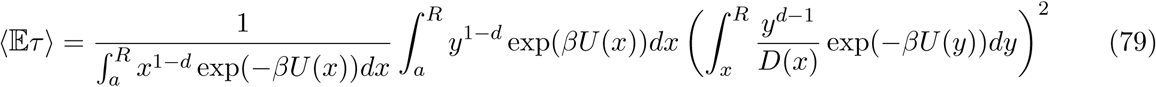

The dynamics of the end-to-end distance *r* = | *R*_*N*_ − *R*_0_| is described by a Smoluchowski-type equation and the overall effect of the polymer can be approximated by harmonic spring potential, i.e., 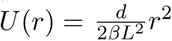 where *L* is the mean distance between the polymer ends. When the radius *R* is large, but *a* is small, the asymptotic expansion of 79 for *d* = 3 is

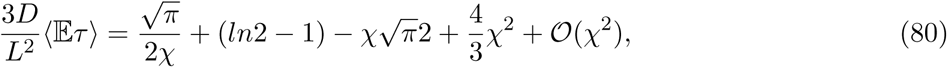

where 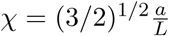 In addition, numerical simulations suggest that the passage time distribution is well approximated by a single exponential, where the rate is 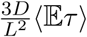 This approach was later on improved in [202], followed by a discussion of the validity of the asymptotic expansion, depending whether the radius *a* is larger or smaller compared to *b* (equilibrium mean length of a single bond) and 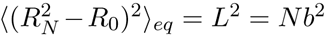. Indeed, the equilibrium distribution p(*r*) of the end-to-end distance r can be approximated by a Gaussian with the mean squared distance *L*2,

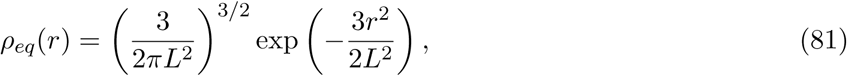

the following radial Laplace equation is solved with a weight _*eq*_ and an absorbing boundary conditions at *r* = *a*:

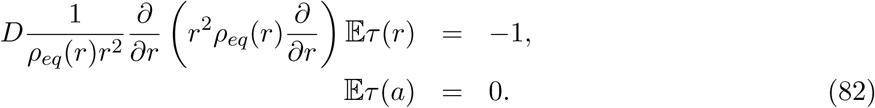

The effective diffusion coefficient *D* was chosen empirically, based on numerical integration *D* = 2 *D*_0_, (*D*_0_ is the diffusion coefficient of a single monomer in the Rouse equation) to account for the diffusion of the two ends, neglecting the effect of all other monomers. Equation 81 assumes that the local equilibrium approximation is valid, but in reality, it is valid only when the rate of approach to local equilibrium is much faster than the rate of the end monomers to meet. The time scale to equilibrium is governed by the largest Rouse mode relaxation time (see eq. 22) 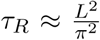 in units (*b* = *D* = 1) and does not depend on the radius *a*. The local equilibrium condition can be satisfied only when the contact distance *a* is very small. In the small *a* limit, the mean first passage time, predicted by eq. 82 is

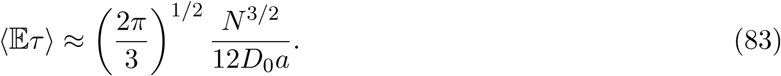

If *a* is much smaller than 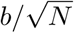, the rate to equilibrium is faster than trapping, for which formula 83 is valid.

In summary, in the limit of small *a* for fixed 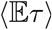 is of the order of *N*^3/2^/ *a*, while in the limit of large *N* at fixed *a*
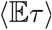 is of order *N*^2^ and independent of *a*. But as we shall see below, in general, the looping time formula combines both behavior in *N*^3/2^/ *a* and *N*^2^. A general computation of the asymptotic behavior for the looping time requires considering the motion of the entire polymer and not simply the two end distance, as we shall see below [13].

### 4.3 Closed loop configuration space for a rod-like polymer

We introduce in sub-section 3.9 rod-like polymer models. We describe now the configuration space of loops. The MFET 〈τ_e_〉 for the two ends of a rod-polymer is equivalent to the one for a Brownian particle *P* (*t*) = (*e*^*i*θ_1_(*t*)^,.., *e*^i *θ*_*N*_(*t*)^) located on the torus *T*_*N*_ in dimension *N* (length of the polymer) to meet for the first time the boundary of the domain

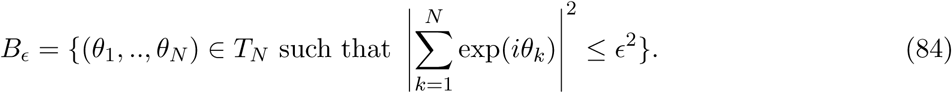

The domain *Bϵ* is a tubular neighborhood of a *N* - 2-dimensional manifold defined by the *B*_0_ (taking *ϵ* = 0 in equation 84). *B*_0_ is of codimension 2, because it is the set of points defined by two independent equations

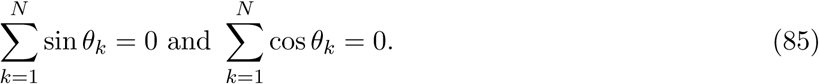

The boundary ∂ *B*_ϵ_ is represented for *N* = 3 and *N* = 4 in Fig.16 for *ϵ* = 0.2 *b*. The boundary ∂ *B*_ϵ_ is analyzed by setting *ϵ* = 0 corresponding to closed configurations (Fig.15): for *N* = 3, the ensemble of closed chains (Fig. 15B) consists of two equilateral triangles (for each orientation) with one vertex at zero. This ensemble is invariant by planar rotations centered at zero. Under this action, the ensemble consists of two orbits and any reflection (preserving zero) permutes these two orbits. For *N* = 4, the three different configurations are shown in Fig. 15C): it consists of a polygon and two non-isomorphic configurations. These structures are connected by continuous deformations. In that case, the boundary ∂ *B*_ϵ_ is made of domains, connected by narrow structures (Fig. 16b). The description of *B*_0_ for large *N* is quite difficult due to lack of parametrization [12].

**Fig. 15:**
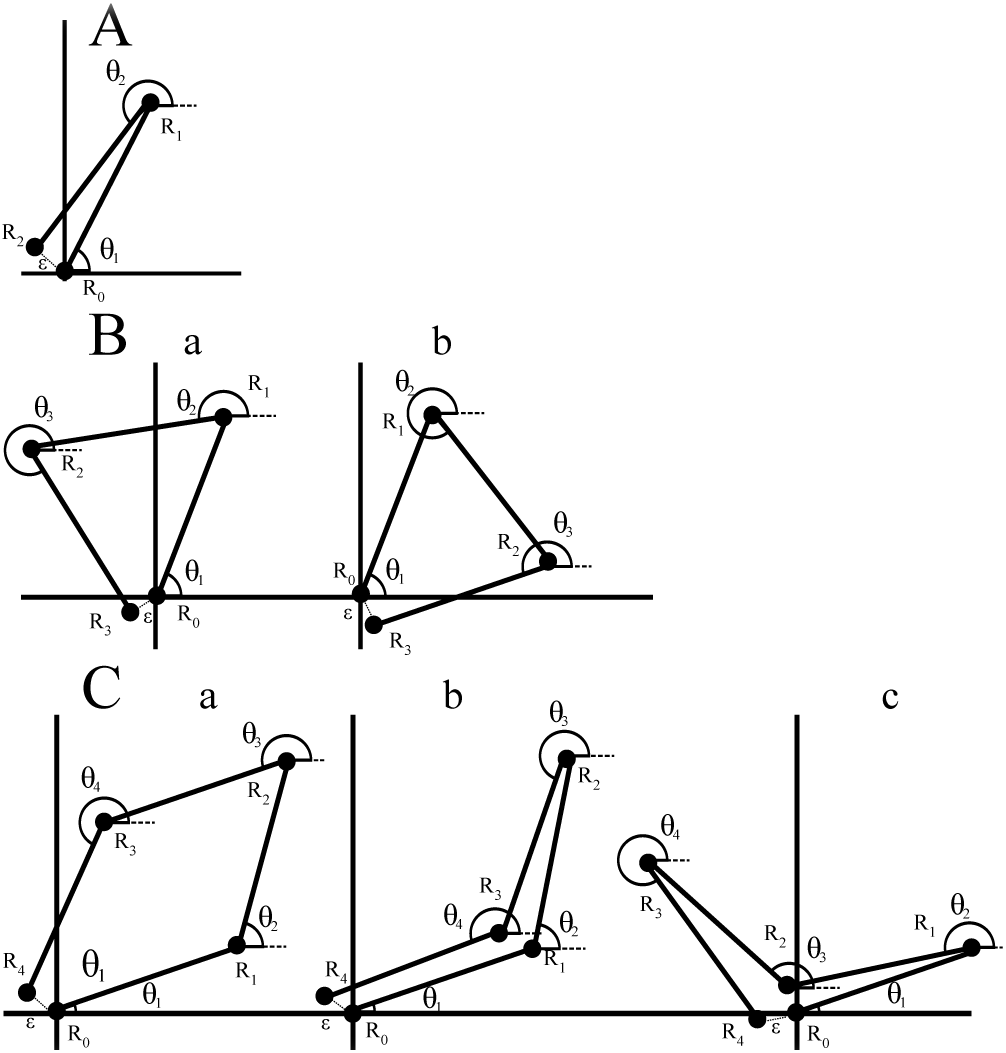
Schematic representation of polymer loops for different lengths. Monomers positions are ***R**_k_* while ***R***_0_ is fixed at zero. *¸_k_* is the angle between ***R**_k_*– ***R***_*k*−1_ and the positive direction of the *x* axis. ε is the absorption distance. **A**: two bonds polymer (*N* = 2): there is only one closed configuration (up to rotation). **B**: three bonds polymer (*N* = 3), showing closed that the only configuration is an equilateral triangle (the loop can be either clock-wise or counter clock-wise). **C**: four bonds polymer (*N* = 4), showing three possible closed configurations [12].

**Fig. 16:**
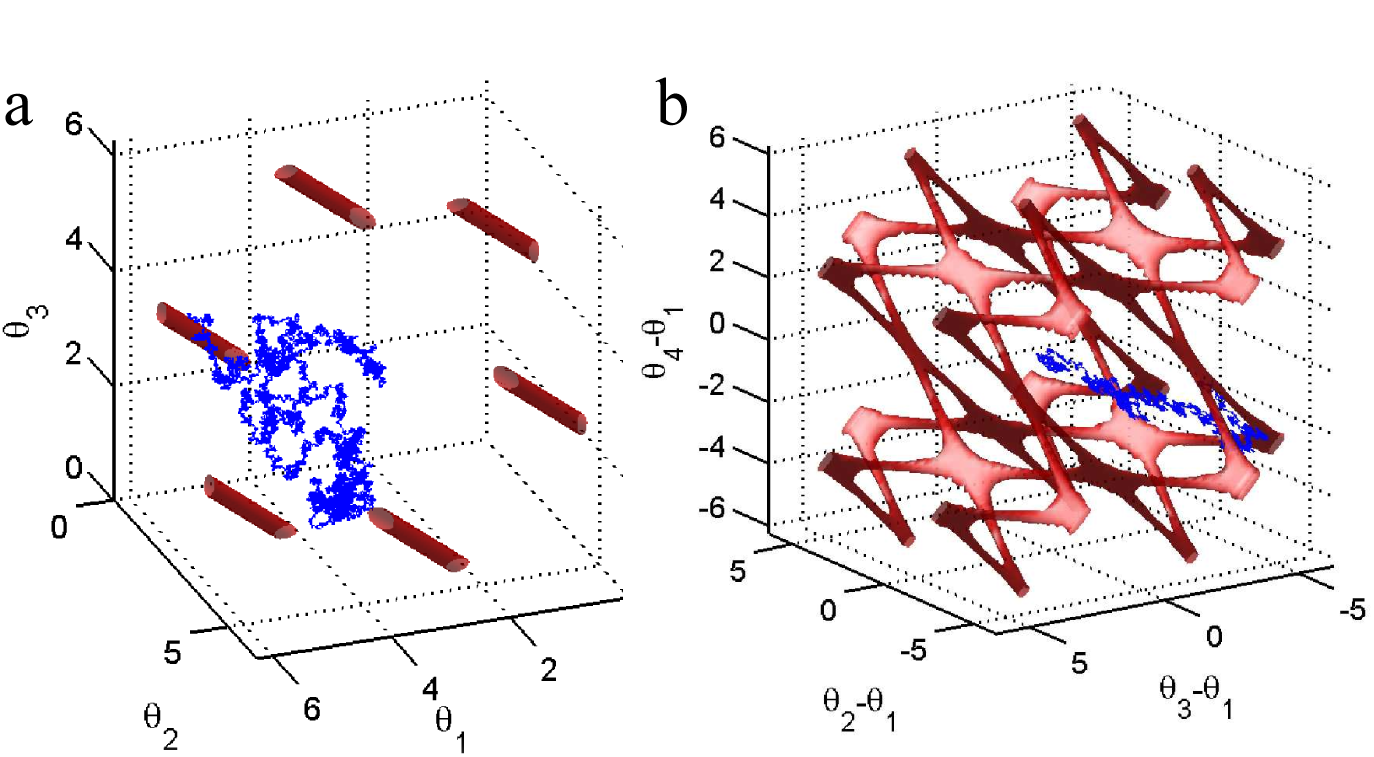
The boundary of *B_ϵ_* for a rod-polymer. **a)**: for *N* = 3 and *ϵ* = 0:2 in [0, 2π] * [0, 2π] * [0, 2π]. Although figure (a) show 6 disconnected, there are only two of them in a unitary torus. A Brownian trajectory (blue) in the angle space moves until it is absorbed.**b)**: for *N* = 4 and *ϵ* = 0:2b. The manifold consists of domains connected by narrow tube, corresponding to deformation of loops [12].

### 4.4 Looping for a Rouse polymer

The two ends *R*_*N*_; *R*_1_ meets when their distance is less than *ɛ < b*, that is

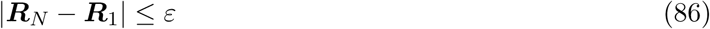

In Rouse coordinates, 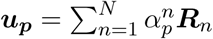 where 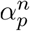 are defined in 18, condition 86 is

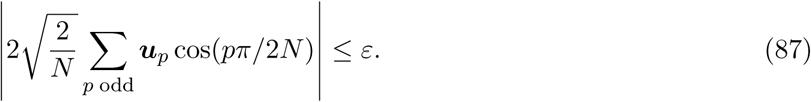

The end-to-end encounter is independent of the center of mass (coordinate *u*_0_ 17). The MFET is the mean first passage time of the (*N* - 1) *d*-dimensional stochastic process

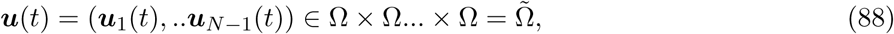

where 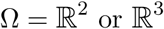 and *u_p_* satisfies the OU-equations 21 to the boundary of the domain

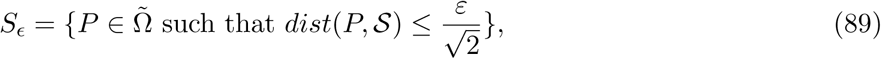

Where *dist* is the Euclidean distance and

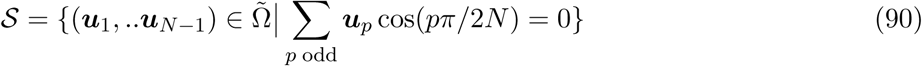

is a submanifold of codimension *d* in 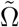. The probability density function (pdf) *p*(*u*(*t*) = *x, t*) satisfies the forward Fokker-Planck equation (FPE) [237]

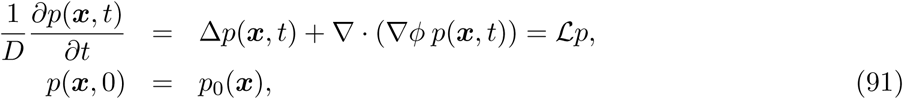

with boundary condition 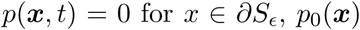 is the initial distribution and the potential 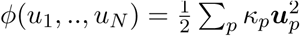 was introduced in 19.

The solution of equation 91 can be expanded in eigenfunctions

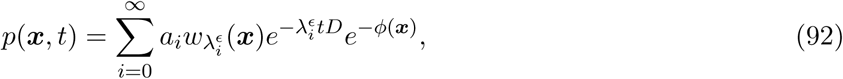

where *a_i_* are coefficients, 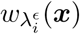 are the eigenfunctions and eigenvalues respectively of the operator 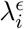 in the domain 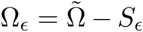. The probability distribution that the two ends have not met before time *t* is the survival probability

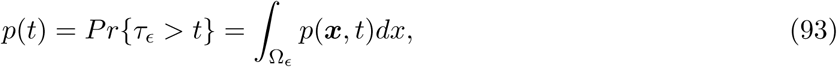

and the first looping time is Using expansion 92, 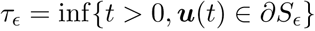 where 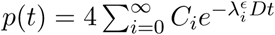 distribution 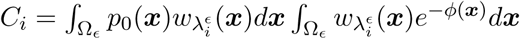

Starting with an equilibrium distribution 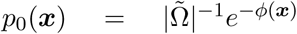 we have 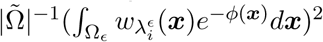 and finally the MFET is given by

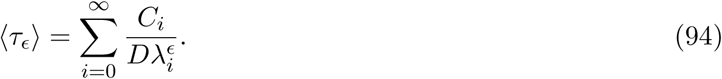

When the polymer distribution is sampled from the equilibrium distribution, we have that the significant contribution is coming from *C*_0_ ≈ 1, while the other terms are *C*_*i*_ = *o*(1). Thus the first eigenvalue is the main contributor of the series.

### 4.5 Computing the eigenvalues of the Fokker-Planck equation and the MFET

The eigenvalues 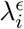of the operator 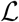 (eq.91) are obtained by solving the forward FPE in 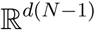, with the absorbing boundary condition on the boundary of the domain *S*_*ϵ*_ (see eq. 89), which is the tubular neighborhood of the (*N* - 1) *d*-dimensional sub-manifold *S*. For small *ε*, the eigenvalues can be computed from a regular expansion near the eigenvalue of the entire space when the small domain *S*_*ϵ*_ is not removed:

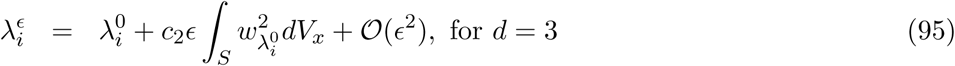

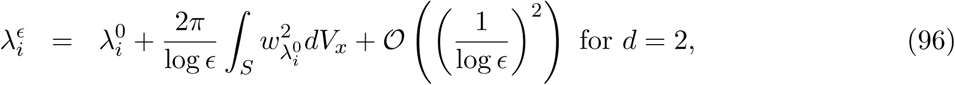

where the eigenfunction 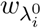 and eigenvalues 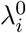 are associated to the non-perturbed operator (no boundary) [46], *d* = 3, 2. In the context of the Rouse polymer, the volume element is 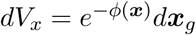 *dx*_*g*_, a measure over the sub-manifold *S* and 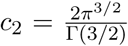[46]. The unperturbed eigenfunctions 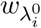 are products of Hermite polynomials [1], that depend on the spatial coordinates and the eigenvalues 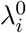 are the sum of one dimensional eigenvalues [13].

The first eigenfunction associated to the zero eigenvalue is the normalized constant

The first eigenvalue for *ε* small is obtained from relation 95 in dimension 3 with 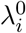

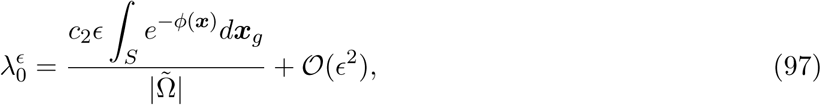

Of the closed loops to all polymer configurations. Using the expression for the potential *ϕ* (de ned in 19), the volume is computed from Gaussian integrals

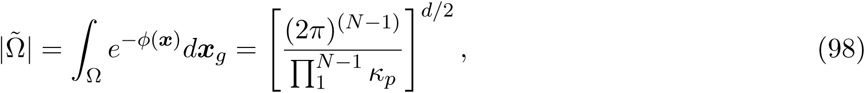

While the parametrization of the constraint 90 leads to [13] after a direct computation

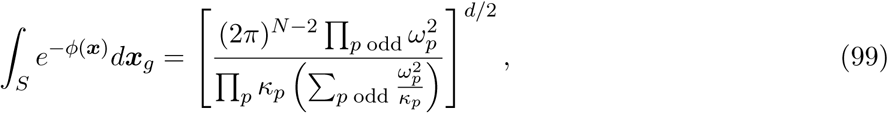

Where *w_p_* = cos(*p*π/2 *N*). We now detail these computations: to evaluate the integration over *S*

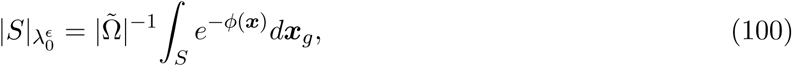

Note that *S* is embedded into 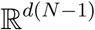 by the immersion (for the detailed computations see [10, 14]),

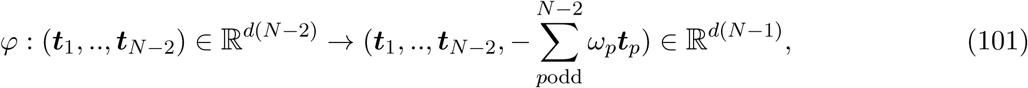

Where

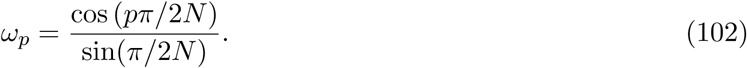

Thus 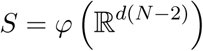is precisely the product of hyperplanes 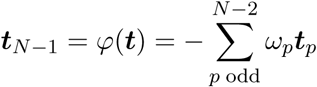The *i*-th element of the normal vector is

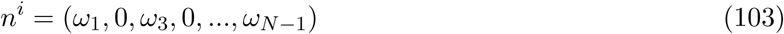

and the normal vector to the submanifold is *n* = *n*^1^ ⊗ *n*^2^… ⊗ *n*^d^. Thus, the metric on *S* is induced by the restriction of the Euclidean distance

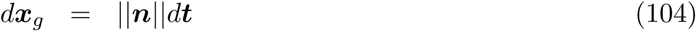

And thus

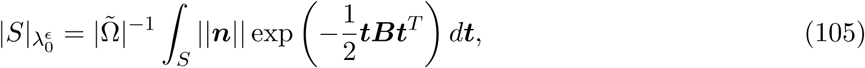

Where *B* is an *d*(*N* - 2) × *d*(*N* - 2) block matrix where the *i*-th block is given by

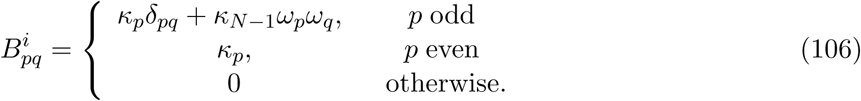

We use that this matrix can be re-written as

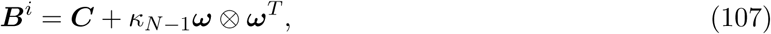

Where *C_pq_* = *κ_p_*δpq and *w* = (*w*_1_, *w*_3_,…, *w*_*N* — 2_). The determinant of *B^i^* is computed using the matrix determinant theorem

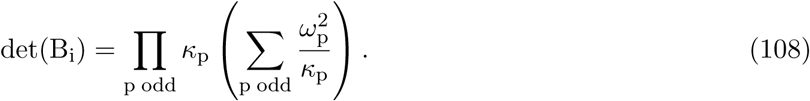

(108)

Finally, we get with

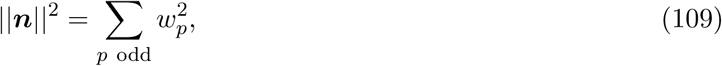

That

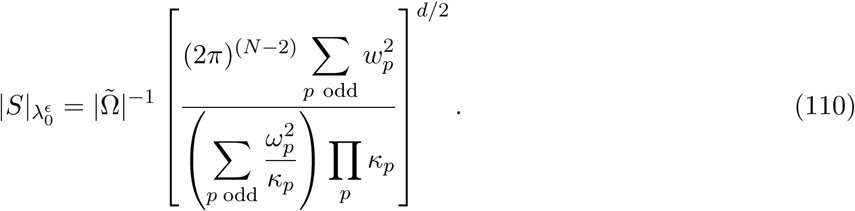

The numerator is

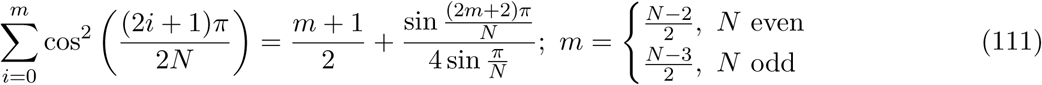

And the denominator

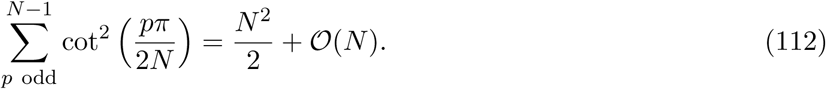

Combining expressions 112 and 111 into equation 110, we get for large *N*

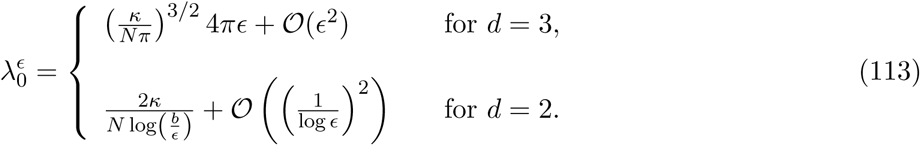

This result shows in dimension three, for small *ε*, the MFET depends linearly on ^1^_*"*_. In summary, for fixed *N* and small *ɛ*,

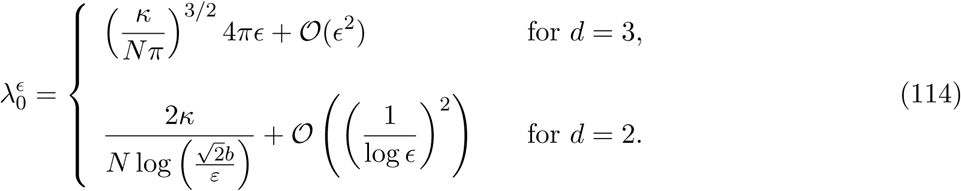

For small *ɛ*, the MFET depends linearly on 
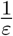 as confirmed by Brownian simulations (Fig. 14e). Because the zero order eigenvalue 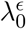 is converging to the zero as *ϵ* tends to zero, it is well separated from the rest of the spectrum. Indeed, the second eigenvalues associated to the first two modes (*p* = 1, 2) are computed from the Hermite eigenfunctions [1], expressed with respect to the coordinates 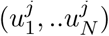 and *j* = 1 *,..d*

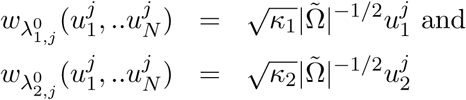

Where 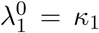 and 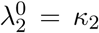 respectively, which are the Rouse first mode (see relation 20). The expansion is [8]

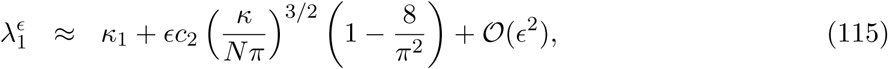

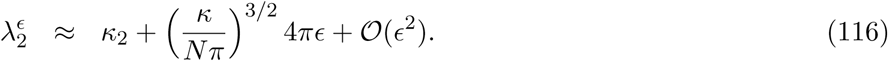

The contribution of 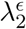 to the first looping time distribution is however quite small as shown in Fig. 14d (small deviation at the beginning), while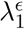 is not negligible. The MFET is well approximated 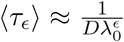 When *N* increases, additional terms are needed in the expansion of 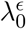, while higher eigenvalues do not contribute signi cantly.

In summary, the zero eigenvalue is sufficient to characterize the MFET, confirming that the FET is almost Poissonian, except for very short time. Moreover, the second term in the expansion of 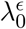 is proportional to 1/ *N* [13]. Using the approximation *C*_0_ ∼ 1 and relation 94, for *d* = 3, the MFET is approximated by

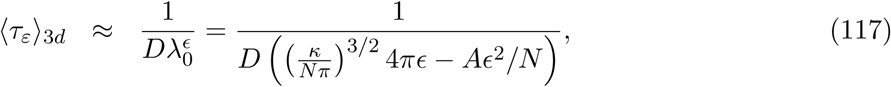

where *A* is a constant that has been estimated numerically. Indeed, with 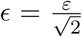 the MFET is for d=3

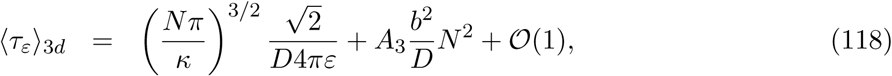

which hold for a large range of *N*, as evaluated with Brownian simulations (Fig. 17). The value of the coefficients are *A*_3_ = 0.053 [13], obtained from fitting, comparable to the coefficient of *N*^2^ obtained in [273], eq.13 (*A*_3_ = 0.053, was estimated from the WF-approximation [287, 288]. These estimates are obtained for fixed *N* and small *ε*. Similar for *d* = 2, a a two dimensional space, the [13]

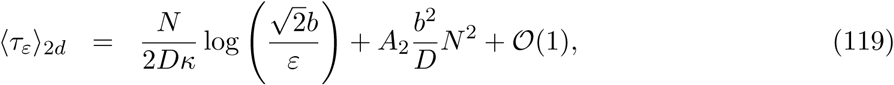

**Fig. 17:**
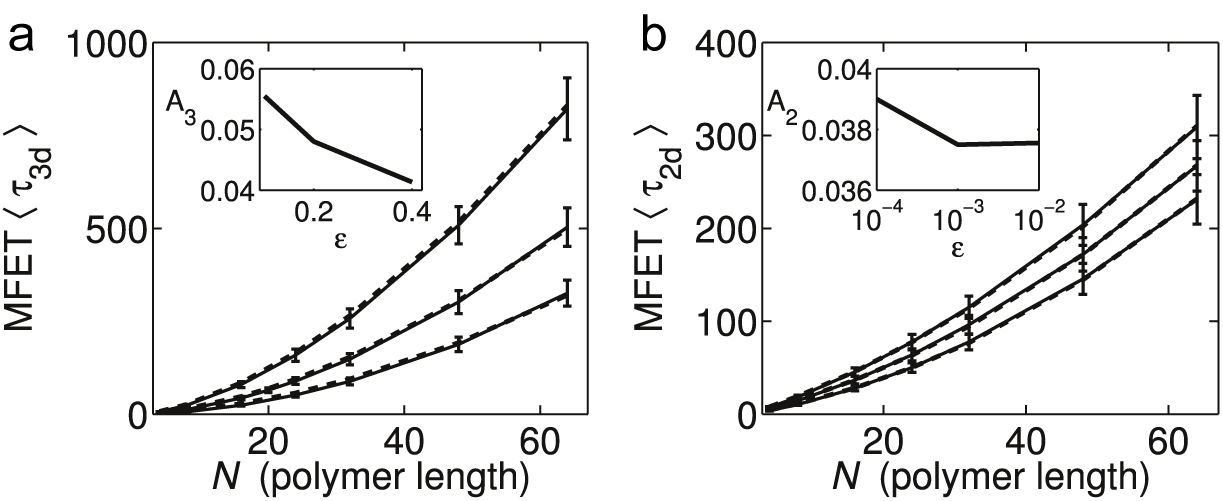
Mean first encounter time for different polymer lengths and different values of *ε*. (a) MFET (three dimensions) estimated from Brownian simulations (full line) and compared to the theoretical MFET (eq.175) (dashed lines). The parameter *A*_3_ is obtained by fitting (*ε* = 0.1, 0.2, 0.4). The unit is in *b*^2^/ *D* (as in fig. 14) (b) The MFET (two dimensions) estimated from Brownian simulation (full lines) and compared to the theoretical MFET (eq.174) (dashed lines). The parameter *A*_2_is obtained by tting (*ε* = 10^-4^, 10^-3^, 10^−2^) (reproduced from [13]).

All these asymptotic expansion are derived for fixed *N* and small *ε*. However, there should not be valid in the limit *N* large, although stochastic simulations (Fig. 17a-b) shows that the range of validity is broader than expected [13].

### 4.6 Looping in a confined domain

The looping time in confined geometries is computed following the same step as in free space, described insection 4.5. The terminology changes and we call this time the mean first encounter time in a confined domain (MFETC). The confined domain can be general but the computations consider the case of a ball. The principle of the computation is indeed to replace the reflecting boundaries for the monomers at the sphere by an external harmonic potential (Fig. 18).

**Fig. 18:**
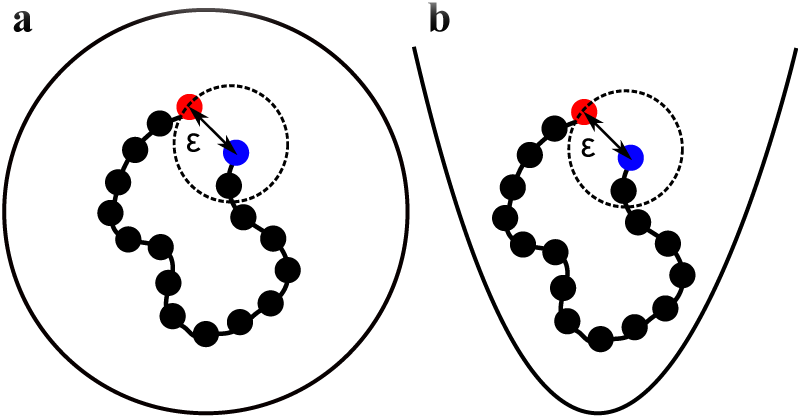
MFETC of polymer ends in a confined domain and in a potential well. (a-b) Schematic representation of the encounter process between the ends of a polymer in a confined ball and in a harmonic potential.

We start with the Rouse polymer [72] containing *N* monomers, with a diffusion constant *D*, located inside a harmonic potential of strength *B*. Monomers are positioned at *R*_n_ (*n* = 1, 2,… *N*). Their Brownian motion is coupled by a spring force to the nearest neighbors (see energy potential relation

5). The energy potential is the sum of the Rouse 5 plus a con ned potential:

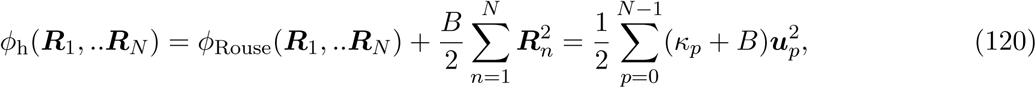

where 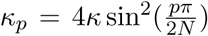 and *u_p_* are defined in 17. The dynamics of monomer *R_n_* for *n* = 1,…, *N* follows the stochastic equations

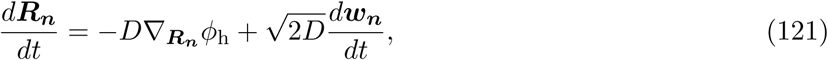

*w_n_* are independent three-dimensional white Gaussian noises with mean zero and variance 1. The MFETC 〈 *τ_e_*〉 is the mean time for the two ends *R_N_*, *R_1_* to enter the distance *ε*, when the polymer evolves according to the new potential *ϕ*_h_. The strength parameter *B* entering into the definition of the potential *ϕ*_h_ is a free parameter, to be calibrated to match a con nement condition [8]: indeed, the parameter *B* is chosen such that the mean square end-to-end distance of the polymer is equal to the radius *A* of the confining ball, that is

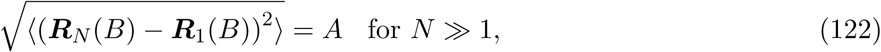

By averaging over the Boltzmann distribution *e*^*-ϕ*^h *dx*, the explicit solution of eq. 122 is [8]

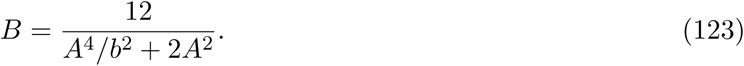

The asymptotic formula for the MFETC 〈τ_h_〉 that two end monomers of a Rouse polymer moving in a harmonic potential meet is [10] for *ɛ* ≪ *b*

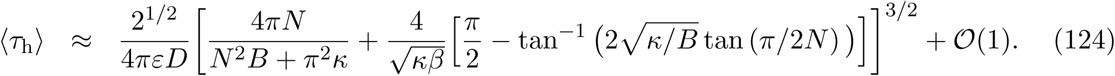

This result is in contrast with previous formula derived for free looping polymers [202,273,82,47,271,254,13] (Fig. 19d): when *N* is large, the MFETC 〈τ_h_〉 converges to a value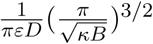. The details asymptotic analysis is presented in the next section.

**Fig. 19:**
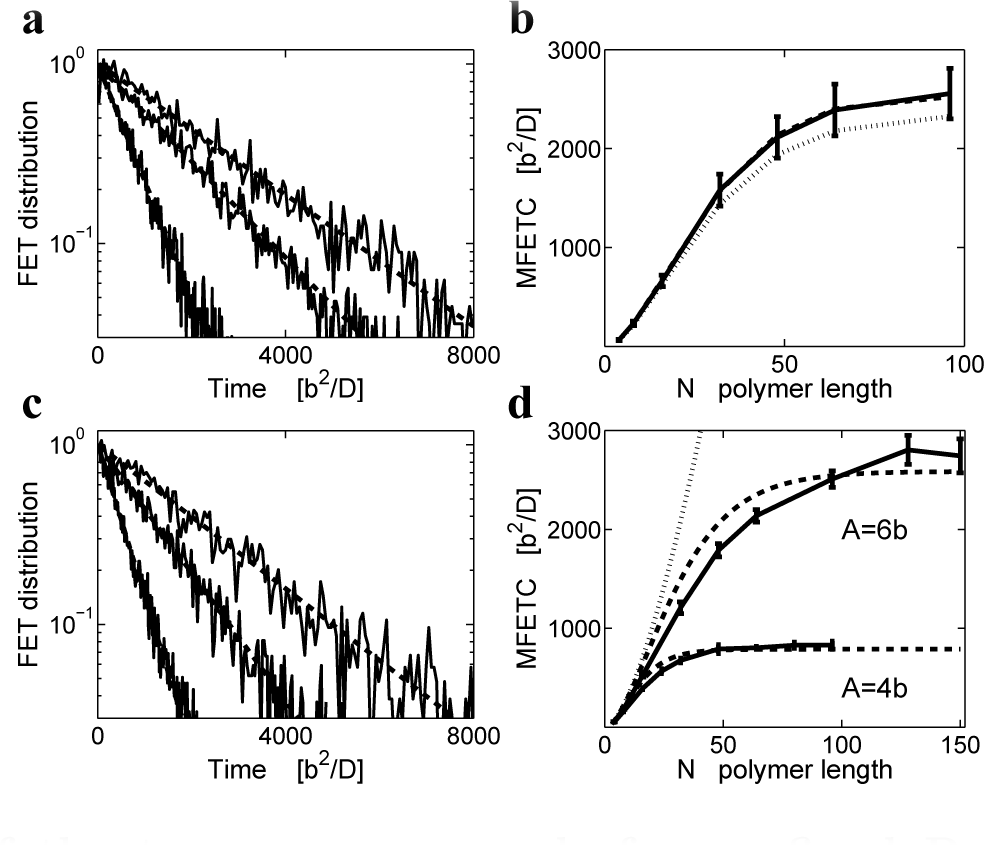
Encounter of the two monomer ends forficon ned Rouse polymer. (a) First encounter time distributions for a Rouse polymer (eq. 14) confined in a harmonic well for various polymer lengths *N* = 16, 32, 64 (left to right) with *B* = 0.01 *b*^2^ and *ε* = 0.01 *b*. A single exponential (dashed line) is enough to describe the process. (b) Brownian simulations (full line) are compared to the MFETC computedeq.128 by taking the rst order only in *ε* (pointed line) and by taking into account the second order correction (dashed line) given byeq.136. The value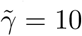 is obtained by tting expression 138 to stochastic simulations. (c) First encounter time distributions for a polymer in a ball of radius *A* = 6 *b*, for various polymer lengths (*N* = 16, 32, 64 left to right). A single exponential (dashed line) is sufficient to approximate the distribution over the entire range. (d) MFETC as a function of the polymer length: Brownian simulations (full lines) in spheres of radii *A* = 4 *b,* 6 *b* with *ɛ* = 0.01 *b*. The MFETC (dashed lines) is estimated by using an harmonic wellapproximationeq.128, where *B* was tted to the simulation results *B*_4_ = 0.0406 *b*^2^ *; B*_6_ = 0.0089 *b*^2^. The MFET is also shown (points) for a freely moving polymer (eq.2 [13]).

### 4.7 Detail computations of the looping time in a confined domain

Formula 182 [13] was derived for a Rouse polymer in confined domain, driven byeq.121 with the potential well *ϕ*_h_ (see 120). The approximation starts with the relation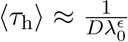 where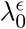 is the first eigenvalue of the operator

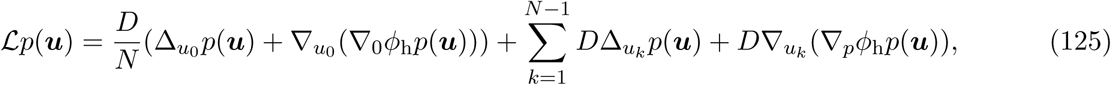

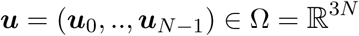 and the absorbing boundary condition is *p*(*u*) = 0 for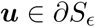 where *S*_*ϵ*_ is the ensemble of closed polymer con gurations

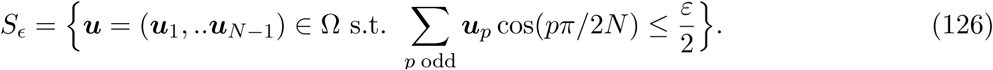

The first eigenvalue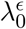 is computed from the perturbation formula 95 (in dimension 3) [46]

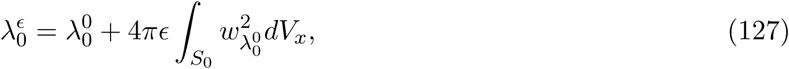

where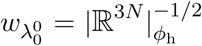 is the constant eigenfunction associated with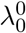 and the volume element is *dV_x_* = e^*-ϕh*^ *dx_g_* and *dx_g_* the Euclidean measure over the sub-manifold *S*_0_ (obtained by taking *ɛ* = 0 ineq.126). With 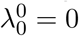 and a direct computation [9] leads to

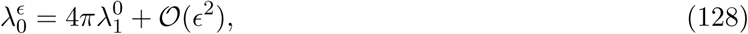

where

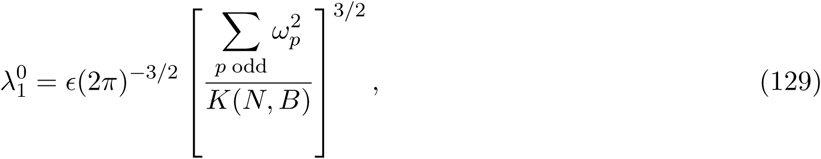

*w_p_* = cos(*p*π/2 *N*) and

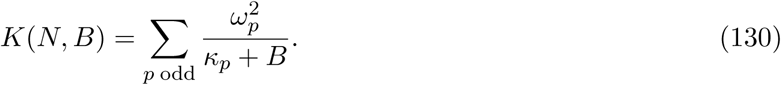

For *N*≫1 relation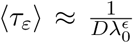 and expanding relation 130 lead toequation 124 [9]. order in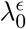 is estimated as follows [14,48]:

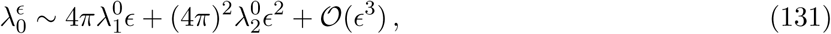

with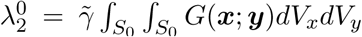 where 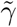 is a constant that depends on the geometry of the boundary. This constant measures the contribution of the second order terpansion of the MFET. The Green’s function *G* is associated to the operator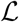 solution of 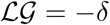 where δ is the Dirac operator. The Green's function is expanded on the eigenfunctions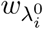 and eigenvalues *λ_i_* of the operator 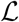 so that

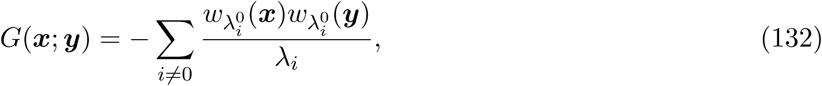

which leads to the approximation [14]

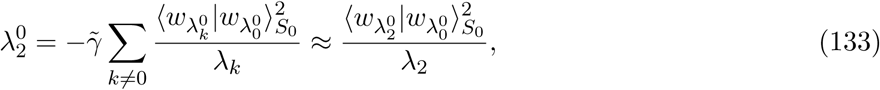

where

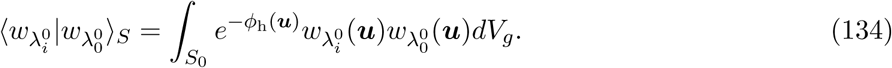

Since the first non-zero eigenfunction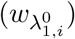 is linear with the coordinates *u*_1 *,i*_, the scalar product is zero and equation 133 is computed from the product associated with the second eigenvalue and *p* = 1 in the spatial directions *i*. The eigenfunctions are given by the second Hermite polynomial

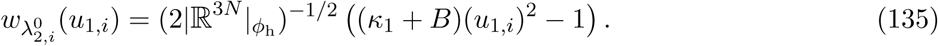

and λ_2_ = 2(*k*_1_ + *B*). A direct computation gives [14]

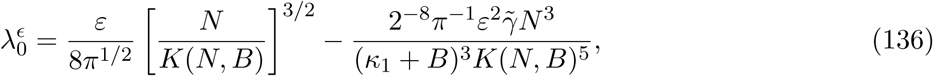

where

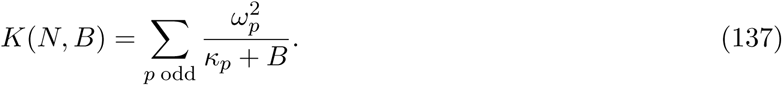

For large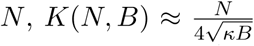 and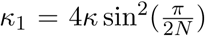. Finally, using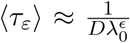 for small *ɛ* and *N* fixed, the approximation 127 is refined, leading to the MFETC expression

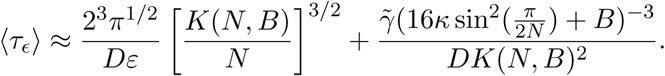

Contrary to the case of a free space (eq. 118) where 〈τ_e_〉 = a_1_N^3/2^/e + *a*_2_ *N*^2^ (a_1_, a_2_ are constants) and the *N*^2^ term dominates for *N* ≫ 1, in the confined case, only the first term is increasing with the length *N* and the two asymptotic limit is bounded by the diameter of the domain.

The looping time distribution in con ned domains is Poissonian. It is also the survival probability *P*(*t*) that a loop is not formed before time *t* (see section 4.4). In this approximation,

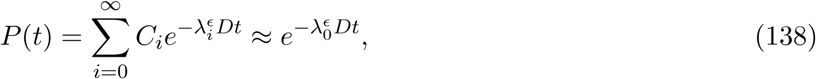

where *C_i_* are coefficients. Brownian simulations of the two ends of a polymer in a harmonic well (Fig. 19a) confirms this Poissonian approximation. Unlike the FET distribution for a free polymer, *P*(*t*) is well approximated by a single exponential, even for long polymers, showing that the higher exponential terms (eq.138) do not contribute.

In summary, the encounter time for a polymer in a harmonic well is Poissonian. Moreover, as observed from matching the theoretical formula 182 and Brownian simulations for the MFETC in a potential well (Fig. 19b), the first order correction in *ε* is enough and this approximation remains valid for large *N* and small *ε*. The true formula for a bounded domain, without using the parabolic confinement is still to be found.

### 4.8 Brownian simulations of looping in a con ned domain

The range of validity of formula 182 for a polymer looping in a con ned domain can be estimated using Brownian simulations of a Rouse polymer in a ball (Fig. 19d) for different radii. Although for small polymers, there is almost no differences between confined and non-confined looping time, for long polymers, the MFETC reaches an asymptotic value that depends on the radius of the domain. When the interacting monomers are not at the ends of polymer chain, the MFETC is reduced in a range between five to thirty percent, due to the interactions of the additional monomers with the boundary [8].

The distribution of arrival times (Fig. 19c) for short and long polymers is well approximated by a single exponential, confirming that the looping event is almost Poissonian. This is in contrast with looping in free space, where for longer polymers, a second exponential is necessary. The MFETC is computed by taking into account only the first order terms in *ɛ* (seeeq.128). The strength *B* is estimated by comparing the analytical formula with Brownian simulations: the calibration formula 123 gives similar values to the fitted ones. For *A* = 4 *b,* 6 *b*, the fitted values are *B*_fit,4_ = 0.0406b^-2^, *B*_fit,6_ = 0.0089b^-2^ respectively, while the calibration formula 123 gives *B*_cal,4_ = 0.0417b^-2^, *B*_cal,6_ = 0.0088b^-2^. Using the fitting procedure (formula 123), it is possible to estimate the radius of a confined sphere. The MFETC has different applications such as estimating the chromosomal interaction forces, the nuclear sub-organization [157,110] and the encounter between two sites that can initiate gene activation or regulation [66,209,76].

In summary, the FLT is well approximated by a single exponential, showing that the associated stochastic process is almost Poissonian. Consequently, looping in the nucleus, chromosome or telomere encounter can be well characterized by the mean looping time. This approximation simplifies heavy numerical simulations or reduce analysis to Poissonian processes, such the analysis of telomere clustering in the yeast nucleus [122]. Another conclusion is that by increasing the radius *ε* or the polymer length *N*, the asymptotic formula for the MFET is completely determined by the first eigenvalue, but not by higher order eigenvalues. Two scales are involved in the MFET, one proportional to *N*^2^ and the other to *N*^3/2^ and both are already contained in the first eigenvalue and do not arise from higher ones. It is surprising that the regular perturbation of the Fokker-Planck operator in *ε* (formula 95) introduces a novel scale with *N* in all eigenvalues. A complete expansion for the MFET has to be found and it would be interesting to find a geometrical characterization of the constants *A*_2_ and *A*_3_.

### 4.9 Looping formula for rod-like long polymer

In this subsection, we analyze a different polymer model called the rod-like model (introduced in paragraph 3.9) which is more elementary than Rouse, but this model is not really used for describing looping. The reason is that this model shows correlation of the last monomer with the rest of polymer chain (see [12] for a complete discussion). However, the analysis of this model is quite simple and contains interesting asymptotic features.

We shall now describe the MFET computation for the rod-like polymer. The analysis starts with approximating the stochastic process of the position *R*_*N*_ (*t*) in the large *N* limit in dimension 2 [12]. Ito's formula for the exponential of the Brownian angle *θ_k_* gives

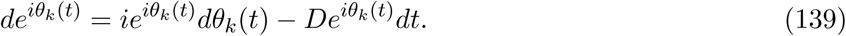

By summing relations 139, using relation 41 for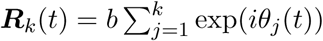 we get

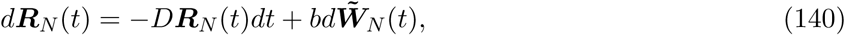

where the new noise term is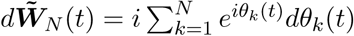The noise source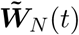 can be approximated by a Brownian motion for large *N* in the following sense: 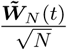 converges asymptotically (in probability) to a Brownian motion [278]. While therst moment of *W*_*N*_ is zero, the second order is computed from the decomposition 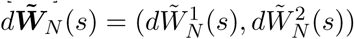 where

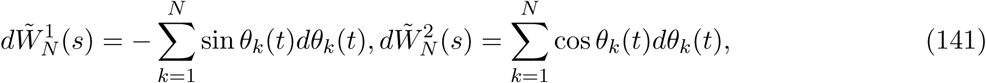

are derived from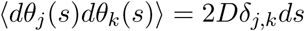 Indeed,

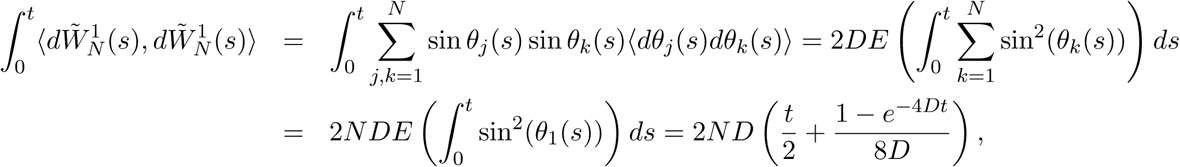

because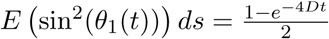 Similarly,

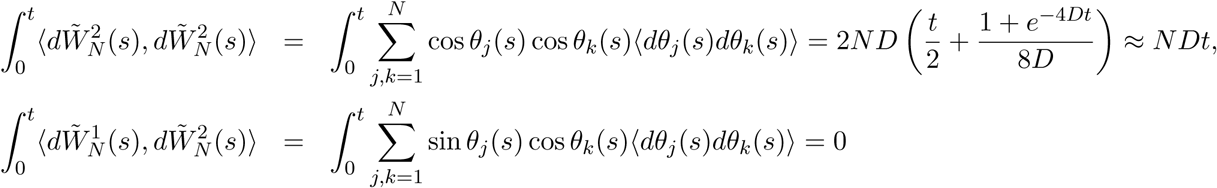

For large times, the leading order term for the second order moments is *N Dt*. Thus to compute the MFET of the end-points of the long polymer, the noise term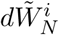 was approximated by a Brownian motion in both x and y-component, with a diffusion constant *ND* and the stochastic equation 140 is approximated by the following Ornstein-Ulhenbeck process

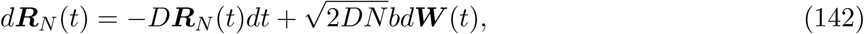

where***W*** is a two-dimensional Brownian motion of variance one. The MFET can be computed by considering the dynamics inside a ring with exterior and interior radii *Nb* and *ε* respectively. The associated partial differential equation is described in 144, where the inner boundary *r* = *ε* is absorbing, while a Neumann boundary condition is imposed on the external boundary, the looping time *u*(*r*), where initially | *R_N_*| = *r* is solution of the partial differential equations

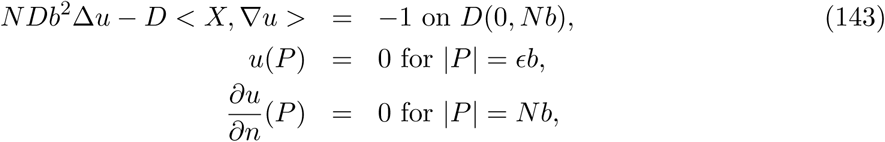

where *X* is the radial vector, *D*(0 *; Nb*) is the disk centered at the origin of radius *R* = *Nb*. The equation is solved in polar coordinates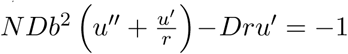 and the solution is

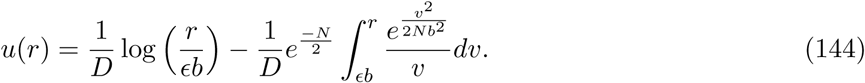

Interestingly, when the initial configuration of the rod-polymer is a straight segment (*r* = *R*), the MFLT is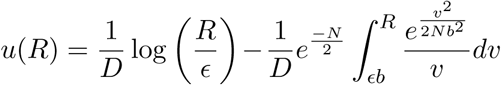 Integrating by parts and using Laplace's method for *N* large, the MFET is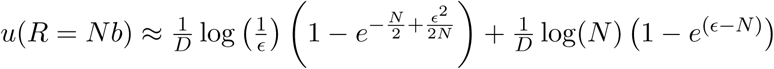 Finally,

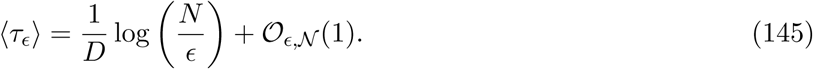

Three regimes can be distinguished depending whether *N* is ≫ ≪ or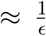. when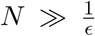, the dominant behavior depends on log(*N*). The MFLT for a chain starting in a uniform distribution is

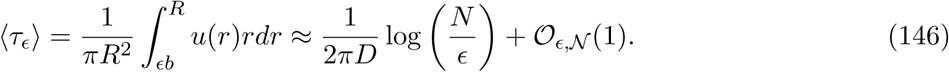

This asymptotic formula is different from the one obtained for a chain initially straight (formula 145). The leading order term depends on the initial configuration of the polymer. This is quite surprising because the prediction of the narrow escape formula for a Brownian particle implies that the leading order term does not depend on the initial configuration [239]. In summary, the initial configuration for a rod-like polymer affects the leading order asymptotic expansion of the MFLT, suggesting that the underlying stochastic process is not Markovian and has a long memory, a phenomena that should be further investigated.

When the initial condition is chosen at equilibrium with a distribution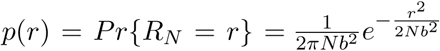 then the MFLT is given by

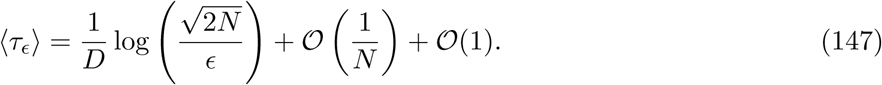

The leading order term of the MFLT depends again on the initial configuration and should be compared with formula 146.

### 4.10 Looping time for a 4^*th*^ -order stiff polymer using a non-Markovian approach

The dynamics of a single monomer from a polymer chain and the mean first looping time was considered using Brownian simulations using a non-Markovian approach [99,100]. In that approach, the parameter of the model are the length (the number of monomer *N*), the same diffusion coefficient *D* for each monomer, the bond length *l*_0_, a target size *a* such that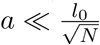 inside a domain of volume *V.* In that case, the leading order term for the mean time recover the classical result of a search to a small target

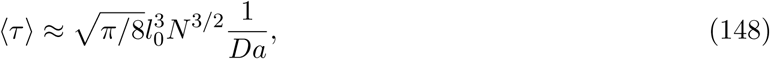

while for 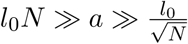

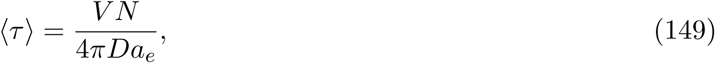

where the effective radius is defined by 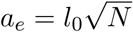 and does not depend on the target size (when *a* is very small). The general formula is derived insection 4.6.

Interestingly, the mean looping time 〈τ〉 for a semiflexible Gaussian chain can be estimated in a polymer model, where the local interactions tends to align successive bonds. A continuous model is discussed in [107,90] for a stiff polymer. The tension along the chain is approximated by a constant *g*, neglecting the long-range hydrodynamic, *b*_0_ is the thickness of the chain, is the bending rigidity. The equation of motion for a segment *r*(*s, t*) located at position *s* at time *t* is

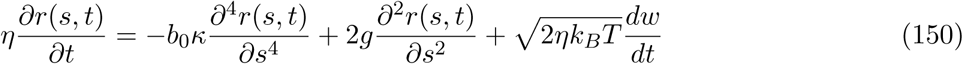

where η is the friction per unit length and *w* the classical normalized Gaussian noise in time and delta-correlated in space. Four reflective boundary conditions are imposed at the boundary of the chain [101]. The evolution of the position can be computed in the Fourier space from an ensemble of OU-processes as follow

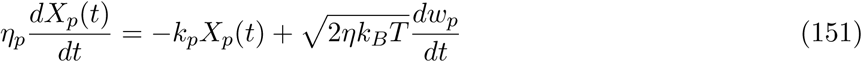

for *p* ≠ 0 *k*_*p*_ = *ap*^4^ + *bp*^2^, *a* = 2κb_0_(π/ *L*)^4^, *b* = 4 *g*(π/ *L*)^2^, *w*_*p*_ are independent normalized Brownian motion, with

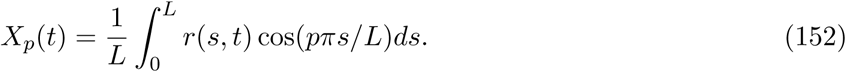

The looping time of a semiflexible chain when the smallest length scale is the bending fluctuation length and the target size is *a* ≪1 has been estimated asymptotically

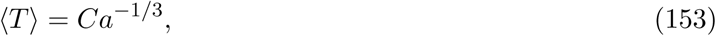

where *C* is constant [101]. This scaling law is a characterization of looping for a stiff polymer. The regime where κ≫ *l*_0_ *N* should be very different from the one discussed in this section and the asymptotic analysis should be clari ed.

### 4.11 Mean First Encounter Time (MFET) between two monomers located on the same polymer

The MFET between two monomers of a β-polymer (see section 3.5) is computed from the first eigenvalue expansion of the associated Fokker-Planck operator. We proceed now following the same steps as in the previous paragraph, neglecting the contributions of all higher eigenvalues, thus

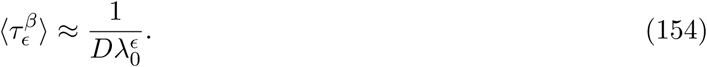

To estimate 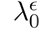 to first order in ɛ, we use formula [14]

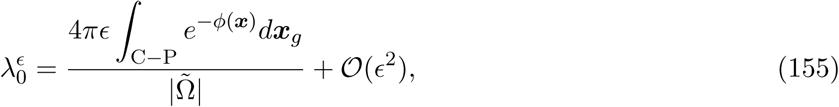

which is the ratio of a Gaussian integrals over all closed polymer (C-P) to the whole polymer configuration ensemble. Integral 155 is computed directly. Indeed, using the expression for the potential 25, we get

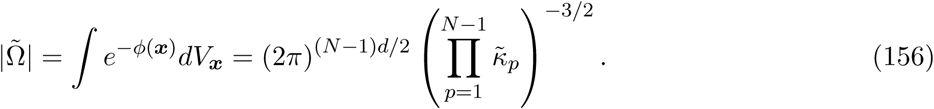

The numerator ofequation 155 involves integrating over the ensemble of closed polymer loops

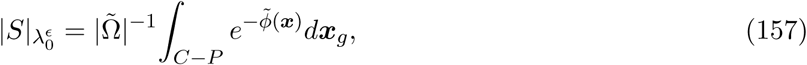

where *dx*_*g*_ is the induced metric of the configuration space onto the set C-P. A direct integration of eq. 157 gives (see paragraph 4.5) [14]

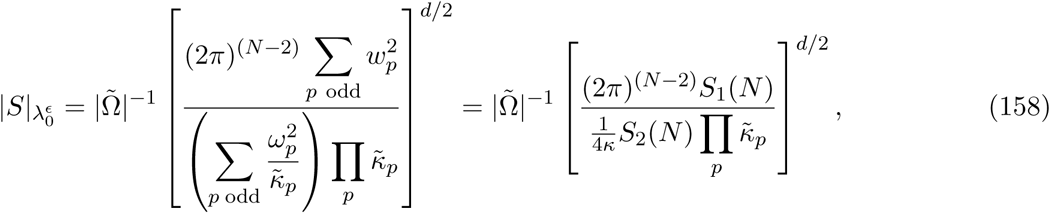

where ω_*p*_ = cos (*pπ /*2 *N*) and the series

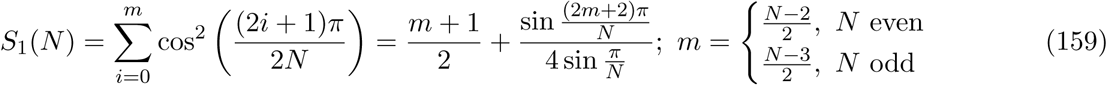

and

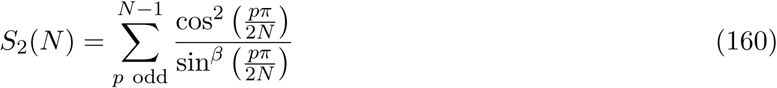

is approximated by the Euler-Maclaurin formula by

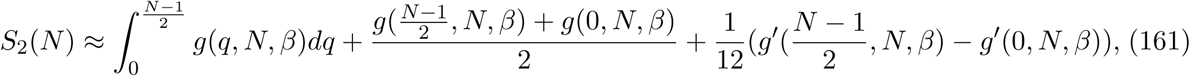

where 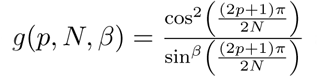 and the integral is

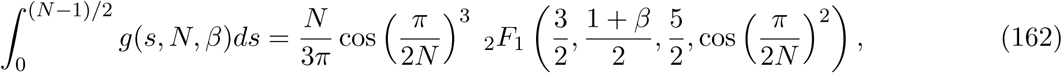

Where_2_ *F*_1_ is the Gaussian hypergeometric function [1]. The terms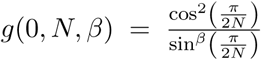 and 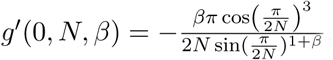 in 161 are of order *O*(*N*^β^). The series *S*_2_(*N*) is approximated by

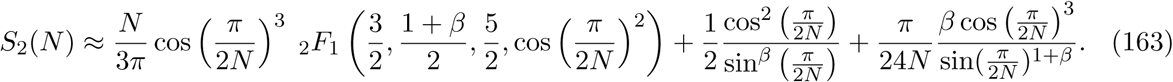

Thus substituting relations 163,159, 158 and 156 intoeq. 155, we get for large *N*,

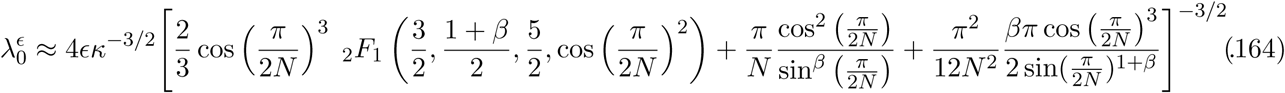

For large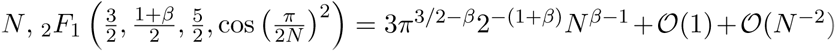, while the two other terms in the parenthesis scale like *N*^1^. Finally, the mean encounter time for the two end monomers is

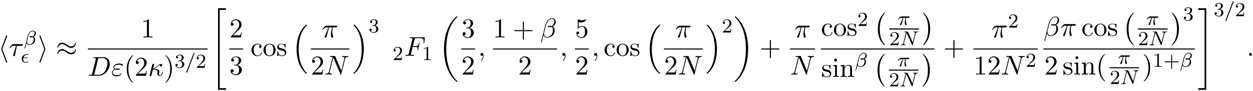

In summary, the MFETs are quite different for a Rouse or a β-polymer: for the latter, it scales with *N*^3/2^, while for a β-polymer in the limit *N* ≫1, it behaves like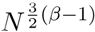.

### 4.12 Search for a small target located on a boundary

In previous sections, we studied the looping time and we now focus on the search process by single monomer for a small target located on the boundary of a ball (nucleus). In a biological context, targets can be small nuclear pores located on the nuclear envelope or active genes that need to be found and activated [5]. Finding a small nuclear pore can also arise in the context of gene delivery, where RNA fragments have to enter the nucleus pores. During the repair of a double-stranded DNA repair, a DNA end has to search for its other broken end in the con ned microdomain generated by the chromatin environment [165] and can also re-localize to the membrane periphery to interact with a nuclear pore.

As there are no analytical formula available for this search process, we present several coarse-grained numerical results of the first arrival time of a monomer from polymer to a small target, called the narrow encounter time for a polymer NETP. In that case, the small target is located on the surface of a bounded microdomain. The polymer model we consider here is a Freely-Joint-Chain (monomers connected by springs with a none-zero resting length *l*_0_).

There are several possibilities to state the search problem:

1. Any one of the monomer or
2. Only one

can find the small target. Simulations reveal that the NETP is an increasing function of the polymer length until a critical length is reached, passed this length, it decreases (see Fig. 20). Interestingly, for the second case, the position of the searching monomer along the polymer that can be absorbed strongly in uences the NETP. Computing the NETP is relevant because it is the reciprocal of the forward activation rate for diffusion limited processes of chemical reactions for a site located on a polymer [299,202]. It may also be used to estimate the first time that a gene is activated by a factor located on the DNA, which is different from the classical activation due to a transcription factor [251,168,24,215]. However, at this stage a final satisfactory analytical formula is still missing (see [117] for an attempt).

**Fig. 20.**
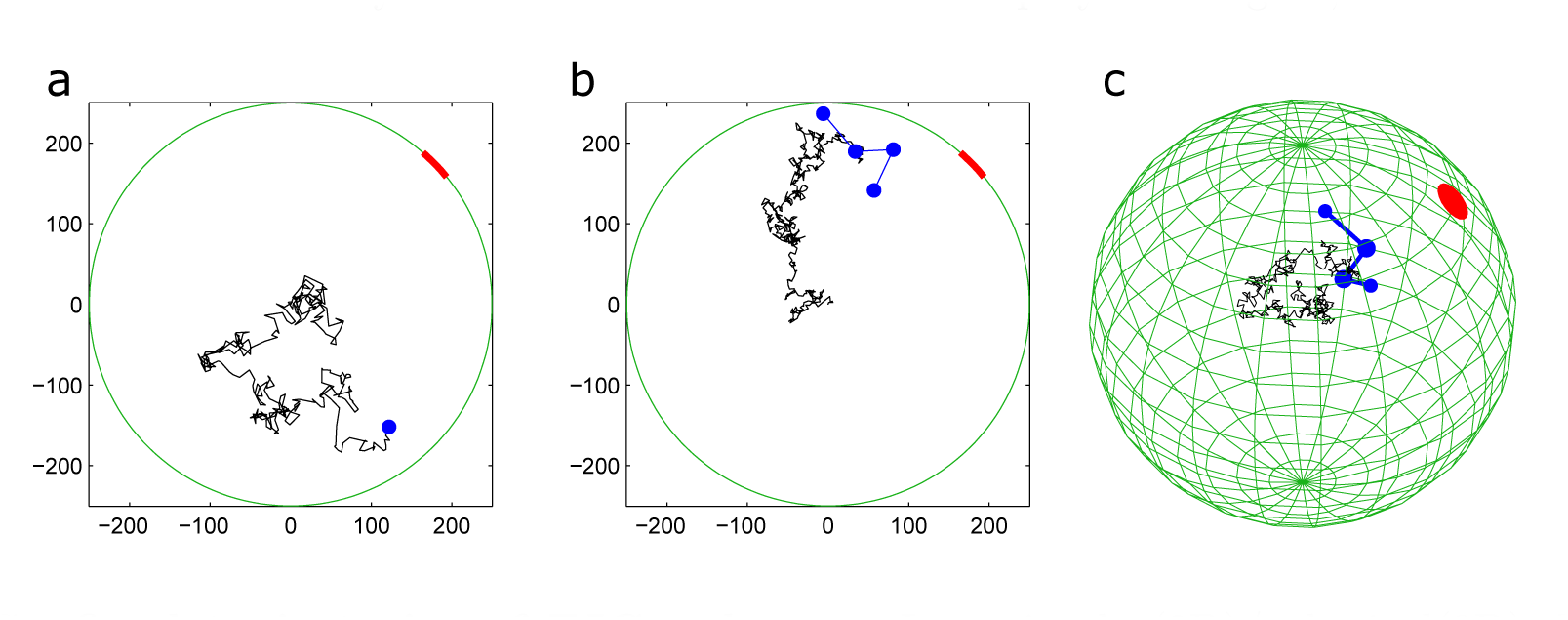
Confined trajectories of FJC polymer. In a circular(2D)/spherical(3D) domain of radius 250 nm, the target size is *ɛ* = 50nm. Trajectories of the center of mass (black) before absorption at the target (red). (a) Simulation with one monomer, moving in two dimensions. (b-c) Simulations of a 4-bead Rouse polymer, moving in 2 dimensions (b) and in 3 dimensions (c).

We now recall some results of Brownian simulations for the two cases mentioned above:

1. When *any* of the polymer monomer can be absorbed at the target region, the search time is designated by 〈τ_any_〉.
2. When only one xed monomer can nd the target region, it is 〈τ_mon_〉.

The key parameters that influence the search time are the polymer length, the monomer’s position along the chain and the bending elasticity [8]. In dimensions two and three, the NETP increasing with the polymer length (number of monomers *N*), until a critical value *N*_*c*_ after which it is decreasing. The radius of gyration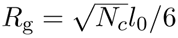 (where *l*0 is de ned as the shortest length between the nearest monomers at equilibrium < | *x*_*k*_- *x*_*k*+1_| >= l_0_) can be used to characterize the coupling between the polymer and the boundary of the microdomain. For short polymer lengths, such that *R*_g_ ≪ 2 *R* (diameter of the microdomain), the NETP is largely determined by the motion of the center of mass. When the polymer is far from the absorbing target, none of the monomers will be able to reach it, until the center of mass has moved close to the target. The NETP thus reflects the mean first passage time of the center of mass to the target. In this limiting case, the center of mass undergoes Brownian motion with a diffusion constant inversely proportional to the number of monomers: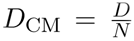. Thus, in the regime *N*≪ *N*_*c*_, the NETP is approximately that of a single Brownian particle, but with a smaller diffusion constant. The expressions for the NET are inversely proportional to *D*_CM_, and thus we obtain the initial linear regime in *N*:

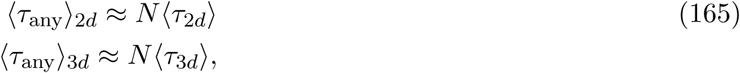

as con rmed by numerical simulations inFig. 21. However for a polymer of length comparable to the size of the microdomain, the location of the center of mass does not determine anymore the NETP. In that regime, smaller subsections of the polymer can be close to the target even if the center of mass is far away. In addition, when any bead can be absorbed, increasing the polymer length results in a decrease in the NETP (Fig. 21). Interestingly, two regimes can be further distinguished for the decay phase. When any bead can be absorbed, the decay phase of the NEPT (Fig 21) can be separated into two different regimes that can be described as followed. In the first one, the polymer moves freely until a monomer hits the absorbing boundary. The NETP is determined by the competitive effects of a decreased diffusion constant for the center of mass and an increased total polymer length. Increasing the polymer length leads to an effective smaller volume of the effective con ning domain in which the polymer has to find the absorbing window.

**Fig. 21.**
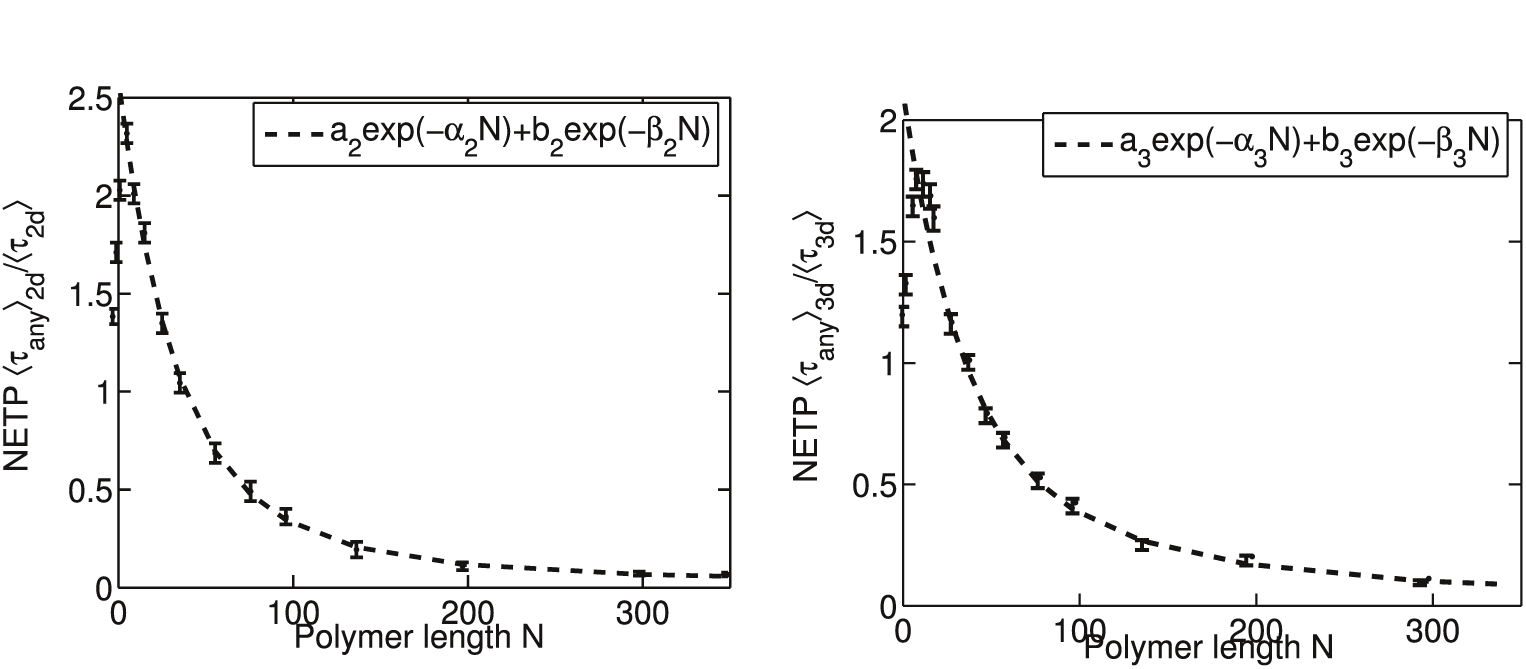
NETP of a monomer to the target for various polymer lengths. Mean time that any one of the monomer to reach a small target located on the boundary (normalized to the NET_0_ for a single monomer) in two and three dimensions (a) and (b). Each point is an average over 2000 runs. Brownian simulations are fitted by a double exponential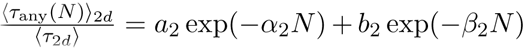 with exponents α_2_ = 0.0075, β_2_ = 0.024 and coefficients *a*_2_ = 0.23, *b*_2_ = 2.17 and in dimension 3 by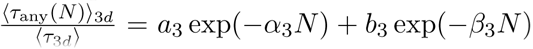 with exponents α_3_ = 0.0082, β_3_ = 0.030 and coefficient *a*_3_= 0.60; *b*_3_= 1.56. (reproduced from [8])

When the length of the polymer becomes long enough, so that at least one monomer can always be found in the boundary layer (of size *ε*) of the absorbing hole [239], the NETP has a different decay as a function of *N* compared to the previous intermediate regime. In that case, the center of mass is strongly restricted due to the interaction of all the monomers with the microdomain surface (Fig. 22). The NETP is determined by mean time for a monomer in the boundary layer of the absorbing window. It is still an unsolved problem to obtain an asymptotic estimate for that time. A fitting procedure shows that the NETP scale with two exponentials

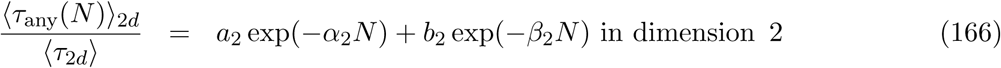

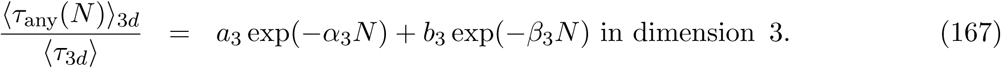

〈τ_3 *d*_〉 and 〈τ_2 *d*_〉 are the NET for a single particle τ_2_, in dimension 3 and 2 respectively. In dimension two, the exponents are α_2_ = 0.0075, β_2_ = 0.024 and coefficients *a*_2_ = 0.23, *b*_2_ = 2.17 and in dimension 3, α_3_ = 0.0082, β_3_ = 0.030 and coefficient *a*_3_ = 0.60, *b*_3_ = 1.56 (Fig. 21). The empirical laws 166-167 have not yet been derived analytically.

The bell shape nature of the NETP can be qualitatively explained using the NET equations: indeed, a small polymer can be considered as a quasi-particle of radius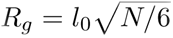 [57], evolving in an effective domain which is the full domain minus its volume. Thus the NETP is related to the mean first passage time of the quasi-particle with diffusion constant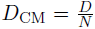 in the apparent domain of volume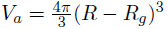 leading to a mean time proportional

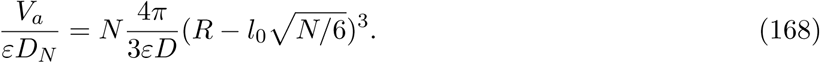

**Fig. 22.**
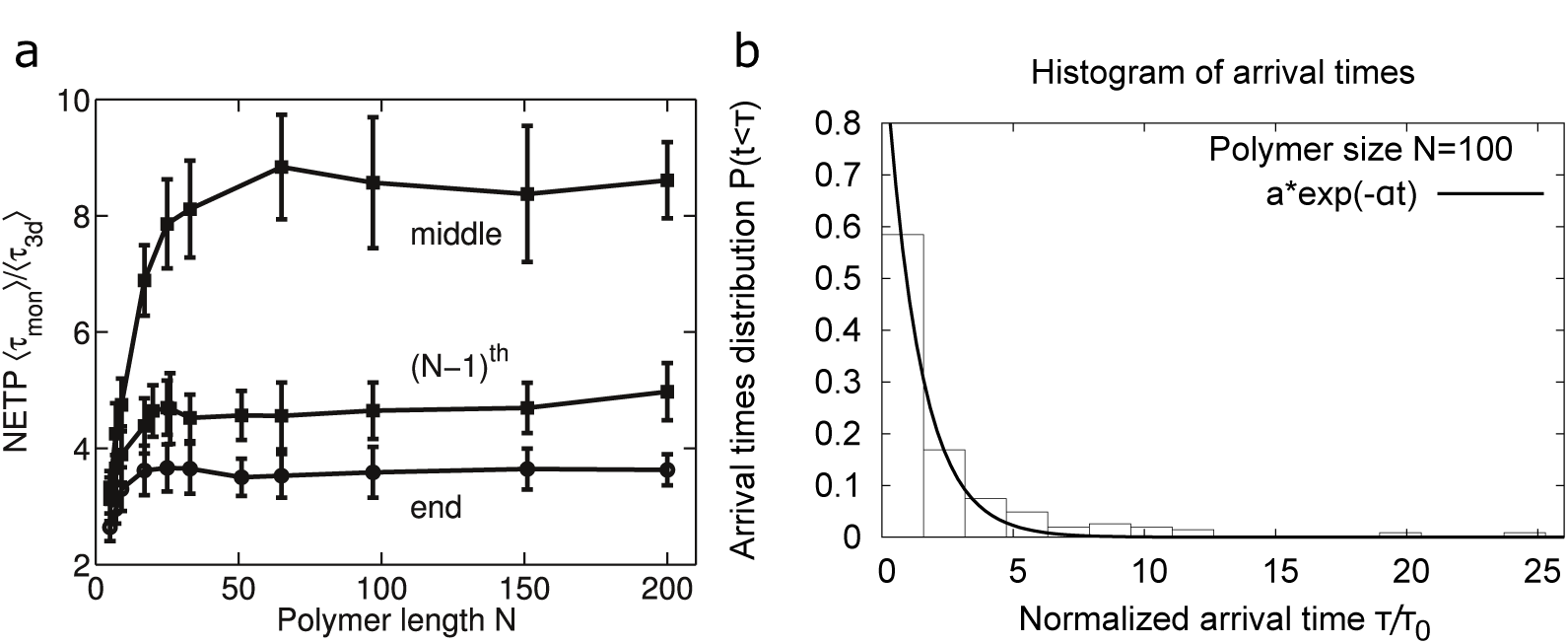
Mean search time of a single monomer (NETP) to a small hole located in a cavity (see also gure 3). (a) NETP for three different monomers: end, middle and N-1: The NETP is plotted as a function of the polymer length *N* in three dimensions (Brownian simulations): The encounter time is normalized to the time τ_0_ (for a single Brownian particle). Parameters are described intable 2. (b) Probability distribution *P* [τ/τ_0_] of arrival times for the end monomer to a small target (in three dimensions). The Probability distribution of the arrival times to a small hole located on the boundary of a sphere in three dimensions is well approximated by a single exponential *Pr*{τ_3 *d*_= *t*}= *a* exp(-λ *t*) with *a* = 1.014, λ= 0.76 (reproduced from [8]).

This phenomenological formula shows that the mean time has a maximum for *N*_*m*_ = 25, which is an over estimation of the empirical value that we obtained from Brownian simulation *N* = 10.

The case when only one monomer can find the target is quite different from the situation where they can all find it: indeed the monomer location along the polymer chain influences the result of the NETP, as shown in Fig. 22a. When a polymer model is con ned in a ball and only one monomer can be absorbed and all others are reflected at the target site, numerical simulations reveal that between the middle and the end monomer, there is a factor 3 reduction in the arrival time (Fig. 22a). Interestingly, already taking the *N*-1th monomer compared to the last one is making a noticeable difference. In addition, the NETP increases to a different plateau that depends also on the monomer location. During the increasing phase, the arrival of the monomer to the target is mainly governed by the diffusion of the polymer center of mass, which can be approximated by a diffusing ball of diffusion constant *D*_CM_ = *D/N*. However, for larger *N*, this approximation is not valid, rather the arrival time converges to a constant value that depends on the stochastic dynamics, which is the one of a correlated particle (monomer) to a target. Complementary results are discussed in [100].

Finally, we have shown in Fig. 22b, that the histogram of arrival time of any monomer to a small target can well be approximated by a single exponential, suggesting that the arrival time is almost Poissonian. This situation is similar to the arrival time of a single monomer to a small target located on the wall of a cavity, as shown in Fig. 19a and c. In both cases (search for a small target by a single monomer for for one end by the other end of a polymer in a con ned domain), the NET converges asymptotically to a value in the large polymer length limit(Fig. 19b and d and Fig. 22a). The most striking difference is due to the apparent potential well generated by the reflections of the monomers on the boundary that increases the search time, which is not the case for Rouse polymers. But in both cases, the Poissonian statistics reflects that the searches for a small target is a rare event, characterized by long time asymptotics [239].

### 4.13 Induced screening potential on a monomer by the other ones at a boundary

The interaction of a monomer with other monomers generates an effective potential through their interaction with the boundary, which is different for the middle and the end monomer, leading to the major differences in the searching time, as shown in Fig. 22a. For *N* large enough, the middle monomer is more con ned than the end one, and by analogy with the diffusing of a stochastic particle in a potential well, the middle monomer has to surmount a higher potential barrier to reach the small target located on the boundary. Indeed, a crude approximation consists in considering that the active monomer motion with position ***X***(*t*) follow a diffusion process in a spherical symmetrical potential *V* inside a con ned spherical domain,

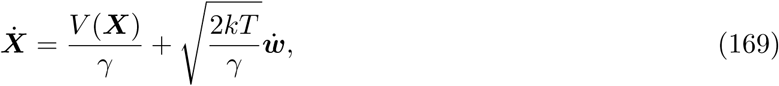

where *w* is the standard Brownian motion. Using the symmetry of the domain, the potential *V* has a single minimum at the center. In a high potential barrier approximation, the mean time to a small target does not depend on the specific shape of the potential, but rather on its minimum and maximum [248]. In that case, the mean time τ(*N*) to a target, which depends on the polymer size is given by:

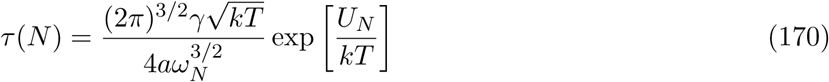

where *a* is the size of the small target, *U*_*N*_ (*r*) is the energy potential generated by the interaction between the polymer and the boundary of the domain at the target site. The key parameter is the frequency ω_*N*_ at the minimum of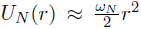 near 0 [248]. The potential *U*_*N*_ (*r*) can be recovered from formula 170 and it is approximated by the numerical simulations described in Fig. 23c. Interestingly, depending on their position along the polymer chain, monomers have also different three dimensional spatial distribution inside a ball as revealed by numerical simulations (see Fig. 23a, b). For example, the middle monomer is more restricted to the center compared to the end monomer, which explores in average a larger area. When the length of the chain increases, the middle monomer position become more restricted, while the end-one does not seem to be much affected (Fig. 23a). The pdf of the center of mass position is comparable to the one of the middle monomer. Today, there are no analytical formula for the mean distance of a monomer to the boundary of a bounded. The extrusion of the end monomer from the center is probably a consequence of a single spring force at the end, compared to two for other polymers.

**Fig. 23.**
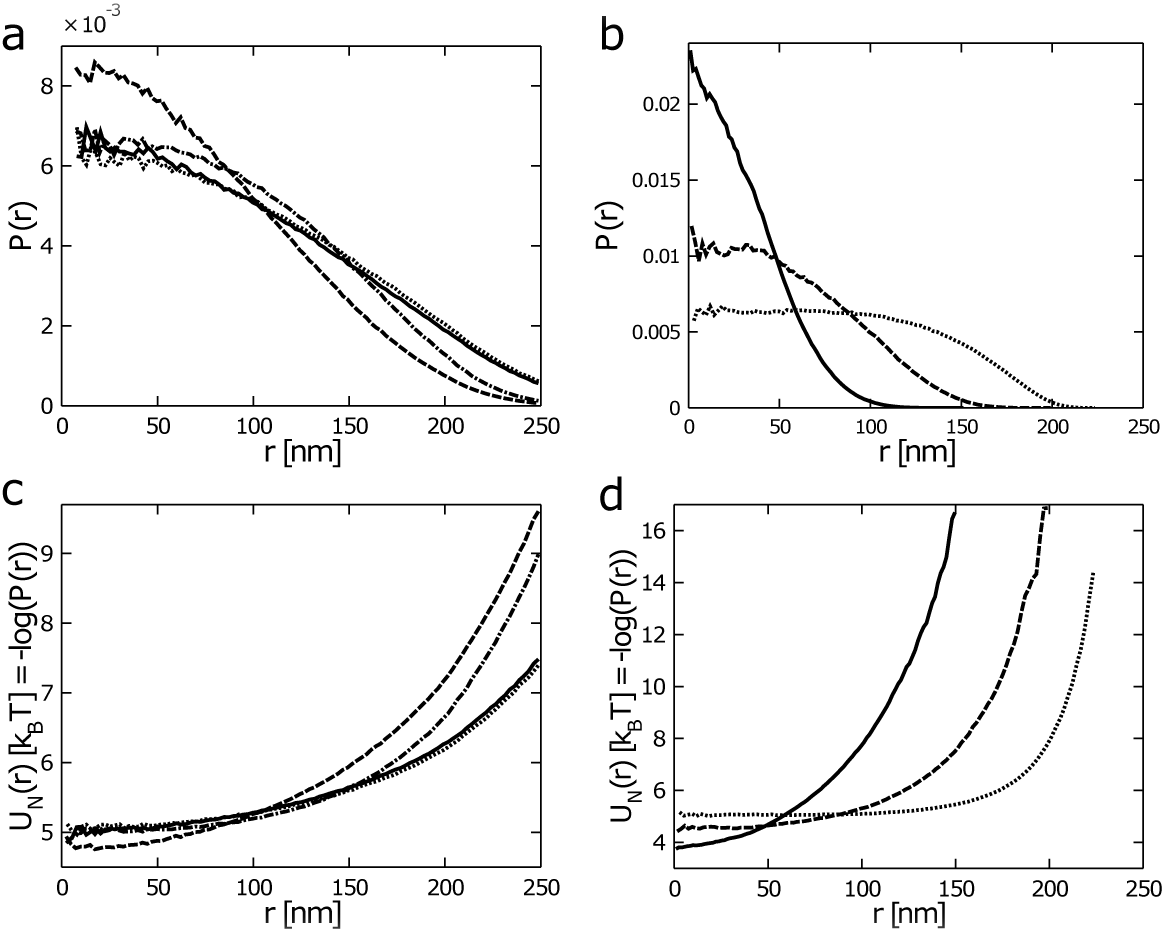
Distribution of monomer position and the in uence of the boundary. (a) Probability distribution function (pdf) of monomer positions. The radial pdf is computed for the end *N* = 16 (points), *N* = 150 (full line), the middle *N* = 16 (points-line) and *N* = 150 (dashed line) monomers. The pdf for the center of mass with *N* = 16 (points), *N* = 48 (dashed line), *N* = 150 (full line). The normalized effective potential *U*_*N*_ (*r*) = *k*_*B*_ *T* log(*P* (*r*)) acting on a monomer is computed from the radial pdf. (c) The potential for the end and middle monomers is shown inFig. 22a. (d) The potential energy for the center of mass. The behavior for the end and the middle monomers are signi cantly different, con rming that the boundary has different effect depending on the position of each monomer. The parameters are summarized intable 2 (reproduced from [8]).

Using the simulated pdf, it is possible to de ne an effective potential *U*_*N*_ (*r*) acting on a single monomer using the relation
 *U*_*N*_(*r*) = - *k*_*B*_ *T* log(*P*(*r*)) (derived from eq. 170). Interestingly, the potential acting on the center of mass is large enough, so that it cannot reach the periphery of a ball during simulations (Fig. 22c, d) [8]. A direct derivation of this potential is laking but would certainly be very useful to study the collective effect of many monomer on one monomer through the boundary.

Finally, similar to the case where all the monomers can nd a small target, the arrival time distribution of a single monomer to a small hole is well approximated by a single exponential. To conclude, the distribution of arrival time of a monomer to a small target is almost Poissonian, however the rate depends on the location of the monomer along the polymer chain, that can be absorbed at the target site. NETP can also be studied when additional constraint such as stiffness is added to the model NETP (see for example [8]).

### 4.14 NETP with additional stiffness

Adding rigidity on the polymer can also in uence NETP. Indeed chromatin stiffness can be modulated by nucleosomes or bound proteins. Stiffness is accounted for by including the bending energy

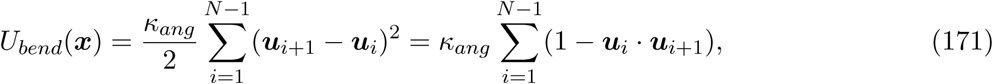

as described in subsection 3.8. With parameter described in that subsection, stochastic simulations for the arrival time of any monomer to the absorbing boundary are shown in g. 24, suggesting that the possible relation in dimension 2 (Fig. 24a)

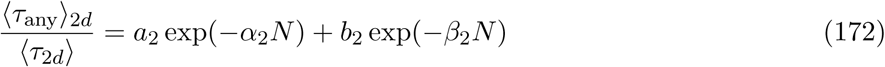

with *a*_2_ = 0.37, *b*_2_ = 2.9 and α_2_ = 0.02, β_2_ = 0.18 and in dimension 3, (Fig. 24b)

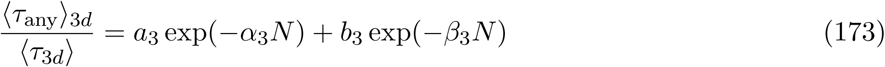

with *a*_3_ = 0.28, *b*_3_ = 1.64 and α_3_ = 0.01, β_3_ = 0.12. Compared to the non exible polymer, the maximum of the NETP is now shifted towards smaller values of *N*: The NETP is an increasing function of *N* for *N* < 6, and for large *N*, it is a decreasing function of *N*.

**Fig. 24.**
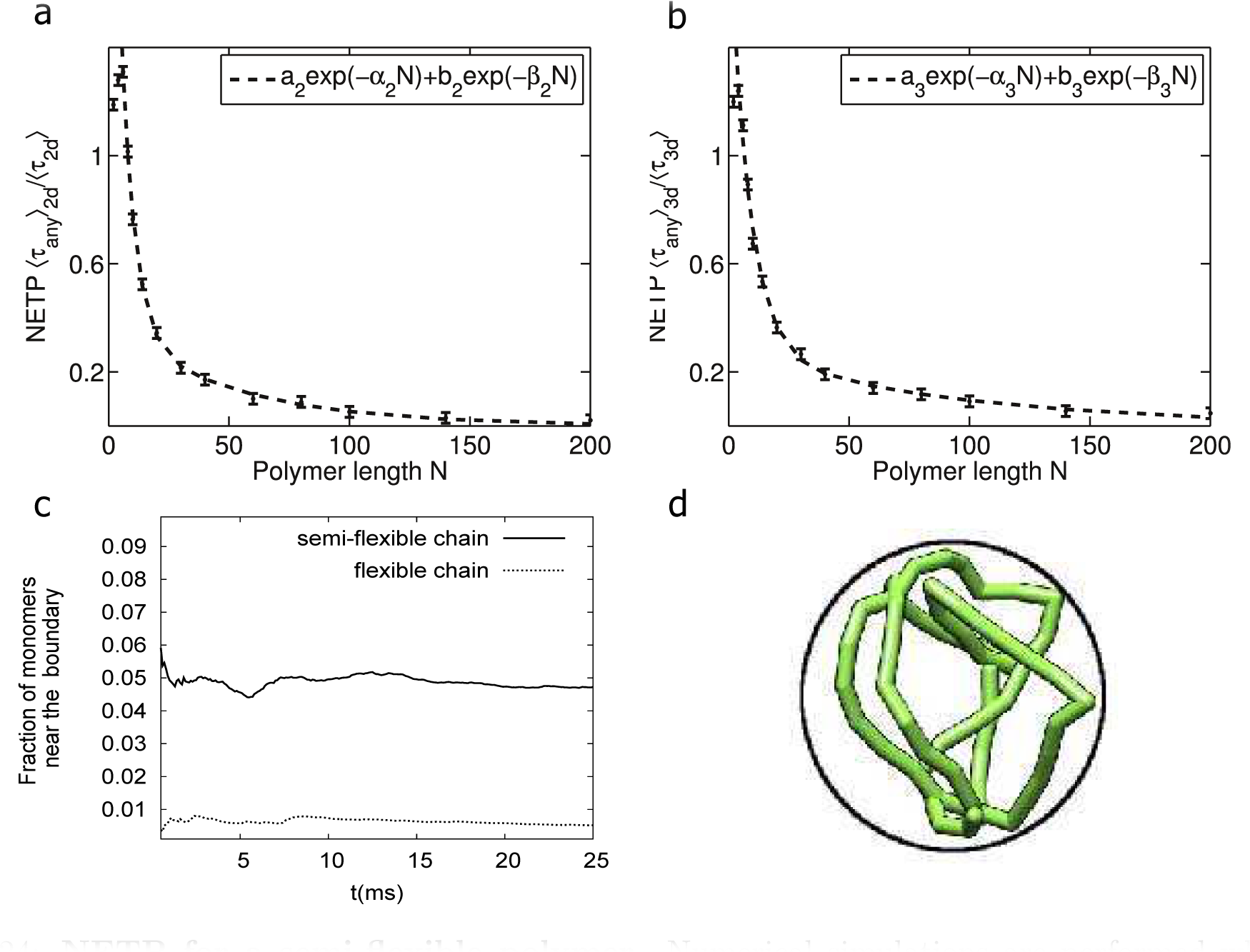
NETP for a semiflexible polymer. Numerical simulations are performed when any of the monomers (of a semi exible polymer) can reach the small target in dimension 2 (a) and 3 (b). The NETP is normalized to τ_0_ (the NET for 1 bead). Each point is an average over 2000. A double exponentialt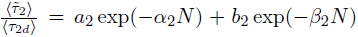 leads to *a*_2_ = 0.37; *b*_2_ = 2.9 and α_2_ = 0.02, β_2_ = 0.18 and in dimension 3,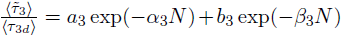. In three dimensions, the parameters are *a*_3_ = 0.28, *b*_3_ = 1.64 and α_3_ = 0.01, β_3_ = 0.12. (c) Fraction of time a monomer spends near the boundary (*>* 0.9 *R*, R is the radius) for a exible and non exible polymer. (d) snapshot of a non exible polymer where a large fraction of its monomer are located near the spherical boundary (reproduced from [8]).

In a con ned spherical cavity, exible polymers have a tendency to fill the available space and the probability of finding a monomer at the center of the cavity is higher compared to the boundary, when the persistence length is of the radius of the ball. On the contrary, for stiff polymers, the polymer chain has to bend abruptly near the boundary, where a large fraction of the polymer is found (Fig. 24c-d). Thus the search time for a target located on the surface by a monomer located on a nonflexible polymer is thus facilitated and the search should be almost two-dimensional, leading to a decreased NETP. To conclude, non exible polymer nd small targets faster than completely flexible ones, this is due to an increase probability to find monomers in the close vicinity of the boundary where the small target is located.

### 4.15 Summary of the looping time formulas

We now summarize the asymptotic formula for the looping time (MFLT) in dimensions two and three respectively, for small *ɛ*.

#### 4.15.1 Mean First Encounter Time formula in free space for a Rouse polymer

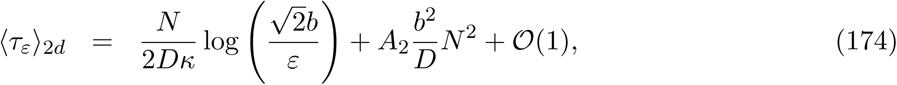

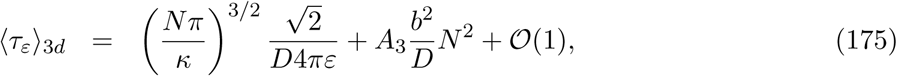

where *ɛ* is the radius centered at one end, *D* is the diffusion coefficient,
κ = *dk*_*B*_ *T*/ *b*^2^ is the spring constant with *d* the spatial dimension, *k*_*B*_ is the Boltzmann coefficient and *T* is the spring the temperature and *A*_2_ and *A*_3_ are constants, fitted to simulations data (see g. 17 for explicit values). Moreover, the distribution of looping time is

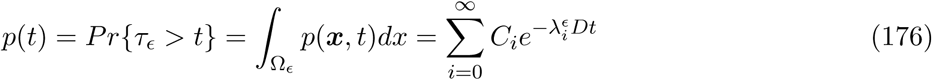

where *C*_*i*_ are constants and is well approximated by a sum of two exponentials. For *N* (*N* = 16 and 32) not too large, a single exponential is sufficient,

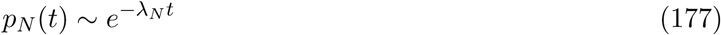

with *ɛ* = 0.1 *b*. Here λ_16_ = 0.0125 *b*^-2^, λ_32_ = 0.0063 *b*^-2^. For long polymers, a sum of two exponentials is more accurate

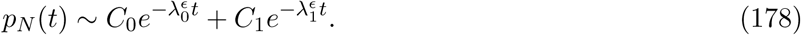

For *N* = 64, the numerical values are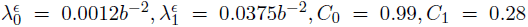 Although the two exponential approximation works well for small *ɛ <* 0.2 *b*, four exponents are needed for larger ɛ (*>* 0.4 *b*). For *N*∈ [4-64], *C*≈_0_1, while *C*_1_ remains approximately constant for a given value of *ε*. For example, for *ε* = 0.1 *b*, *C*_1_ varied with *N* from 0.2 to 0.28.

#### 4.15.2 Mean First Encounter Time formula in a con ned domain for a Rouse polymer

The MFET in a con ned ball of radius *A* is estimated when the boundary is accounted for by a parabolic potential, added to the Rouse potential *ϕ*_Rouse_ such that the total energy is the sum

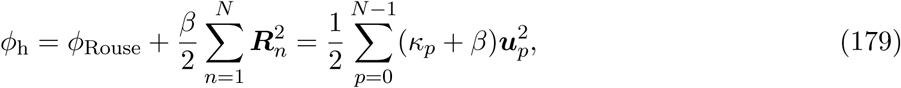

where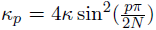
 ***u***_*p*_ are the coordinates in which *ϕ*_Rouse_ and the strength *B* is calibrated to the radius of the ball by

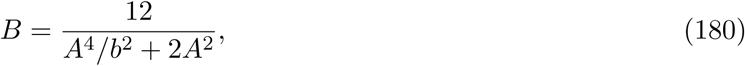

So that the root mean square end-to-end distance of the polymer in the potential field is equal to the square radius of the con ning ball domain *A*, that is [14]

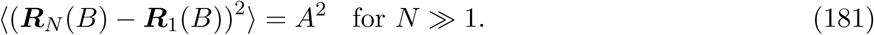

In that case, the MFET in a con ned ball of radius *A* 〈τ_h_〉 for two end monomers of a Rouse polymer to meet is given for *ɛ*≪ *b*,

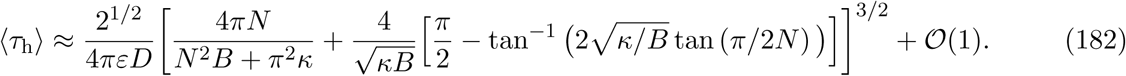

Note that as *N* tends to infinity, the mean looping time 〈τ_h_〉 does not diverge to infinity, but converges to an asymptotic value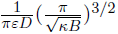. Finally, the distribution of looping times is always well approximated by a Poissonian law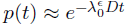 in contrast to looping in a free space.

### 4.15.3 Mean First Encounter Time formula for a *β*-polymer.

The mean encounter time for the two end monomers of a *β*-polymer is given by

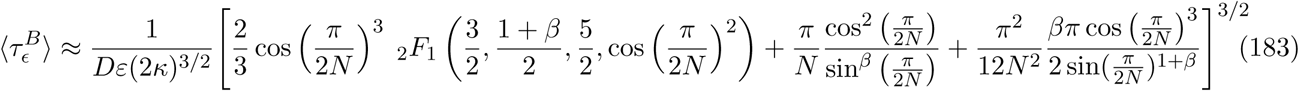

and for large *N*, it is possible to use the estimation

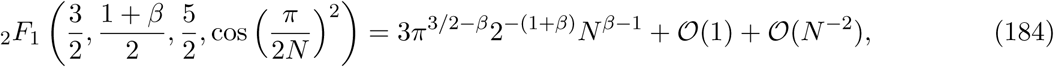

Where_2_ *F*_1_ is the Gaussian hypergeometric function [1]. The two other terms in 165 scale as *N*^β−1^. This asymptotic result is quite different for the Rouse polymer which scales as *N*^3/2^, while for a *β*-polymer, for *N* ≫ 1, the MFET scales as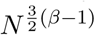

### Arrival time of a monomer to a small target located on the boundary of a bounded domain

Although, we are still missing an analytical derivation for the mean arrival time of a single monomer of a Rouse polymer to the boundary of a bounded domain, the approximation of this process as a single stochastic particle trapped in a single well lead to (seeeq. 170)

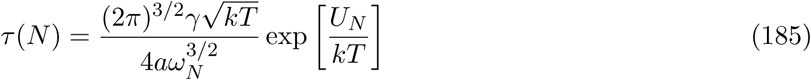

Where *a* is the size of the small target, *U*_*N*_(*r*) is the energy barrier, generated by the polymer due to the presence of the boundary of the domain at the target site and ω_*N*_ is the frequency at the minimum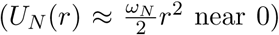 [248,8] and ω_*N*_ ∼ *N*. Additional research is expected to derive this statement from polymer model analysis.

## 5 Analyzing chromatin organization and dynamics using polymer model

This section is dedicated to large-scale simulations of chromosomes and telomeres [274,291,272,124] based on self-avoiding polymer models. The output of these simulations is the exploration of the polymer configuration space, the genesis of statistics that agree with experimental data. We shall see here that polymer models can reproduce chromosomal organization, observed in live cell imaging data in various species such as yeast or mammalian cells. We insist in subsection 6 on local chromatin reconstruction for the X-chromosome inactivation in female mouse embryonic stem cells using sophisticated polymer models.

Polymer looping remains the central event of chromatin organization and in parallel of presenting large scale simulations, we discuss the role of asymptotic analysis presented in previous sections 4 to interpret and extract information from empirical data. Asymptotic formula facilitated even further compared to simulations the exploration of the large parameter space. Indeed, the polymer parameter space remains too complicated to be continuously described by simulations. These simulations allow exploring and sampling small subregions of the configuration space, which is largely insufficient to classify structures, because the underlying spatial dimensions of the polymer space is very high and robust structures require high-dimensional geometrical parametrization. At this stage, simulations and asymptotic analysis provide complementary tools in searching for features in data. Interestingly, passage time formula between two sites reveals some key geometrical polymer features, without examining separately the topology or the possible polymer con gurations.

### 5.1 Numerical simulations and chromatin dynamics

Polymer simulations are used to describe DNA and chromatin folding inside the interphase nucleus of eukaryotic cells, which usually involves multiple length scales (from few to hundreds nanometers). To access higher-order structure, folding in confined nuclear space or distinct territories, polymer models reveal how chromatin looping is used in transcriptional regulation and how chromatin organization mediates long-range interactions [109,30]. Transient loop formation can originated from thermal uctuations. Looping interactions that do not directly involve an enhancer-promoter pair can modulate their interactions, as shown in Fig. 25 [74]: the simulation results show a 3 – 5 fold facilitation of enhancer promoter (E-P) contact frequency, comparable to observed changes in gene expression. In a chromatin model [74] where monomers are connected by harmonic bonds, with a 15 nm diameter representing 500 bps (approximately three nucleosomes), a permanent loop is formed when two monomers are connected with a harmonic bond of the same strength. Two such loops are formed in the simulations and a bending energy is imposed to account for the rigidity of the chromatin ber. Monomers interact via a Lennard-Jones potential (purely-repulsive potential, truncated at an energy *U* = 3 *kT*). Polymers are confined to a ball and initialized from an un entangled conformation. The probability distribution of loops for random walk and self-avoiding random walk was studied for chain lengths of size *N* =64, 128, 256 and 512, using Monte-Carlo simulations on a cubic lattice (boxes were of width *L* 64) and a density of *r* 12: 5%, which is similar to the conditions of interphase nuclei [30]. With a total of 4096 monomers, chromosomes are initially equilibrated as self-avoiding walks. After an initial equilibration step, the Monte-Carlo algorithm samples different loop configurations. A loop is formed with a certain probability *P* when two monomers co-localized. The life time of a loop is Poissonian. Results of simulations are shown in Fig 26. The contact map is similar to one obtained by 5C data, for a polymer with different looping probabilities: in the self-avoiding walk polymer model (Fig. 26A), there are only few contacts between beads located far apart. Increasing the looping probability (Fig. 26B and C) results in a strong increase of both the number of loops and the abundance of large loops [30].

**Fig. 25:**
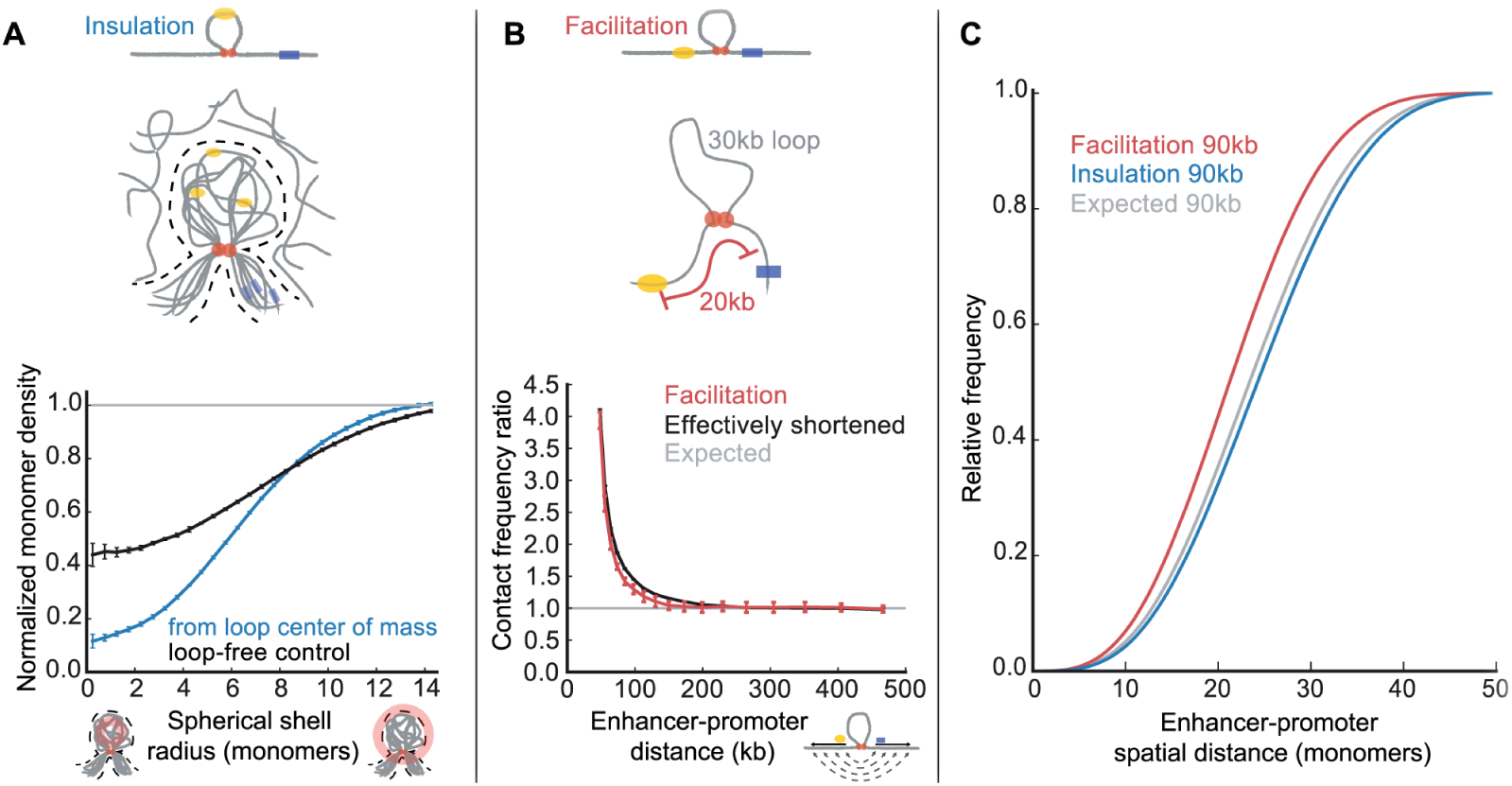
Mechanisms of insulation and facilitation. (A) (top) Insulation mechanism: steric exclusion by a chromatin loop from the superposition multiple loops(grey, with enhancer and promoter) and their sterically excluded region (dashed lines), surrounded by other distal regions of chromatin (grey). (bottom) Density of distal monomers (i.e. outside the loop and > 10 kb from the loop base) as a function of radial distance from the center of mass of the loop. The loop-free control exactly repeats this procedure for an equivalent region without a loop. Both are normalized using respective radial-position dependent spatial density. (B) (top) Facilitation mechanism: an E-P pair flanking a loop has an effectively shorter genomic distance; here an E-P pair with 50 kb separation and a 30 kb loop behaves similarly to an E-P pair separated by 20 kb in a region without a loop. (bottom) Comparison of contact frequency ratios for the above situations, as a function of E-P distance. (C) Simulated cumulative distribution of spatial distances for an E-P pair with a genomic distance of 90 kb (reproduced from [74]).

**Fig. 26:**
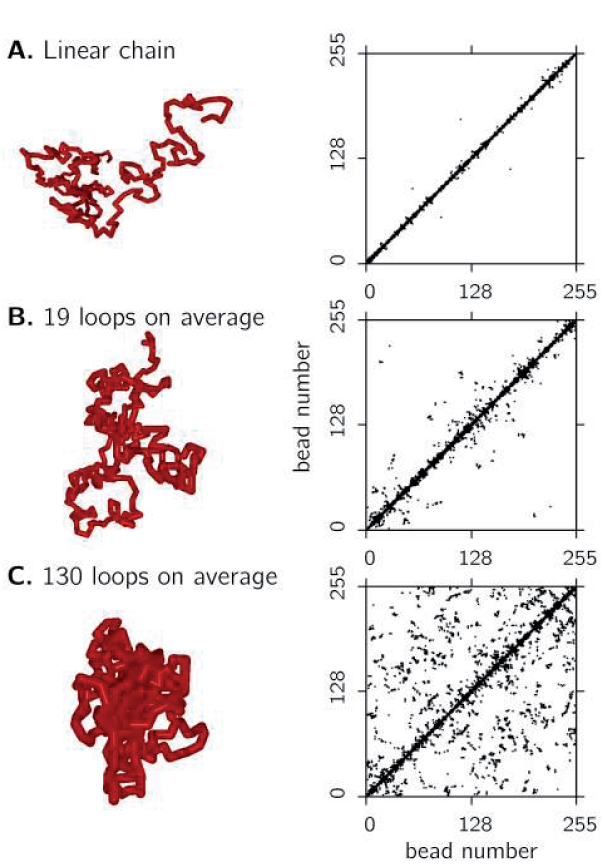
Intra-chromosomal contacts of isolated model polymers. Equilibrated polymers with different looping probabilities. (A. linear chains (no loops), B. on average 19 loops per conformation and C. on average 130 loops per conformation) co-localized beads were determined and marked by a black square. For each image, the contacts of 4 independent polymer conformations are plotted. Linear chains (A) have not so many contacts between beads which are widely separated along the contour of the polymer. Increasing the probability of functional loops (B and C) results in a boost of contacts both between close-by segments as well as between segments having a large genomic separation (reproduced from [30]).

The role of looping in large-scale (super Mega base pairs) folding of human chromosomes was studied using self-avoiding walk polymer model that can generate transient looping [268]. Chromatin compaction was shown to relate to a reduction of the concentration of two looping proteins (knocked down the gene that codes for CTCF and the one coding for Rad21, an essential subunit of cohesin). This effect can be explained by selectively decreasing the number of short-range loops, leaving long-range looping unchanged [268]. Polymer configurations for different looping probabilities at short and long-ranges are shown in Fig. 27. Higher order loops involving more than two monomers was considered to model discrete spatial transcription foci: the formation of these foci resulted from the encounter of sparse interacting sites located on the same chromosome [135]. In these foci, several genes related to the same transcription system, come physically in close proximity in order to share regulatory and transcription proteins.

**Fig. 27:**
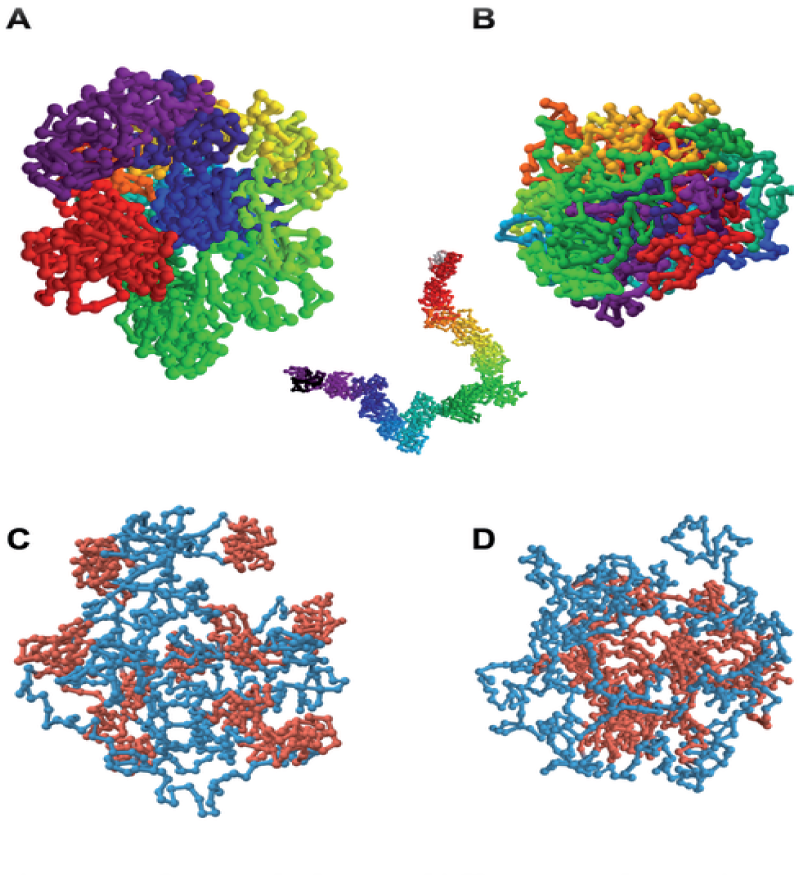
Polymer conformations of model at different looping regimes. [268]. (A) Conformation with high short-range (pshort = 0.12) and low long-range (plong = 0.04) looping probabilities and (B) the same polymer with low short-range (pshort = 0.04) and high long-range (plong = 0.12) looping probabilities. The color code labels the monomers of the polymer according to the visible spectrum along the length of the polymer. The inset in panel A displays the same situation as in after abolishing all long-range looping (plong = 0), showing that a uniform thick ber is formed. Conformation of a polymer with high short-range (pshort = 0.16) and low long-range (plong = 0.02) looping probabilities and (D) the same polymer with low short-range (pshort = 0.02) and high long-range (plong = 0.16) looping probabilities. Here, topological domains are labeled red and non-looping linker regions blue (reproduced from [30]).

In summary, polymer models with additional short and long-range looping can account for chromatin organization at various scales and can be used to explain some of the modulation of gene activation.

### 5.2 Characterizing chromosomes and nucleus architecture using large scales simulations

Chromosomal territories are nuclear regions occupied by a single chromosome and are found in most higher eukaryotes, observed in live cell imaging data [32]. But lower eukaryotes, such as the yeast Saccharomyces Cerevisiae, lack these chromosome territories and their chromosomes seem to be loosely arranged. Interestingly, in lower eukaryotes such as plants and flies, chromosomes are often polarized, with the ends of the arms (telomeres) on one side of the cell nucleus. The two arms are anchored at a single point (the centromere) located on the opposite side of the nuclear surface.

These structures are associated with gene territories, where genes of similar metabolic function are together [25]. Territories are revealed by microscopy imaging and also chromosome capture experiments, and are reproduced using large scales simulations [274,291,272,212] of self-avoiding polymers. However, the output of these simulations remains difficult to compare to experimental data except at equilibrium, where the probability distribution function of chromosomes and moments can be estimated. Detailed simulations of higher eukaryotes are still lacking due to the complexity of the simulations, the number of degrees of freedom and the lack in understanding of the basic molecular mechanisms involved in establishing and maintaining these territories.

Transient re-organization of the genome is also hard to simulate, because it is difficult to identify relevant time scales. Although different coarse-graining simulations of the nucleus have been developed [83,290], a universal model for chromatin is still to be found because it is difficult to identify steady or quasi-steady polymer states from experimental data (such as Hi-C). By taking into account constraints acting on the chromatin ber, polymer simulations can reproduce measured physical quantities that scale with distances between genomic loci. Polymer models can also reproduce chromosomal territories, and probabilities of contacts between loci measured by chromosome conformation capture methods [83].

Simplified polymer model lead to various striking conclusions: classical models have ignored basic DNA identity such as sequences, but they accounted for the centromere, telomeres, and the ribosomal DNA (rDNA) to model the interphase yeast nucleus chromosomes. The first output is that the large-scale architecture of the yeast nucleus is dominated by random forces moving polymers, rather than by a large number of DNA-specific deterministic factors [290]. Recently, chromosomes were modeled as semiflexible polymers with freely jointed chains of non-intersecting rigid segments (Fig. 28). This coarse-grained model neglected chemical bound details of the chromosomes, DNA sequence and possible histone modi cations, but could reproduce chromosomal organization.

**Fig. 28:**
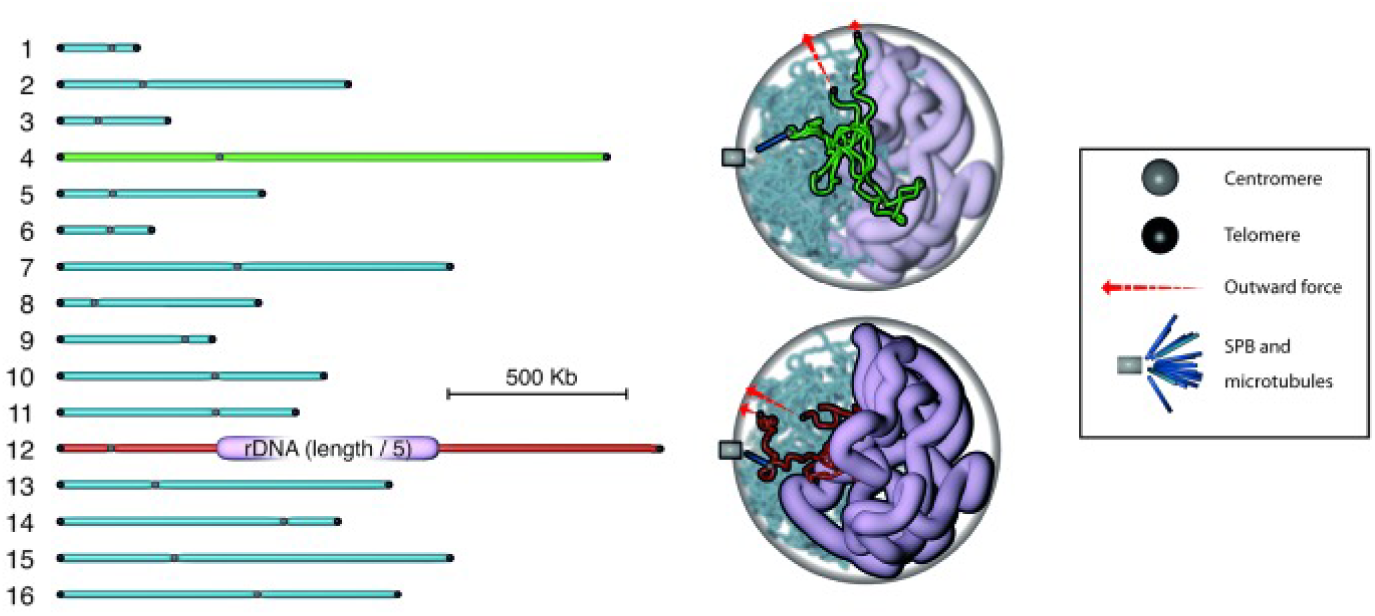
Computational model of chromosome architecture in the budding yeast nucleus [290]. The 16 chromosomes of the haploid yeast genome (shown stretched out on the left), are modeled as random polymer chains undergoing Brownian motion and con ned to a spherical nucleus (right). Only three DNA sequence-specific constraints are included: centromeres, which are attached to the SPB via short rigid microtubules; telomeres, which are tethered to the nuclear envelope by an outward force; and the rDNA locus, which is modeled as a chain of increased diameter. Chromosomes 4 and 12 are highlighted in the 3D snapshot of the simulation shown on the right (reproduced from [290]).

Using freely-jointpolymer chains requires specifying the following parameters: the length of each segment, equals to 50 *nm* accounting for the chain rigidity, the number of base pairs corresponding to each segment (DNA compaction equal to 5 kbs), and the segment diameter (20 nm), see Fig. 28. Other values used in modeling and simulations are possible (persistence length of dsDNA is chosen around 50 nm and for chromatin ber it is 150 nm [106]). A second output is that such model can replicate many of the steady-state statistics measured by contact frequencies across the genome (genome-wide chromosome-conformation capture), showing that the main statistical organization of chromosomes in the nucleus results from elementary stochastic polymer dynamics. This model was used to examine the impact of reducing the volume of the nucleolus: the model predicted a displacement of telomeres further away from the spindle pole body, in agreement with prior observations [269].

### 5.3 Interpreting chromosome capture data inside nuclear territories from polymer looping

Looping properties of polymer model provide constraints on the statistics of long-range interactions and the anomalous exponent that can be extracted from chromosome capture (CC) data. Indeed, analytical expressions for the MFET predict a decay exponent presented in eq.165. For example, long-range interactions introduced in the *β*-polymer model can be due to the condensin proteins, part of the Structural Maintenance of Chromosomes (SMC) protein family [292]. These proteins are capable of generating large loops that hold together sites located far apart along a chromosome. During mitosis, the concentration of condensin molecules increases, resulting in an increase of the Young modulus of the chromosome [169].

We now summarize the CC-data. They contains statistics of contact frequencies that require a physical model for interpretation. In a reconstruction procedure, sites that have high encounter frequencies should be positioned in closer proximity inside the nucleus domain. Given a set of encounter probabilities *P*(*x*_1_, *x*_2_), *P*(*x*_1_, *x*_3_),‥, *P*(*x*_1_, *x*_N_), *P*(*x*_2_, *x*_3_),…, *P*(*x*_N_, *x*_N_), where *x*_1_, *x*_2_,…, *x*_N_ are genomic sites, the following reverse engineering problem should be resolved: how to reconstruct the spatial positions *x*_1_,…, *x*_*N*_ from encounter probabilities? Is this reconstruction possible? This construction remains an open problem and requires additional information.

When sites are located on the same chromosome chain, using a Rouse polymer, the distances between any site in a free space increase with the distance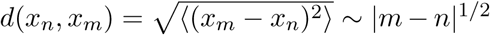, while the encounter probability decays with the genomic distance as

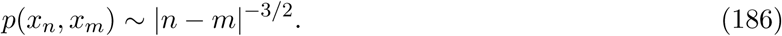

When sites are not located on the same chromosome, the encounter probability is a characteristic feature of the long-range nuclear organization and a reconstruction procedure should account for this decay. We should assume that the genome is at equilibrium. By averaging over millions of cells, the CC-data is averaged over population [152]. In reality, the chromatin is never at equilibrium, as it is constantly pulled by different proteins and sometimes open during transcription. In this case, interpretation of the data should take into account the dynamics and the encounter probability given by

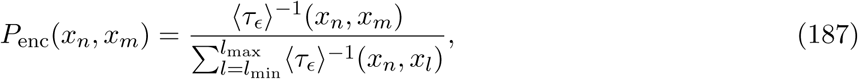

where 〈τ_e_〉^−1^ (*x_n_*, *x_m_*) is the rate of encounter between sites *x*_*n*_ and *x*_*m*_ [9].

To conclude, a procedure for reconstructing the chromatin organization should assume that polymer equilibrium is reached. So far, there are no generic procedures to reconstruct chromatin organization from data. As we shall see in the next sections, not only a polymer model should be chosen, but also a spatial scale (distance between two monomers) and some constraints on the empirical statistics to be prescribed. For example, it is possible to construct a polymer model such that the encounter probability of the simulations and the empirical data is the same [246]. Other criteria may be chosen as well. These constraints insure that the reconstructed model contains some of the key statistical characteristic of the data.

### 5.4 Interpreting CC-data using polymer models

At a resolution of 1Mbp, chromosomes can be divided into segments, with higher inter-segment contact frequencies between different segments [157]. These sub-segments form blocks in the frequency matrix. The encounter probability decays as a power-law of the genomic distance *P*_enc_ ∼ *n*^−1^ (see fig.29a). An exponent decay of 1.5 suggests an equilibrium polymer and corresponds to a flexible polymer model. However, the power law extracted from data differs from 1.5, leading to a constraint for the choice of polymer models. Various models have been proposed to explain qualitatively these observations, as we shall now review.

#### 5.4.1 Fractal polymer globule

The compact conformation frequency data suggests that the polymer conformation of chromosomes followed a “fractal globule” image [96,97]: this non-equilibrium state polymer consists of non-entangled structure, where beads are crumpled in small non-overlapping domains, which gradually fill a domain according to a auto-similar rule, without crossing [183] (Fig. 29b). The DNA may be compacted in a non-entangled manner in the nucleus, such that its opening during transcription would be not impaired by any nodes. This representation is now clearly insufficient to account for the chromatin organization [196].

**Fig. 29:**
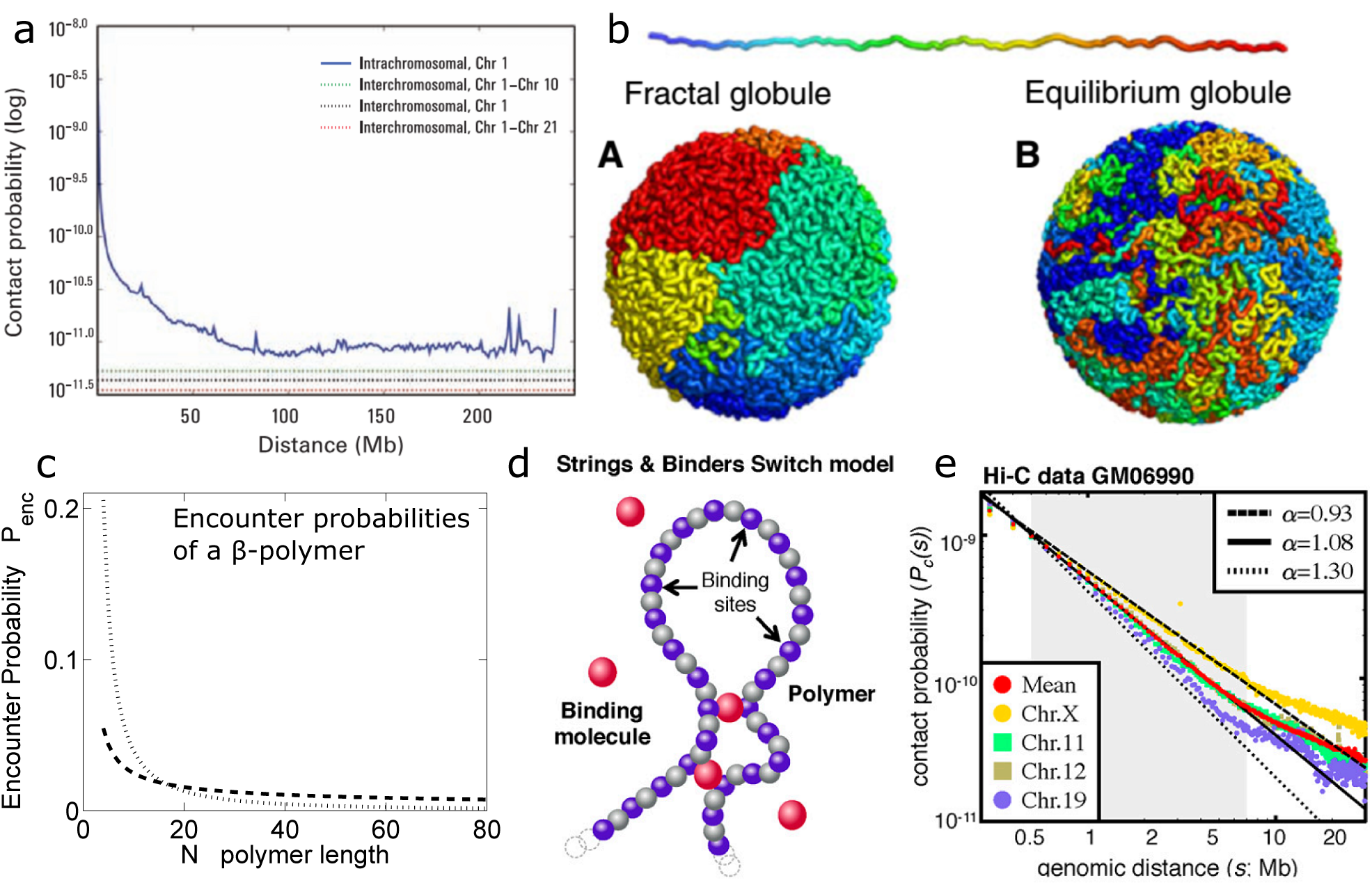
Encounter frequencies of genomic sites inspired chromosomes polymer models: (a) Probability of contact decreases as a function of genomic distance on chromosome 1, eventually reaching a plateau at 90 Mb (blue). The level of interchromosomal contact (black dashes) differs for different pairs of chromosomes; loci on chromosome 1 are most likely to interact with loci on chromosome 10 (green dashes) and least likely to interact with loci on chromosome 21 (red dashes). Interchromosomal interactions are depleted relative to intrachromosomal interactions. [157] (b) Conformations of the fractal (A) and equilibrium (B) globules. The chain is colored from red to blue in rainbow colors as shown on the top. The polymer globule has a striking territorial organization, which strongly contrasts with the mixing observed in the equilibrium globule. (c) The normalized encounter probability for the polymer ends for the Rouse polymer for which *β* = 2 (points, *P(N) ~ N^-1.5^* and a β polymer for which *β* = 1:5 (dashed line, *P(N) ~ N^-1.5^* [10]. (d) Schematic of the string binder and switch model. The chromatin is modeled as a SAW polymer [19]. A fraction of the monomers (purple) can bind binder molecules (magenta) diffusing in the solution, which transiently fixed polymer looping events. (e) Contact probability, *P*_*c*_(*s*), was calculated separately for different chromosomes from published Hi-C dataset in human lymphoblastoid cell line GM06990 [157]. Chromosomes 11 and 12 follow the average behavior reported (reproduced from [157]) in the 0.5-7 Mb region (shaded in grey), with exponent of approximately 1.08. Chromosomes X and 19 deviate from the average, with exponents of approximately 0.93 to approximately 1.30, respectively. In a given system, different chromosomes can have different exponents.

#### 5.4.2 Interpreting CC-data using the *β*-polymer model

The spatial organization of the chromatin can also result from specific protein interactions. In the context of the *β*-polymer model, the encounter probability *P*_enc_ can be computed from relations 154 and 155. The normalized sums 159 and 160 for the encounter probability

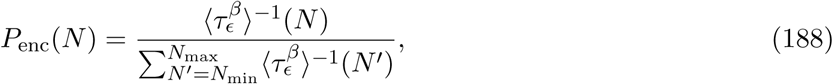

reveal the decay rate shown in Fig. 29c for *N*_min_ = 4 and *N*_max_ = 80: the encounter probability *P*_enc_ does not necessarily decay with *N*^−3/2^ nor *N*^−1^, but *N*^−3/2(*β*−1)^, where *β* is identi ed by tting a curve to data [157].

To conclude, it is possible to relate the empirical decay rate to the intrinsic properties of the chromatin, incorporated in the value of the parameter *β*.

#### 5.4.3 String binder and switch model (SBS) model

Although chromatin is quite compacted, another approach to interpret the CC data is provided by the string-binder and switch model [19,20] (Fig. 29d), where loops are generated randomly insidea ball. The chromatin is constraint with many binding sites that represent different DNA binding proteins [171]. To account for protein interactions, chromatin is represented as a flexible chain with self-avoiding interactions (Fig. 29d) [194,195]. A fraction *f* of the polymer sites can bind particles (binders) with a binding energy *E*_*X*_. The binder molecules can bind to more than one monomer, allowing transient connections to be formed between polymeric sites. These binders can represent chromatin molecules such as the CTCF Zinc finger protein [136], known to mediate inter and intra-chromosomal contacts [293].

Polymer conformation were studied using Monte-Carlo simulations for a concentration *c*_*m*_ of binder molecules. Adding binders affects the polymer size and can lead to collapse at a specific binder concentration *C*_*tr*_. Interestingly, changing the concentration of the binders can reproduce long-encounter in chromosome capture experiments (Fig. 29e).

In summary, the SBS model accounts for the compaction observed in CC data based on specific interaction acting between distance monomers. However, the exact location and numbers of binders discussed in [194,195] is not yet constraint and the empirical encounter probability cannot be recovered. Interestingly, it should be possible to account for these binding interactions that can further change during development or transcription.

With the increasing richness extracted from CC data, their interpretation has moved from general mean field models (fractal globule, *β*-model, SBS) to more precise but heterogeneous polymer models. This change allows exploring the reorganization of mitotic Chromosome [192] or the relationship between transcription and chromosome conformation of the X-inactivation center, where the regulatory element are located on the same chain (*cis* regulation [91]), see below.

### 5.5 Using CC-data and encounter probabilities to describe nuclear subcon nement

The encounter probabilities (relation 188) between chromosomal sites at steady state is used to extract the confinement radius from measured looping distribution obtained from 4C data of yeast *Saccharomyces cerevisiae* [75]. Indeed, the frequency of encounter is inversely proportional to the MFETC, thus the encounter frequency can be obtained from the MFETC formula (eq. 165). Although the MFETC can vary with the monomer position along the polymer chain that it will encounter, the normalized rate (the reciprocal of the MFETC divided by its integral over the length) of monomer interaction (as experimentally measured) is independent on the encounter rate radius *ε* (as it cancels out in the normalization). The encounter probability depends only on the distance between two monomer pairs along the chain and is in fact not much affected by the remaining polymer tails.

The chromatin molecule can be organized in higher order structure such as stable separated domains that result from local specific interactions. At the current stage, the analysis of the model does not take into account the effects due to higher order structure in chromosomes. Another limitation of this model when applied to yeast, is due to the Rabl chromosomal configuration where the centromeres are connected to the mitotic spindle body. This interaction breaks the radial symmetry. The above approach was applied to 4C data in yeast, neglecting higher order structures and considering that the radius of yeast nucleus is fairly small (1 μm).

The encounter probability data was computed from data presented in [75] and for a locus at position 99kbp from the right-end of chromosome II in the yeast (Fig. 30d) and fitted with the encounter formula eq.128, where *β* is the only free parameter, leading to *β* = 3.91 × 10^−6^nm^−2^. Using formula 180, an effective confinement radius was extracted [9] of *A* = 230nm, representing a subdomain of the yeast nucleus [269]. Moreover, for large distances along the strand, the encounter probability reaches an asymptotic value rather than going asymptotically to zero (Fig. 30d). This decays shows that the encounter of loci pairs is affected by nuclear confinement. Extending the analysis of polymer models, beyond the Rouse linear chain would be relevant to account for higher-order organization of the chromosome. Geometrical constraints of chromatin loci are classically obtained by single particle tracking of a florescent loci on the chromatin. Extracting spatial information from CC data, which can be later on compared to single particle tracking experiments would certainly lead to novel results about the organization of local subnuclear compartments.

**Fig. 30:**
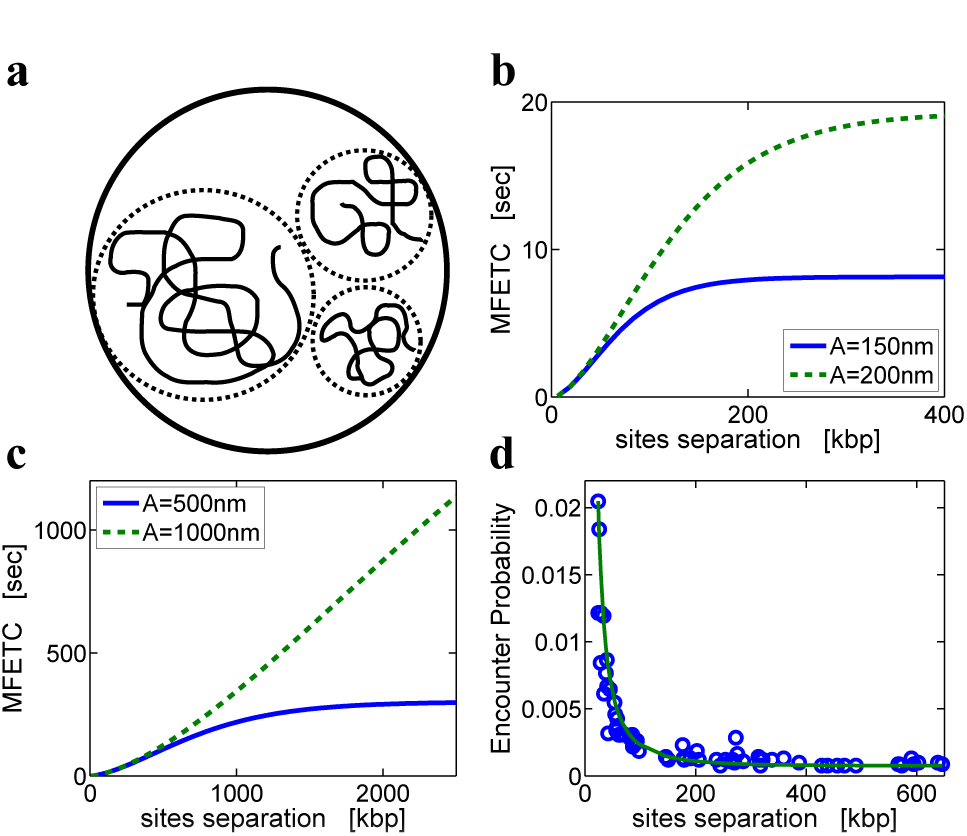
Encounter between two DNA loci inside a domain. (a) Schematics representation of gene territories inside the nucleus. (b-c) MFETC of two sites located on the same chromosome as a function of their separation along the DNA strand. Chromosomes are con ned in domains of radii *A* = 150 ; 200 ; 500 ; 1000nm. The encounter radius is *ε* = 5nm. (d) Encounter probability of a locus on chromosome II in the yeast and fitting of our looping probability formula. The effective confining radius extracted using our calibration formula is *A* = 240nm. Parameters are *b* = 30 *nm* and *D* = 310^3^ *nm*^2^ *s*^-1^ (reproduced from [9]).

## 6 Chromatin modeling using polymer

Polymer models are used to extract specific features from chromosomal capture data, presented in section 5. There is a large diversity of genome folding. For example, metazoan genomes are folded in a hierarchy of structures, while in mammals, each chromosome is folded independently of each other and occupies a well-defined territory (in interphase). The ability of higher Eukaryotes to regulate and differentiate depends precisely on different chromatin states. Chromatin domains vary in length between kilo to Megabps, but are separated by boundary-imposing-elements called insulators. Eukaryotes have chromatin domains with distinct functionality. The open chromatin called the euchromatin less dense than the rest permits a higher accessibility of proteins to gene sites. However, the dense heterochromatin region, much less accessible to transcription machinery is associated with gene silencing.

Inside each chromosome territory, some sub-compartments have emerged, corresponding to large (multi-Mbps) active and inactive chromosomal regions, which tend to interact with regions of similar transcriptional activity and are segregated from each other [157]. Interestingly, these compartments were found to be cell-type and developmental-stage specific, as they reflect the transcriptional state of a chromosome. Recently, sub-compartment folding units called topologically associating domains (TADs) or topological domains [71,197,121,245] were characterized as sub-megabases (around 800kbs) domains with increased three-dimensional interactions, imposing a genome partition in adjacent regions of preferential chromatin contacts. TADs are further partitioned into sub-TAD domains and loops near binding factor (CTCF) sites [214,253,280] and co-factor cohesin [20]. Gene expression depends on distal regulatory regions called enhancers, which can be located mega bps away from the controlled gene [240] and TADs regulate the interactions of distal regulatory elements and enhancers [164,260].

Understanding how TADs are established and maintained, how is the three-dimensional conformation of chromatin within single TADs the mechanisms has bene ted from polymer models. Indeed, mean-field approaches described in section 5 have been used to reconstruct the average three-dimensional conformations of genomic regions starting from 3C-based datasets [22,75]. While these approaches can be useful to represent the conformation of a genomic region, they do not provide any mechanism to explain neither conformational changes nor can they account for the cell-to-cell variability. The statistical and dynamical properties of the chromatin fiber within single TADs should depend on genomic elements/DNA binding factors. To account for cell-to-cell variability and population heterogeneity, various classes of phenomenological polymer models were proposed [91,270,132,246] and the goal of this section is to summarize them.

### 6.1 Chromatin modeled by copolymer

Starting with a semi-flexible chains and self-avoiding interactions (see section 3.7), the configuration of a copolymer is by de nition driven by thermal motion and repulsive interactions. The resulting structure is a homogeneous chain, except at the polymer ends. Heterogeneous structures are generated by adding interactions in the energy term

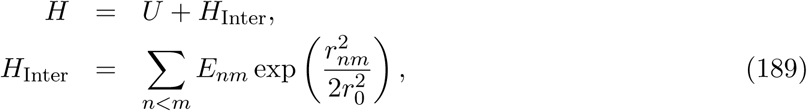

where the potential *U* includes spring forces and LJ-interactions (eq.32), while the energy *H*_Inter_ accounts for specific interactions between different chromatin sites. The Gaussian potential *H*_Inter_ depends on interaction strength *E*_*nm*_ between monomers *n* and *m* and *r*_0_ and defines the range of interactions [132] (see also [274] for a Gaussian potentials).

To reproduce chromatin organization [44], Langevin dynamics simulations and Monte Carlo methods [198] are used to explore the configuration space [132]. Monomers with the same states interact directly following eqs.189 (Fig.31a). In that case, the interaction energy is the sum of two terms: *E*_*mn*_ = *U*_ns_ +_*mn*_ *U*_*s*_, where the energy *U*_ns_ between every pairs of monomers accounts for the confinement into the nucleus and *U*_*s*_ is an attractive interaction between monomers having the same (epigenetic) state. Results of simulations show immiscibility between monomers of different types and phase separation of the polymer. This model was tested on region of Drosophila melanogaster chromatin [79], where five typical epigenomic states can co-exist: two euchromatic states and three heterochromatic states. The experimental Hi-C map (Fig.31bA) shows the internal folding (black) and active domains with almost uniform intra-domain contact probabilities, where inter-domain contacts are especially numerous (between black domains). A region is modeled by a chain of 131 beads with 10 kb per bead. The encounter probability is shown in Fig.31bB: the pattern of inter-black domain interactions and some long-range contacts between active domains can also be reproduced (Fig.31bC). The polymer section (black chromatin) forms a compact metastable globule that transiently dissociates, and small active domains are expelled at the periphery (Fig.31bD). While the model could account for some experimental properties, the initial hypothesis that different epigenomic states attract each other remains to be validated.

**Fig. 31:**
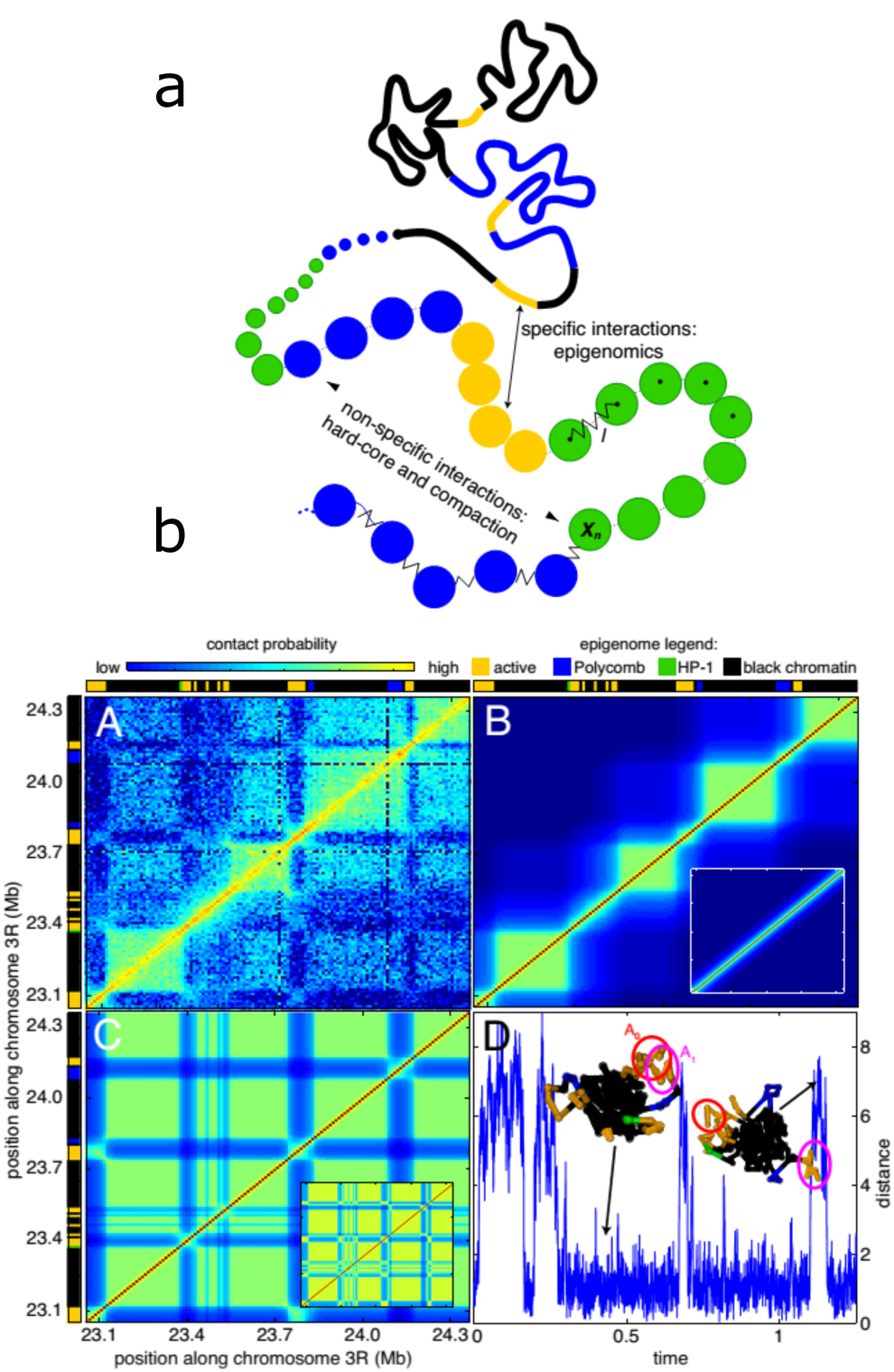
Chromatin modeled as a co-polymer. (a) Block copolymer model: the chromatin is modeled as a self-avoiding bead-spring chain where each monomer represents a portion of DNA (10 kb) and is characterized by an epigenetic state: yellow (active), green (HP1-like heterochromatin), blue (Polycomb-like heterochromatin), black (repressive chromatin) [79]. The model integrates non-specific and specific short-range interactions to account respectively for the effective compaction of the chain and for epigenomically related affinities between monomers [132]. (b) (A) Experimental Hi-C contact map for the chromatin region located between 23.05 and 24.36 Mb of chromosome 3R (from [245]). Domains [79] represented at top and left borders are active (orange), Polycomb (blue), HP-1 (green) and black chromatin. (B and C) predicted contact maps inside the multistability region (*U*_*ns*_ = -25kBT, *U*_*s*_ = -63kBT) starting from a coil (B) or microphase separated (C) configuration (see insets). (D) Evolution of the distance between the centers of masses of the active domains A0 and A1 along a simulated trajectory. Insets: conformations of the chain (reproduced from [79]).

### 6.2 Chromatin condensation/aggregation depending on chromatin state

Chromatin de-condensation was recently visualized in Drosophila Kc167 cells [227,28] revealing multiple genomic domains. These domains are divided into three major epigenetic states: transcriptionally active, inactive, and polycomb-repressed. This classification is made on the basis of histone modi cation and regulatory protein enrichment (Fig. 32a).

**Fig. 32:**
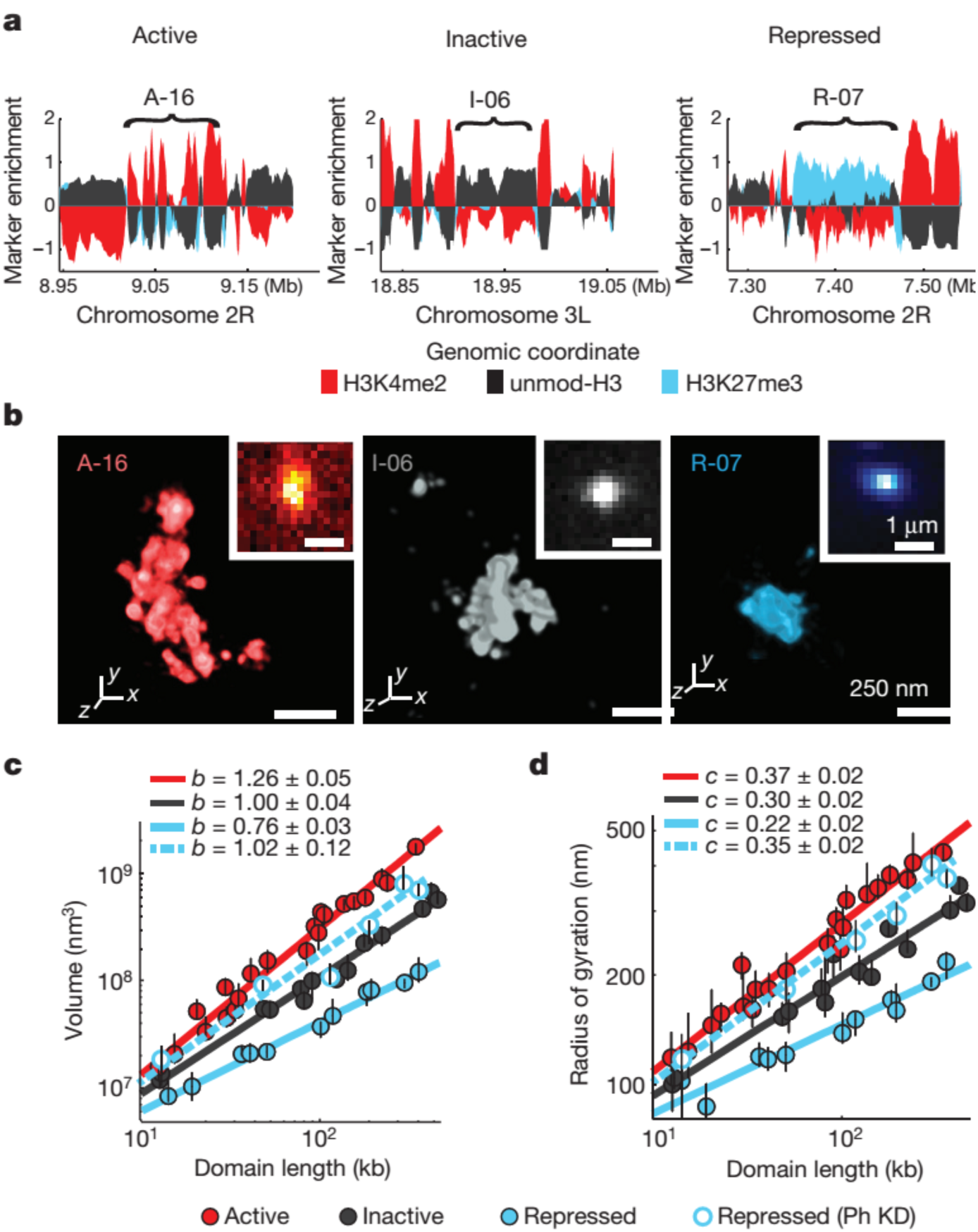
Chromatin in different epigenetic states exhibits distinct packaging and power-law scaling. (a) Enrichment profile of H3K4me2 (red), H3K27me3 (light blue) and unmodi ed H3 (black) in three genomic regions, each harbouring an example Active, Inactive or Repressed domain (indicated by brackets). Marker enrichment, as defined in Supplementary Methods, was determined from ChIP-seq data20. (b) 3D-STORM images of three distinct epigenetic domains in a, labeled by in situ hybridization with DNA probes conjugated to the photoswitchable dye Alexa-647, shown with their corresponding conventional images in the inset. Each epigenetic domain appears as a single region in nearly all cells due to homologous pairing in the tetraploid Kc167 cells. (c) loglog plot of the median domain volume as a function of domain length for Active (red solid circles), Inactive (black solid circles) and Repressed (light blue solid circles) domains, as well as for Repressed domains in Ph-knockdown (Ph KD) cells (light blue hollow circles). Error bars represent 95% resampling (*n* ≈ 50 cells). The lines indicate power-law fits, with the scaling exponent *b* shown in the legend. (d) As in c but the radius of gyration depends on the domain length with the scaling exponent *c*.

Active domains scaled differently in size than the repressed/inactive ones (see Fig.32b) and fitting a power-law for the relation between volume *V* and domain length *L* leads to *V ~ L*^*b*^ where *b* is largest for the active state, decreasing for inactive and is the smallest for silence domains (Fig.32c,d). This result suggests that packaging density controls genetic activity. While active and inactive chromatin had a self-similar structure, which is a uniform power-law across large genomic length, the radius of gyration computed inside subdomain of the imaged chromatin behaves differently (see Fig.32): repressed chromatin is composed of smaller sub-domains with a 60% - 80% overlap.

### 6.3 Constructing an effective polymer model from Hi-C maps to sample the space of chromatin configuration

Chromosomal long-range interactions with an effective potential was recently used to describe chromatin dynamics [91]: this model permits extracting single-cell information from population-averaged 5C or Hi-C maps. Despite its descriptive power, as we have shown in a previous section, 5C and Hi-C are population-averaged assays in which the contact probabilities is averaged over populations of millions of cells. These data do not permit access to the cell-to-cell and temporal variabilities of the chromatin conformation. However, these data can be used to constrain polymer simulations based on the contact frequencies measured experimentally. Two key regulators of X-chromosome inactivation in female mouse embryonic stem cells [207] were analyzed using polymer simulations and used to reconstruct TADs, identified in the X chromosome inactivation system (Fig.33A). The promoters of Tsix and Xist are shown in Fig.33B.

**Fig. 33:**
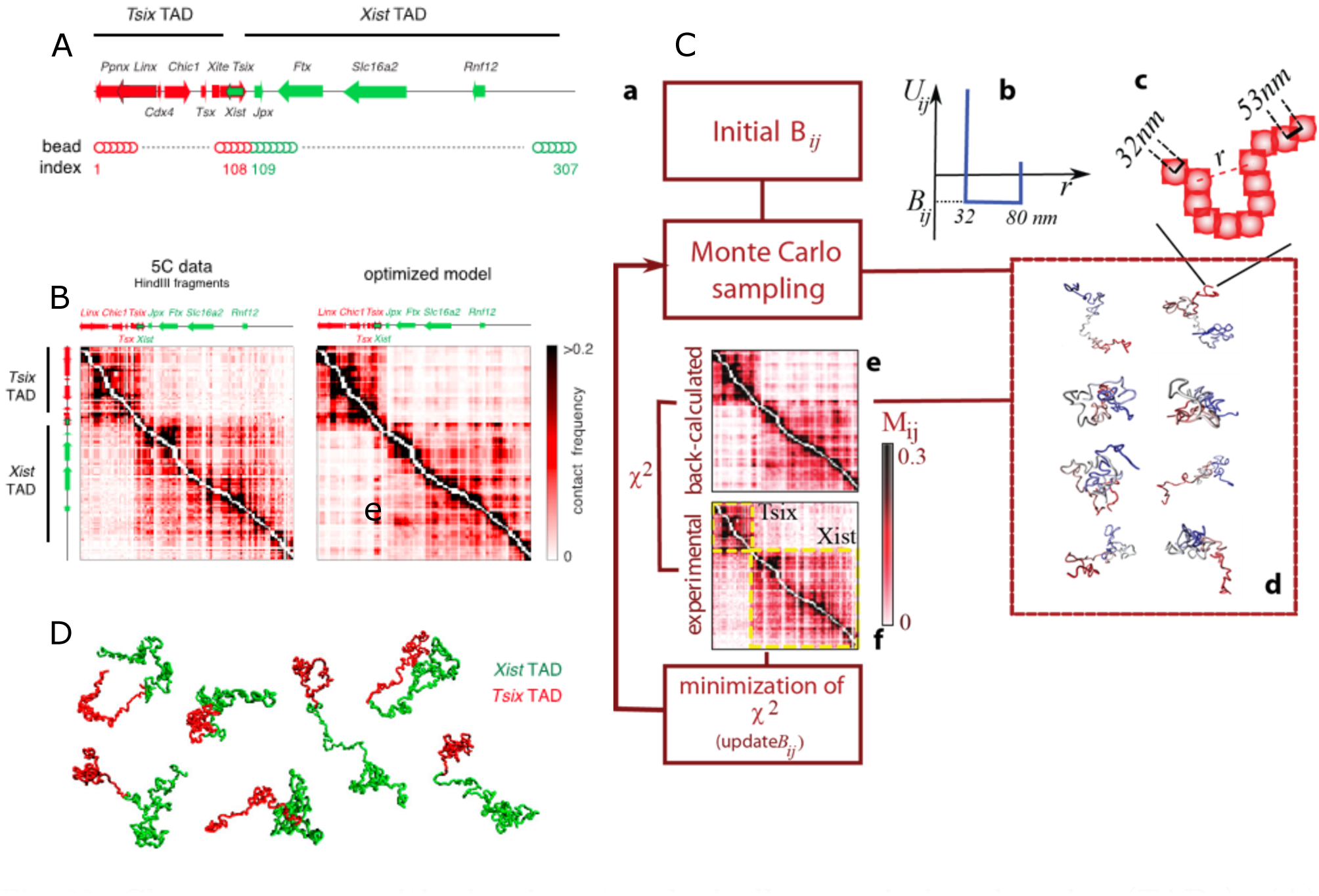
Chromosome partitioning into topologically associating domains (TADs). **(A)** Model of polymer ber for simulating the Tsix and Xist TADs together (red and green respectively).**(B)** Experimental and simulated contact frequencies for the Tsix and Xist TADs. The model reproduces the two TADs and the weak contact frequencies between them [197].**(C)**: (a) Computational algorithm to determine the potentials that reproduce the experimental 5C contact map. (b) The interaction potential is the sum of two terms shaped as spherical wells of energy *B*_*ij*_. (c) the chromatin ber is modeled as a chain of beads, each representing a segment of 3 kbp along the chromatin ber. following each iteration of the algorithm, a Monte Carlo sampling records the conformations computed from the energies *B*_*ij*_. (e) Contact map back-calculated from the sampled conformations. Experimental contact map returned by the 5C experiments, where the Tsix and Xist TADs can be identified and compared with the back-calculated map through evaluation of *χ*^2^ (the squared dis-tance between the experimental and numerically estimated contact probabilities). *χ*^2^ is minimized by updating the energies *B*_*ij*_ and re-injection to compute the conformations sampled in (d). Monte Carlo sampling is carried out and the algorithm is iterated until the sequence of *χ*^2^ converges. (g) Simulated vs. experimental contact maps for the Tsix and Xist TADs in mouse ESCs (adapted from [91]). Arrowheads indicate the frequent interactions between Linx, Chic1 and Xite described in [197] (adapted from [270]).**(D)** Sample conformations from the optimized simulation shown in (B), compartmentalization of the model ber into two separated domains corresponding to the Tsix and Xist TADs despite extensive structural variability (adapted from [91]).

In this model, the chromatin fiber is coarse-grained as a chain of identical beads (108 for the Tsix TAD and 199 for the Xist TAD), separated by distance *a* = 53nm, which represents 3kb of genomic sequence (Fig.33A) [270]. This length is the average size of HindIII restriction fragments in the 5C data set in [197]. The original 5C data, based on pairs of interacting forward/reverse restriction fragments, is converted into a list of interacting bead-pairs sequences. The chromatin fiber is assumed to be at equilibrium (at least locally on the genomic length scale of a TAD), so that the probability of a chain configuration is a Boltzmann distribution. Each pair of monomer interacts with a spherical-well potential of hardcore radius *r*_HC_ = 0:6 *a* and range *R* = 1.5 *a*. The value of *R* and *r*_HC_ were determined to give the best agreement between calculated and experimentally observed contact probabilities during the optimization procedure of the potentials *B*_*ij*_ (Fig.33C).

The output of the simulations based on the potentials constructed from the coefficients *B*_*ij*_ [270] is used to produce polymer configurations compatible with experimental data (DNA FISH)[91]. Cell-to-cell variations in chromosome structure are correlated at a single cell level with the ones in transcription levels of Tsix and other transcripts within the same TAD [91]. These variations predict genetic modifications such as the deletion of beads from the polymer, which are verified experimentally. Finally, interactions within each TAD contribute to the stability of the boundary between them. To conclude, a common effective long-range potential acting on dispersed binding monomer sites along a polymer chain is sufficient to generate micro-segregation, TAD and sharp boundaries [241]. However, this long-range interaction do not have yet a direct physical support and TAD can also result from long-range interactions due to binding proteins position at random position inside TADs.

It might be possible that TADs can be generated by different physical mechanisms. Supercoiling induced during transcription [147] or unconstrained supercoiling in chromatin [191,146] can be generated in models by taking a rod, twisting it ∆ *Lk* times and gluing the two ends together. The resulting loop has to coil in order to relieve tension [36]. TADs could also emergence from supercoiling and in between boundary elements, there are sections of unconstrained chromatin which is super-coiled. In this model, the axial rotation of chromatin fibers is limited, although the anchoring nuclear granules are free to move. Strong supercoiling with ∆ *Lk* = -8 per 400kbp reproduces contact probabilities occurring within individual TADs and between neighboring topological domains in undifferentiated embryonic stem cells [197]. Furthermore, weak supercoiling (∆ *Lk* = -1 per 400kbp) is used to reproduce the contact probabilities in differentiated cells [23].

### 6.4 Chromatin loops extrusion as a mechanism for chromosome segregation and TADs

Mitotic chromosomes are condensed by enzymatic proteins (condensin) that actively extrude chromatin loops [190,6]. Condensin and cohesin complexes play a central role in chromosome compaction [231,115,185], but direct experimental evidences are still lacking. The enzymes link two adjacent sites and move both contact points along the chromosome in opposite directions, so that they progressively bridge distant elements stalling at specific sites (CTCF). A polymer model was constructed based on loop-extruding factors (LEFs), where two heads bound to nearby sites along the chromatin ber can slide away from each other [92], leading to a loop extrusion ( g.36a). When the heads of two neighboring LEFs collide, they block each other and stop. LEF can dissociate with a certain rate independent of their state and location and rebinds immediately at a random site elsewhere on the chromosome, where it resumes extruding a new loop (Fig.36a). Thus, LEFs can generate a tightly stacked loop array.

Numerical simulations shows that the steady-state does not dependent on initial configurations.

Using this scenario, loops, sub-TAD contact domains and TADs are generated through loop-extruding factors (Fig.36). Upon deletion of a domain boundary, topological domains extend beyond their boundary. The encounter probability map of CTCF binding sites contains also isolated peaks [214,280]. This scenario is in contrast with the looping interaction models, mediated by protein binding used to model TADs formation, as discussed in previous sections see also [84,92].

Furthermore, numerical simulations of polymer with LEFs reveal two distinct steady-states of loop-extrusion dynamics: at low LEF concentration, polymers are poorly compacted, where loops are formed by single LEFs and separated by gaps. At high concentration, the chromosome is compacted into an array of consecutive loops, having multiple LEFs (Fig.36c). For a LEF abundance of one condensin per 10 − 30kb, the loop sizes are consistent with the ones inferred from Hi-C data [192]. These results further suggest that chromosome compaction occurs in the dense regime (Fig.36d). Large scale Hi-C data of contact maps of the human genome with a 1kb resolution [214] reveals the existence of 10,000 loops with an average size of 185 kb, occurring between CTCF motifs. The encouter probability *P*_enc_ peaks at 5kb and thus the Kuhn length is less than 5kb, measured experimentally in [231]. The decay of *P*_enc_ ~ *n*^-1.27^ is computed from data, which is faster than previously suggested [157]. Within a topological domain, the decay is slower (*P*_enc_ ~ *n*^-0.75^) as expected due to higher interactions (see 5.4.2).

Recently a new method was developed to construct a minimal polymer model from the encounter probability (EP) distribution between genomic sites represented in a large matrix [246]. Although this matrix is obtained by averaging the EP over cell population, the TAD diagonal blocks contains hidden information about sub-chromatin organization. To construct the polymer, the EP decay is used in two steps: first to account for TADs, random connectors are positioned inside a restricted region defining the TADs. In the second step, the long-range frequent specific genomic interactions in the polymer architecture are taken into account for deriving from the EP matrix the strength of the interaction. Interestingly, only a small number of randomly placed connectors are required to reproduce the EP of TADs. The mathematical difficulty solved in [246] is to determine the strength and the number of the connectors, so that the EP of the data and the simulations have same decay exponent *α* in the formula *P*_enc_ ~ *n^α^* (Fig. 34). The advantage of this model is that connectors can be directly interpreted as direct molecular binding.

**Fig. 34:**
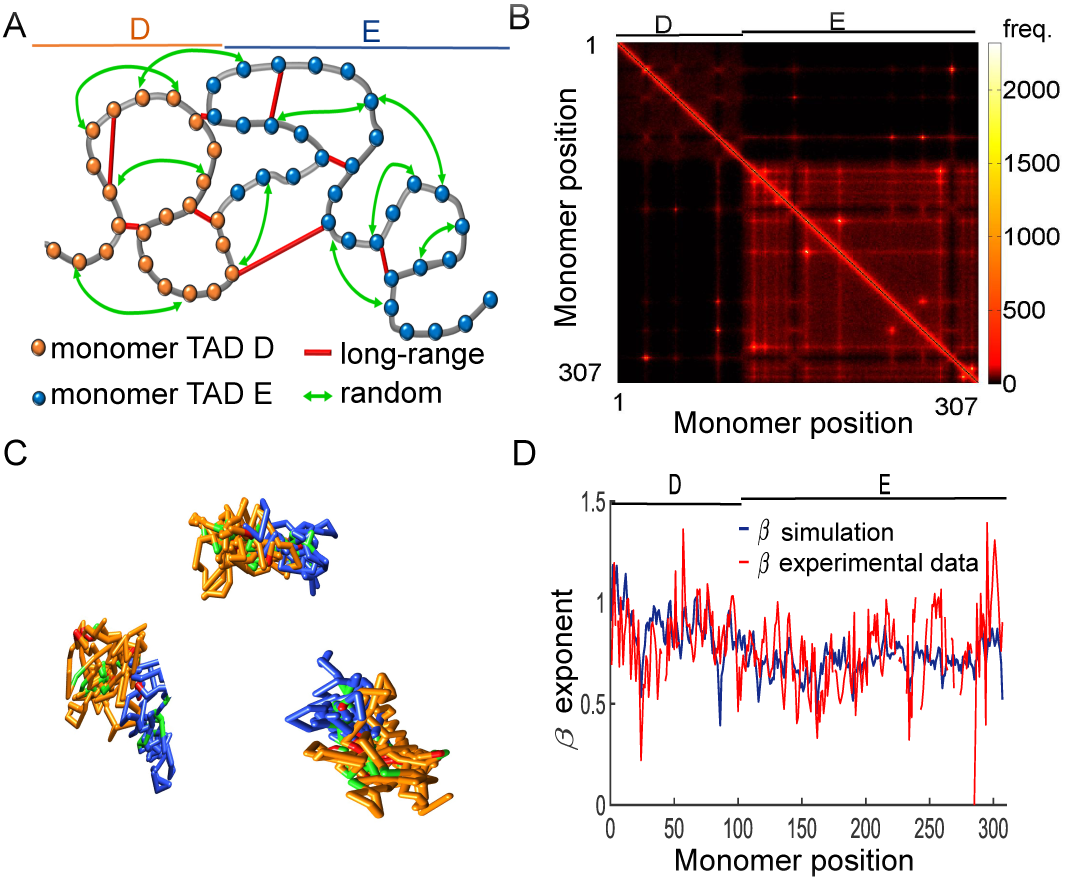
Coarse-grained reconstruction of chromatin using extracted random loops and connectors peaks present in 5C data. **A.** Schematic polymer model where TAD D (orange, monomers 1-106), and TAD E (blue, monomers 107-307) are recovered from random loops (green arrows) according to the connectivity *ζ* and fixed connectors (red bars) corresponding to the specific peaks of the 5C data.**B**. Three realizations of the polymer model.**C** Encounter frequency matrix of the simulated polymer model, showing two TADs, and the off-diagonal points correspond to the xed connectors.**D**. Comparison between *β* computed from experiments and simulations data, con rming that the present polymer model accounts for the statistics of encounter frequency distribution.

Following the construction of a polymer model, stochastic simulations can be used to compute the distribution of first time and the conditional encounter probability of any genomic sites to meet. Using a polymer with 307 monomers, the encounter time and the probability were estimated for monomer 26 (position of the Linx) to meet monomer 87 (Xite) before monomer 64 (Chic1). In this coarse-grained model, each monomer represents three key sites on the X chromosome [91] located inside TAD D. Three polymer realizations indicate the location of the three sites inside TAD D (yellow) and E (blue), that do not mix (Fig. 35). Numerical simulations reveal that the encounter probability is *P* = 0.55, while the mean times are quite comparable of the order of 131s (see table C in Fig. 35). Interestingly this approach can be used to check the impact of TAD E on the mean encounter time inside TAD D: after removing TAD E, stochastic simulations (Fig. 35D) show that the encounter probability was inverted compared to the case of no deletion: the mean encounter has increased by almost 50% to 195s (Linx to Chic1) and 205s (Linx to Xite). This result suggests that TAD E contributes in modulating the interaction probability and the mean time and thus further indicates that the search time inside a TAD depends on neighboring chromatin configuration, due to interaction between TADS present in the data. These encounter times reveal how chromatin can self-regulate. The present polymer construction is generic and can be used to study steady-state and transient properties of chromatin constrained in 5C data.

**Fig. 35:**
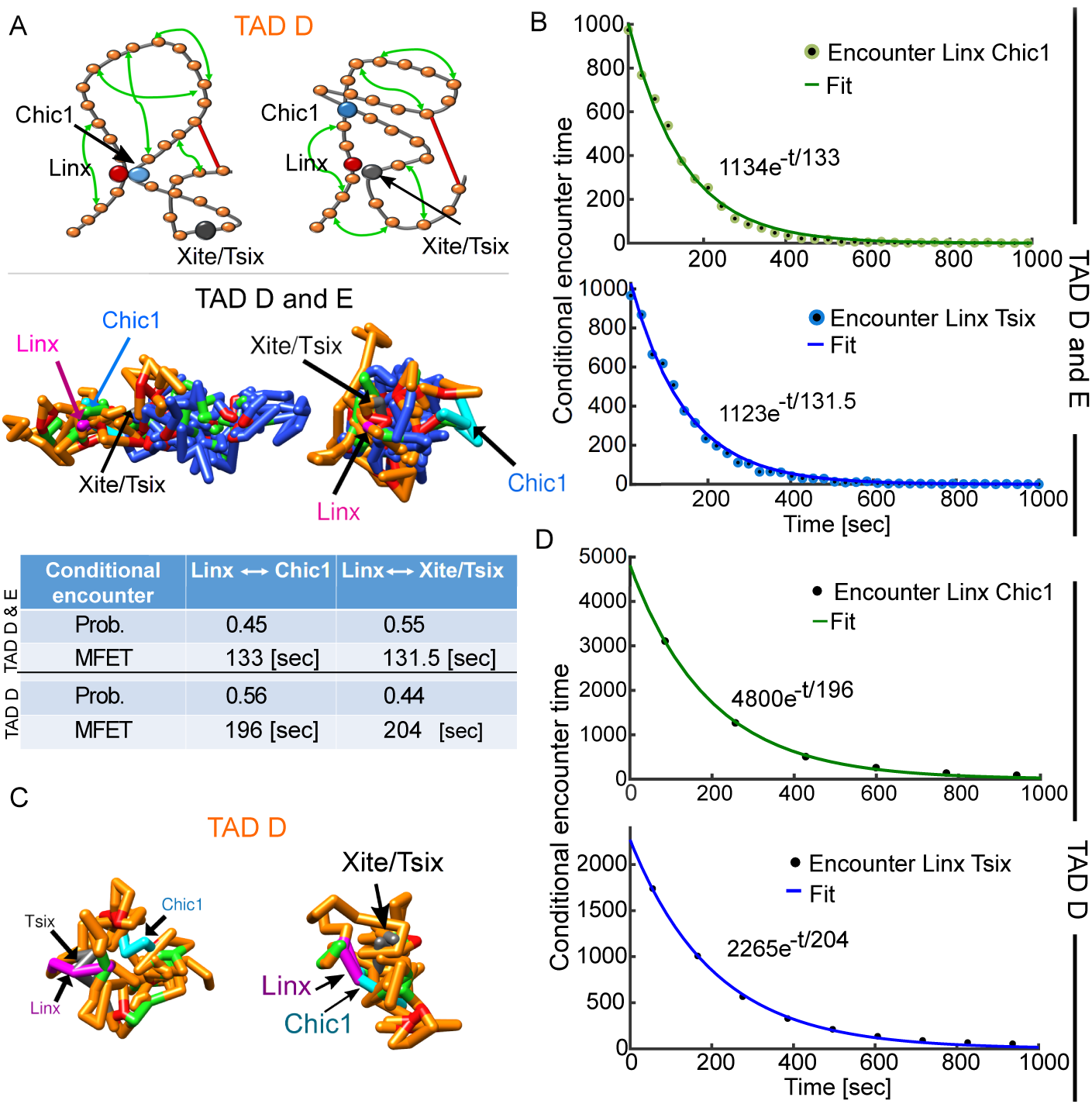
Transient properties of the chromatin: Conditional mean time and probability for three sites to meet. **A**. (upper panel) Representation of the polymer model for TAD D (orange, monomers 1-106), where loci Linx (bead 26, red) meets Chic1 (bead 67, cyan) and Xite/Tsix (bead 87, gray) respectively. Random loops (green arrows) and specific long range-connectors (red bar) are added following the connectivity recovered from data [246]. Fixed connectors (red bars) correspond to the specific peaks of the 5C data. Three realizations (bottom panel) of the polymer model containing TAD D and E, showing the encounter between Linx (magenta) and Chic1 (cyan) and Xite/Tsix (gray) respectively. The color code is from the upper panel.**B**. Histogram of the conditional encounter times between Linx and Chic1 (upper panel, green), and Linx and Xite/Tsix (bottom panel, blue) with TAD D and E.**C**. Two polymer realization with a single TAD D (bead 1-106, orange) showing the encounter between Linx (magenta) and Xite/Tsix (gray, left panel), and the encounter between Linx and Chic1 (cyan, right panel).**D**. Histogram of the conditional encounter times for a polymer with only TAD D, showing an exponential decay as in sub gure C.

To conclude, the models described in this section assume that the polymerber is at equilibrium. However, one fundamental difference between metazoan and yeast genomes is that while yeast chromosomes can equilibrate during one cell cycle, metazoan chromosomes are so long that they are unable to unmix in the time interval between mitosis [223]. Hence, it is possible that TADs are structures generated during mitosis. The prediction of the models discussed here remain to be tested, especially for the formation and maintenance of TAD boundaries. Coarse-grained polymer models certainly provide a theoretical basis to analyze Hi-C data. As these data become more ubiquitous and the spatial resolution increases, more accurate models would be required to describe this increasing complexity.

**Fig. 36.**
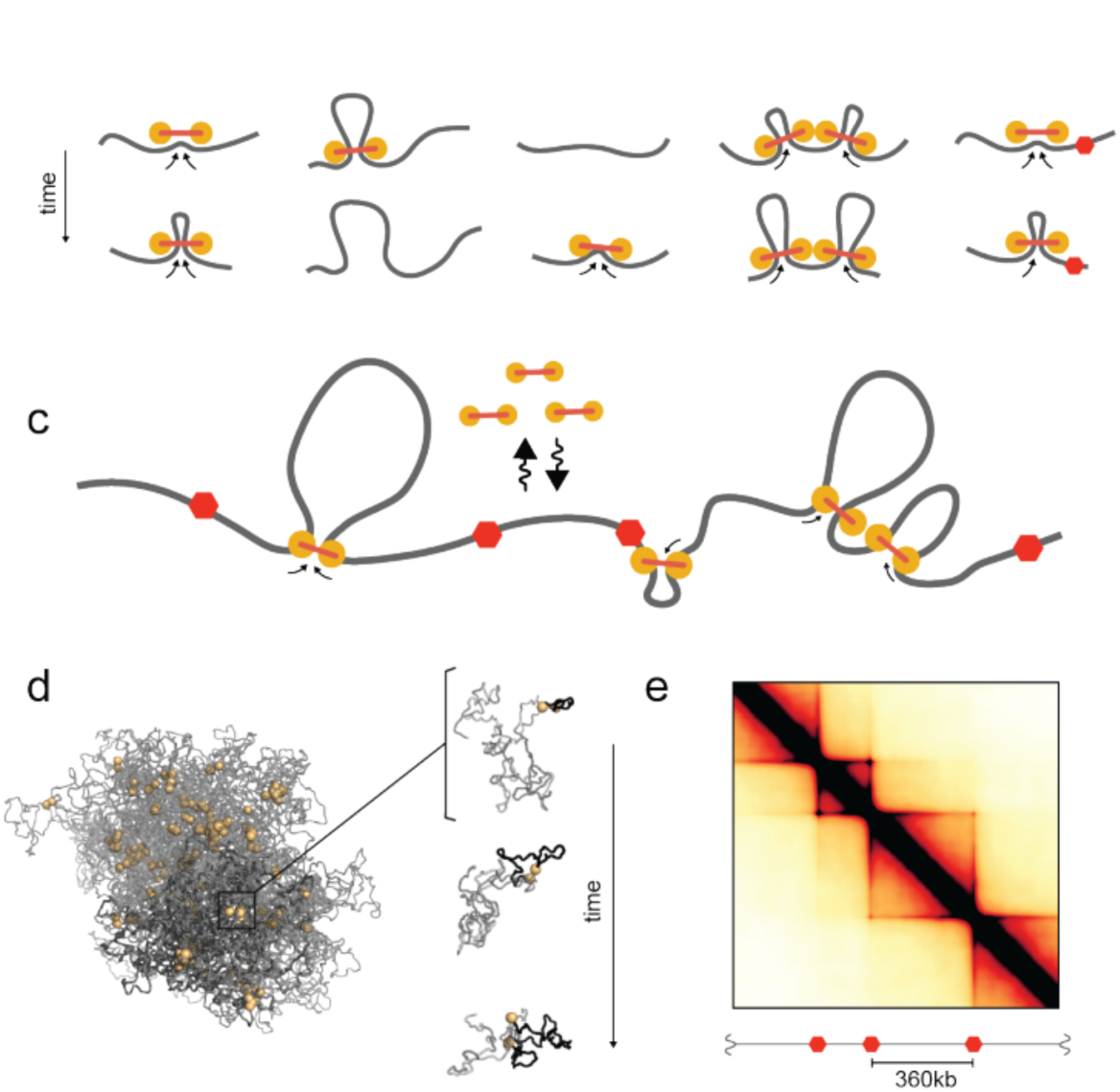
Chromosome partitioning into TADs. (a) Model of LEF dynamics, LEFs shown as linked pairs (yellow circles), chromatin ber (grey). From left to right: extrusion, dissociation, association, stalling upon encountering a neighboring LEF, stalling at a BE (red hexagon). (b). Schematic of LEF dynamics. (c) Conformations of a polymer subject to LEF dynamics, with processivity 120kb, separation 120kb. Left: shows LEFs (yellow), and chromatin (grey), for one conformation, where darker grey highlights the combined extent of three regions of sizes (180kb, 360kb, 720kb) separated by BEs. Right: progressive extrusion of a loop (black) within a 180kb region. (d) Simulated contact map for processivity 120kb, separation 120kb. (adapted from Fudenberg et al. [84]).

## 7 Coarse-graining polymer models for analyzing telomere organization in yeast nucleus

Telomere organization is a specific characteristic of the nucleus, in addition to the nucleolus, centromeres and different bodies (PML, Cajal) [265]. In budding yeast, the 32 telomeres of a haploid cell can associate in several clusters [94,263,225,124] as shown inFig. 37. Although a single telomere cannot be continuously monitored, the formation of telomere clusters observed *in vivo* have opened the debate of their role, function and whether these clusters resulted from random encounter without physical interactions or from a dissociation-aggregation process.

**Fig. 37.**
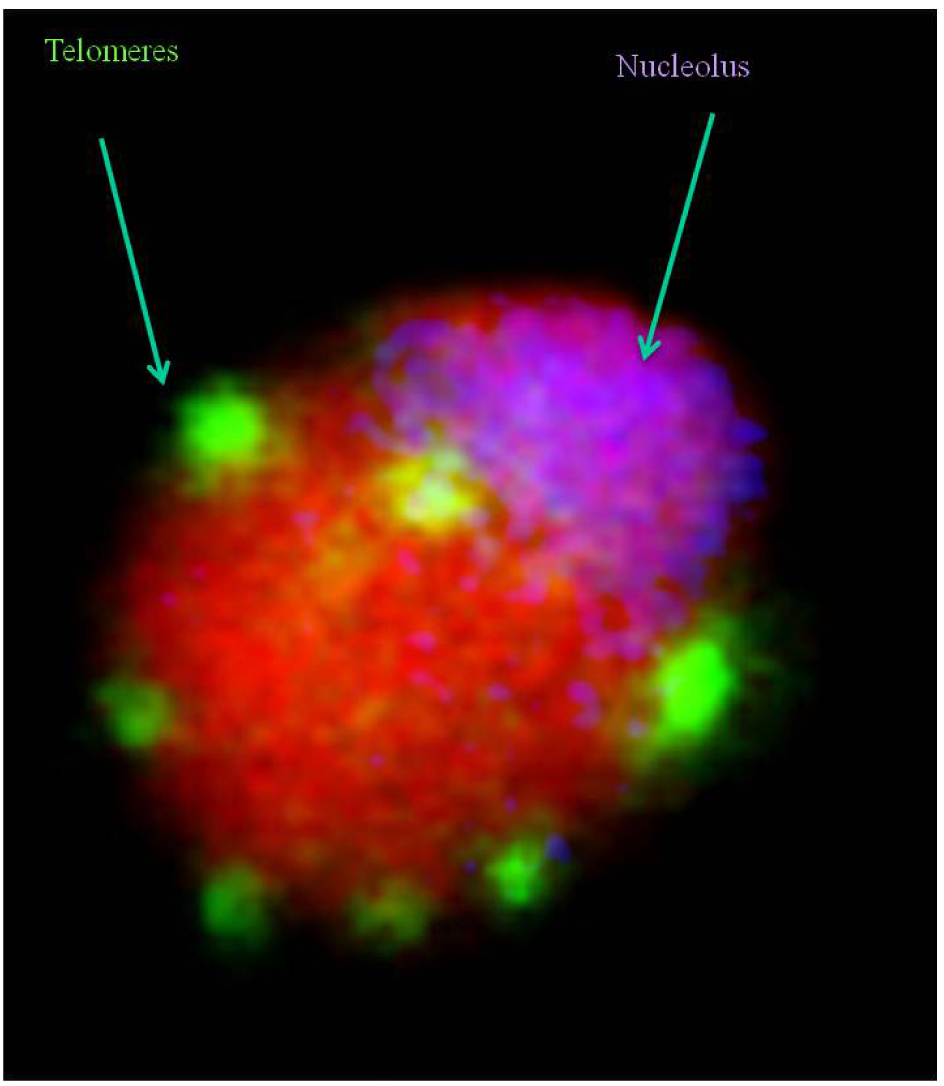
64 telomeres of diploid cells. Telomeres are clustered of 8 to 10 perinuclear foci on which silencing factors concentrate (reproduced from [94]).

In yeast, telomere dynamics lead to the formation of two to six clusters, composed of an average of four telomeres [225]. Telomere foci undergo fusion and fission events over a time-scale of minutes [235]. Numerical simulations underlying telomere dynamics are realized by adding structural constraints, including chromosome structure, attachment to the spindle pole body (SPB), and nuclear crowding [269,297], suggesting that clusters are due to transient encounter.

Although telomere foci observed *in vivo* could result from random encounters of telomeres at the nuclear periphery, stochastic simulations of non-interacting telomeres have shown that this hypothesis would contradict the stability of foci observed invivo, where telomere foci are stable over minutes ([235] and Fig. 1G, Movie S1 and S2 of [124]). In numerical simulations, telomere dynamics can be coarse-grained to 32 independent Brownian particles, diffusing on the two-dimensional nuclear envelope of radius 1μm with a diffusion constant *D* = 0:005 *m*^2^ *=s* [39]. The nucleolus region which occupies about 1/3 of the total surface is excluded. This result is in contrast with previous simulations based on microscopy images of fluorescently tagged telomeres, where two telomeres located at a distance *<* 0:3 *m* were considered to be part of the same focus [297]. Thus, if telomere clusters cannot result simply from the transient encounter of independent telomeres moving by random motion, clustering dynamics and stability are due to aggregation-dissociation processes (see discussion below).

### 7.1 Encounter rate of telomeres

Coarse-graining the motion of one end of a Rouse polymer (Fig. 38A) into a single Brownian particle moving on the surface of a sphere was recently used to study telomere clustering. Indeed the arrival time of a single telomere moving on the surface of the nucleus to a cluster (which occupies a small fraction of the surface) is a rare event, taking a long time [239]. This coarse-graining approximation was validated by Brownian simulation of a Rouse polymer, which models the chromosome dynamics (Fig. 38). In this simulation, the telomere motion occurs on the surface of a sphere, which represents the nuclear surface, whereas the remainder of the monomers in the chain evolves inside the ball (modeling the nucleus). The distribution of telomere arrival times to a small target, which can be another telomere or a small cluster, is well approximated by a single exponential (Fig. 38B). The encounter time of telomeres at the nuclear periphery is characterized by the arrival time only, even though telomere motion involves complex polymer chains accounting for the physical chromosomal chain. Consequently, telomere clustering is mediated by the arrival of a chromosome to a small cluster and this process is modeled as Poissonian, as long as the chromosome arm does not restrict the motion of the telomere on the nuclear surface.

**Fig. 38.**
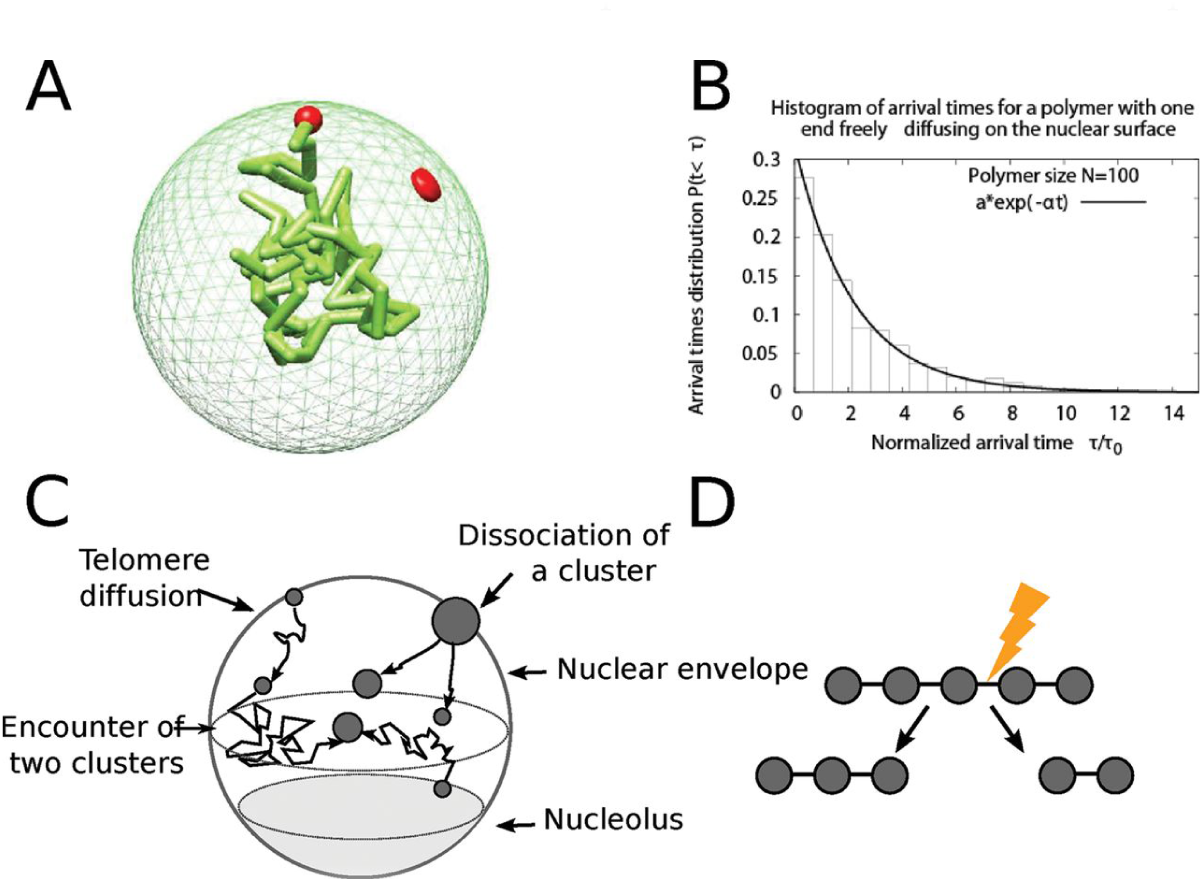
Computational model of telomere cluster formation. (A) Snapshot from a Brownian Dynamics simulation of a polymer with one end anchored on the nuclear surface. The polymer is composed of 100 monomers with average distance between monomers *l*_0_ = 50 nm, the nucleus is a reflecting sphere of size *R* = 250 *nm*. (B) Histogram of the arrival times for a polymer of size 100 monomers freely diffusing in the nucleus and one end constrained to the surface. A fit of the form *f*(*t*) = *a* exp (*bt*) gives *a* = 0:3, *b* = 0:44 *s*^1^. (C) Scheme of the diffusion-aggregation-dissociation model of telomere organization. Telomeres are represented as Brownian particles diffusing on the nuclear surface, and two telomeres coalesce with a rate *k*_*f*_ and a cluster of *n* splits at a rate (*n* − 1) *kb*. Schematic representation of the cluster dissociation model, where a cluster of *n* telomeres has *n* − 1 bounds. Any of these bounds can break at a rate *k*_*b*_and the cluster effective dissociation rateis (*n* − 1) *k*_*b*_ (reproduced from [124]).

In summary, the arrival time of a telomere to a cluster can be simulated using a Poissonian distribution approach. To study 32 telomeres (Fig. 38C), it is thus enough to run molecular dynamics simulations of Brownian particles on the surface of a sphere, except on the nucleolus region, where there are re ected. This process is characterized by the reciprocal of the mean encounter time *k*_*f*_ between two telomeres and dissociation process explain later on. The arrival time of a telomere to a cluster is Poissonian (Fig. 38C).

Molecular dynamics simulations of two stochastic particles on the surface of a sphere shows provide a mean of estimating the encounter rate of the order *k*_*f*_≈ 1.910^−3^ *s*^1^ r= 0.015 *μm*. Thus the encounter rate for two telomeres is *k_*f*._* for a target which is a disk radius Following an encounter, although telomeres form a cluster that in principle varies in size, because this size remains small, possible change in the scattering cross-section and motility, which could modify the forward binding rates [122] can be neglected. Thus the encounter rate between clusters or telomeres is approximated by a constant *k*_*f*_, independent of the size.

### 7.2 Dissociation of a telomere from a cluster

The dissociation of a telomere from a telomere cluster is still not clearly understood, because there are many possibilities for a particle to dissociate from an ensemble. This process has been described using the hypothesis that dissociations are single independent Poissonian events [122,124], where a cluster containing *n* telomere dissociate with a rate (*n* − 1) *kb*, where kb is the dissociation rate between two telomeres (Fig. 38D). In addition, dissociation gives rise to two clusters of random size p and n-p, and the probability is uniform. In the absence of speci c information about the molecular organization of a cluster, it is difficult to account for the local molecular interactions of telomeres in a cluster. This simple model of dissociation is independent of any polymer model, but could be added to any future polymer models. A dissociation model based on polymer model is still to be formulated, but should involve the SIR family of proteins (A. Taddei, private communication).

The clustering of telomeres was studied using numerical simulations as follow: free telomeres can bind together to form clusters with a forward rate *k*_*f*_ or dissociate with a backward rate *k*_*b*_. The ratio of these two constants defines an equilibrium, characterized by the ratio parameter *a* = *k*_*b*_ */k*_*f*_. When association and dissociation rates are Poissonian, the classical Gillespie's algorithm can be used to simulate telomere dynamics and study the cluster distributions. By comparing experimental distribution [225] with simulations [124], under the dissociation model described above, the dissociation rate of a telomere pair was found to be *k*_*b*_ = 2.310^−2^ *s*^−1^. Thus, the mean lifetime of a bond between two telomeres is 1 */k*_*b*_ = 43.5sec. Interestingly, under a genetic perturbation where a fundamental protein to link telomeres together (the silencing factor Sir3) has been overexpressed by 6- or 12-fold the endogenous level, then the backward rate changed to *k*_b_^−1^ = 109min for GALSp and *k*_b_^−1^ = 154min for GAL1p [124]. The exact molecular changes underlying these differences is still lacking.

### 7.3 Characterization of telomere exchanged between clusters

We have seen in the previous section that a combination of polymer models with live cell imaging led to the conclusion that telomere organization in wild-type cells results from several physical processes: aggregation mediated by direct interactions between telomeres, dissociation resulting from the separation from a cluster, and telomere random motion located on the nuclear envelope. Telomere Clusters are constantly reshaped due to binding and unbinding, leading on average to three to five detectable foci of four telomeres [94,225,122,124]. Although, cluster dynamics is characterized by the exchange rate of telomeres, two quantities are conserved in this process: the first one is the co-localization time *T*_*C*_, which is the time that two telomeres spend in a cluster (including cluster of two telomeres), when other telomeres can be exchanged (detached or attached). This time is not equal to the classic dissociation time, which represents the lifetime of a bond between two telomeres, because it accounts for all events of binding and unbinding leaving the two telomeres inside the same cluster. Using Brownian simulations in yeast, the co-localization time was estimated *T*_*C*_ 23.4 *s* [124].

The second conserved quantity to characterize clustering is the recurrence time *T*_*R*_, defined as the mean time for two telomeres to meet again in a cluster after they have just been separated. Two telomeres return in the same cluster in a mean time *T*_*R*_ (*T*_*R*_ is the sum of the time that both telomeres spend separately in different clusters plus the time to travel between clusters until they meet again forthe st time) of the order of 480sec, whereas they stay together in a given cluster for 23sec only [124], consistently with experimental observations [269]. The probability *P*_2_ to find two given telomeres in the same cluster (including in a pair) is equal to the ratio of times, *P*_2_ = *T*_*C*_ */*(*T*_*C*_ + *T*_*R*_), where *T*_*C*_ is the mean time that two telomeres spend in the same cluster. This probability is *P*_2_ = 0.047, consistent with the result of [269], where the probabilities for two telomeres to belong to the same cluster were determined experimentally to be mostly in the range 0.04–0.09 (Fig. 39).

**Fig. 39:**
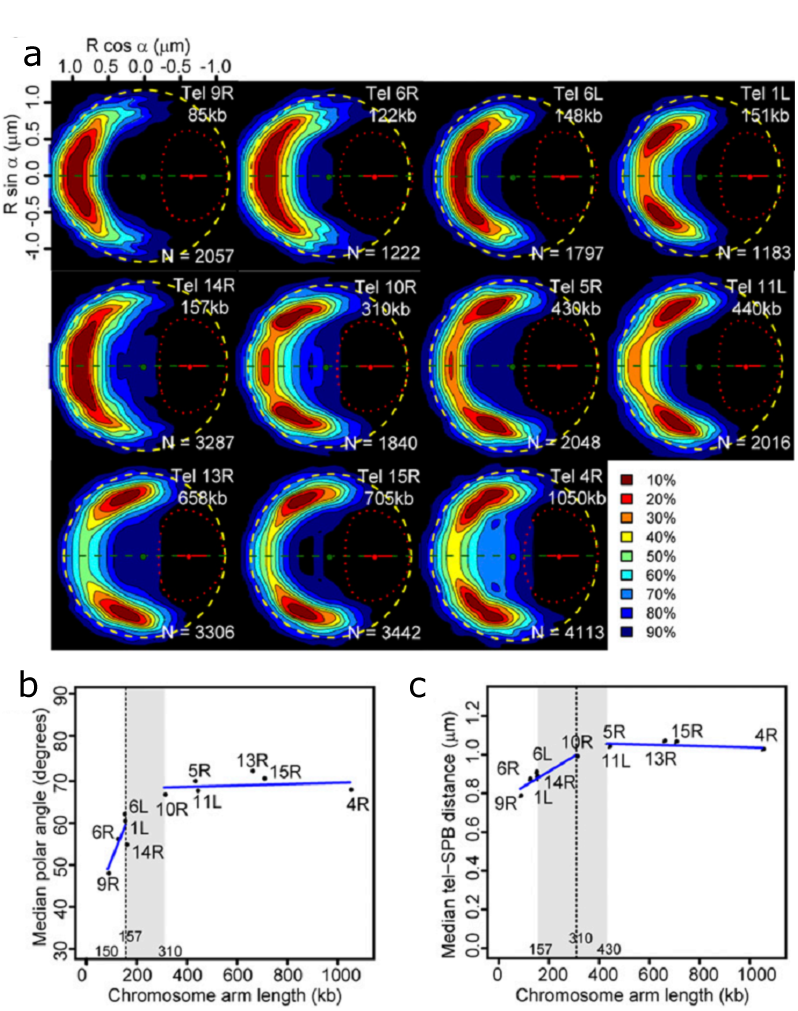
Arm length influences localization of subtelomeres at the nuclear periphery. (a) 2D localization probability maps for different telomeres. Top and bottom halves are mirrored (around the green dashed line) for visual convenience. Yellow circle and red dotted outline represent a median nuclear envelope and nucleolus, respectively. The probability to find a locus inside different regions of the nucleus is indicated by the percentage of the enclosing contour. Hot colors indicate higher probability densities. Chromosome arm sizes are indicated on the right (kb). *N*, number of cells analyzed. (b) Median polar angle vs. chromosome arm length, annotated with the subtelomere label. Dashed vertical line represents the change point value, gray area represents its 95 percent CI. Linear regression relationships are *Y* = 46.3 + 6.110^−2^ L (kb), for arm length *L* = 430kb and Y= 85.5+1.910^−2^ L (kb), for L larger than 430 kb. (c) Median subtelomere-SPB distances calculated as dist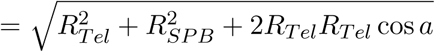where *R*_*Tel*_ is the distance of subtelomere to the nuclear center, *R*_*SPB*_ is the distance of SPB from the nuclear center = 900 nm, and *a* is the angle with the horizontal axis. Dashed vertical and gray areas are defined as in b (reproduced from [269]).

In summary, coarse-grained polymer models with live cell imaging led to the conclusion that telomere organization in yeast wild-type cells results from three physical processes: aggregation mediated by direct interactions between telomeres, dissociation resulting from the separation from a cluster, and telomere random motion located on the nuclear envelope. Clusters are constantly reshaped due to binding and unbinding, leading on average to three to five detectable foci of four telomeres. An exact model of telomere dissociation in yeast is still missing. Including super-resolution imaging information about the local geometry would be necessary. It would also be interesting to study telomere dynamics in other species.

## 8 Single particle trajectories of a chromatin locus

In this section, we describe the statistical analysis of Single Particle Trajectories (SPTs) of a chromatin locus. Many of the basic cell functions such as transcription, repair and division can now be monitored in vivo, but the interpretation of the trajectories remains difficult due to all mechanisms involved in generating the motion. There is also a large unexplained variability. We first describe how SPTs data are acquired and in the second part, we present recent polymer models and their statistics used to interpret data and extract biophysical parameters.

### 8.1 Live imaging of nuclear elements

The data acquisition for the motion of a chromatin locus is based on using a green uores-cent protein (GFP), extracted from luminescent jellyfish Aequorea victoria. This protein is excited using wavelengths 395nm and 475nm *in vivo*. In that case, processes such as transcription [43] can be monitored in cells. This discovery had a huge impact in cell biology, revealing cell function at a molecular level, as described by The Tsien's 2008 Nobel lecture http://www:nobelprize:org/nobelprizes/chemistry/laureates/2008/tsien-lecture.html).

A genetic construction designed to visualize a given locus is to integrate a Lac operon at different positions in the yeast genome [220] and using the LacI protein bound to a GFP (making a LacI-GFP complex). This type of molecular construction (Fig. 40a) permits a chromatin locus to become visible (Fig. 40b) and for example sister chromatin separation in yeast can be visualized [180]. When a GFP is fused to nuclear pore proteins (Nup49p) located in the nuclear membrane [111], the motion of the nuclear envelope can be monitored. When both nuclear pores and a locus are labeled, their relative motion can be observed. Once nucleus elements are labeled, high resolution microscopy [243] is used to monitor DNA locus motion at tens of nanometer precisions. Another issue of image acquisition is the possible motion, division of cells or their change to an environment, that should be accounted for in single locus analysis. For example, when a protein diffuse with a diffusion coefficient of the order of 1 *μ*m^2^ */*sec [126] (although there is large variability depending on the protein), the diffusion of a chromatin section can be three orders of magnitude slower [170]. Thus, capturing cellular processes requires a time resolution of seconds or below.

**Fig. 40:**
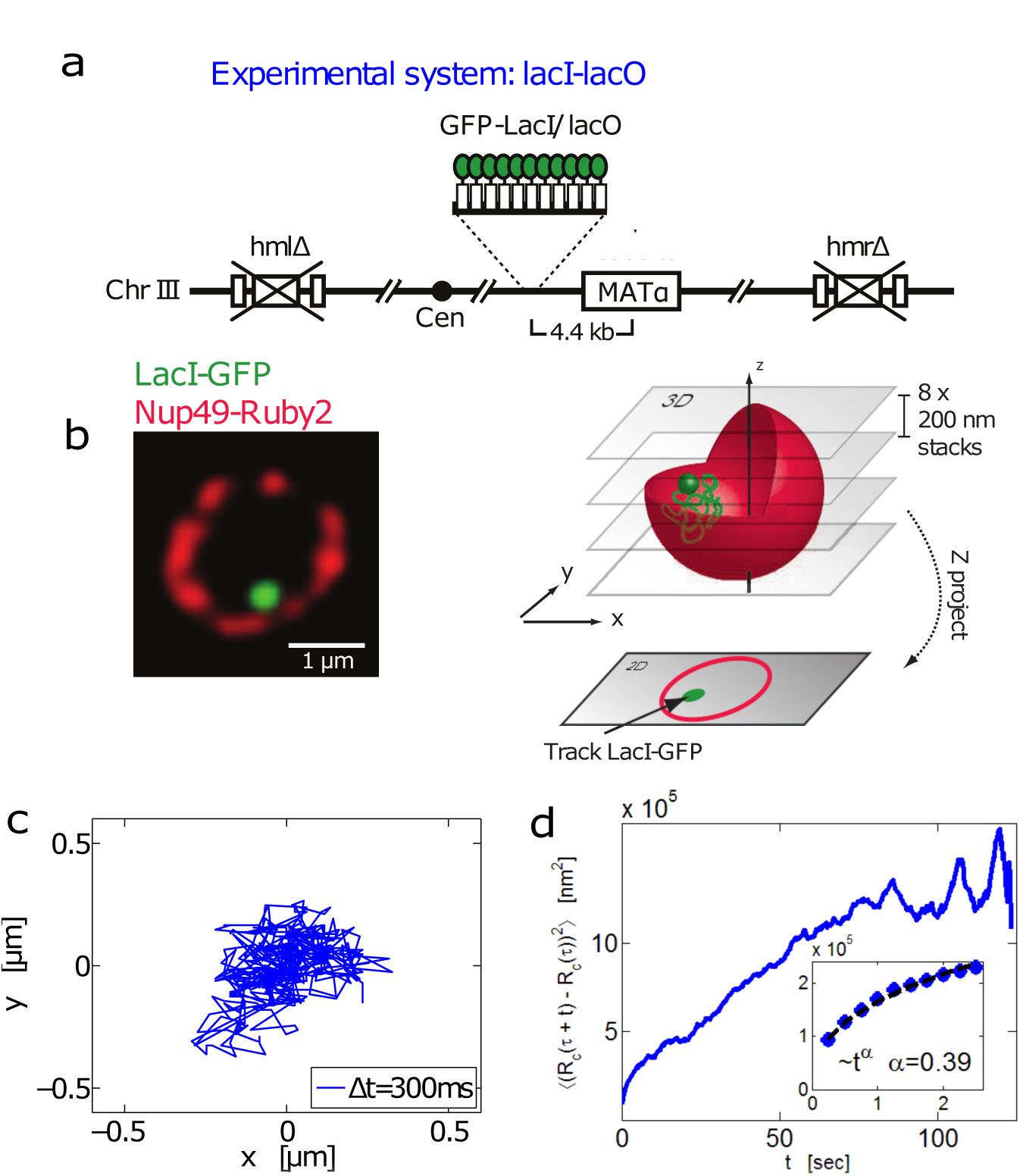
Imaging a chromatin locus. (a) LacI-lacO system [188]: Galactose induces a DSB at MAT locus in yeast in a haploid strain that bears lacO sites 4.4 kb from MATa and expresses GFP-lacI and Nup49-GFP fusion proteins (GA-1496). (b) MAT locus in yeast (green) while the nuclear membrane (red) was marked with the nup49-mCherry fusion protein (left). The nucleus is imaged in 8 stacks along the z-axis and trajectories are projected on the xy-plane. (c) A trajectory of the MAT locus taken at a time resolution of 300msec during 300 frames. (d) Correlation function *C*(*t*) eq.198 of the position depicted (from c). At short times, the sub-diffusion regime is characterized by an anomalous exponent α = 0.39 (inset magni cation at short times) (reproduced from [15]).

Once nucleus elements are labeled, high resolution microscopy [243] is used to monitor DNA locus motion. Using deconvolution procedure based on the Point Spread Function (bright fixed spot spread of a Dirac) [86,177], single particle trajectories are acquired, but contain a noise due to the imprecision in the location. Deconvolution procedures further introduce various localization errors that must be differentiated from physical effects [116]. Although tracking procedures remain an image processing challenge, some routine algorithms are now implemented in freely available softwares such as ICY [56], ImageJ [234,232] or Volocity [151].

### 8.2 Statistics analysis of SPTs

The ability to follow single locus located at different positions on chromosomes [39] revealed the heterogeneity of the nuclear organization. In the yeast *Saccharomyces Cerevisiae*, chromosomes are non homogeneously distributed, but organized in a Rabl con guration [213] (Fig. 41a), where centromeres are attached together. Similarly, telomeres (ends of the chromosomes) forms clusters on the yeast membrane [265]. In these conditions, the statistics of SPTs of chromatin locus re ect the nuclear heterogeneity. Trajectories can be restricted to a small region ( g.41b-i where the con ned ball has a radius of 221nm). In fig.41b-ii, although the recording duration is the same as in fig.41b-i, the trajectory was contained in a smaller ball of radius 164nm. The large heterogeneity of loci behavior across a cell population suggests that the loci can experience different interaction over a time scale of minutes. Analysis of chromosome loci SPTs reveals (Fig. 41b) the local nuclear organization: subtelomeres are positioned at the nuclear periphery, depending on the chromosome arm length, centromeres are attached to the microtubule organizing center (the SPB), and the volume of the nucleolus, which acts as a physical barrier.

**Fig. 41.**
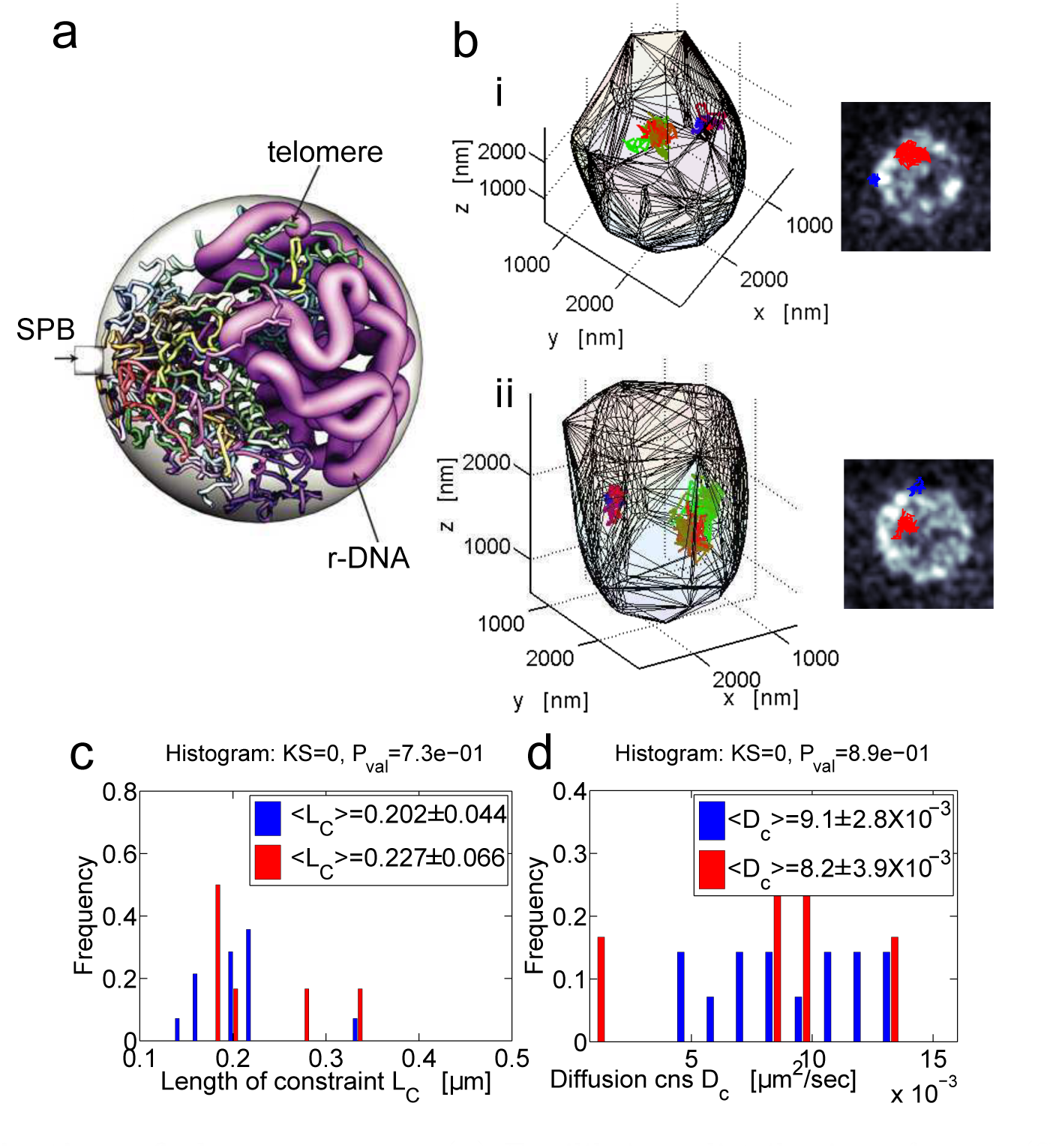
Organization of chromosomes. (a) Equilibrium distribution of yeast (Saccharomyces Cerevisiae) chromosome, from Monte-Carlo simulations of polymer [291]. (b) Recorded for 100 seconds of two trajectories of a chromatin locus (upper gures-i, ii) and (bottom gures-iii, iv). Time resolution ∆ *t* = 0:33 *s*. At the beginning the trajectory is red and gradually becomes green (locus), while the SPB starts red and gradually becomes blue. The trajectory of the locus (red) and the displacement of the SPB trajectory (blue) inside the nucleus is projected on a plane. The nuclear membrane is reconstructed based on the nup49-mCherry fusion protein. The position and dynamics of the MAT locus in yeast nucleus: (c) Distribution of the constrain length (*L*_*C*_) estimated from eq.201 (100 sec) trajectories when the cells grow in glucose (blue) or galactose (red). (d) Distribution of the diffusion coefficients for different cells (eq.202) (reproduced from [18]).

To extract biophysical information from SPTs of a chromatin locus, a physical model of the locus motion is needed, based on the local organization of the medium, paved with obstacles and local forces. The classical Langevin'sequation 12 and its overdamped approximation 13 usually provides such framework.

The most common statistical estimators to study stochastic trajectories are the Mean Squared Displacement (MSD) and the cross-correlation function, defined for a time series *R*_*c*_(*t*) (locus position at time *t*) by [237,211]

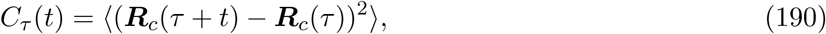

where 〈.〉 denotes the average over ensemble realization. When the data contains enough recursion, the ergodicity assumption says that the function *C*(*t*) can be also computed along a trajectory from the empirical estimator

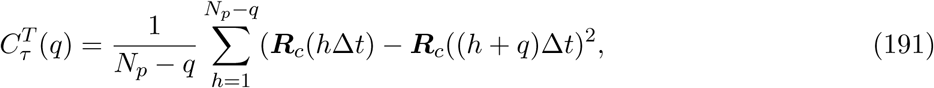

for *q* = 1 *,..,N*_*p*_ −1, where *t* = *q*Δ *t* and *T* = *N*_*p*_Δ *t* is the duration of the entire trajectory.Fig.40c–d show the function *C*(*t*) computed for a chromatin locus. Similarly the MSD of a trajectory is de ned by

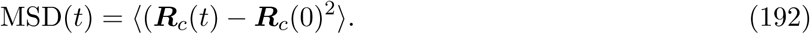

The MSD is by definition computed by averaging over *M* different trajectories and by taking the limit

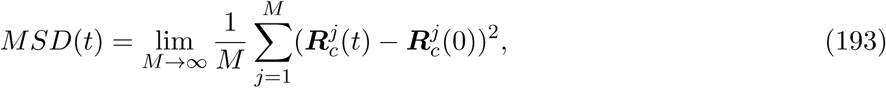

Which again assumes that the space is homogeneous. For a Brownian particle moving in R^*d*^ space, characterized by the diffusion coefficient *D*, the MSD is MSD(*t*) = 2 *dDt* [237]. While *C*(*t*) is linear in time for a free Brownian particle, it reaches an asymptotic value when the particle is moving in a con ned space Ω, with an exponential rate. Indeed, the MSD is expressed in terms of the pdf *p*_*t*_, solution of the Fokker-Planck equation with re ecting boundary conditions on *−*Ω and equals to the Dirac-distribution at *t*=0: *p*_0_(*x*, *R*_*c*_(0)) = δ(*x*− *R*_*c*_(0)),

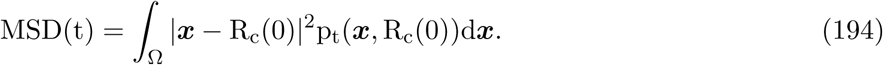

Using an expansion in eigenfunctions 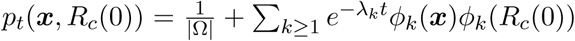 where λ_*k*_are the eigenvalue of the Laplacian,

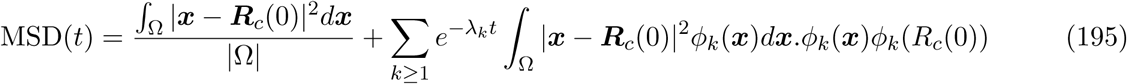

For a disk, 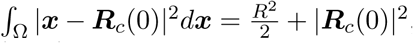. The *α*_*nm*_are the ordered m-positive roots of 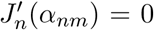 which is the Bessel function of the first kind of order *n*[26]. The long-time asymptotic for the average MSD(t) over the initial point is approximated by truncating the sum to the first term:

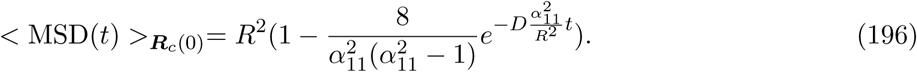

For a general domain, the influence of the domain geometry on the MSD is far more complicated.

For a particle con ned in a harmonic potential by a spring force with constant *k* [237] and friction coefficient γ, the stochastic description is OU-process

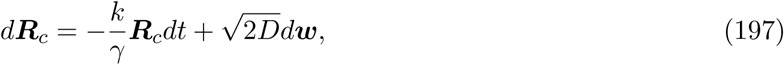

and the cross-correlation function is

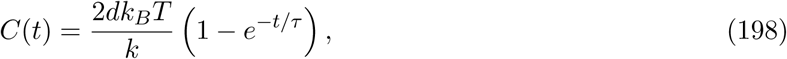

with τ= γ/κ, while

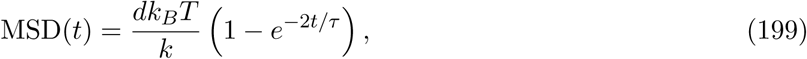

where τ = k_B_ *T* / *Dκ*. We note that the auto-correlation function for an OU process is

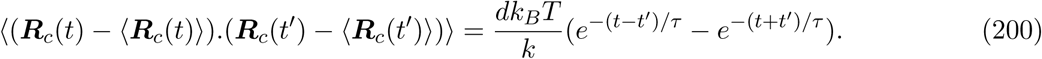

In summary, when the correlation function *C*(*t*) converges to an asymptotic value, the underlying motion can either be restricted by a tethering force or by obstacles.

### 8.3 Second moment statistics of SPTs

The MSD was historically computed by averaging the MSD over a cell population and the effective diffusion coefficient is obtained by evaluating the tangent of the MSD at the origin. Due to the large heterogeneity across cells, averaging over a cell population does not reflect diffusion in a single cell. The con nement radius *R*_*C*_ of an abstract disk, where the locus is con ned can be extracted from formula 195 as the maximum of the MSD. The two parameters (*D* and *R*_*C*_) have been estimated with and without a dsDNA break [69] and also when some proteins have been deleted [170]. For the yeast *HIS3* locus [193], the diffusion coefficient is 〈 *D*〉 = 2.1 ± 0.2 × 10^−3^μm^2^/sec, while the radius of confinement is 〈 *R_*c*_*〉 = 0.61 ± 0.3μ *m*. After treating the E. Coli DNA with sodium azide, which inhibits ATP synthesis, the diffusion coefficient of a DNA locus is reduced [283]. Thus, these estimators can be used to characterize the dynamics and localization of a chromatin locus. The extraction methods of the two parameters are mostly relevant to analyze particles described as Brownian, but is it insufficient for a DNA locus moving a con ned environment? we shall explore these questions in the next sections.

#### 8.3.1 Constraint length of a locus

A locus localization can be estimated using the standard deviation (SD) of the position with respect to its mean averaged over time. This SD is a characteristic length, called the constraint length *L*_*C*_, estimated by the empirical sum

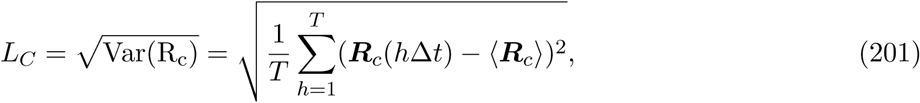

Where 〈 *R_*c*_*〉 is the average position of the locus along the trajectory. For a MAT locus in yeast, where trajectories are recorded for hundreds of seconds, the distribution *L*_*C*_ is shown in g.41c: the histogram of *L*_*C*_ shows a large variability across cells with an average 〈 *L*_*C*_〉 = 0.202±0.444μm, when the yeasts grew in glucose and higher in galactose.

#### 8.3.2 Effective diffusion coefficient of a chromatin locus

The effective diffusion coefficient of a locus can be computed directly from the time series [237] using the estimator

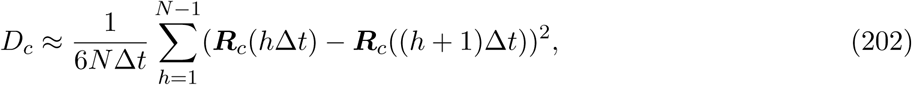

Where *N* is the number of time frames. For short time Δ *t*, the locus motion is described as Brownian (subsection 3.10.2) and the diffusion coefficient is computed from eq.202 [11]. Reducing the time step Δ *t* provides a better estimate of the local diffusion coefficient *D*_*c*_. Applying formula 202 on MAT locus trajectories leads to a distribution of diffusion coefficients (Fig. 41d) with a mean over cell population 〈 *D*_*c*_〉 = 9.1 ± 2.8 × 10^−3^μm^2^/sec, when the cells grow in glucose [18]. When the motion is not driven by diffusion only, the estimator 202 should be replaced by the correlation function (relation 191) and other parameters can be extracted by tting a curve *Atα*. The interpretation of *A* remains unclear, while characterizes the anomalous behavior (see section 3.10.2).

### 8.4 Anomalous diffusion of a chromatin locus

The motion of a tagged monomer, part of a polymer, is described as subdiffusion for an intermediate time scale (section 3.10.1). As we discusses above, this description applied to a tagged locus located in a chromosome chain. The correlation function (eq.191) of a DNA locus, fig. 40d) shows the anomalous diffusion behavior, characterized by an exponent *α* < 1, [178, 139, 284].

The anomalous exponent of a locus contains physical information about the underlying polymer model describing the chromatin. However, there is still a debate about the precise meaning of the anomalous exponent computed from empirical data. The anomalous exponent of many loci along chromosome XII show (Fig. 42a) a mean value around 0.5 (Fig. 42b). Consequently, in [105] it was suggested that the Rouse flexible chain model is sufficient to describe chromatin dynamics in yeast. However, a cell-to-cell analysis reveals that there is a large distribution of the anomalous exponent, suggesting that the chromatin structure changes over time and across cell populations [105, 4, 18].

**Fig. 42.**
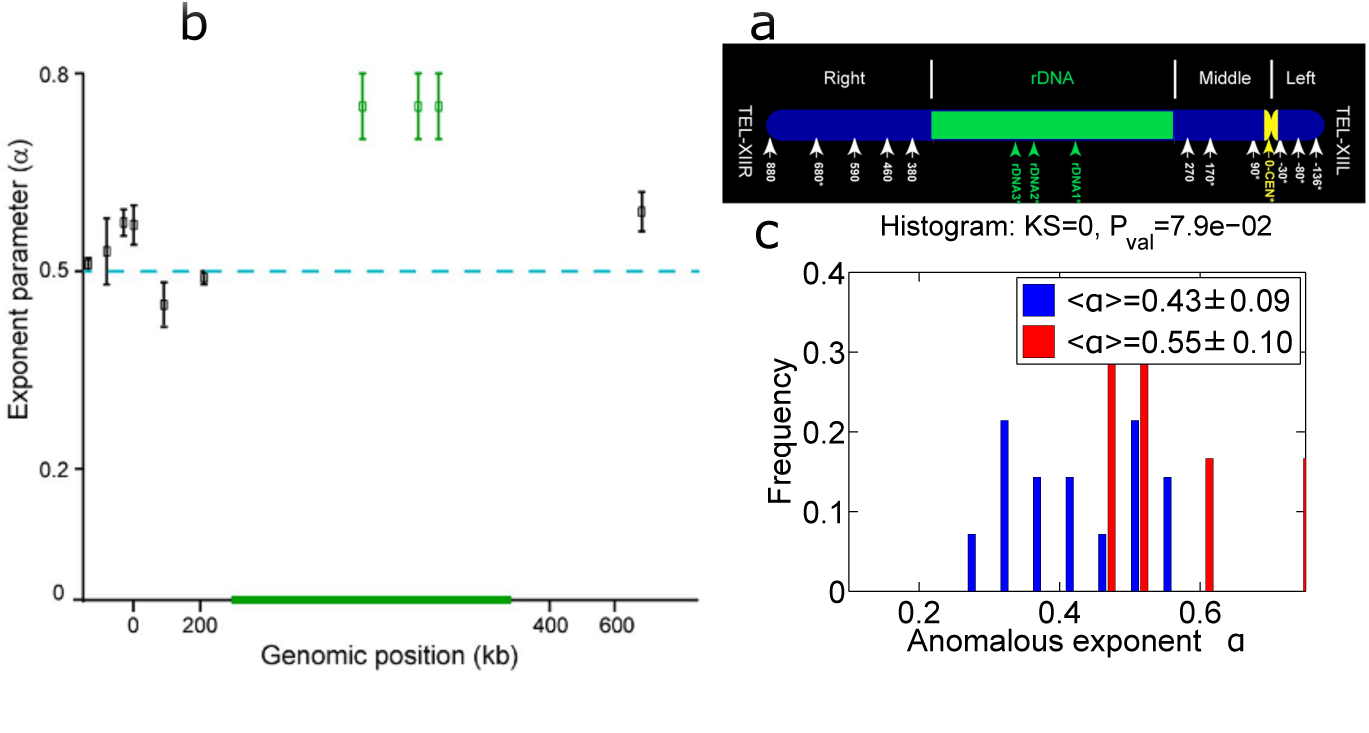
Anomalous exponent of chromatin loci in yeast. (a) Color-coded statistical mapping of positions of 15 loci along chromosome XII. Schematic representation of chromosome XII with the 15 FROS-labeled loci. The rDNA of 1.8 Mb is depicted as a 1-Mb segment. Loci studied by particle tracking are marked with asterisks [4]. (b) Anomalous parameter versus the genomic position for 10 loci [4]. (c) Distribution of the anomalous exponent from the MAT locus (chromosome III) for chosen cells with a xed SPB when the cells grow in glucose (blue) or galactose (red) (reproduced from [18]).

#### 8.4.1 Variability of the anomalous exponent over yeast cell population

The anomalous exponent is estimated from the projection of SPTs on the axis of an orthogonal frame, but which axis should be chosen? Although the anomalous exponent vary significantly across cell population, its value is often reported above 0.5 (Fig. 42b,c for yeast). To compare the DNA locus motion with respect to the nucleus, a reference point should be chosen, such as the SPB. Since the SPB is supposed to be fixed at the periphery, any precession (rotational) motion can be detected (Fig. 43a).

**Fig. 43.**
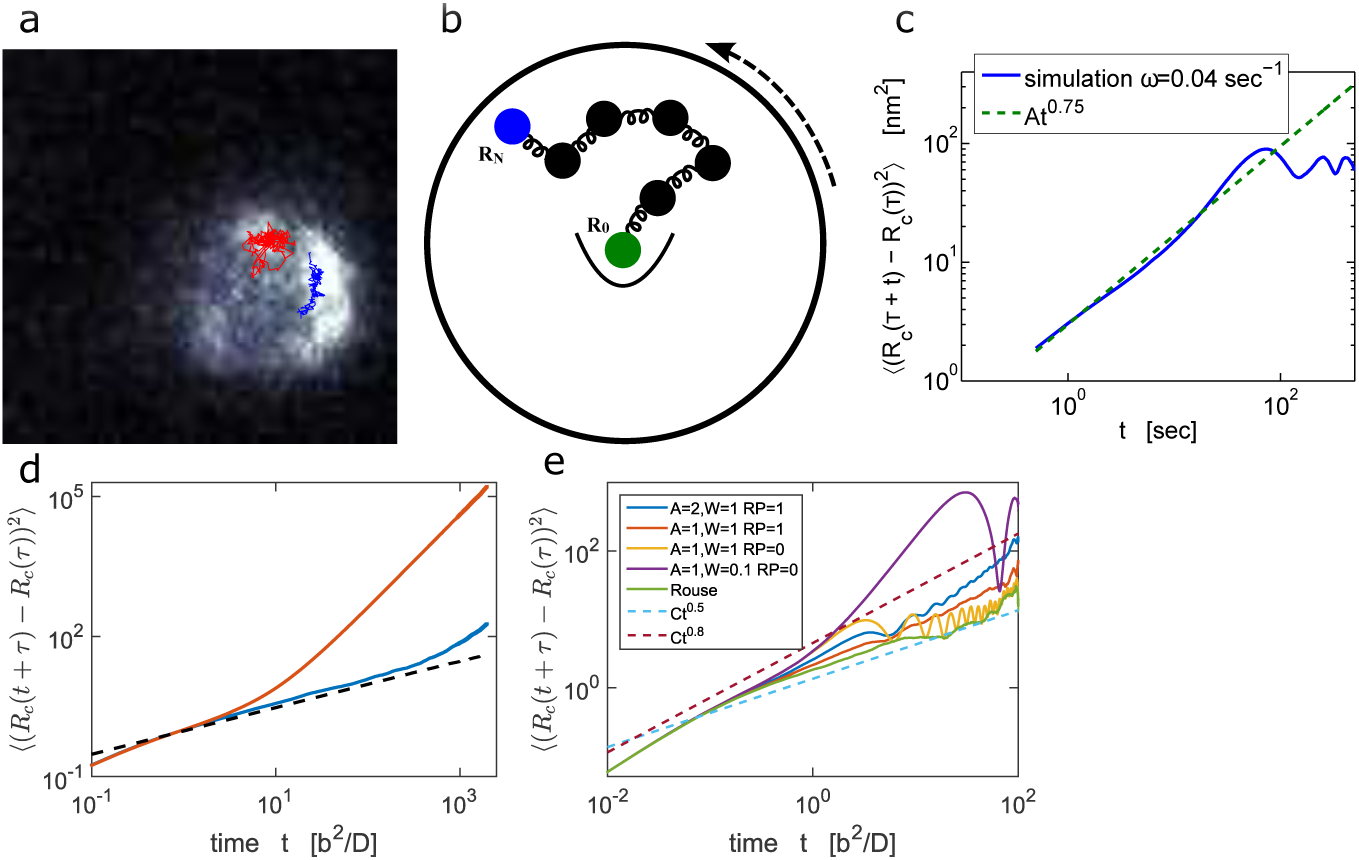
Effect of external drift on the anomalous exponent.α. (a) The nucleus is monitored and the SPB (blue trajectory) is performing a precession motion, compared to a tagged locus trajectory (red). (b) a Rouse polymer model in a spherical domain, with one end anchored at the center, while following the other end. (c) Cross-correlation function and anomalous exponent of the tagged monomer in (b), with *N* = 33 and the domain is rotating in an angular velocity of ω= 0.04sec^1^. The anomalous exponent is approximately equal to 0.75 (and not 0.5 for Rouse). (d) Effect of an external drift on the anomalous exponent α: the cross-correlation function was estimated from Brownian simulations of a Rouse polymer (β = 2 blue line), where the position of the first monomer was followed. At intermediate time, the particle performs anomalous diffusion with α= 0.66, which should be compared to α= 0.5 (black dashed line). (e) The time cross correlation function of the middle monomer position of a Rouse polymer (*N* = 33), where an oscillating force is acting on all monomers (Eq. 207). The phase of forces can be the same for all monomers (RP=0) or random (RP=1). We estimated the cross-correlation function for several values of the amplitude (*A*) and frequencies (*W*). Also shown is the result for the Rouse polymer, with no oscillating force, for which the anomalous exponent is 0.5 (green). Two trend lines (α = 0.5 and α= 0.8) (reproduced from [15]).

The effect of precession on a locus motion is evaluated using Brownian simulations of a polymer confined in a rotating sphere. A polymer end is attached to the center of the sphere. When the angular velocity is *ω* = 0:04sec ^1^ (Fig. 43b), the anomalous exponent increases from 0.5 (Rouse polymer) to 0.75 [18] (Fig. 43c). This result provides a criteria for selecting trajectories originating from different cells: if the SPB is not moving, the anomalous exponent is lower compared to cases where the nucleus is driven by an external motion (Fig. 42c,α = 0.43 ± 0.09). To conclude, the Rouse model characterized by α = 0.5 cannot account for the large distribution of anomalous exponent α observed across cell population, which reveals the heterogeneity of the chromatin organization at a given locus site. For α ≤ 0.5, β-polymer model can be used to interpret SPTs and suggests different chromatin organization for a given locus across cell population (see also [246]). Possibly, nucleosomes local remodeling can change the local short-and long-range forces re ected in the distribution of the anomalous exponents [18].

Directed motion has also been reported to affect the anomalous exponent [50] by following telomere motion in ALT (alternative telomere maintenance) cancer cells. In these cells, telomeres elongate via homologous recombination, where they encounter and use each other as a template for elongation: Telomeres perform long-range movement to actively search for recombination partners. Telomeres perform anomalous diffusion with α = 0.8 [50] on average. Similarly, the motion of the TetO locus of Tsix TAD in the X chromosome perform intrinsic motion characterized by an exponent α = 0.56 [270]. Similar subdiffusive motion are found in pro-B cells [162].

#### 8.4.2 Increasing the anomalous exponent by applying an external drift

Actin network in the cytoplasm can pull the nucleus membrane leading to random oscillations that could potentially affect the internal motion and organization of the chromatin. To explore how a deterministic force acting on a polymer in confined domain affects a monomer motion, numerical simulations have been used where a deterministic force is added to the polymer model. Starting with a Rouse polymer and adding a constant drift in a given direction of amplitude ∣ *v*∣ = 0.2 *D/b*, the monomer motions are now described by

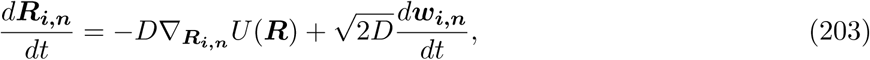

where

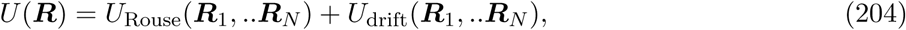

Where *U*_Rouse_ is de ned by eq. 5 and

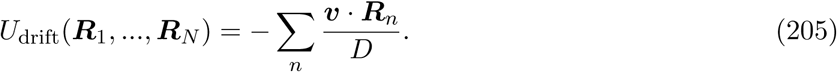

There are no boundaries in the simulations. How this additional motion affects the statistics of a single locus? this question is addressed by computing the anomalous exponent α in the intermediate time regime (Fig. 43d). Contrary to the classical Rouse model or β-polymer models, under a constant pulling force, the anomalous exponent increases to α = 0.66 (Fig. 43d). In the long-time regime, the motion of the center of mass becomes ballistic. Thus, adding a deterministic drift on the motion of a polymer results in an increase of the anomalous exponent [18, 15]. Deriving an analytical formula would certainly help to clarify how the anomalous exponent α depends on an external force. There are almost no analytical results in that direction.

#### 8.4.3 Oscillating forces acting on a monomer modify the anomalous exponent

Adding an oscillating force, that can represent oscillation of the nuclear membrane, affects monomer motion and eventually the anomalous exponent. This effect can be studied by adding an oscillatory potential to a Rouse polymer so that the total energy in eq. 203 becomes

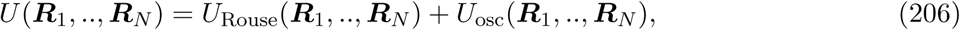

and

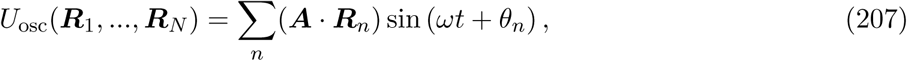

where *A,* ω are constants and compared to eq. 205, the oscillating velocity is written as *v* = *A* sin (ω *t* + _*θ_n_*_), with phases _*θ_n_*_. There are two extreme cases rst when all _*θ_n_*_ = 0 for *n* = 1 *… N* or second _*θ_n_*_∼ *U*(0, 2π) are random variables chosen uniformly distributed. By increasing the amplitude *A* or the frequency ω the anomalous exponent increases (Fig. 43e). Numerical simulations shows that for *A* = *kT* / *b* = 1 and *W* = *b*^2^/ *D*, where θ_*n*_ are randomly chosen, the anomalous exponent increases from α = 0.5 to 0.8. [18, 15]. In summary, adding a deterministic motion, which may be an artifact of the measuring system, increases the anomalous exponent α This reason may explain the very high values of α estimated for loci within of chromosome XII [4] of yeast.

Similarly, the rDNA locus is an active chromatin area and is transcribed by RNA Polymerase (Pol I) and Pol III [204], where about 600 protein coding genes are transcribed Such a domain is expected to be under physical stress, where chromatin is repeatedly re-opened. This dynamics suggests that some external forces might be applied on the nucleus. In [90], an alternative approach to model the effect of ATP-dependent active fluctuations was taken, by considering exponentially correlated colored noise acting on the monomers of a semiflexible polymer. In this case the colored noise also resulted in anomalous motion with an exponent much higher than the one obtained with a Gaussian noise (α= 3 */*4 for a semiflexible polymer). If this activity is modeled by an oscillatory motion (Eq.207), high anomalous exponent (α~ 0.8) are expected. While values higher than 0.5 can be explained by a deterministic motion and forces acting on the chromatin, values smaller than 0.5 reflect chromatin condensation, modeled by the β-polymer model (section 3.5) or viscoelastic nuclear medium (section 3.10.3).

### 8.5 Probing chromosomal structure and dynamics in bacteria using SPTs

Chromosomal structure and dynamics in bacteria can be explored using fluorescently labeled chromosomal loci in E. coli at a timescale of 0.1100 s [129, 128]. A large heterogeneity in trajectories has been reported containing diffusing and ballistic epochs, with abnormally and elongated regions characterized by superdiffusive motion (Fig. 44). These directed movements were analyzed by computing the anomalous exponent α and the diffusion coefficient along single trajectories. The statistics reveals a large spectrum of anomalous exponent, centered around α= 0.4, indicating a higher order connectivity compared to Rouse (see subsection 3.10.2). In contrast with sub-diffusion, the ballistic motions reflect active mechanisms or directed forces, and support the heterogeneous chromosome motion on a viscoelastic background for the bacterial nucleoid or local forces. To conclude, the variability in the anomalous exponent α reveals the local effect of nuclear proteins on the chromosome motion.

**Fig. 44.**
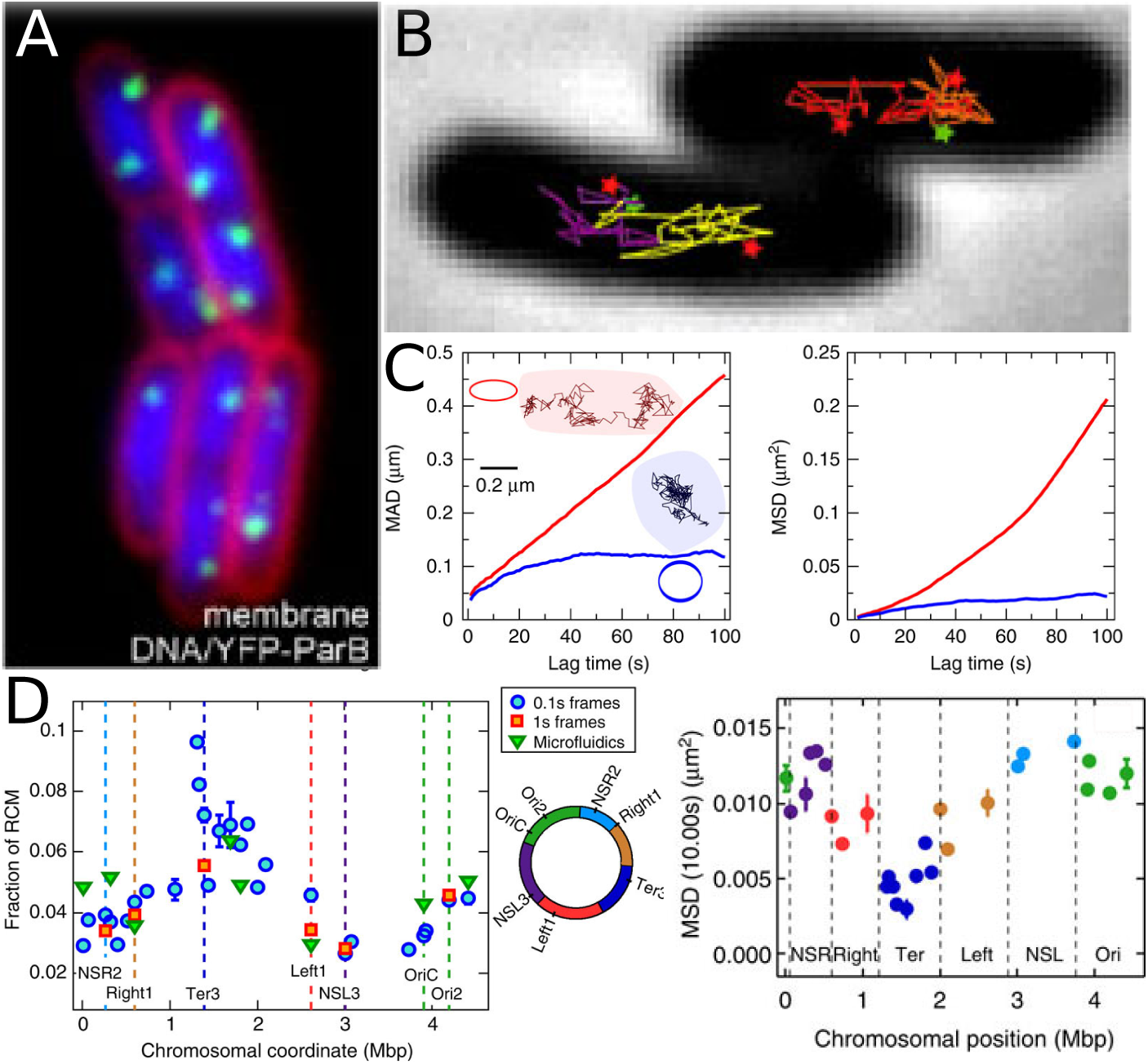
SPTs to probe chromosomal structure and dynamics in bacteria. (A) Cell sub-compartment visualization, membrane (FM 4-64), DNA (DAPI)and foci dynamic measurement (YFP-ParB) stainings. (B) Trajectories of fluorescent dots of foci dynamics compared to territories (stars) in which genetic loci are labeled [175]. (C) Analysis of chromosomal loci trajectories: a ballistic (red) and a typical (blue) track (Left) are represented with their elliptic convex-hulls. The time-averaged MAD (left) and mean square displacement (right) of the two tracks reveals heterogeneous motions. (D) Fraction of rapid movements (RCM) as a function of chromosomal coordinate: Squares correspond to the loci belonging to different macrodomains. Experiments on 27 loci with frame separation of 0.1s, 1s and in a micro uidics device (triangles). Right: Means square Displacement (MSD) showing the intrinsic differences with the chromosomal location (reproduced from [129] and [128]).

Interestingly, there is also a relation between dynamics and geometrical position: Indeed the loci with the highest variability belong to a macrodomain found near the cell periphery [129], possibly subject to tethering interactions at least during certain parts of the cell cycle. Direct computation using formula 226 should clarify the nature of the ballistic motion.

### 8.6 Local interactions acting on chromatin

External and internal forces are constantly remodeling the chromatin driven by protein complexes that can compactify and reorganize the DNA. Chromatin fragments interact with many partners such as the nuclear membrane, other chromosomes or nuclear bodies, but the resulting forces cannot be directly measured *in vivo*. However, these forces impact chromatin dynamics and should be reflected in particular in the motion of a single locus (Fig. 45a). It is now possible to extract tethering forces applied on chromatin from the statistics of a single locus trajectories imaged. Indeed, DNA locus plays the role of a probe to reveal the local forces (Fig. 39c). The motion of a chromatin locus can be driven by local diffusion and/or forces between monomers of the model polymer [138, 291, 274]. Monomers motion is highly correlated due the polymer hierarchy of relaxation times [72, 254], leading in particular to anomalous diffusion [137]. Much of the chromatin dynamics is reflected in the motion of a single chromosomal locus and conversely, a locus motion allows probing the chromatin dynamics [154, 177] at tens of nanometers and millisecond scales resolution [297]. Measuring these forces from a single locus trajectories can reveal genomic organization, information about chromosome arm interaction and obstacle organization [279, 15].

When a local interaction is applied on a given locus, it affects the polymer chain, and conversely any monomer motion will be affected by possible interactions exerted along the chain (Fig. 45a).

**Fig. 45.**
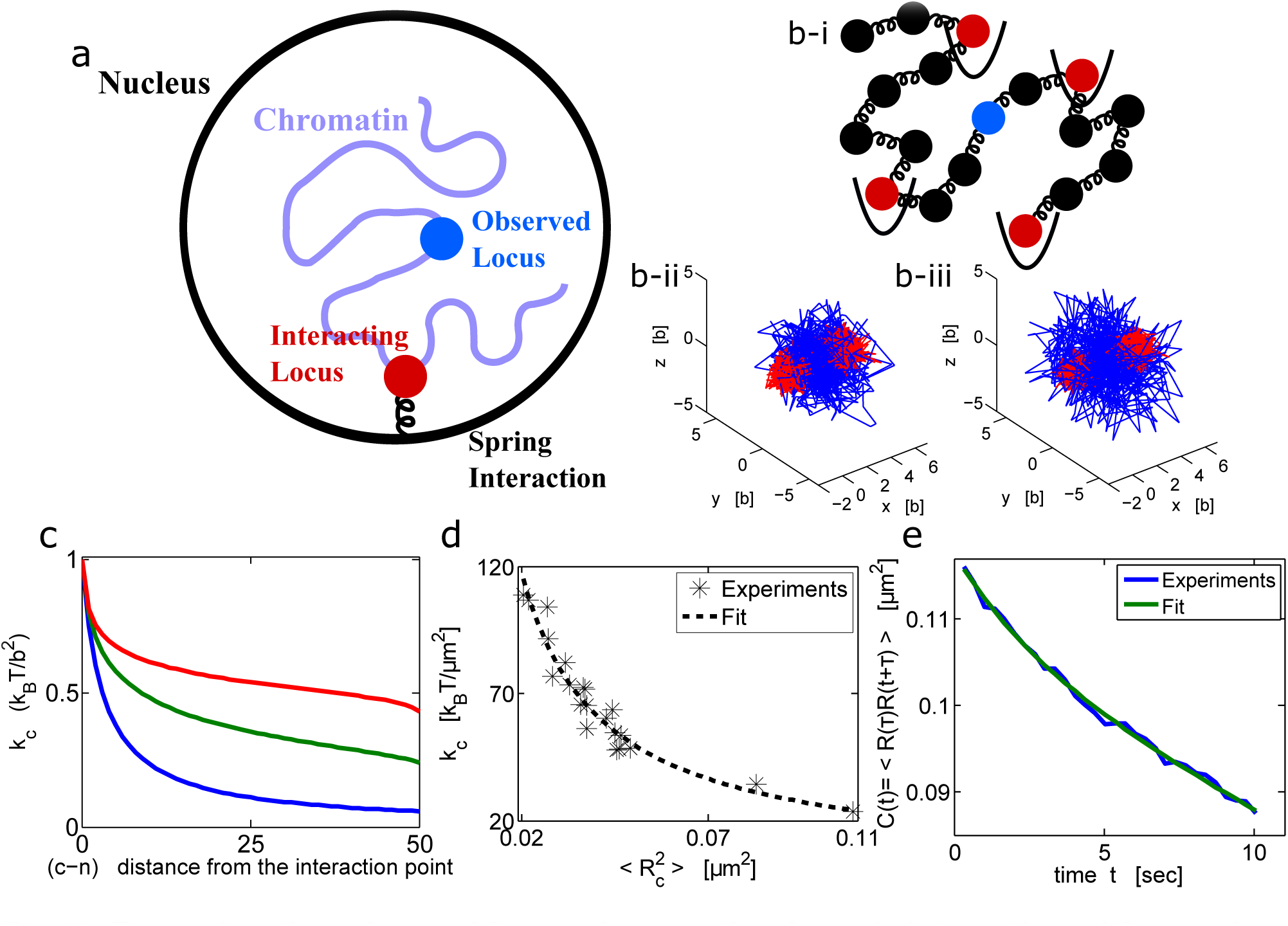
Dynamics of an observed locus when another locus is interacting with a nuclear element. (a) Schematic representation of the nucleus, where one locus is observed and followed with orescent label while another (non-visible) chromatin locus is interacting with another nuclear element. (b-i) Schematic representation of a polymer, where some monomers (red) interact with xed harmonic potential wells. Monomer *c* (blue) is observed. (b: ii, iii) Stochastic trajectories of three monomers, part of a polymer where the two extremities interact with potential wells xed at the origin and at position *μ* = (5 *b;* 0; 0) respectively. The middle monomer trajectory (blue) is more extended than the two others, shown for a polymer of length *N* = 21 (ii) and *N* = 41 (iii). (c) The apparent spring constant *k*_*c*_ is computed from formulas 218 and 222, for a polymer of length *N* = 100, where monomer *n* = 50 interacts with an harmonic potential (eq.214) with *k* = 2 *k*_*B*_ *T/b*^2^, while *k* = 3 *k*_*B*_ *T*/b^2^. *k*_*c*_ is computed for increasing distances | *c* - *n*| between the observed and the interacting monomers for β= 2 (Rouse polymer) (blue), β= 1.5 (green) and β= 1.2 (red). (d-e)Analysis of trajectories of the chromatin MAT-locus (located on chromosome III) in the yeast Sc. (c) Scatter plot of the effective spring coefficient *k*_*c*_ and the variance (*R*_*c*_^2^) of the locus trajectory estimated in two-dimensions, extracted for 21 cells. The constant *k*_*c*_ is estimated using formula 226, fitted to a power law, 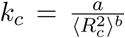 with *a* = 3.03 ± 1.05 *k*_*B*_ *T* and *b* = 0.94 ± 0.1. (d) Autocorrelation function computed using formula 191 for the trajectory shown in a. The t uses the sum of two exponentials: C(t) = a_1_e −t/T_1_ + a_2_e −t/T_2_ with τ_1_ = 45.7 —± 0.005 *s* and τ_2_ = 2.4 ± 0.35 s, *a*_1_ = 109 ± 5×10^-3^ *μm*^2^, *a*_2_ = 8.38 ± 4.94×10 ^-3^ *μm*^2^ (reproduced from [16]).

When a far away monomer is anchored, the motion of the observed monomer is certainly restricted. Indeed, when applied forces are stationary over the time course of a trajectory recording, the empirical velocity distribution of a locus is related to forces applied to a distant single monomer. It is possible to distinguish external forces applied on a single monomer from intrinsic forces acting on monomers. The principle and the difficulty of the method can be understood as follows: for a single stochastic particle modeled by the Smoluchowski's limit of the Langevin equation, the velocity of the particle v is proportional to a force f applied on the particle plus an additional white noise, summarized as

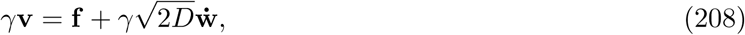

Where γ is the friction coefficient, *D* the diffusion coefficient and *w* is the normalized wiener process. Thus by averaging over the ensemble of velocity realizations, it is possible to recover the first moment, which is the force field. However, for a polymer chain, there are internal forces between monomers and thus, as the data are measured at a single monomer, the internal forces acting on the measured monomer need to be separate from the external ones acting on a monomer further away. Analytical formula are constructed to recover the force acting on the polymer model. When the external applied force is the gradient of a quadratic potential. The method is presented in in the next sections, we review recent effort to recover far away interactions from the analysis of SPTs at a given locus.

#### 8.6.1 Recovering a tethering force (external potential) for a Rouse polymer

To extract a tethering force acting on a Rouse polymer from single locus trajectories, we recall first how a force *F* (*X*) can be recovered from the trajectories of a single particle, modeled by the overdamped Langevin equation

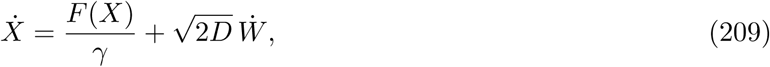

Where *W* is a Gaussian white noise and *γ* is the viscosity [238]. The source of the driving noise is the thermal agitation [238, 236]. The present model and analysis assumes that the local environment where the particle evolves is stationary. Moreover, the statistical properties of the trajectories do not change over time. In practice, the assumption of stationary is usually satisfied as trajectories are acquired for short time, where few statistical changes are expected to occur. The parameters in the coarse-grained model 209 are recovered from computing the conditional moments of the trajectory increments Δ *X* = *X*(*t* + Δt) − *X*(*t*) (sec[236]),

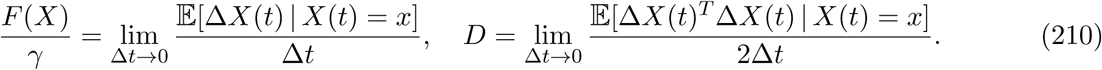

Here the notation 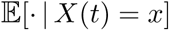means averaging over all trajectories that are at point *x* at time *t*.

To generalize relation 210 to a Rouse polymer embedded in an external force, we define the potential *U*_ext_(*R*) and described the total energy of the interacting polymer by

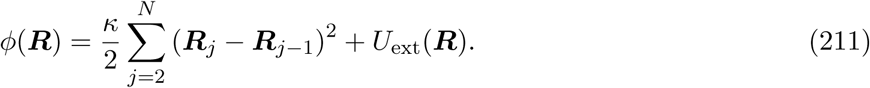

The monomer dynamics follows stochastic eq.14. When monomer *R*_*c*_ is monitored and trajectories are collected, the first moment of the monomer increment position R_c_(*t* + τ *t*) − R_c_(*t*) is proportional to the velocity. Computing the increment involves averaging over all polymer realizations and accounts for the entire polymer configuration [17]. The following relation was derived in [17]

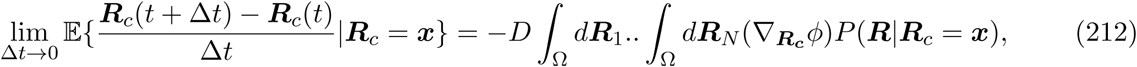

where 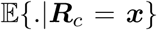 denotes averaging over all polymer configuration under the condition that the tagged monomer is at position *R*_*c*_ = *x*. Formula 212 is generic and generalizes relation 209. Furthermore, it does not depend on the particular expression of the external forces acting on the polymer. No restrictions are imposed on the domain Ω where the polymer evolves, but the polymer is re ected on the boundary *@*Ω. The conditional probability *P*(*R*| *R_c_* = *x*) is computed from the equilibrium probability distribution function (pdf) *P*(*R*_1_, *R*_2_,…, *R*_N_) which satisfies the Fokker-Planck equation (FPE) in the phase space 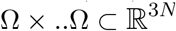

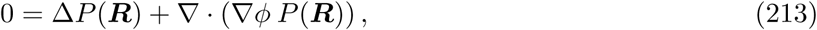

with boundary condition

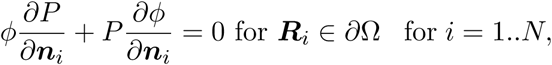

Where *n*_*i*_ is the normal vector to the boundary *∂*Ω at position *R*_*i*_. The model for the external force acting on monomer *n* and located at position *R*_*n*_ is the gradient of a harmonic potential

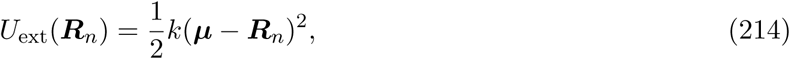

Where *k* is the constant force and we choose *n < c*. This external potential well affects the dynamics of the entire polymer and specifically the observed locus *c*.

#### 8.6.2 Extracting a deterministic force from an ensemble observed locus trajectories

To extract the strength of the potential well applied on monomer *n* from the measured velocity of locus *c*, we need an explicit expression for formula 212. First, the force acting on monomer *c*, when its position is *x* is given by

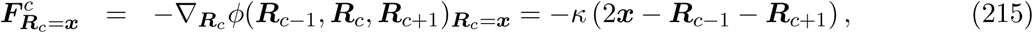

Where *ϕ* is defined in 211. When the potential well is localized at the origin ( μ= 0) in eq.214, the equilibrium pdf is the Boltzmann distribution conditioned on *R*_*c*_ = *x*. It is given by

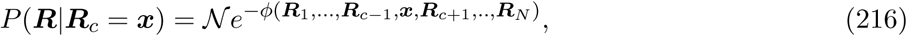

Where *N* is a normalization factor 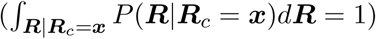, involving Gaussian integrals. The result is

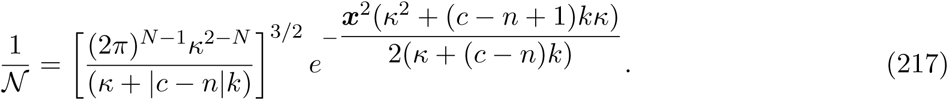

Substituting eqs. 215 and the normalization expression for *N* into eq.212 [17], the mean velocity of monomer *c* is computed and given by

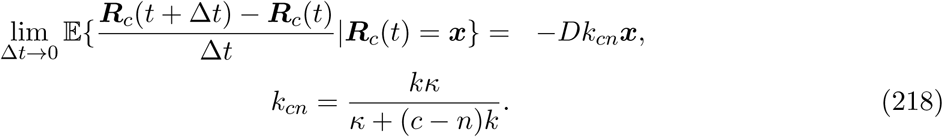

Expression 218 links the average velocity of the observed monomer *c* to the force applied on monomer *n*. The coefficient *k*_*cn*_ depends on the harmonic well strength *k*, the inter-monomer spring constant κ and it is inversely proportional to the distance *∣n-c∣* between monomers *n* and *c* along the chain. Furthermore, the steady state variance 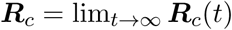of the monomer‘s position is given by

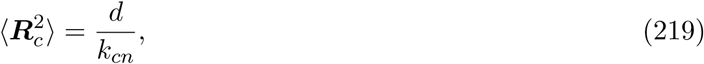

When 〈 *R_c_*〉 = 0 Relation 219 is reminiscent of long time asymptotic of classical Ornstein-Uhlenbeck processes. The dynamics of monomer *R*_*c*_ generated by Brownian simulations is shown in fig.45b-c. In the limit of large *k* (pinned monomer), an analogue of formula 219 was used for analyzing chromatin organization [279] and DNA [158]. Inversion formula 218 assumes the Boltzmann distribution for the single monomer and that the entire polymer has reached equilibrium at the time scale of the simulation or the experiment (from Eq. 216). Finally, formula 218 reveals how internal and external polymer forces are mixing together and are in uencing the monomer velocity. It also shows the explicit decay of the force amplitude with the distance between the observed and forced monomer.

In summary, the coefficient *k*_*cn*_ can be estimated from Brownian simulations of a tethered polymer with self-avoiding interactions. The comparison with the experimental radius of confinement measured for chromatin loci [181, 193, 111, 39], reveals that the distance from the centromere is inversely correlated with the effective spring constant. Similar results are found for a tethered DNA molecule [158].

#### 8.6.3 Polymer dynamics constricted by two tethering forces

The analysis of a single force can be generalized to the case of two monomers *n* and *m* (*n < m*) that are interacting with two distinct potential wells applied at positions _*μ_n_*_ and _*μ_m_*_ (Fig. 45b). The total potential energy of the Rouse polymer is

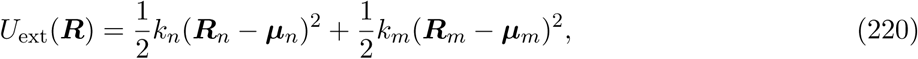

and the average steady state position of the tagged monomer *c* is given by [17] 

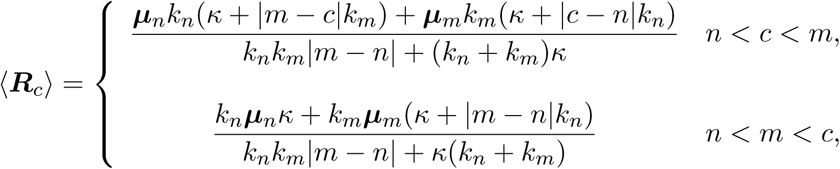

and 

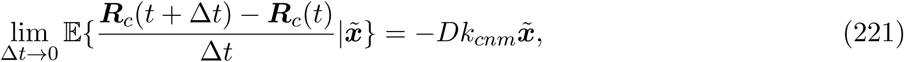

where 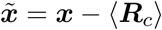 The effective spring coefficient is given by [17]

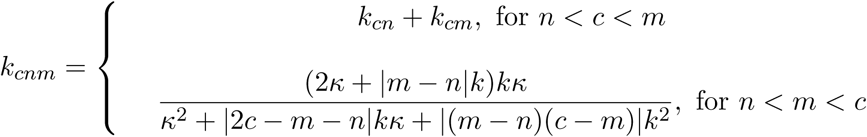

Where *k*_*cn*_ and *k*_*cm*_ are given by eq.218. For *n < m < c*, in the limit *m − n* ≫ 1, then *k*_*cnm*_∼│ *C-M*│^-1^ *k*. The spring coefficient depends on the distance to the anchoring point only. When *n < c < m*, the effective spring coefficient depends on the positions of the two wells [279]. Finally, the variance of the monomer position with respect to its mean position (eq.221) is 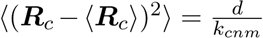 Thus, the distance scanned by the observed monomer is proportional to the distance to the anchoring point. (see fig.45b-ii, iii).

#### 8.6.4 Analyzing tethering forces for a β-polymer

The polymer model [10] introduced in section 3.5, captures long-range interactions between monomers, that decay with the distance along the chain 3.5. Chromatin dynamics with an anomalous exponent less than 1/2 can thus be described by β-polymer models. When a gradient force (see eq.is 214 applied on monomer *R*_*n*_ of a polymer, the expectation of the velocity of monomer *c* (*c > n*) is given by [17]:

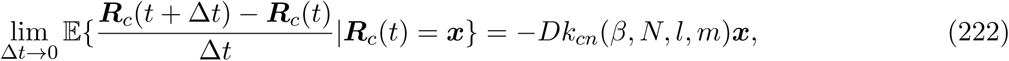

where μ= 0 and 

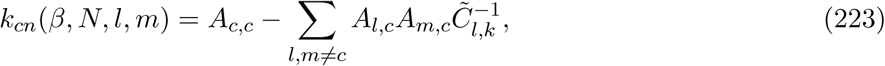

where 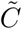 is a block matrix, the *i*-th block of which is

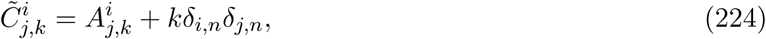

and [10]

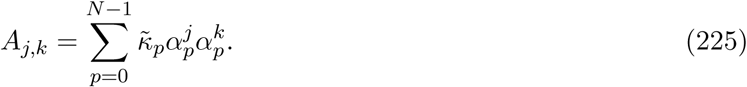

Where the matrix α _*p*_
^*j*^ is de ned in18. The inversion formula 222 for a βpolymer is similar to the one derived for a Rouse polymer (eq. 218), but the dependency with the parameters is now implicit [17]. Computing numerically the parameters in equations 223-225 reveals that the apparent spring constant *k*_*cn*_(β *β, N, l, m*) decays slower with the distance │ *c-n*│ (between the interacting and the observed monomer) and for smaller β (Fig. 45c). In summary, when the anomalous exponent is α<0.5, monomer trajectories are influenced by both the interacting monomers and the anomalous dynamics, characterized by the exponent α, that reflects the intrinsic chromatin organization.

#### 8.6.5 Extracting forces acting on a locus from SPTs

We now present the empirical estimator for computing the effective spring coefficient *k*_*c*_ (eq. 218) from locus trajectories R_c_(hτ *t*) (*h* = 1..N_p_). We define the following empirical estimator

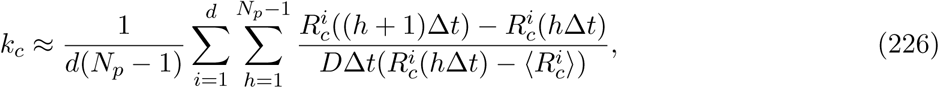

where *N*_*p*_ is the number of points. The first step is to extract the mean position of the locus. Once the steady state is reached, the time average of the locus position is 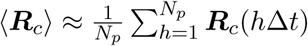

This estimator can also be used when the polymer is not in equilibrium, but in this case 〈Ri_c_〉 should be estimated separately. The diffusion coefficient can also be estimated using eq.202.

While *k*_*c*_ is estimated using eq.226, there are two unknown parameters: the spring force *k* and the distance | *c*– *n*|. For a strong anchoring (*K ⪢ k*) and the approximation k_c_ = k|c − n|^−1^ is valid. The empirical effective spring constant can be used to estimate the distance to the interaction monomer.

For a long enough sampled trajectory and a force derived from a stationary potential well, the effective spring coefficient can be recovered directly either from the empirical estimator 226 or by using the reciprocal of the variance (eq.219). However, trajectories are often measured with a small sampling time Δ *t* allowing probing the fine behavior of the chromatin and recovering accurately the diffusion coefficient. The total length of a trajectory is however limited by photobleaching effects [217]. Thus, the length of a trajectory may be shorter than the equilibration time scale, and thus acquired before equilibrium is reached. In that case, formula 226 can still be applied to recover the parameter *k*_*c*_, while formula eq.219, which implies equilibrium, cannot be used.

#### 8.6.6 Recovering applied forces from the auto-correlation function

The auto-correlation function C(c,t1,t2) = 〈[R_c_(t_1_) − R_c_(t_1_)〉][R_c_(t_2_) − R_c_(t_2_)〉] of a tagged monomer *c* is changed when a force is applied. Indeed, we decompose the energy potential of a polymer into the sum of internal Rouse plus an external component (eq.214) in the eigenvalue basis *u*_*p*_ that diagonalizes the Rouse potential 18. The external energy is

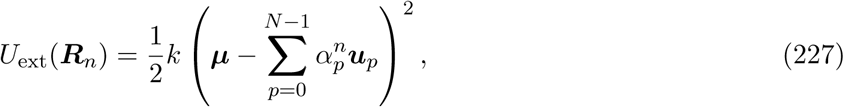

and the Langevin equations for the polymer are given by

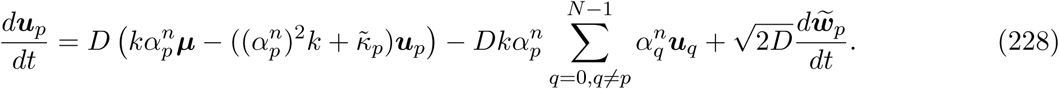

For *p* = 0… *N* – 1. When the strength of the coupling term is relatively weak 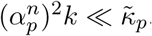 the coupling term can be neglected as well as higher modes for *K < k* and *N* large. Thus the auto-correlation function is

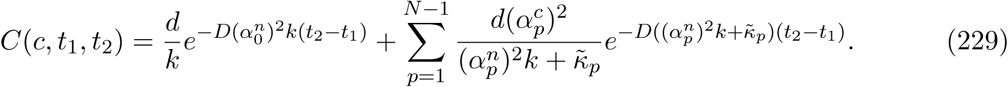

The long-time asymptotic exponential decay can be used to extract the spring constant *k* of the applied force, when the diffusion coefficient D is known.

#### 8.6.7 Distribution of forces of MAT-locus from live cell imaging SPT in yeast

We now discuss the implication of formula 226 to study the dynamics of a moving chromatin locus. We will present data from the MAT-locus acquired during 100sec with a time resolution of Δ *t* = 0.33sec. Trajectories are covering a certain portion of the nucleus. Several predictions have been made based on formula 226: the computations on 20 cells revealed a large heterogeneity of *k*_*c*_ (Fig. 45d), where. This heterogeneity indicates that in different cells the locus and the local chromatin interact differently with other nuclear elements. Plotting the force constant *k*_*c*_ for each cell with respect to the locus position averaged over the trajectory shows that (〈R^2^_c_〉) satisfies relation 218 (Fig. 45d), confirming that chromatin locus localization is governed by local interactions. The relation between *k*_*c*_ and is found empirically to be

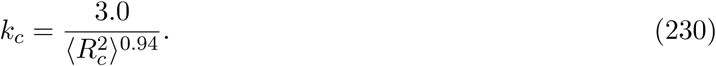

In addition, the auto-correlation function [237] for the MAT-locus position is computed from the estimator

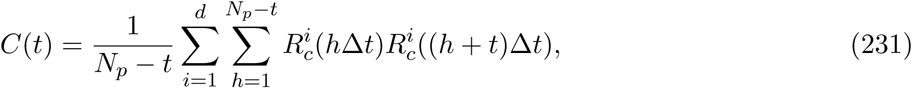

And it is represented in fig.45e. Fractional Brownian motion has been previously used to model the dynamics of chromatin loci [138, 285, 127]. For fractional Brownian motion, the auto-correlation function *C*(*t*) decays as a sum of power laws [130]. Thus the empirical correlation function *C*(*t*) cannot be fitted by a power law, suggesting that the description of the locus motion as a fractional Brownian motion alone is not sufficient. However, a good fitting of the function *C*(*t*) is obtained using a sum of two exponentials

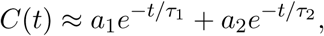

with τ_1_ = 45.7 ± 0.005sec, τ_2_ = 2.4 ± 0.35sec and *a*_1_ = 109± 5 × 10 ^-3^ μm^2^, *a*_2_ = 8.38± 4.94 × 10 ^-3^ μm^2^. This fit suggests that the auto-correlation function for the locus position is well described by a sum of exponentials, as predicted by formula 229 derived for general polymer model.

In summary, polymer models such as classical Rouse or polymer accounts for the dynamics of a chromatin locus. In that context, it is possible to extract characteristics of the DNA locus, its dynamics, external forces and some properties of the polymer model from SPTs. At this stage, the origin of the anchoring force is not clear and may occur at the centromere. However, the large heterogeneity of the spring constant *k*_*c*_ characterizing an external applied force suggests (Fig. 45d) that the nearest anchoring points of that force is different for the same locus in different cells. Finally, the motion of the chromatin is driven by both thermal fluctuations and by active ATP-dependent forces [295]. While the above analysis is relevant to extract an interaction that does not change during the time acquisition of the trajectory, the spring constant *k*_*c*_ that would be extracted during an active chromatin motion could be differentiated from the thermal one by projecting the dynamics perpendicular to the direction of motion. Extracting active force elements may perhaps be done by considering the random force spectrum [90] or through the analysis of the correlation function, which are in uenced by external forces.

## 9 Physical principles underlying DNA Repair

The nucleus organization can be revealed by the dynamics of loci at speci c locations, such as nuclear pores at the membrane periphery, transcription regulation hot spots [184, 75, 32, 267], DNA repair foci [70], transient foci of proteins [51] or chromatin loci [244]. The life-time of these transient structures certainly determines their efficiency. Although the formation of these structures is still unknown, the search of homology during DNA repair is of particular interest and it consists of a random search by the broken strand for a homologous DNA template to repair the break. This section is dedicated to the polymer modelings describing the successive steps involved in the search of small targets and in particular during homologous recombination (HR).

### 9.1 Encounter loci at the nuclear envelope

Although the role of the nuclear envelope is to separate chromosome from the rest of the cytoplasm, in differentiated nuclei, part of the chromatin (heterochromatin) is relocated along the inner face of the nuclear membrane, while open chromatin is found near nuclear pores. Evidence from yeast suggests that the nuclear envelope controls also gene transcription, DNA break repair and differentiation [3].

Yeast telomeres interact with the nuclear membrane through two redundant and cell cycle regulated pathways, involving telomere bound proteins Sir4 and Ku [265]. The perinuclear localization of this telomere can be increased upon activation of the HXK1 gene located 15kb upstream of the telomeric repeats. Transcriptional activation of the HXK1 in the presence of galactose increases its interaction with the nuclear pore complex and hence its perinuclear localization [266, 263] (Fig. 46a). In the presence of glucose, this gene becomes silent and on average it is relocated further away from the periphery. The localization of this particular telomere is mediated by the yKu protein in the G1 phase of the cell cycle. Interestingly, tagging a locus located near the subtelomeric region is used to visualize its reduced mobility (Fig. 46b), shown by the decrease of the length *L*_*C*_ (Fig. 46c). This reduction results from the interaction between the telomere and the nuclear periphery, con rmed by the increase of the coefficient *k*_*c*_ (Fig.46d). The diffusion coefficient is also reduced upon activation [266, 18].

**Fig. 46.**
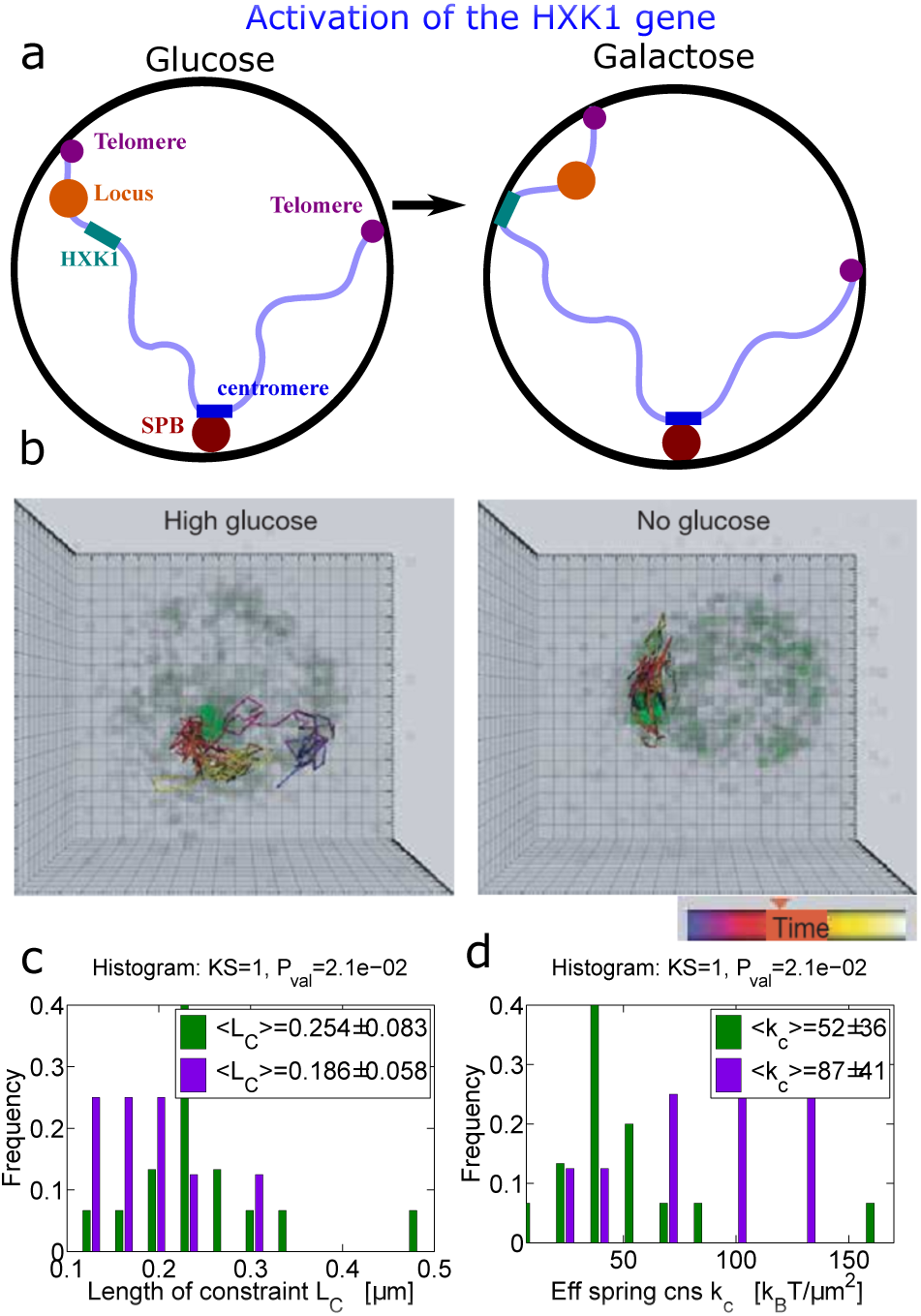
Transcription is regulated by gene re-positioning. (a) The sub-telomeric locus ARS609 next to the right hand of telomere VI (TelVIR) in different growth conditions is compared: glucose (green) and galactose (purple). In both conditions the cells are in G1 [18]. (b) Four-dimensional time-lapse imaging of wild-type cells bearing lacop-tagged HXK1 as described [86] on glucose or galactose-containing media. Image stacks were collected at 1.5-s intervals for 7.5 min and tracking was performed with the Imaris Time program. The darker grey background shows the peripheral pore signal. The path of the locus is shown with a temporal color code (blue represents early and white represents late time points) [266]. (c) Distribution of the constrain length (*L*_*C*_) found using eq.201. (d) Distribution of the apparent force strength (*k*_*c*_) histogram for all cells as measured on the visible monomer *c* using eq.226.

To conclude, gene regulation can involve nuclear positioning and the spatial position of some genes is correlated with the expression level. In addition, some actively transcribed genes (Gal1-10, Ino1) contact the nuclear envelope through interactions with the nuclear pore complexes [263].

### 9.2 Physical constraints underlying double-stranded DNA break repair

Double-stranded DNA breaks (DSBs) can be caused by internal and external agents. Ionizing radiations lead to DSBs, while within the nucleus or promiscuous enzymes can cut both DNA strands. Cleavage of the DNA does not necessary implies that the two strands will separate [112]. In most cases, the two ends remain in close proximity with the assistance of repair proteins such as the MRN complex [93]. DSBs are the most serious and dangerous genetic lesions and if left un-repaired or repaired inaccurately, they can cause chromosomal translocations, cell death, loss of heterozygosity and can lead to the development of a cancerous cell.

DNA double-strand breaks can be repaired by one of two major classes of mechanisms: either by Non-Homologous End Joining (NHEJ), whereby the broken ends of the DNA are simply religated [93], or by homologous recombination (HR), which entails a physical search for a homologous DNA template by the broken strands, which is then used to repair the break [21, 112]. The template for repair may be the chromosomal homologue if the cell is diploid, or the sister chromatid if the break occurs after DNA replication. When a broken and resected DNA end comes close to the appropriate double-strand sequence, it invades the undamaged template and primes DNA synthesis. A subsequent mechanism called Holliday junction resolution leads to an accurate repair of the broken strand [102, 262, 2].

The search for the template is the rate-limiting for HR [182, 2, 289], but the exact mechanism is still unclear. It involves several spatial-temporal scales, from the molecular to the nuclear level, which are starting to be resolved experimentally and theoretically [88, 181, 69]. Several physical scenarios are possible to estimate the search time of the HR. The encounter time depends on the chromosome length, because the probability distribution of loci positions are determined by their length and distance from the centromere. The chromosome length can thus affect the probability of encounter: shorter chromosomes would tend to repair more amongst themselves, as would long chromosomes [2]. How can a single chromosomal locus find a copy of itself? How does it explore the territory of another chromosome in order to nd a small sequence target [54]? The search for homology requires scanning of millions of base pairs for the correct sequence, but this process takes only between tens of minutes to a few hours [2, 49]. How is this possible? there is no final answer to that question. The surprising fast rate at which HR occurs is even more remarkable when considering that it is driven by diffusion and must overcome physical barriers and forces associated with the invasion of a chromatin [15].

The physical paradigm of HR is based on an experimental model for this search process: in these experiments, a well-characterized endonuclease mediated DSB at the MAT locus in haploid budding yeast cells can be induced by adding galactose (Fig. 47a). Following the expression of the endonuclease, the locus is cut with high efficiency. The locus is visualized by the binding of a fluorescent (LacI-GFP) protein at a Lac operator array (LacO), which is integrated in the chromosome at a distance 4.4kb from the cut site (Fig. 47a). Live-cell imaging studies in yeast have shown that a DSB scans a larger area of the nucleus than the unbroken locus, measured by the radius of con nement [181, 69, 15] (Fig. 47b). Equivalently, the value of the length of constraint (de ned by eq.201), increases signi cantly after a break (Fig. 47c).

**Fig. 47.**
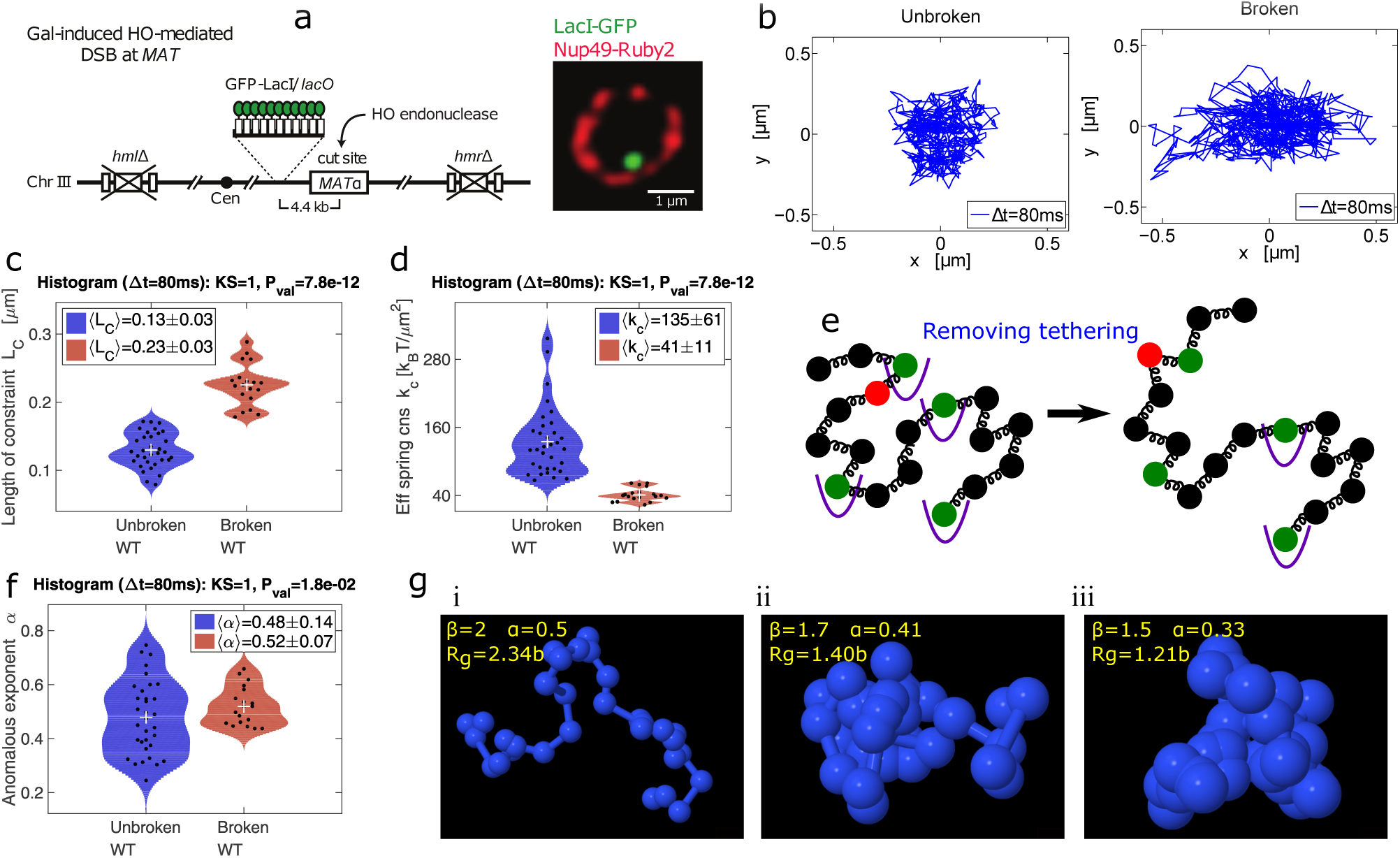
The dynamics of a double stranded DNA break in the MAT locus. (a) Schematics of the experimental system. (b) (left) The MAT locus trajectory during 60 seconds (Δ *t* = 80ms). (right) The MAT Locus trajectories 2 hours after the induction of a break, during repair with the homologous recombination pathway. (c) The length of constraint calculated from the trajectories using eq.201 of a broken locus (red) vs. the unbroken locus (blue). In the title we present the result of a Kolmogorov-Smirnov and the *P* value. (d) Quanti cation of the effective spring coefficient *k*_*c*_(eq.226) before and after break induction.(e)Loss of chromatin sites interactions: Schematicrepresentation of two homologous polymers, where a fraction of monomers (green) participate in a steady state external interaction, modeled by a xed harmonic potential wells (purple), while single trajectories are recorded from monomer *c* (red). (f) The anomalous exponent is extracted from the cross-correlation function of trajectories for a broken (red) vs. unbroken (blue) locus, with time steps of 80msec. (g) Three examples of polymers with (a) β = 2, (b) β= 1.7 and (c) β= 1.5, where the inter-monomer interactions are modi ed, resulting in different condensation state. The radius of gyration *R*_*g*_ (red) measures the degree of compaction associated with, which is related the to anomalous exponent of a monomer by the relation α = 1 - 1/β. Balls (blue) represent the monomers of radius 0.3 *b*. After break induction increase, which indicates the chromatin expansion (reproduced from [15]).

The DSB and the immediate surrounding chromatin can interact with the nuclear environment, the chromosomes, the nucleolus and known DSB anchored sites located at the nuclear envelope (nuclear pores and the SUN-domain protein Mps3 [265, 120]). The accumulated effect of these interactions impact the motion of a tagged locus. These interactions can be modeled as external forces acting on the monomer (discussed in section 8.6) and the strength of the force can be extracted from SPTs. From MAT locus trajectories, the effective spring coefficient *k*_*c*_ (formula 226) was shown to decrease signi cantly after break induction (Fig. 47d). This result shows that the chromatin structure around a DSB undergoes physical modi cations, re ected by a change in the forces applied on the chromatin. This effect suggests that a tagged locus scans a larger nuclear area, emphasized by an increase in the length of constraint *L*_*C*_ (Fig. 47c). Following a DSB [258], the attachment of the centromere to the kinetochore is relieved, resulting in a reduction in *k*_*c*_.

To conclude, the physical changes after break induction in yeast and engagement into the HR repair pathway, support the theory that the chromatin is locally reorganized, nucleosomes are first rapidly acetylated, H2A is phosphorylated, and then evicted around the break, coincident with end-resection [276].

### 9.3 Increasing DSB dynamics reduces the MFET

What is the consequence of reducing local interactions following a DSB motion? it might change the HR search time. Indeed, during HR repair of a DSB, chromatin structure may expand [252], facilitating both the search process [208], and the binding of the repair machinery [298]. Yet, how the chromatin structure is modified to favor this search process is unclear [203, 98]. One possible scenario is the release of forces associated with the relaxation/decondensation of chromatin. This expansion of the chromatin was characterized using the β–polymer model. The anomalous exponent increases following break induction (Fig. 47f). This increase re ects a change in the local property of the polymer model (the chromatin): when it is small (β → 1), all monomers are highly interacting and the polymer has a compacted shape, while for large values of β (β → 2) the polymer is more relaxed (Fig. 47g). After DSB induction β increases, leading to a wider distribution of locus (monomer) position. The increase of the anomalous exponent could also be due to active mechanisms as described in section 8.4.3.

Brownian simulations of polymer models suggest [15] that chromatin relaxation accelerates the search for homology by factor 4 to 8, by increasing the locus mobility and reducing the screening barrier, imposed when the broken site remains hidden in the chromatin structure. When a monomer stays inside the chromatin domain, the surrounding monomers generated expulsion forces between themselves and any other monomers belonging to another polymer (see also subsection 4.12). When a monomer located inside a chromatin domain needs to get in close proximity to a template site, modeled as a monomer inside another polymer, it would have to penetrate into that domain. The interactions between the search monomer and monomers surrounding the target generate an effective potential barrier, that the searcher needs to overcome in order to encounter its target. This potential is due to the local effect of all interacting monomers (see Fig. 4 in [8]). However, when a monomer is robustly positioned at the periphery of a polymer globule, when the chromatin is open, the effective potential is reduced, accelerating the search time [15]. This situation is similar to the scenario of classical chemical reaction where a catalyst reduces the activation potential barrier, accelerating the reaction. It would be possible to use polymer models to estimate the timescales of relocation a DSB toward the nuclear envelope, although precise information would be needed about the chromatin condensation for various distances to the boundary.

In summary, releasing tethering forces around a DSB enhances its motion and signi cantly reduces the homology search time. Thus, the DSB interaction capacity (the external forces applied to the DSB region) with other chromosomes is reduced, allowing an efficient search for a template.

### 9.4 DSB re-localization as a protection mechanism

In budding yeast persistent DBSs that do not have a homologous donor migrate to the nuclear envelope [188, 120] (Fig. 48a), where they are anchored either at nuclear pores or at the SUN domain protein, Mps3. The relocalization of the HO-induced DSB to the nuclear envelope is not immediate, occurring between 30 minutes and two hours after cut induction, yet it persists for over four hours. This re-localization can serve as a protection from translocation with other chromatin sites.

**Fig. 48.**
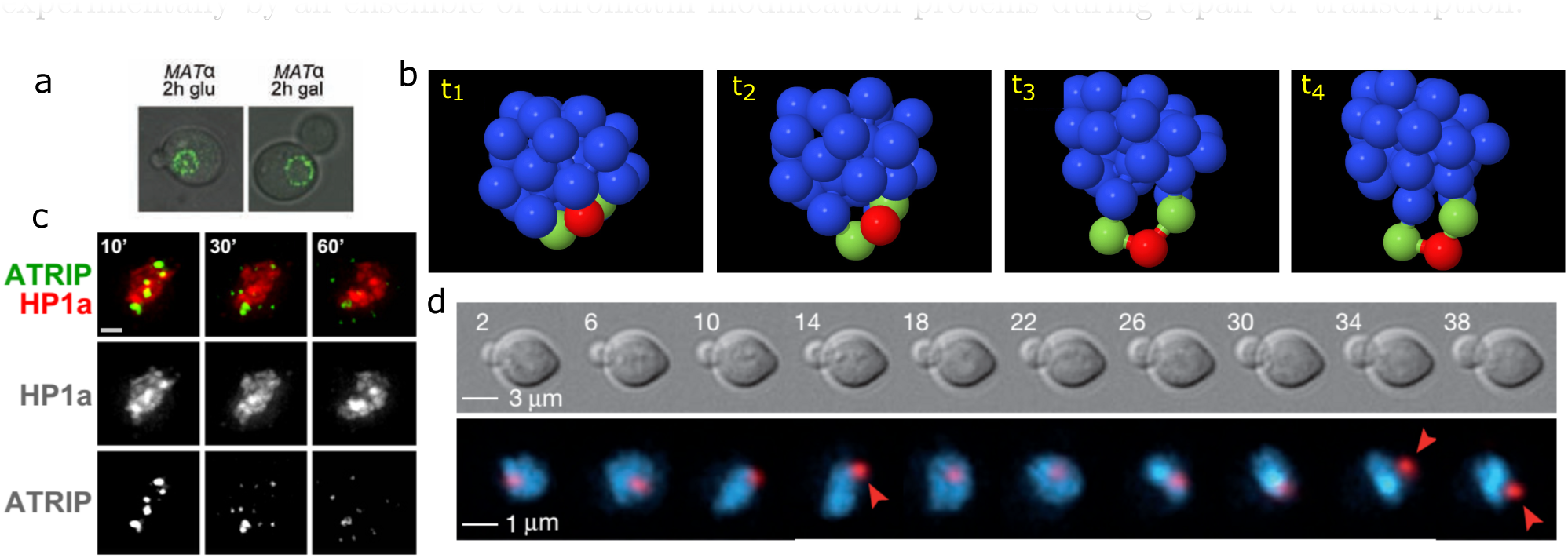
DSB motion can serve to increase repair efficiency and reduce the change for chromosomal translocations. (a) Un-repairable DSB move to the periphery: When a DSB cannot repair itself for a long time, it will be anchored an the nuclear pores. Before break induction (growth in glucose), the locus is randomly distributed in the nucleus. The MAT locus relocate to the periphery between 0.5 and 2 hours after the HO-induced DSB (using galactose) [188]. (b) Upon the induction of a DSB in condense chromatin domain, the DSB is gradually extruded to the periphery of a domain. The process is modeled using a β polymer, where at *t*_1_, the long-range connections for monomers *n* = 16, 17, 18 (in a chain of length *N* = 33) are cut. As a result monomer *n* = 17 is slowly pushed(*t* = *t*_2_ *, t*_3_ *, t*_4_) to the periphery of the polymer globule. (c) DSB move outside of heterochromatin after irradiation of Drosophila nuclei [49]: ATRIP foci (ATRIP is a repair protein which is recruited to the DSB) form within the HP1a (heterochromatin/condense chromatin) domain and relocate outside of this domain. Stills images of cells expressing GFP-ATRIP and mCharry-HP1 (HP1 is a heterochromatin associated protein) are shown, in which DSBs were produced by ionizing radiation (IR). Maximum intensity projections of one cell at 10, 30, and 60 min after IR show the relocalization of ATRIP foci to the periphery and outside of the HP1a domain. (d) A DSB in rDNA locus is moved outside of the nucleolus [276]: Time-lapse microscopy of a TetImRFP-focus (red) next to a DSB marked rDNA repeat in a yeast cell expressing Nop1CFP (blue region). Arrowheads mark the exit of the marked rDNA repeat with the DSB from the nucleolus. Time is indicated in minutes. Selected time points of a representative cell are shown (reproduced from [15]).

The changes in the motion of a DSB after break induction are characterized by an increase in the diffusion coefficient, the anomalous exponent and the con nement radius, con rming physical changes at the chromatin site (Fig. 47b-c-d-f). These changes depend on repair proteins [181, 69, 15] and can be caused by nucleosome repositioning [15]. In some cases, the broken site is extruded from the interior of a dense chromatin territory, and it relocates to the periphery of the local chromatin domain (Fig. 48b). This reconfiguration is due to a reduction of the internal chromatin constraints and result in the extrusion of the DSB. Consequently, it may facilitate sequence-sequence encounters in a condensed chromatin environment. Several observations support this concept: the homologous recombination protein Rad51 is recruited to and forms foci at DSBs. Because heterochromatin may have repetitive sequences, there is a danger that repairing by NHEJ with the domain will lead to chromosome translocation. Thus, these foci move from the interior of the heterochromatin domain to the periphery to complete recombination [49] (Fig. 48c). Similarly, in *S. cerevisiae*, to form a Rad52 focus on broken rDNA, the locus must move out of the nucleolus [276] (Fig. 48d). Rad51 dependent telomere movement was involved in the alternative lengthening of telomeres (ALT) [50]. These modi cations depend on the activity of repair proteins Rad51 and Rad54 which facilitate the extrusion of the DSB. A deletion of either one decreases the anomalous exponent α, allowing a strong increase in local intrinsic chromatin tethering forces [15]. In the absence of Rad51 or Rad54, a DSB becomes more constrained.

In summary, local condensation or compaction of nucleosomes around the break site is due to an increase in local tethering forces. This effect may either be a protective mechanism to avoid translocations or it can influence the microhomology mediated end-joining pathway [62]. If HR isnot functional, the damage is sequestered in a compact nanodomain, possibly to facilitate end-joining. To conclude, in a crowded environment such as the nucleus, self-avoiding interactions have a dominant effect on the encounter time and probability between two chromatin locus. These interactions can be regulated in numerical simulations by changing the monomer density and dynamics and is mediated experimentally by an ensemble of chromatin modi cation proteins during repair or transcription.

## 10 Conclusion and perspective

Polymer models have a signi cant role in clarifying chromatin and nucleus organization. These models are used for extracting biophysical parameters and interpreting single particle trajectories of a chromatin locus. They further serve for reconstructing the chromatin organization from large data sets such as chromosomal capture data. Re ned modeling, coarse-grained stochastic simulations, analysis of the models are now possible for major nuclear events such as double stranded DNA repair and differentiation, but also for the smaller scale function of a cell involving transcription, gene activation or nucleosome translocation. We expect much progress in the next decade in integrating three dimensional super-resolution data with a nanometer precision into polymer models. Indeed, with the improvement of real-time, high resolution imaging, we shall be able to study the manner at which proteins interact with the chromatin.

DNA and chromatin spatial scales span one to two orders of magnitude between bacteria and mammalian cells: while yeast chromosomal organization is fairly well described by classical polymer models based on anchoring, self-avoiding interaction and confinement, metazoans chromosome structure would require more sophisticated models. It remains unclear how to interpret numerical simulations of polymers reconstructed from Hi-C data, because these data represent an average of many realizations, where geometrical distances to the nucleus surface are disregarded. As discussed here, the interpretation of chromosome capture experimental results and their finer details remain unclear. These data represent a frozen conformation of a cell and it is not clear how far they have distorted in-vivo configurations. Developing novel analytical and algorithm methods to reconstruct the chromatin organization should certainly continue clarifying the local chromatin re-organization. Understanding and quantifying the nucleus dynamics requires also novel development in the asymptotic analysis of the polymer models, to extract relations between parameters. These formula will be a fundamental tool to summarize the laws of epigenetic-physics.

### 10.1 Which polymer model to use and when?

Today, the overall space of polymer physics is certainly not completely understood or described. But the increasing need for interpreting data has push to chose a correct (or not) appropriate polymer model for any given situation. We shall now provide some indications about which polymer model should be used to analyse data. Indeed, several statistical quantities can be analytically computed, large and heavy simulations are possible but they all depend on a chose of a polymer model so which one to chose?

In general, there are no consensus about using one polymer model rather than another. For example, the Hi-C data remains difficult to analyse and as discussed in sections 5.4.3, but the compact chromatin is taken into account by the string binder and switch model [19, 20] using random loops and binding sites. Recently, the binder model was extended to impose link between monomers that are highly connected in the Hi-C matrix [246], so that the decay of the encounter probability matrix is recovered. The method allows to prescribed the number of random connectors positioned to reconstruct TAD.

The classical Rouse model was used to interpret single particle trajectories where the anomalous exponent of a given Chromatin locus is α= 0:5. For bacteria, where the exponent is distributed with a mean at 0:4 [128], another possibility is to use polymer model, discussed in section 5.4.2 to generate a polymer model with a prescribed anomalous exponent (α ≦0:5). However the interpretation of exponents α ≥0:5 remains difficult as it certainly involved a mixture of diffusion and active transport. For modeling telomere dynamics in yeast, polymer models were based on spring forces and repulsion force between beads [291] (see also section 7). In all cases, polymer model can be used to obtain statistics about transient events and reconstruct the histogram of arrival time between any two sites of interest (see section 6.4). Another use of polymer model is to study some properties of the search of homologous side during HR (see section 9.3). Classical Rouse or with some modi cations (betapolymer or with repulsion) have been used to study the influence of the nucleus membrane on the the search.

In the context of interpreting single locus trajectories, polymer models serve for predicting the statistics to be expected from data such as the anomalous exponent, radius of con nement, etc. (summarized in section 8). But the statistics are not necessarily polymer model specific. For example, the first and second moments (see formula 221) of the velocity can always be computed, but their interpretation require a specific polymer model.

### 10.2 Possible directions and open questions

We end this section with some open directions where modeling and experimental approaches that should combine:

#### 10.2.1 Extracting forces from multiples tagged loci

What can be extracted from several single locus trajectories. For example how use the properties of the correlation function? the goal would be to extract intrinsic mechanical properties of the chromatin and applied forces. This question is now accessible due to the possibility of tagging at least two loci with a known distance [65, 108]. The theory remains to be worked out and could be based on generalizing eq. 221 to two known locus positions. The cross-correlation function would provide information the rigidity of the interaction. The difficulty is to get long enough trajectories.

#### 10.2.2 Is there a best model to describe Hi-C data and the encounter probability matrix?

What should be the correct model to describe Hi-C data. As discuss in the previous subsection, there are a few models [19, 20, 91, 196]. Possible improvements may by based on using simultaneously random loops and local peaks in the data distribution [246]. It would be also interesting to reconstruct the polymer model by including the geometrical organization of euchromatin and heterochromatin and the nucleus membrane. No such data are yet available, but the application of this approach would be to have a precise location of specific gene in space. We expect that new finding is expected about the organization of gene activated together.

#### 10.2.3 How to reconciliate SPTs and Hi-C statistics?

At the present moment, statistics of SPTs and Hi-C have not been really compared. However, polymer models can be used to compare the decay of the encounter probability of a given locus and the anomalous exponent of that locus. Both quantities are related for example through the *β*- polymer model (see section 5.4.2). However, the deviation from the formula 71 (section 3.10.4) should reveal local multiple chromatin interactions. It should be possible to start with computing the anomalous exponent from numerical simulations.

#### 10.2.4 Proteins remodeling during DNA break repair

Double-stranded DNA break repair involves many repair proteins at different stage of the process (see sections 9.2 and 9.4). For example Rad51 and Rad54 were shown to contribute to the increased movement of DSBs in yeast [68] and Rad51 might facilitate the extrusion of a DSB. Nucleosome reorganization is critical when repairing a break in heterochromatin where nucleosome remodelers invade and pair. In budding yeast, Arp8 is a key component of the INO80 nucleosome remodeling complex and is required for efficient recombination [2] and for increasing the mobility for DSBs and undamaged sites [242, 68]. How shall we model the effect of these proteins during DSB break repair? for example, how the local forces (spring constants) on a Rouse model should be locally modified and to what extend? The polymer model can be modified by considering multiple states for the local monomers near the break. It would be interesting to generate numerical polymer simulations of single locus and estimate various quantities of interested such the diffusion coefficient, interaction forces, the anomalous exponent and so on. A comparison with SPTs should reveal re ned repair steps [15].

#### 10.2.5 Revisiting the classical question of the search process for a promoter site by a transcription factor

With the unprecedent resolution of the chromatin structure and live cell organization, the classical search process scenario of a promoter site by a TF, studied by von Hippel and colleagues (see also [167] for an analytical approach) should be revisited. Indeed, it would be now possible to take into account the relative positions of binding sites to reconstruct the local chromatin environment using the encounter probability matrix of the Hi-C data. Super-resolution data would further refined the exact geometry for stochastic simulations to explore how the chromatin organization can modulate the stochastic search process and in particular, what is really the time spends in three dimension versus the time spend scanning along the DNA.

#### 10.2.6 Computing the probability of a translocation during Non-Homologous End-Joining (NHEJ)

We summarize here one aspect of repair, called HR (setion 9). However, another repair pathway is the NHEJ where two broken end simply relegate. However, when there are several break, some can relegate incorrectly (with another broken arm). This process is call translocation. It would be important to estimate the probability of such translocation event. No much simulations or analysis have been pushed in that direction.

#### 10.2.7 Modeling the random initiation of gene factories

The looping time between any two monomers is now clarified (see section 4), however the mean first time for k (*k* ≥ 2) monomers of a long polymer to enter into a ball of radius *ε* has not yet been computed. This encounter may represent the process of initiating a gene factory regulation in a subdomain. Indeed, it may conceivable that several promoters and repressors interact in small regions to regulate gene expression [209]. The difficulty in the computation of the encounter time is due to additional constraint in the computations of the Gaussian integrals (see relation 99). It would be important to derive analytical expressions as computer simulations of this process should converge slowly due to the singular perturbation: indeed, the encounter time diverges to in nity when the radius *"* tends to zero.

#### 10.2.8 Nucleosome repositioning following DNA damages

Nucleosomes are repositioned following DNA damages, but there is no model for this changes because both the chromatin and the nucleosome are changing their organization. It would be to difficult to account for the molecular detail of the nucleosomes, so a coarse-grained model is necessary. The difficult is chromatin and nucleosomes do not have te same spatial scale, which render the modeling and simulation difficult. Such simulations would help exploring how nucleosome and chromatin are reoganized following break and how the re-organized during repair.

#### 10.2.9 Limitation of statistical analysis for High C-data

Hi-C data have been so far analyzed by estimating statistical parameters such as the decay law for the encounter probability predicted by polymer models [196]. It is however possible to go one step further and design realistic stochastic simulations of chromatin constraint on chromosomal High C-data. We are still facing the limitation of the data that only represent an average configuration over a large cell population, but single cell Hi-C is now possible. Simulations could be used to explore the probability and the conditional mean time for particular sites to meet and also to explore and classify the ensemble of polymer configurations (see also [246]for a first step).

#### 10.2.10 Can statistical analysis of Hi-C data reveal novel features hidden in the genome organization of cancer cells or cells infected by viruses

We speculate that the nucleus organization in cancer cells might reveal clear difference with normal cells, although there no available data generated so far in that direction. Similarly, viral DNA inserted at specific random locations in the chromatin of the host cell might lead to changes in the encounter probability of the Hi-C data. SPT dynamics with polymer reconstruction might lead to a better understanding of the consequences host-virus interactions.

## Acknowledgment

We would like to thank the anonymous reviewers for constructive remarks and A. Taddei, K. Dubrana, A. Seeber, S. Gasser, X. Darzacq, Z. Schuss, L. Atkinson, E. Laue, I. Kupka, J. M. Victor, E. Fabre, L. Mirny, Z. Schuss, Y. Kantor, C. Lavelle, P. Ciccuta, L. Georgetti, O. Shukron, R. Metzler, M. Barbi, E. Heard, A. Bancaud, I. Sokolov, S. Redner, M. Kardar, C. Lavelle, Y. Garini, C. Vaillant, I. Cisse, C. Zimmer and G. Almouznie for stimulating discussions over the past ten years. D. H. thanks the Mathematical Institute, University of Oxford for hospitality, this research was partially supported by a Simons fellowship from the Newton Institute.

